# Additive, synergic and antagonistic interactions between maternal immune activation and peripubertal stress in cocaine addiction-like behaviour, morphofunctional brain parameters and striatal transcriptome

**DOI:** 10.1101/2021.10.11.463760

**Authors:** Roberto Capellán, Javier Orihuel, Alberto Marcos, Marcos Ucha, Mario Moreno-Fernández, Marta Casquero-Veiga, María Luisa Soto-Montenegro, Manuel Desco, Marta Oteo-Vives, Marta Ibáñez-Moragues, Natalia Magro-Calvo, Miguel Ángel Morcillo, Emilio Ambrosio, Alejandro Higuera-Matas

**Author notes:** **Corresponding authors:** Alejandro Higuera-Matas, Department of Psychobiology. School of Psychology. National University for Distance Education (UNED). C/Juan del Rosal 10, Madrid. Spain., Emilio Ambrosio, Department of Psychobiology. School of Psychology. National University for Distance Education (UNED). C/Juan del Rosal 10, Madrid. Spain.

## Abstract

Substance use disorders are more prevalent in schizophrenia, worsening its course and prognosis. Here, we used a double-hit rat model, combining maternal immune activation (MIA) and peripubertal stress (PUS), to study cocaine addiction and the underlying neurobehavioural alterations. We injected lipopolysaccharide or saline on gestational days 15 and 16 to pregnant rats. Their male offspring were then subjected to 5 episodes of unpredictable stress every other day during adolescence (from postnatal day 28 to 38). When rats reached adulthood, we studied cocaine addiction-like behaviour, impulsivity, conditioning processes and several aspects of brain structure and function by MRI, PET and RNAseq. MIA facilitated the acquisition of cocaine self-administration while PUS reduced cocaine intake, an effect that was reversed by MIA. MIA increased motivation for cocaine and reversed the effects of PUS during extended access. Incubation of seeking was unaffected. Neither hit alone nor their combination impacted Pavlovian or instrumental conditioning or impulsiveness. At the brain level, PUS reduced hippocampal volume and hyperactivated the dorsal subiculum. MIA+PUS altered the structure and function of the dorsal striatum increasing its volume and interfering with glutamatergic dynamics. MIA did not affect the gene expression of the nucleus accumbens but, when combined with PUS, modulated specific genes that could account for the restored cocaine intake. PUS had a profound effect on the dorsal striatal transcriptome however, this was obliterated when PUS occurred in animals with MIA. These results describe a complex interplay between MIA and stress on neurodevelopment and in the susceptibility to develop cocaine addiction.

## INTRODUCTION

Despite impressive advances in our understanding of the aetiology of autism spectrum disorders (ASDs) and schizophrenia, the complex interplay between the factors causing these conditions is far from being fully understood. A preponderant view in the field is that ASDs and schizophrenia have a neurodevelopmental origin (1– 3) however, the variables contributing to the progression and prognosis in both conditions still need to be thoroughly investigated. An important aspect to be considered is the presence of substance-related disorders among people suffering from schizophrenia or ASDs. Indeed, substance use in people with schizophrenia is associated with increased morbidity and mortality, worsening the overall course of the disease, as suggested by increased rates of hospitalization, lower treatment adherence, increased suicide rates, etc. (4). This is especially worrying in the light of the fact that substance-use disorders are up to five times more prevalent than in the general population, affecting almost 50% of patients (this figure rises to 90% for nicotine use disorder) (5). In the specific case of cocaine, in a European sample, the risk of developing cocaine abuse was 7 times higher in those with schizophrenia than in the general population (6). Concerning substance use-related problems in ASDs, they have traditionally been considered rare probably because the core features of the condition seemed to lower the risk of using drugs of abuse; however, recent data suggest that the prevalence of substance use among those with ASDs may be slightly higher than in the general population, even after a comorbid diagnosis of attention-deficit hyperactivity disorder was considered (7).

In considering the neurodevelopmental origin of schizophrenia and ASDs, maternal immune activation (MIA) models have arisen as a key tool to experimentally investigate the comorbidity of neurodevelopmental disorders and addiction. MIA models recapitulate several aspects of these diseases (8,9) and have been used to determine if behaviours reminiscent of specific aspects of addiction are increased in animals exposed to MIA (see Menne and Chesworth 2020 and Ng et al. 2013 for excellent reviews). In the case of cocaine, MIA induced by the synthetic viral RNA analogue polyinosinic:polycytidylic acid (poly I:C) increased cocaine-induced conditioned place preference (but decreased cocaine-induced locomotor activity) (11) and also potentiated cocaine cross-sensitization after an amphetamine challenge (12). However, when we tested if MIA induced by the bacterial endotoxin lipopolysaccharide (LPS) would potentiate cocaine self-administration, we found no evidence of such effect even in the presence of sensorimotor gating and cognitive deficts (13) or sensorimotor gating and immune alterations (14). However, despite their translational validity, MIA models alone may not fully capture the complexity of the interactions between causative factors and, more recently, these models are incorporating a second hit during adolescence to increase their validity (15,16).

In the present work, we aimed to expand our previous findings combing LPS-induced MIA and PUS and perform an extensive characterization of the addiction-like behaviour of these animals (acquisition of cocaine self-administration, motivation for cocaine, compulsivity, loss of control over drug intake and incubation of seeking) (17,18) and extensively study the neurodevelopmental alterations present in this model by taking advantage of preclinical neuroimaging (structural magnetic resonance imaging -MRI-, diffusion tensor imaging -DTI-, magnetic resonance spectroscopy - MRS-positron emission tomography -PET-) and next-generation sequencing (RNAseq).

## MATERIALS AND METHODS

A complete description of the materials and methods followed can be found in the Supplementary Methods provided in the Supplementary Information.

### Experimental animals

Experiments were performed on the male offspring of 14-week-old male and 12-week old female Sprague-Dawley rats obtained from Charles River (France) following the EU Directive 2010/63/EU and approved by the Bioethics Committee of UNED and the Autonomous Community of Madrid (PROEX 078/18).

### Maternal immune activation

LPS (from *Escherichia coli* 0111: B4 [Sigma-Aldrich]) dissolved in 0.9% NaCl was intraperitoneally injected to pregnant rats at a dose of 100 mg/kg/ml on gestational days (GD) 15 and 16.

### Peripubertal unpredictable stress

Between PNDs 28 and 38, we exposed the male offspring to peripubertal unpredictable stress (PUS) consisting of 1) stress by agitation. 2) Stress by immobilization; 3) water deprivation; 4) forced swimming and 5) constant changes of the home cage, each one applied every other day in a counterbalanced order. Non-stressed controls received handling by the same researcher and on the same days as the stressed subjects.

### Experimental Design

All the studies that will be described hereafter were carried out in independent batches of animals, except for the PPI test, which was performed in all of them. Figure S1 shows an outline of the experimental design employed.

### Experimental Procedures

#### Prepulse inhibition of the acoustic startle response (PPI)

At PND70-73, PPI of the acoustic startle was measured in a non-restrictive Plexiglas cage (28×15×17 cm). The session was composed of ten 120 dB pulse-alone trials, five null trials with no stimulus and twenty pulses preceded by a prepulse of 69- or 77-dB intensity (4 or 12 dB above the background noise, respectively) with an interval of 30 or 120 ms (13).

#### Cocaine self-administration

At PND80 animals underwent surgery to be implanted with a jugular vein catheter that allowed intravenous cocaine self-administration. On PND90, cocaine self-administration (0.5 mg/kg i.v.) program began with six different phases: (1) acquisition (12 sessions 2-hour daily sessions under an FR1 schedule); (2) motivation test (6 sessions 2-hour daily sessions under a progressive ratio program); (3) stabilization (3 sessions of identical conditions as those used in the acquisition phase); (4) compulsive seeking test (a single session 1-hour session under an FR3 schedule where the animals randomly received an infusion or a mild plant shock of 0.5 mA for 0.5 seconds); (5) extended access (10 6-hour daily sessions under an FR1 schedule). At the end of this phase, a forced abstinence period was imposed, to evaluate cue-induced relapse, leading to the sixth phase; (6) drug-seeking test (a 2-hour session with no drug on days withdrawal days 1, 30, 60 and 100).

#### Pavlovian and instrumental conditioning

The Pavlovian learning protocol consisted of eight daily sessions where a CS predicted the delivery of a food pellet and another CS the absensce of the reward. We measured the head entries to the food magazine after the presentation of each type of stimulus and calculated a ratio between them as a learning index. The instrumental learning protocol was performed after the Pavlovian training and consisted of seven daily sessions where presses of an active lever led to the presentation of a food reward under different reinforcement schedules. An inactive lever was present but had no effect on food delivery.

#### Two-choice serial reaction time task (2-CSRTT)

We used an adaptation of the widely used 5-CSRTT (19) to measure sustained visual attention and waiting impulsivity. Rats had to withhold their responses on a specific lever until the appropiate stimulus appeared marking the onset of the interval when responses on that marked lever were rewarded.

### Neuroimaging studies

#### Magnetic resonance imaging (MRI) and diffusion tensor imaging (DTI) and *in vivo* proton magnetic resonance spectroscopy (^1^H-MRS)

At PND90, MRI and DTI studies were performed at the Biomedical Research Institute “Alberto Sols” (CSIC-UAM, Madrid, Spain) using a 7.0 Tesla Pharmascan MRI scanner (Bruker, Germany) to acquire T2-weighted (T2-W) spin-echo anatomical images. Diffusion-weighted images were acquired with a spin-echo single-shot echo-planar imaging (EPI) pulse. Fractional anisotropy (FA), mean diffusivity (MD), trace, eigenvalues and eigenvector maps were calculated with a homemade software application written in Matlab (R2007a). Values were extracted from maps using regions of interest (ROIs) with Image J software.

Immediately after conducting the MRI and DTI analysis, an *in vivo* ^1^H MRS study was also performed in the Biomedical Research Institute “Alberto Sols” (CSIC-UAM, Madrid, Spain). Two brain regions were selected for this study: cortex and striatum. A Point-REsolved Spatially Spectroscopy (PRESS) was used, combined with a variable power radiofrequency (VAPOR) water suppression.

#### Positron emission tomography/computed tomography (PET-CT)

Once PND90 was reached, PET-CT studies were performed at the Radioisotopes for Biomedicine research group of the Center for Energy, Environmental and Technological Research (CIEMAT) in Madrid, Spain, using a small-animal Argus PET/CT, SEDECAL, Madrid, Spain). PET studies (energy window 400–700 KeV and 45 minutes static acquisition) and CT (voltage 45 kV, current 150 μA, 8 shots, 360 projections and standard resolution) were performed 30 minutes after administration of 34.7±3.02 MBq of [^18^F]-2-fluoro-2-deoxy-d-glucose (^18^F-FDG) via the tail vein in fasting rats anesthetized by inhalation of 2–2.5% isoflurane in 100% oxygen. PET-CT images reconstruction was accomplished using a 2-dimension ordered subset expectation maximization (2D-OSEM) algorithm (16 subsets and 3 iterations), with random and scatter correction. Statistical Parametric Mapping (SPM) software was used for voxel-based analysis (http://www.fil.ion.ucl.ac.uk/spm/software/spm12/).

#### RNAseq

Once PND90 was reached, animals were sacrificed and tissue from the nucleus accumbens (NAcc) and dorsolateral striatum dissected out. RNA extraction was performed using the RNeasy Mini Kit (Qiagen). RNA-Seq analysis was carried out in the Genomics Unit of the Madrid Science Park. Libraries were prepared according to the “NEBNext Ultra Directional RNA Library Prep kit for Illumina” (New England Biolabs) instructions and sequenced using a “NextSeq™ 500 High Output Kit”, in a 1×75 single read sequencing run on a NextSeq500 sequencer. Differential expression analysis was carried out using the CUFFDIFF tool. We then used Metascape (https://metascape.org/gp/index.html#/main/step1) to analyze the enrichment in specific gene ontologies for each comparison.

### Statistical Analysis

We used 2 way ANOVAs or non-parametric statistic when necessary.

## RESULTS

The main results obtained in all the studies are summarized in Table 1.

**TABLE 1:** Summary of the main experimental findings.

| SUMMARY TABLE |  |  | LPS | PUS | LPS & PUS interaction |
| --- | --- | --- | --- | --- | --- |
| Behaviour | PPI |  | = (↓ trend) | = | = |
|  | Cocaine self-administration | Acquisition | = | ↓ | = |
|  |  | Motivation | ↑ | = | = |
|  |  | Compulsivity | = | = | = |
|  |  | Extended access | = | = | ↑ PUS-exposed animals by LPS |
|  |  | Cue-induced seeking | = | = | = |
|  | Food reinforcement | Pavlovian conditioning | = | = | = |
|  |  | Instrumental conditioning | = | = | = |
|  | Motor impulsivity |  | = | = | = |
| Neuroimaging | Volumetry |  | ↓ Whole brain | ↓ Right hippocampus<br>↑ Left cortex<br>↑ Dorsal striatum | = |
|  | DTI | MD | ↑ Hippocampus | = | = |
|  |  | FA | = | = | = |
|  | PET-CT |  | = | ↑ Dorsal cortex<br>↑ Dorsal hippocampus<br>↓ Brainstem | = |
| Neurochemistry | *In vivo* <sup>1</sup>H MRS | Choline compounds | = | = | ↓ Cortex by PUS; reversed by LPS |
|  |  | *N*-acetylaspartate compounds | = | = | ↓ Striatum<br>LPS + PUS synergy |
| Gene expression | RNA-Seq |  | ↑↓ Functionality and development<br>• Nucleus accumbens<br>• Dorsolateral striatum | ↑↓ Functionality and development<br>• Nucleus accumbens<br>• Dorsolateral striatum (large effect) | ↑↓ Functionality and development<br>• Nucleus accumbens<br>• Dorsolateral striatum |

### Prepulse inhibition of the acoustic startle response

We only found a trend in LPS-exposed animals to show impaired PPI in the 12 dB 30 ms condition (p=0.056) suggesting a latent deficit, but, contrary to our expectations, no interaction was observed between MIA and PUS (see Table S1). Other than this, no significant global effects of MIA, PUS or their interaction were observed in any of the experimental conditions used in the PPI test (Figure S2).

### Cocaine-related behaviours

#### Acquisition patterns and initial cocaine self-administration

Even in the absence of overt sensorimotor deficits, we asked whether these animals that show a latent vulnerability would be at increased risk of developing a cocaine use disorder. We first performed a categorical analysis of the rats, according to whether they successfully acquired cocaine self-administration. (Figure 1, A2). Although there were no significant differences between groups in the overall analysis (X^2^_6_=9.553; p=0.1450), we observed that the group of LPS-exposed animals (without stress) had a strong trend (X^2^_2_=5.887; p=0.053) to have a higher proportion of rats displaying robust cocaine self-administration (greater “Normal acquisition” percentage). Within the stressed rats (non exposed to LPS), no significant differences were obtained (X^2^_6_=3.625; p=0.163), suggesting that stressful experiences during peripuberty do not affect the percentage of individuals acquiring cocaine self-administration.

**Figure 1:**
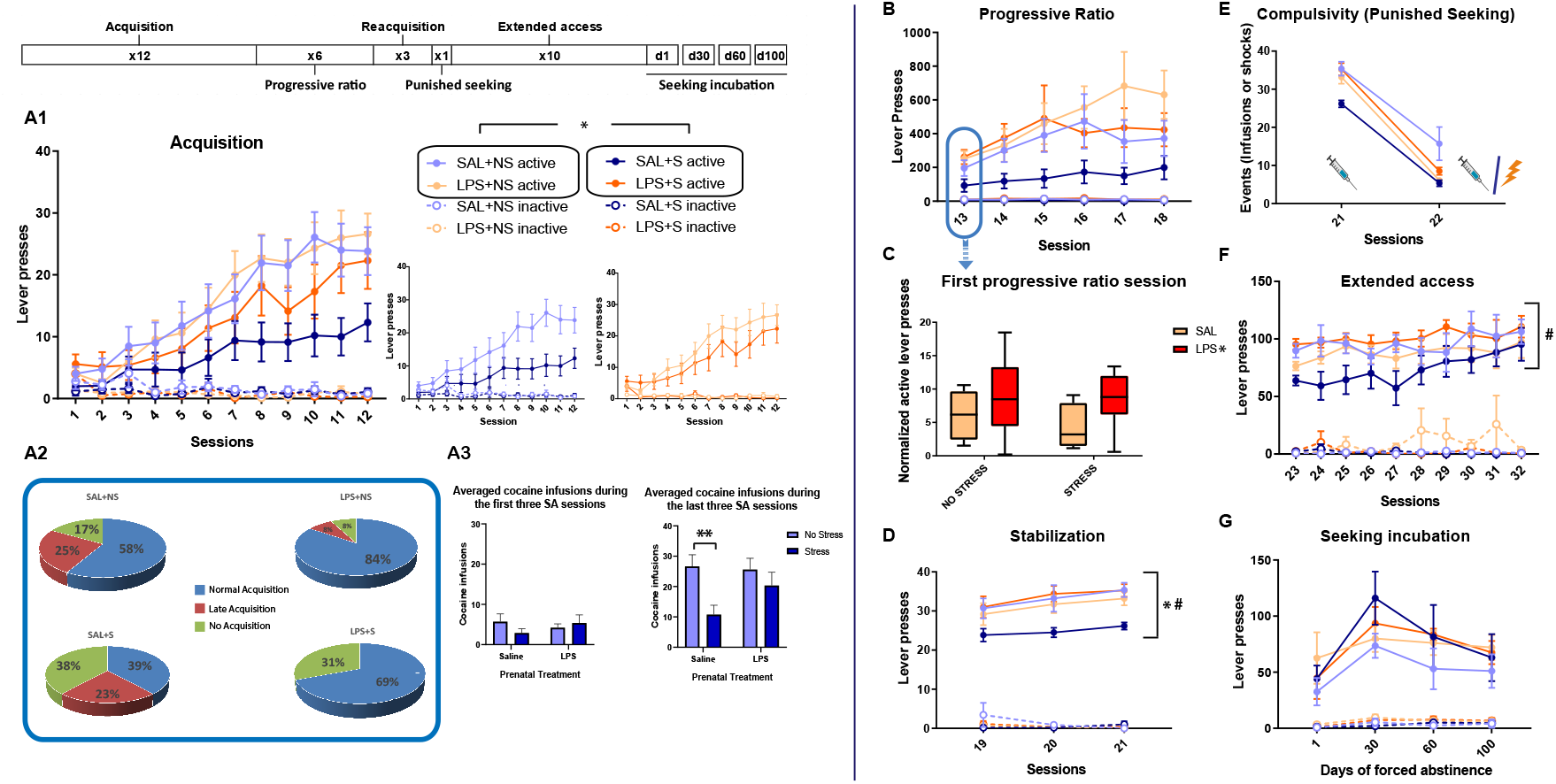
Cocaine self-administration program. (A1) Active and inactive lever presses during the acquisition phase (SAL+NS: n=13; SAL+S: n=13; LPS+NS: n=13; LPS+S: n=15). All groups showed an effective discrimination between active and inactive lever presses, as shown by the significant Lever factor (F_1,47_=72.233; p=0.000; η^2^p=0.606) and the Lever * Sessions factors interaction (F_1,47_=41.403; p=0.000; η^2^p=0.468). Importantly, the Lever * prenatal immune activation or the Lever * PUS interactions were not significant. In addition, no significant effects were found on inactive lever presses during this phase. (A2) The results of the categorical analysis showing the percentage of animals with regular, delayed or absent cocaine self-administration acquisition. To carry out this qualitative analysis of the acquisition phase, animals were categorized as to show: “Regular acquisition” (animals that showed cocaine self-administration behaviour i.e. more than 5 infusions over the last three days of acquisition), “Late acquisition” (animals that took more than 12 sessions to present a stable behaviour of cocaine self-administration) and “No-acquisition” (animals that by the end of the experiment never showed a cocaine self-administration behaviour) (A3) Number of cocaine infusions over the first or last three self-administration sessions. (B) Active and inactive lever presses during the progressive ratio phase (SAL+NS: n=9; SAL+S: n=8; LPS+NS: n=12; LPS+S: n=9). Discrimination between active and inactive lever presses was significant in all groups, as shown by the Lever factor (F_1,34_=40.674; p=0.000; η^2^p=0.610) and Lever * Sessions factors interaction (F_1,34_=5.276; p=0.002; η^2^p=0.169). (C) Normalized active lever presses in the first progressive ratio session (active lever presses in in the first progressive ratio session / active lever presses in the last acquisition phase session). (D) Active and inactive lever presses during the stabilization phase (SAL+NS: n=9; SAL+S: n=6; LPS+NS: n=11; LPS+S: n=8). Discrimination between active and inactive lever presses was significant in all groups, as shown by the Lever factor (F_1,29_=853.870; p=0.000; η^2^p=0.967) and Lever * Sessions factors interaction (F_1,29_=7.935; p=0.002; η^2^p=0.215). (E) Reduction in drug-seeking due to foot shock represented as a decrease in the events (infusions or shocks) registered during the compulsive-seeking session (SAL+NS: n=7; SAL+S: n=6; LPS+NS: n=11; LPS+S: n=8) regarding those registered in the last stabilization session; (F) active and inactive lever presses during the extended access phase (SAL+NS: n=8; SAL+S: n=6; LPS+NS: n=11; LPS+S: n=8). Discrimination between active and inactive lever presses was significant in all groups by the Lever factor (F_1,23_=244.412; p=0.000; η^2^p=0.914). (G) Active and inactive lever presses during the seeking incubation phase (SAL+NS: n=5; SAL+S: n=5; LPS+NS: n=10; LPS+S: n=8). Discrimination between active and inactive lever presses was significant in all groups, as shown by the Lever factor (F_1,24_=81.650; p=0.000; η^2^p=0.773) and Lever * Sessions factors interaction (F_1,24_=9.199; p=0.001; η^2^p=0.277). * denotes a p<0.05 while ** denotes a p<0.01 difference compared to the stressed group among saline-exposed rats; # denotates a p<0.05 difference compared to the respective stressed group among LPS-exposed rats.

All rats progressively increased the number of lever presses during training (significant effects of the Sessions factor (F_1,47_=43.071; p=0.0001; η^2^p=0.478)); however, as seen in Figure 1 A1, PUS significantly reduced cocaine self-administration (F_1,47_=4.409; p=0.041; η^2^p=0.086), an effect that was more evident at the end of the training, as revealed by the significant Sessions * PUS interaction (F_1,47_=3.718; p=0.007; η^2^p=0.073). Interestingly, the effect of PUS was restricted to SAL-exposed animals while LPS-rats seemed to be resistant to the effects of PUS on cocaine self-administration acquisition (Figure 1, A3).

Other critical parameters that indicate a tendency to develop compulsive cocaine intake, such as the number of lever presses during the time out period, the mean latency to the first lever press or the mean inter-response time, were not affected (data not shown).

For the analysis of the motivation to consume cocaine, we used a progressive ratio schedule. We first focused on the performance during the first session as it is devoided of effects of the progressive behavioural adaptation to the schedule. Active lever presses values in the first session were normalized against those of the last acquisition session to rule out a possible influence of the consumption pattern exhibited in the previous phase. Using this index, we observed that LPS-exposed animals showed greater motivation for cocaine consumption (F_1,34_=6.628; p=0.015; η^2^p= 0.163) in the first session of the progressive ratio phase (day 13) (Figure 1, B and C). A significant effect of the Sessions factor was found in the active lever presses (F_1,34_=5.206; p=0.002; η^2^p=0.167) (Figure 1, B) and in the breaking points (F_1,34_=5.096; p=0.002; η^2^p=0.164) (Figure S3) reached during the progressive ratio phase, however, no significant differences were obtained between groups. Inactive lever presses were not significantly altered during this phase (Figure. 1, B).

When animals were returned to FR1, we observed a progressive increase in active lever presses (F_1,29_=7.432; p=0.001; η^2^p=0.204) and also significant differences between the groups (as revealed by the interaction between prenatal immune activation and PUS (F_1,29_=7.316; p=0.011; η^2^p=0.201). Prenatal LPS treatment reversed the reduction in cocaine self-administration induced by PUS (indeed, LPS-stressed rats self-administered more cocaine than SAL-stressed rats (F_1,29_=6.21; p=0.018) and SAL-stressed rats displayed lower intake than SAL-non-stressed rats F_1,29_=7.54; p=0.010) (Figure 1, D). No significant effects were found in inactive lever presses during this phase (Figure 1, D). Compulsive cocaine-seeking was not affected by MIA or PUS. Indeed, all rats equally reduced their drug-seeking when the random electric shocks were introduced (significant effect of the Sessions factor (F_1,28_=388.172; p=0.000; η^2^p=0.933) when comparing the drug infusions earned during the last stabilization phase session the number of infusions/shocks in the compulsive seeking test session (Figure 1, E). During the extended access phase, we observed significant effects of the Sessions factor (F_1,23_=4.945; p=0.004; η^2^p=0.177) and the interaction between MIA and PUS (F_1,23_=4.893; p=0.037; η^2^p=0.175). Prenatal LPS-treatment normalised cocaine self-administration among stressed animals (F_1,23_=4.889; p=0.035). No significant effects were found on inactive lever presses during this phase (Figure 1, F). When we analysed drug-seeking after different withdrawal times, we detected a significant effect of the Sessions factor (F_1,24_=10.989; p<0.001 η^2^p=0.314), suggestive of the incubation of seeking phenomenon. However, no significant differences were obtained between groups (Figure 1, G).

These results cannot be attributed to alterations in Pavlovian or instrumental learning processes (see supplementary results and Figure S4 A and B). Moreover, there were no changes in impulsive behaviour that may have contributed to generate part of this cocaine self-administration phenotype (Figure S5).

### Neuroimaging studies

#### MRI-assisted volumetry

Having characterized cocaine addiction-like behaviour in this model, we decided to use MRI to search for brain alterations that could be underlying the behavioural patterns obtained. Whole-brain volume was reduced as a consequence of prenatal immune activation (F_1,27_=6.520; p=0.017; η^2^p=0.195) (Figure 2, A). On the other hand, PUS increased both the left cortical volume (F_1,27_=5.709; p=0.024; η^2^p=0.175) (Figure 2, B) and the dorsostriatal volume bilaterally (Right: F_1,27_=5.289; p=0.029; η^2^p=0.164 / Left: F_1,27_=6.856; p=0.014; η^2^p=0.203) (Figure 2, D), but decreased the right hippocampus in the LPS animals (F_1,27_=4.417; p=0.045; η^2^p=0.141) (Figure 2, C). A MIA * PUS interaction was evident in the right cortical volume (F_1,27_=4.484; p=0.044; η^2^p=0.025) (Figure 2, B), however, no significant differences were observed in subsequent simple effects analysis. This interaction was also observed in the volume of the left dorsal striatum (F_1,27_=6.105; p=0.020; η^2^p=0.184), whereby LPS-exposed stressed rats showed a significant increase in volume compared to non-stressed (F_1,27_=12.519; p=0.001) and saline-exposed (F_1,27_=5.018; p=0.034) controls (Figure 2, D).

**Figure 2:**
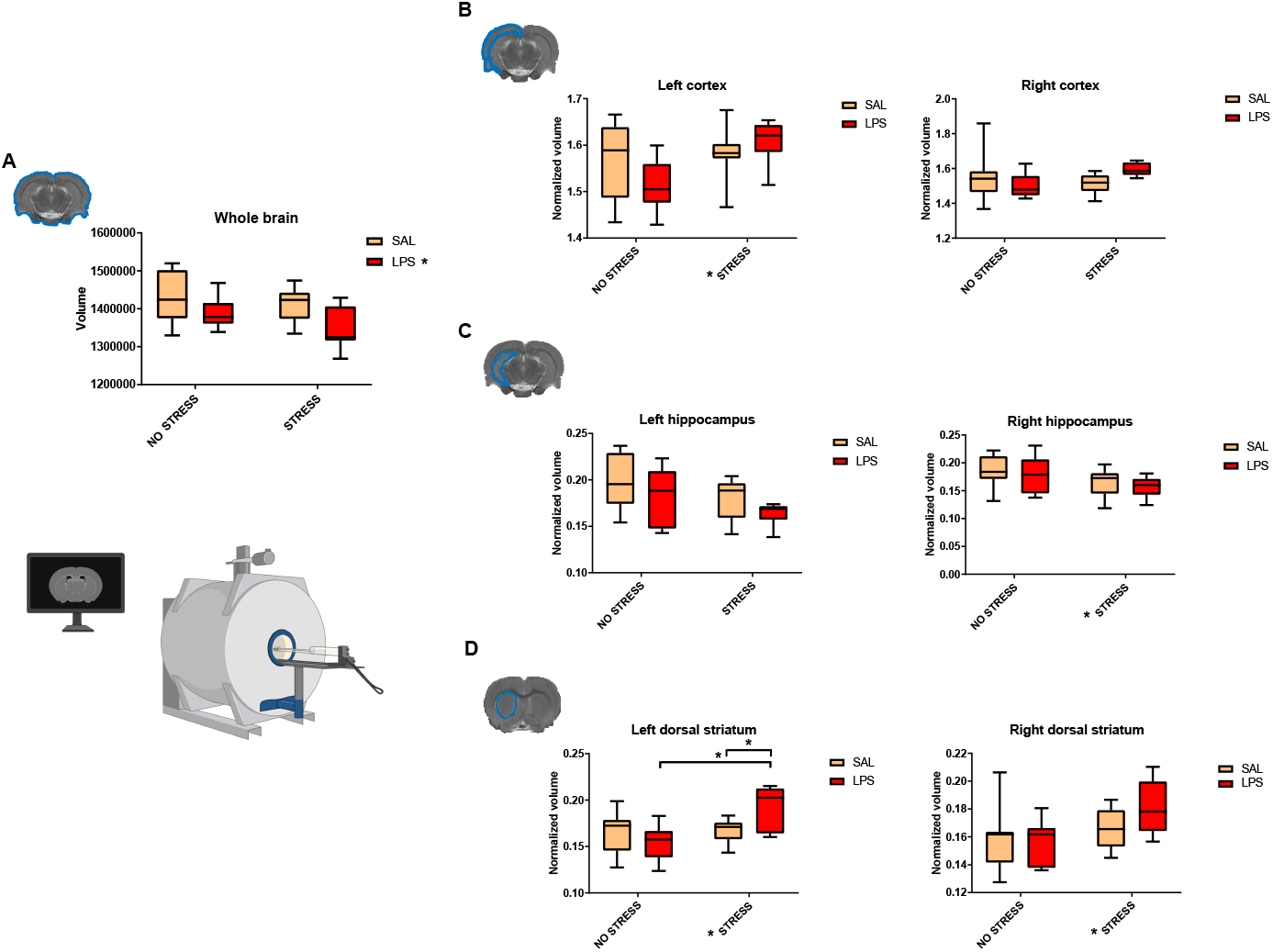
Brain Regional Volumetry. The figure shows the volume of the whole-brain (A), left and right cortex (B), left and right hippocampus (C), left and right dorsal striatum (D). The volumetric quantification of each brain structure was normalized by whole brain volume in the different MRI sections. (SAL+NS: n=8; SAL+S: n=8; LPS+NS: n=8; LPS+S: n=7). * in a boxplot denotes a p<0.05 difference compared to the corresponding saline group. * in the legend denotes a significant main effect (p<0.05) of the maternal immune activation factor.

No significant effects of prenatal immune activation, PUS or their interaction were found in cerebellar, amygdalar and NAcc volumes (Figure S6, A-C), nor in the fourth ventricle, third ventricle, lateral ventricles, cerebral aqueduct and total ventricular volumes (Figure S7, A-F).

#### DTI

MD was significantly increased in the right (U=60, p=0.0177) and left (U=58, p=0.0143) hippocampi of LPS-exposed animals (Figure 3, C). Regarding this parameter, no significant differences were observed between groups in the cortex or the dorsal striatum (Figure 3, A and B). No alterations in white matter integrity were observed in our analysis. Indeed, there were no changes in fractional anisotropy in any of the structures mentioned (Figure S8, A-C) or in the main cerebral tracts (corpus callosum, internal capsule and anterior commissure) (Figure S9, A-C), as a consequence of MIA, PUS or their interaction.

**Figure 3:**
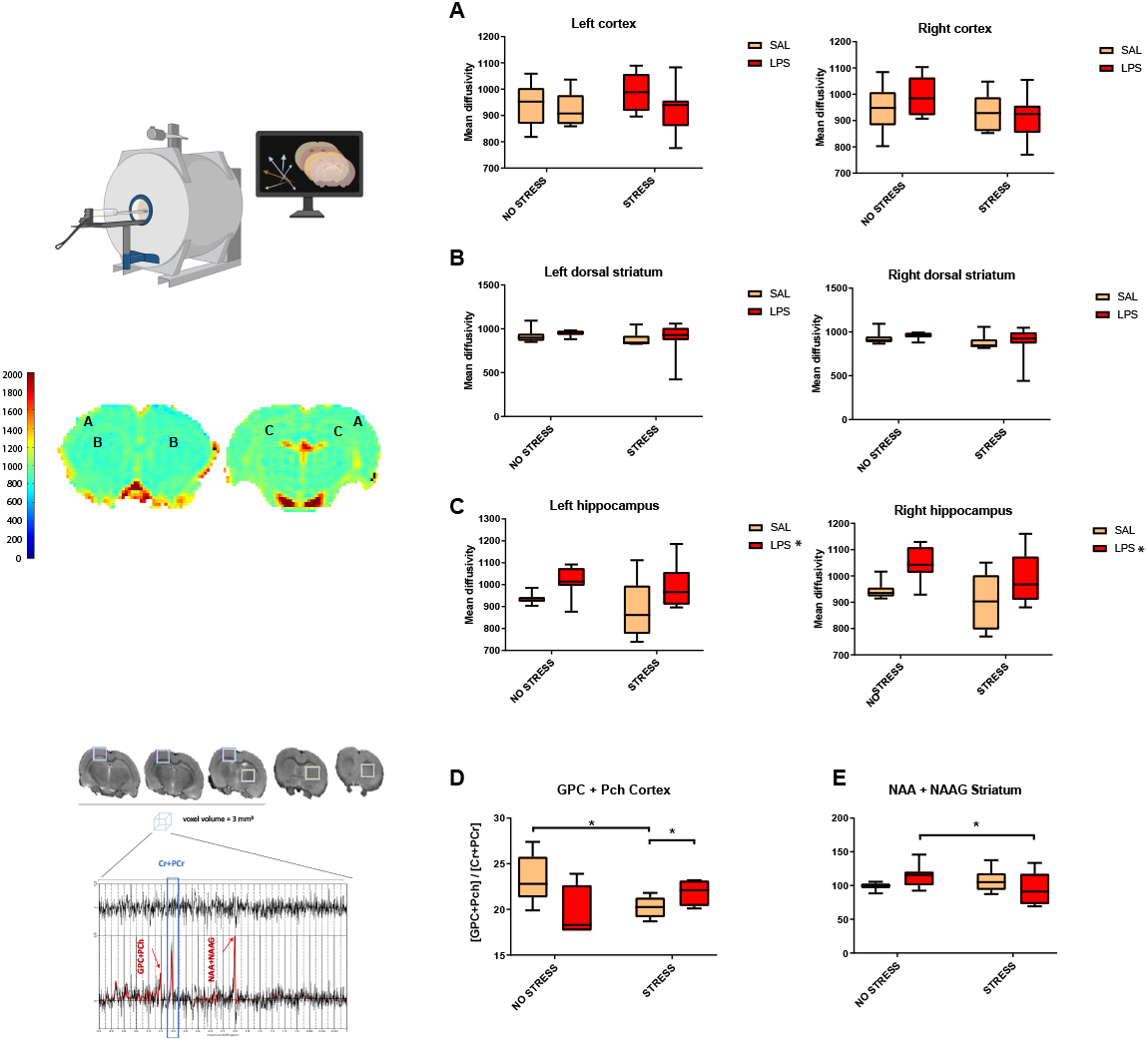
DTI: Mean diffusivity measurements in specific brain areas and ^1^H-MRS: normalized [GPC+PCh] levels in the cortex and [NAA+NAAG] levels in the striatum. The figure shows mean diffusivity values in the cortex (A), dorsal striatum (B) and hippocampus (C), in both hemispheres. (SAL+NS: n=8; SAL+S: n=8; LPS+NS: n=8; LPS+S: n=8). * boxplot denotes a p<0.05 difference compared to the corresponding saline group. The figure shows (A) normalized [GPC + PCh] levels in the cortex (SAL+NS: n=6; SAL+S: n=6; LPS+NS: n=4; LPS+S: n=4) and (B) normalized [NAA+NAAG] levels in the striatum (SAL+NS: n=7; SAL+S: n=8; LPS+NS: n=8; LPS+S: n=7). * indicates a significant difference (p<0.05) between groups.

#### ^1^H-MRS

We obtained a MIA * PUS interaction in the levels of choline compounds (glycerophosphorylcholine + phosphorylcholine -GPC + PCh-peak) in the cortex (F_1,16=_6.940, p=0.018, η^2^p=0.303). While PUS reduced the relative amount of these metabolites (F_1,16_=6.246, p=0.024), the combination with MIA reversed this effect and slightly increased the levels of this metabolite among stressed animals (F_1,16_=6.481, p=0.022) (Figure 3, D).

With regard to the N-Acetylaspartate + N-Acetylaspartylglutamic acid (NAA+NAAG) peak levels, we also obtained a MIA* PUS in the striatum (F_1,26_=4.776, p=0.038, η^2^p=0.155) whereby PUS reduced the relative amount of these metabolites only in LPS-exposed animals (F_1,26_=4.495, p=0.044) (Figure 3, E).

#### PET

PUS alone caused basal hypermetabolism of the cortex and hippocampus at the dorsal level, while brainstem regions showed basal hypometabolism (Figure 4, A). When combining this factor with prenatal immune activation, the hypermetabolism shifted to the cortex and the brainstem hypometabolism disappeared (Figure 4, B), suggesting that prenatal LPS exposure modulates the effects of PUS on brain metabolism. Furthermore, cortical hypermetabolism by such a combination was notably stronger as compared to the group that was not exposed to any hit (Figure 4, C).

**Figure 4:**
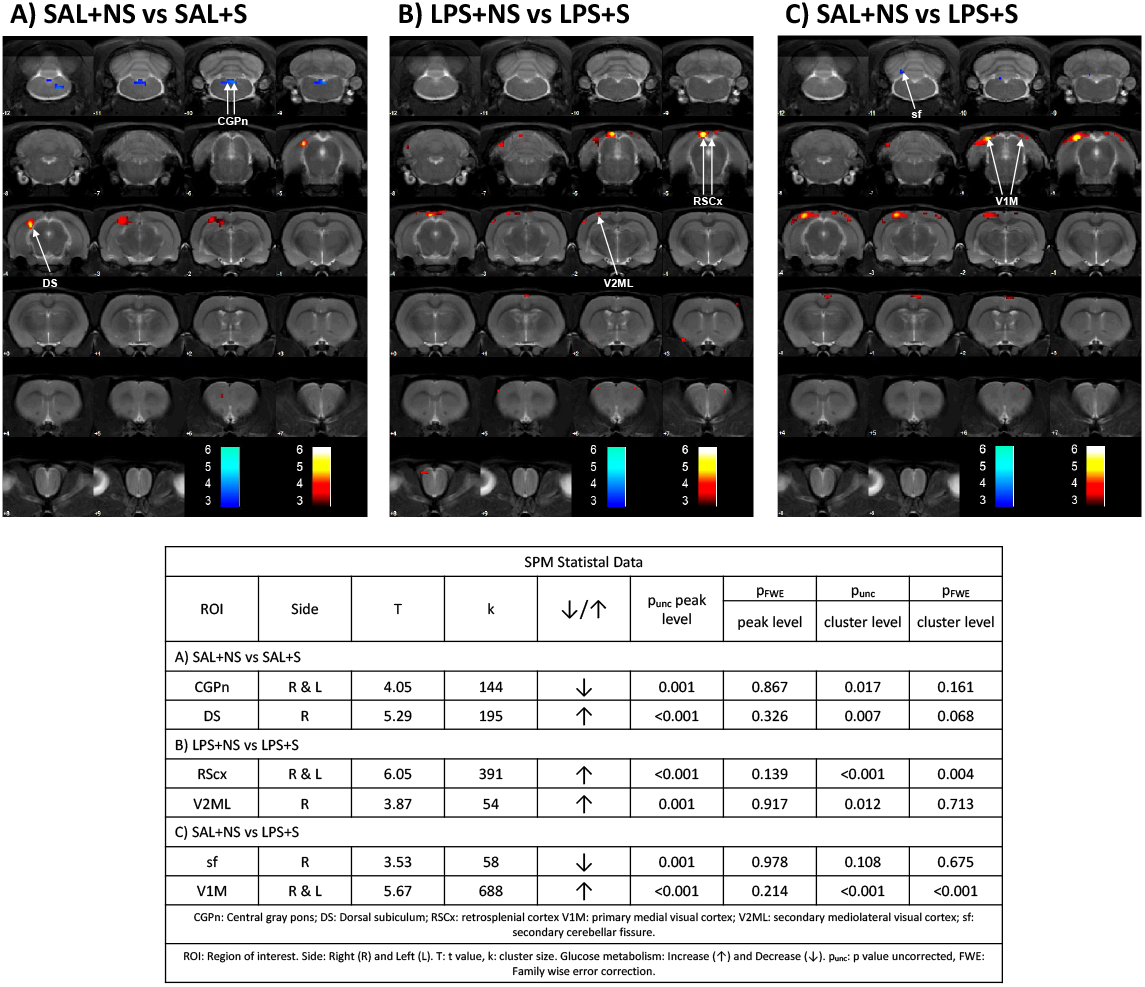
SPM analysis of [^18^F]-FDG for the metabolic activity. Colored **PET images: T-maps overlaid on an MRI template** indicate reduced (blue) and increased (red) FDG uptake. The color bars represent **p level of the voxel-wise comparisons of the brain metabolic activity**. The figure shows the effects found in the SAL+S compared to SAL-NS rats (A), the effects found in the LPS+S compared to LPS+NS rats (B) and effects found in LPS+S rats compared to SAL+NS animals(C). The statistical significance threshold between groups was set at p<0.01 (uncorrected) and k>50 voxels. CGPn: Central gray pons; DS: Dorsal subiculum; RSCx: retrosplenial cortex V1M: primary medial visual cortex; V2ML: secondary mediolateral visual cortex; sf: secondary cerebellar fissure. (SAL+NS: n=5; SAL+S: n=5; LPS+NS: n=4; LPS+S: n=5).

No noticeable effects of prenatal immune activation were observed on brain metabolic activity, neither in the absence (Figure S10, A) nor in presence of PUS (Figure S10, B).

### RNAseq Analysis

#### Nucleus Accumbens

Illumina’s RNA sequencing reports showed that MIA induced the differential expression of 60 genes in non-stressed animals and this amount was reduced to 53 among stressed rats. Surprisingly, only three genes were shared between the two comparisons (*Aurkb, Car3* and *Tnnt1*, down-regulated in all cases), suggesting that PUS changes the gene expression programmes induced by MIA in the NAcc. PUS did not affect gene expression in control rats but was associated with the differential expression of 63 genes among animals subjected to MIA (Figure S13). There were 36 differential expressed genes (DEGs) that responded to one hit (MIA or PUS) on specific levels of the other hit: 35 shared DEGs between the SAL+S vs LPS+S comparison and the LPS+NS vs LPS+S comparison, and 1 DEG shared between the SAL+NS vs LPS+NS comparison and the LPS+NS and LPS+S comparison (Figure S13). Concerning specific DEGs and gene ontologies, MIA increased the expression of the *Sync* gene (coding for Syncoilin) (Figure 5, A) and reduced the expression of several genes (*Car3* -carbon anhydrase-, *Fcrl2* - Fc Receptor Like 2- and *Folr1* -Folate Receptor Alpha-being with the ones with highest fold-change). The analysis of the gene ontologies affected showed that MIA reduced the expression of different genes that clustered in separate groups of ontologies. On the one hand, it affected genes involved in the development of the nervous system such as those regulating axoneme assembly, pattern specification or left/right symmetry. On the other, it affected several gene ontologies related to behavioural regulation and central nervous system function, such as ‘locomotory behaviour’, ‘movement’ and ‘long-term synaptic potentiation’, (Figure 5, B, C). As for the effects of MIA among stressed animals, the affected ontologies included synaptic functionality, neurodevelopment and cognition (Figure S11 A-C).

**Figure 5:**
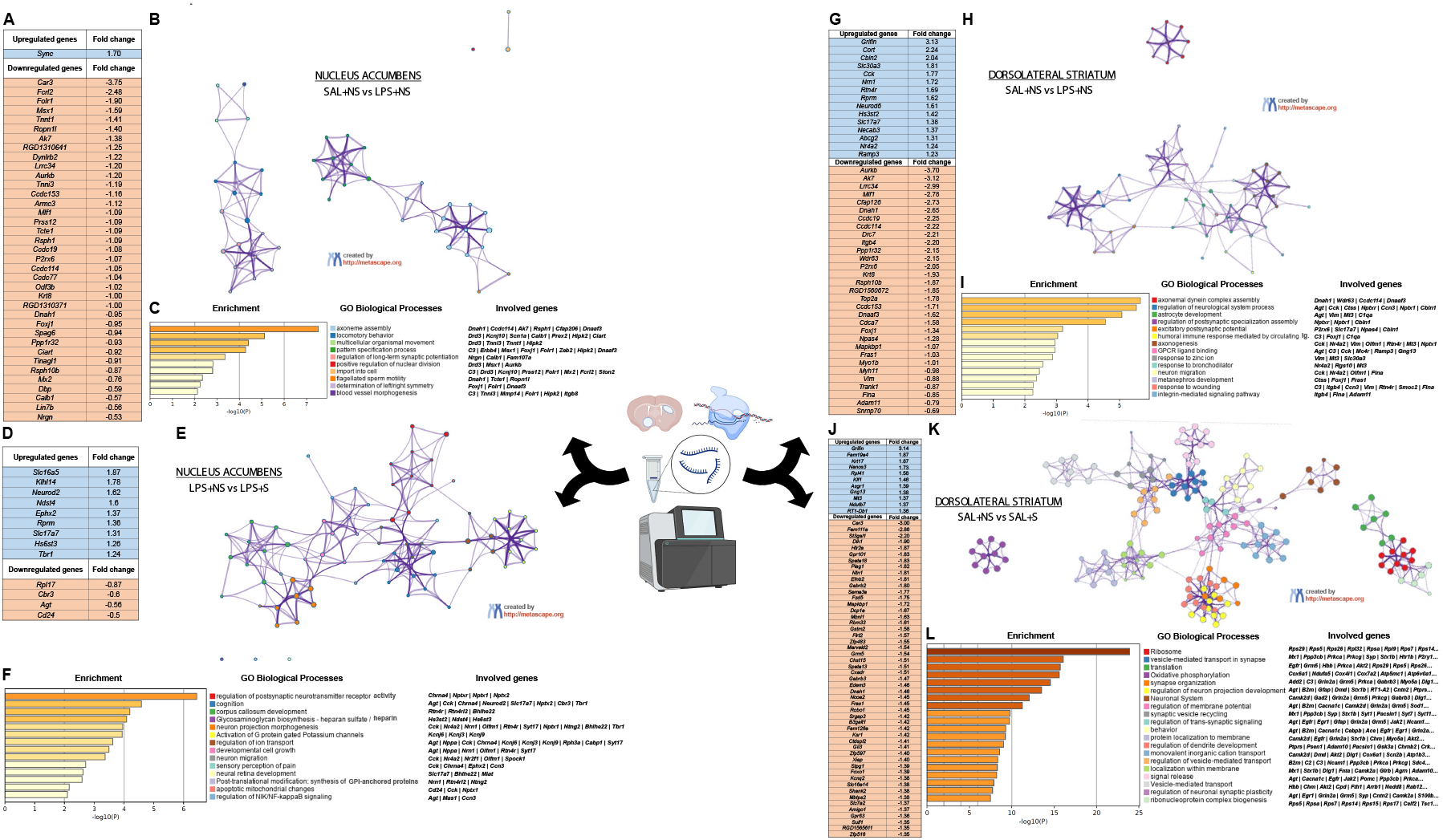
RNA-Seq analysis in the NAcc and dorsolateral striatum. The figure shows (A, D, G, J) main upregulated (Fold change ≥ 1.2) and downregulated (Fold change ≤ -0.5) genes (B, E, H, K) enriched terms network coloured by “GO biological processes” cluster, where nodes that share the same cluster are represented close to each other (C, F, I, L) enriched terms bar graph coloured by p-value (where terms containing more genes show more significant p-value), GO biological processes and involved genes. (SAL+NS: n=4; LPS+NS: n=4; SAL+S= 4; LPS+S=4).

Interestingly, while PUS had no effects on its own, it changed the pattern of MIA-induced transcriptional alterations. Indeed, among LPS-exposed animals, PUS altered the expression of genes involved in the regulation of postsynaptic receptors activity, neurodevelopmental processes such as cell growth, neuron migration or neuron morphogenesis, and even in more complex processes such as ‘cognition’ and ‘sensory perception of pain’ (Figure 5, D-F). Tables S3-S5 provide more detailed information on the genes that belong to each ontology.

#### Dorsolateral Striatum

In the dorsolateral striatum, MIA induced the differential expression of 69 genes while this effect was completely obliterated among animals subjected to PUS. Surprisingly PUS alone induced the expression of 1938 DEGs, but this effect was almost completely abolished in rats with a history of MIA (only 15 DEGs were found in the LPS+NS vs LPS+S comparison) (Figure S14). Only five DEGs associated with PUS were shared among MIA-exposed and non-exposed rats (*Aurkb, Ccded153, Dnah1, P2rx6*, and *Rsph10b*). Noteworthy, these five genes were down-regulated in PUS rats but changed the direction of their expression (to up-regulation) when PUS occurred in MIA-exposed rats, suggesting profound epigenetic modifications. There were 37 shared DEGs among those induced by MIA (SAL+NS vs LPS+NS comparison) and PUS (SAL+NS vs SAL+S comparison). Lastly, 14 DEGs were found both due to MIA in non-stressed rats and due to PUS in MIA-exposed animals, but in opposite directions (Figure S14). In general, it seems that the dorsolateral striatum is more vulnerable to the effects of PUS than the NAcc and that it is more sensitive to the bidirectional interactions between MIA and PUS.

The gene ontology analysis showed that MIA induced and repressed the expression of several genes that clustered in networks that, unlike the NAcc, had no obvious separation. In general, the top five ontologies that were regulated were involved in the development of the nervous system such as ‘axoneme assembly’, ‘astrocyte development’, ‘axonogenesis’ or ‘neuron migration’. It also altered the expression of genes involved in biological processes relevant to the functionality of the nervous system, such as the regulation of postsynaptic specialization assembly or the generation of excitatory postsynaptic potentials (Figure 5, G-I). Interestingly, these MIA-induced gene expression alterations were no longer observed among stressed animals. On the other hand, PUS alone altered the expression of nearly 2000 genes involved in a wide variety of biological processes, mainly at the ribosomal and translational machinery level, synaptic organization and function and behaviour, among others (Figure 5 J-L; Table S2). However, MIA completely modulated these PUS-induced effects and, among LPS-exposed animals, PUS only affected the expression of genes involved in axoneme assembly (Figure S12, A-C). Tables S6-S8 provide more detailed information on the genes that belong to each ontology.

## DISCUSSION

In addition to its disabling consequences, schizophrenia is usually associated with higher rates of substance use disorders (20). To shed some light on this issue, in this work we have studied the presence of cocaine addiction-like behaviour and the underlying alterations in brain structure and function in a two-hit animal model relevant for schizophrenia and other neurodevelopmental disorders.

### 1.- Cocaine-addiction like behaviour and associated processes

In order to evaluate cocaine addiction-like behaviour (and not just drug intake as captured by standard FR1 self-administration programs), we deployed a multicomponent self-administration regime aimed at evaluating the initial drug-intake (FR1 sessions), motivation to consume the drug (PR sessions), compulsive seeking (punished seeking), loss of control over seeking (extended access) and risk of relapse (incubation of drug-seeking).

In our categorical analysis, MIA tended to increase the likelihood of acquiring cocaine self-administration; however, it did not affect drug intake among those animals that did acquire the self-administration behaviour. This absence of effects on cocaine intake is consistent with our previous reports that showed MIA had no effects on cocaine self-administration, extinction, progressive ratio performance, dose-response curves or cue-induced reinstatement (13,14). However, PUS had a remarkable effect on drug intake over the acquisition sessions, lowering the amount of cocaine earned by animals exposed to stress. Previous reports in the literature have already suggested that social defeat stress during adolescence might delay the discrimination between active and inactive lever presses (21), but no studies have indicated that the actual amount of self-administered cocaine could be lower than in non-stressed animals. Quite the contrary, experiments with unpredictable stress during adolescence showed a slightly higher preference for cocaine in adult rats in a two-bottle choice test (22).

This decreased cocaine self-administration due to stress was, however, absent in LPS-exposed rats which displayed a similar amount of cocaine intake regardless of their stress experience. These counteracting effects of MIA are consistent with the recent notion that a prior hit may confer resilience or render subjects insensitive to the actions of a second hit later in life). For example, some of the effects of social isolation (the second hit in the following studies) are reduced by prior maternal separation (23), high-fat diet (24), an immune challenge in adolescence (25), or, more recently, a set of behavioural and biochemical alterations (including locomotor hyperactivity, impaired associative memory and reversal learning, elevated hippocampal and frontal cortical cytokine levels, and increased mammalian target of rapamycin (mTOR) activation in the frontal cortex) (26).

After analysing acquisition, we examined other aspects of cocaine addiction-like behaviour such as motivation for the drug (in a PR schedule) or compulsive seeking (17). MIA increased the motivation for cocaine in the first session of the PR schedule which is the one arguably not affected by the actual learning of the contingencies of the task and properly reflects shifts in the motivational status of the animal. We then returned animals to FR1 and again observed the same pattern as in the acquisition phase whereby PUS decreased cocaine intake, but MIA reversed this effect, suggesting that these interactions, being long-lasting, only emerge under low-effort conditions. We then tested compulsive seeking that was not affected by any of our experimental factors. Extended access conditions in self-administration paradigms induce a loss of control over cocaine intake and the emergence of several addiction-related phenomena (27). Under these conditions (6-hour FR1 sessions) we again observed the effects of PUS (a decrease in cocaine intake) and the modulatory effects of MIA on this phenomenon. Interestingly, this pattern of results did not extrapolate to the incubation of seeking.

Summarizing our results so far, MIA tends to facilitate the acquisition of cocaine self-administration, increases the motivation for cocaine and prevents the lowering effects of PUS on cocaine intake, an interaction which is observed in the last days of acquisition, after returning to low-effort conditions following PR and during extended access (all of them FR1 sessions). MIA or PUS had no effects on compulsive seeking or incubation. Importantly, the reported effects do not seem to be due alterations in Pavlovian or instrumental learning *per se*, since we examined the performance of the animals on tasks involving these cognitive functions and found no changes due to MIA, PUS or their interaction.

The specific pattern of addiction vulnerability observed in MIA animals (increased likelihood of acquiring cocaine self-administration and increased motivation) was not paralleled with changes in impulsivity, which was somewhat expected because impulsivity seems to predict the emergence of a full addiction phenotype characterized not only by increased motivation but also by compulsive seeking and increased seeking when the drug is not available (28) (which in our design is captured by lever presses during signalled time-out periods, a variable that was not affected by MIA, PUS or their interaction).

### 2.- Neuroimaging studies

After characterising the addiction-like behaviour of the animals in this two-hit model, we turned to imaging techniques to gain further insight into the associated brain alterations. Given the relevance of this two-hit model to schizophrenia and other neurodevelopmental disorders, we provide a supplementary discussion of the implications of our findings in the context of these diseases (see supplementary information). Table 1 shows a summary of the main results obtained.

PUS decreased the volume of the right hippocampus. Interestingly, PUS also increased the metabolic activity in the right dorsal subicular compartment of the hippocampal formation, an effect that was prevented in MIA rats. Considering that the dorsal subiculum is necessary for cocaine-seeking and taking (29), this abnormal activity of this structure could explain to some extent the persistent decrease in self-administration behaviour observed across most phases of the addiction-like behaviour examination that we have performed, and also the normalization of this behaviour in rats with MIA experience. An additional explanation may rely on the observed increased dorsal striatal volume due to PUS evident only in rats with a history of MIA (especially in the left hemisphere), an effect that was concomitant to a decrease in NAA+NAAG levels in MIA+PUS rats. NAAG is an agonist of the mGluR3 receptor and negatively modulates glutamate levels (30) therefore, the reduced levels of this peptide may result in increased glutamate levels, which is what we observed in a previous study in MIA+PUS rats where we used *ex vivo* 11T MRS (which allowed us to have a separate measure of glutamate from glutamine peak) (31). Considering all these pieces of evidence, it could be suggested that there might be an acceleration of habit formation in MIA+PUS rats after the initial contact with cocaine that may render their responding for cocaine divorced from the (decreased) rewarding actions of the drug (as observed in PUS rats). This assertion is based on the striatal alterations we observed (increased glutamate levels due to decreased inhibition of glutamate release paralleled with increased striatal volume) which would also be consistent with a recent report that suggests that glutamate dynamics in the left putamen of people with a diagnosis of cocaine use disorder are predictive of their reported habit-like behaviour (i.e. the automaticity dimension of the Creature of Habit Scale) (32).

Cortical choline compounds were reduced in PUS alone animals and an effect that was absent in MIA+PUS rats. The GPC+PCh peak has been traditionally interpreted as reflecting membrane turnover or myelin dynamics (33). In the context of cocaine addiction, it has been shown that cocaine users have higher cortical choline levels, however, in that study, a potential confounding effect of recent cocaine use cannot be ruled out (34). Our data may suggest that these metabolites could actually be driving cocaine use rather than being a consequence of it. Indeed, PUS decreased the levels of this peak in our MRS study and also decreased cocaine self-administration while MIA experience prevented both effects of PUS. This possibility opens an interesting avenue for further research that should be explored in the future.

Some of the brain alterations observed are also relevant to schizophrenia and other neurodevelopmental disorders and, as such, are discussed in the Supplementary Discussion.

### 3.- Transcriptomic studies

Rats with LPS exposure during prenatal development had a tendency to acquire cocaine self-administration faster. Moreover, these rats also showed increased motivation for cocaine. We have observed that MIA affected a gene ontology with potential implication for this pattern of results in the NAcc: ‘regulation of long-term synaptic potentiation’, which may affect the capacity for plasticity of the NAcc upon exposure to cocaine. For example, the gene for Neurogranin (*Nrgn*), present in this ontology (Figure 5 A, C; Table S3), responds to cocaine in several ways: it is induced by cocaine in neural progenitor cells (35) and it is related to anhedonia in cocaine users (36). Moreover, neurogranin in the NAcc also regulates the sensitivity to other drugs such as alcohol (37). Another gene present in the ontology, *Calb1* coding for the protein calbindin is also responsive to cocaine self-administration in the NAcc (38). Accordingly, the expression of these cocaine-responsive genes may be suggested to render these individuals more vulnerable to acquire cocaine self-administration.

On the other hand, PUS did not affect gene expression in the NAcc, which contrasts with its strong effects on cocaine self-administration in those animals that acquired the behaviour. We argue that other brain regions which are more susceptible to stress such as the bed nucleus of stria terminalis (39) or the central amygdala (40,41) may be having direct or indirect influences on cocaine reward. However, the abolishment of the PUS-induced reduction in cocaine intake observed in rats exposed to MIA and PUS may be related to transcriptomic changes in the NAcc that are induced by the combination of both hits. Indeed, as opposed to the absence of effects in controls, PUS modulated several gene ontologies in MIA-animals that could account for the reduction in cocaine intake. These categories include ‘regulation of postsynaptic neurotransmitter receptor activity’, ‘regulation of ion transport’ (including ‘activation of G protein gated potassium channels’) and ‘cognition’. Some members of the pentraxin gene family that we found modulated here (*Nptxr* | *Nptx1* | *Nptx2*) (Figure 5, F; Table S5) are involved in anxiety and responses to stress (42), which support their modulation in stressed animals, and more importantly, they are involved in synaptic processes related to cocaine self-administration, cocaine-induced rearrangements of AMPA receptor trafficking and also in the modulation of cocaine-associated memories (43–45). According to these data, this family of proteins could be partially responsible for the increase in self-administration observed in LPS-exposed stressed rats compared to their controls. Other ontologies affected such as that involving potassium channels could also be relevant in the explanation of the restoration of cocaine intake in LPS-exposed stressed animals. Indeed, this family of potassium channels are also involved in cocaine responses (46–48).

PUS profoundly affected the dorsal striatal transcriptome. 1938 DEGs were found in this area in animals that underwent PUS, however, this transcriptomic ‘scar’ did not form in rats that had MIA. The analysis of the gene ontologies affected by PUS points to deep modifications in the translational machinery of striatal cells. Indeed, the ‘Ribosome’ ontology showed a remarkable impact in PUS-exposed rats, together with the ‘Translation’ category and other plasticity-related ontologies such as ‘Synapse organization’. It seems that PUS affects striatal function by perturbing protein translation of synaptic-related proteins. Given that in addition to the NAcc, the dorsal striatum also regulates cocaine intake under fixed-ration 1 conditions (49), it is tempting to speculate that this structure could be aberrantly imposing a break, or decreasing its goal-directed influence (see discussion of neuroimaging findings above) on cocaine-taking responses. In support of this notion, several addiction-associated genes were down-regulated by PUS in the dorsolateral striatum and that could account for the effects of PUS on cocaine self-administration at this early stage. Some examples of these genes include *Oprk1* (Kappa opioid receptor) (50), *Gabra2*, (GABA_A_ Receptor Subunit Alpha2) (51), *Grin2A* (Glutamate Ionotropic Receptor NMDA Type Subunit 2A) (52), *Chrnb2* (Cholinergic Receptor Nicotinic Beta 2 Subunit) (53) or *Gabra4* (Gamma-Aminobutyric Acid Type A Receptor Subunit Alpha4) (54). Interestingly, these gene alterations were obliterated in stressed animals with a MIA background, which could also explain the normalization of cocaine intake when both hits are combined.

We also found interesting patterns of gene expression data as a consequence of PUS and its interaction with MIA in the NAcc or dorsolateral striatum that are relevant to schizophrenia or ASDs and that are discussed in the Supplementary Discussion.

### Caveats and Concluding remarks

There is one caveat to our study that we would like to acknowledge: we only focused on the male population, mainly because our previous data suggested that MIA only had effects in the male sex and also due to the complexity of our design. Our PET and sequencing data also have a reduced sample size so these results should be replicated in future studies.

The results of the present work suggest that the interactions between prenatal immune activation and stress around puberty are complex, bidirectional and, in some instances, unanticipated. Concerning the important question of cocaine addiction in individuals suffering from neurodevelopmental disorders such as schizophrenia, we argue that stress around puberty, instead of unmasking the latent vulnerability to addiction, may induce an anhedonic status that would be prevented by MIA experience. The neural networks responsible for these interactions likely involve several distributed hubs but the neuroimaging and sequencing techniques employed here suggest an important role for the hippocampal formation and the dorsal striatum. From a translational point of view, these results suggest that the increased rates of cocaine use and addiction observed in individuals suffering from schizophrenia would not be induced by the biological mechanisms of the disease but would rather be a consequence of the experiential factors associated with the condition.

## CONTRIBUTIONS

*Conceptualization*: Alejandro Higuera-Matas and Emilio Ambrosio.

*Formal analysis*: Roberto Capellán, Alejandro Higuera-Matas, Miguel Ángel Morcillo, Marta Oteo, Marta Casquero-Veiga, María Luisa Soto-Montenego.

*Funding acquisition*: Emilio Ambrosio and Alejandro Higuera-Matas.

*Research*: Roberto Capellán, Javier Orihuel, Alberto Marcos, Marcos Ucha and Alejandro Higuera-Matas.

*Methodology*: Roberto Capellán, Javier Orihuel, Alberto Marcos, Marcos Ucha, Marta Casquero-Veiga, María Luisa Soto-Montenegro, Manuel Desco, Marta Oteo Vives, Marta Ibáñez Moragues, Natalia Magro Calvo, Miguel Ángel Morcillo and Alejandro Higuera-Matas.

*Project administration*: Emilio Ambrosio and Alejandro Higuera-Matas.

*Supervision*: Alejandro Higuera-Matas and Emilio Ambrosio.

*Writing – original draft*: Roberto Capellán and Alejandro Higuera-Matas.

*Writing – review & editing*: Roberto Capellán and Alejandro Higuera-Matas with the help of the rest of the authors.

## ACKNOWELGMENTS

We thank Rosa Ferrado, Luis Carrillo, Gonzalo Moreno for their excellent technical assistance. We also thank Daniel Capellán for developing the software used to analyse the microstructure of cocaine self-administration.

