## Supplementary Information for "Additive, synergic and antagonistic interactions between maternal immune activation and peripubertal stress in cocaine addiction-like behaviour, morphofunctional brain parameters and striatal transcriptome"

Index

Supplementary Materials and Methods

Supplementary Results

Supplementary Discussion

Supplementary Figures S1-S14

Supplementary Tables ST1-ST8

**Supplementary Materials and Methods**

*Experimental animals*

Experiments were performed on the male offspring of 14-week-old male and 12-week-old female Sprague-Dawley rats obtained from Charles River (France). Animals were kept in a temperature and humidity-controlled environment (23 °C/50–60%), artificial light (12 h/12 h light/dark cycle, lights on at 8 p.m.), *ad libitum* access to food (commercial diet for rodents A04: Panlab, Barcelona, Spain) and tap water, unless otherwise specified. Rats were housed in transparent Plexiglas cages (48.3 cm length x 26.7 cm width x 20.3 cm height). All the procedures performed were compliant with European Union guidelines for the care of laboratory animals (EU Directive 2010/63/EU governing animal experimentation) and approved by the Bioethics Committee of UNED and the Autonomous Community of Madrid (PROEX 078/18).

*Maternal immune activation*

Male and female rats were mated one week after arrival to the animal facility. Vaginal smears were taken daily from the breeder females, and pregnancy was determined by the presence of sperm in the vaginal smear (day 0 of pregnancy). LPS (from *Escherichia coli* 0111: B4 [Sigma-Aldrich]) dissolved in 0.9% NaCl was intraperitoneally injected to pregnant rats at a dose of 100 mg/kg/ml on gestational days (GD) 15 and 16. This dose was chosen based on previous studies showing that it does not significantly alter the percentage of dam survival and that it only has marginal effects on PPI (1–3). Moreover, in a previous study from our group, we showed that this dose had no consequences on social interaction or cocaine self-administration (4). In considering all these pieces of evidence it seems clear that this dose is suitable for a two-hit study where a single insult should not have strong effects on their own. The control group consisted of pregnant rats submitted to the same treatment schedule with saline injection instead of LPS. We imposed a limit of 12 pups per dam, culling the animal surplus. We marked the pups according to their prenatal treatment with a tattoo in the paw and ensured that each dam had an equal number of LPS and saline-exposed pups. In doing so, we homogenized any potential effect of prenatal treatment on maternal behaviour that could affect the development of the offspring.

*Peripubertal unpredictable stress*

Litters were left undisturbed until postnatal day (PND) 21 when they were weaned and grouped in sets of 2-3. Each set belonged to the same litter and treatment. Between PNDs 28 and 38, we exposed the male offspring to peripubertal unpredictable stress (PUS). This stage of development is a critical period known to be highly sensitive to the disrupting effects of traumatizing events relevant to neuropsychiatric disorders (5). The stress protocol included five distinct stressors, that were sequentially applied every other day, in a randomized order: 1) Stress by agitation (30 min in an orbital shaker at 100 rpm). 2) Stress by immobilization (45 min in a cylindrical restrainer under bright light). 3) Water deprivation for 16 h. 4) 10 min of a forced swimming session in a water tank of 40 cm high x 18 cm in diameter at a temperature of 22 ± 1 ºC and a depth of 30 cm. 5) Constant changes of the home cage (five cage changes, with new sawdust, during the dark cycle at random intervals). Non-stressed controls received handling by the same researcher and on the same days as the stressed subjects. This protocol was adapted from the work of Giovanoli et al., (2013) who showed that PUS unmasked several psychological and neurobiological consequences of prenatal immune activation such as latent inhibition and PPI deficits or dopaminergic hyperactivity and increased immune reactivity in the hippocampus.

*Experimental Design*

Four different experimental groups resulted from the manipulations described above: saline exposed animals (SAL+NS), saline & PUS exposed animals (SAL+S), LPS exposed animals (LPS+NS) and LPS & PUS exposed animals (LPS+S). All the studies that will be described hereafter were carried out in independent batches of animals, except for the PPI test, which was performed in all of them. The exact number of litters per experiment is as follows: Self-administration experiment: Saline: N=9; LPS; N=9; Pavlovian and instrumental conditioning experiment: Saline: N=9; LPS; N=12; 2-CSRTT experiment: Saline: N=4; LPS; N=3; MRI experiments: Saline: N=5; LPS; N=5; PET experiment: Saline: N=4; LPS; N=4; RNAseq experiment: Saline: N=4; LPS; N=4. Figure S1 shows an outline of the experimental design employed. The final sample size of each of the four experimental groups is indicated in the figure footnotes.

*Experimental Procedures*

Prepulse inhibition of the acoustic startle response (PPI)

At PND70-73, PPI of the acoustic startle was measured in a nonrestrictive Plexiglas cage (28x15x17 cm) containing a vibration-sensitive platform (Cibertec), enclosed in a sound-attenuating chamber, and a set of two speakers located above the cage. Rats were habituated for 7 min with a background noise of 65 decibels (dB), which continued throughout the session. Animals were exposed to 6 pulse-alone trials at the beginning and the end of the session, to stabilize the startle response and to calculate habituation percentage (these pulses were not included in the PPI calculations). The session was composed of 35 different trials: ten 120 dB pulse-alone trials, five null trials with no stimulus and twenty pulses preceded by a prepulse of 69- or 77-dB intensity (4 or 12 dB above the background noise, respectively) with an interval of 30 or 120 ms (4). The duration of the test was 20 min approximately. Prepulse inhibition is expressed as the % PPI and calculated using the following formula: 1 − [startle amplitude on prepulse + pulse trial/mean startle amplitude on pulse-alone trials]) x 100. The percentage of habituation is expressed as 100 x [(Mean of first pulse-alone block − Mean of last pulse-alone block) ⁄ Mean of first 6-pulse alone block].

Cocaine self-administration

At PND80 animals underwent surgery to be implanted with a jugular vein catheter that allowed intravenous cocaine self-administration. After this, they were single-housed in independent home cages. Surgeries were performed under isoflurane anaesthesia (5% for induction and 1.5-2% during maintenance) and buprenorphine analgesia (0.05 mg/kg s.c.). An incision was made to implant a polyvinyl chloride catheter (0.064 i.d.) into the jugular vein at the atrium level. This catheter was implanted subcutaneously and reached the exterior in the middle scapular region. Another incision was made on the back of the animal, where a mesh attached to a dental cement-made pedestal and a screw (Plastics One) was placed. After surgery, the rats were allowed to recover for 10 days and a nonsteroidal anti-inflammatory drug (meloxicam, Metacam™: 15 drops of a 1.5 g/mL solution per 500 mL of water) was added to the drinking water during the first five recovery days and the antibiotic Marbocyl^TM^ (marbofloxacin) was administered during the first three recovery days (2 mg/kg s.c.). After the surgery and during the experiment, they were infused daily through the catheter with 0.5 ml of a saline solution with heparin (1.5 IU/ml) and gentamicin (40 mg/ml) to prevent possible catheter infections and to ensure patency. On PND90, the cocaine self-administration program began with six different phases as described below. All sessions (one per day) were carried out in operant conditioning boxes (Coulbourn Instruments) which had two levers, an “active” lever that was associated with the drug infusion and another “inactive” lever without programmed consequences. The active lever triggered an infusion pump that delivered the infusion of cocaine (Alcaliber) at a dose of 0.5 mg/kg throughout a tubing that connects with the catheter previously implanted in the animal. The tubing reached the animal within a thin spring theter attached at the upper end of the conditioning box to prevent breakage. The end of the spring was attached to a screw that was implanted on the back of the animal. At the beginning of the sessions, a cue light was activated on the active lever, which was used as a discriminative stimulus signalling the availability of the drug. When the animal obtained an infusion, a time out period of 7 seconds began (same length as the infusion) in which the discriminative stimulus disappeared, and the active lever presses had no consequence. The first phase of the cocaine self-administration program, the Acquisition phase, consisted of 12 sessions, of 2 hours each in an FR1 program, in which an active lever press initiated the infusion of the drug by the pump. Rats were categorised as showing: “Regular acquisition” (animals that showed cocaine self-administration behaviour i.e. more than 5 infusions over the last three days of acquisition), “Late acquisition” (animals that took more than 12 sessions to present a stable behaviour of cocaine self-administration) and “No-acquisition” (animals that by the end of the experiment never showed a cocaine self-administration behaviour). The second phase, where motivation for consumption was studied, consisted of 6 sessions of 2 hours each in a progressive ratio program, in which the lever presses required to obtain an infusion increased after each infusion according the following sequence: 1, 2, 3, 4, 5, 6, 7, 8, 10, 12, 14, 16, 18, 20, 22, 24, 28, 32, 36, 40, 44, 48, 52, 56, 64, 72, 80, 88, 96, 104, etc...). In the third phase, Stabilization, the aim was to rebaseline the lever press, to do this, we used three sessions of identical conditions as those used in the acquisition phase. The fourth phase, where cocaine compulsive consumption was measured, consisted of a single session of one hour in which under an FR3, the animals randomly received an infusion or a mild plant shock of 0.5 mA for 0.5 seconds. The fifth phase, Extended Access, consisted of 10 days with sessions of 6 hours in FR1. At the end of this phase, a forced abstinence period was imposed, to evaluate Cue Induced Relapses, leading to the sixth phase. Drug-seeking sessions occurred on days 1, 30, 60 and 100 after the beginning of abstinence, in the same conditions as those described in the Acquisition phase but without any solution flowing throughout the system. The day before starting the self-administration program and after the last Extended Access session, catheter functionality was checked by a 0.1-0.2 mL infusion of a thiopental solution (Northia, 20 mg/mL, NaCl 0.9%). The catheter was considered to be functional if the animal showed an immediate loss of consciousness or motor coordination impairment after the infusion.

Pavlovian and Instrumental conditioning programs

At PND90, food was removed, and body weight was controlled so that it remained between 90-95%. Food was provided after each experimental session. The Pavlovian learning protocol consisted of eight daily sessions. Each session consisted of 4 cycles of 12 minutes. In each cycle, animals were exposed to 4 minutes of a continuous “tone” stimulus, 4 minutes of an intermittent “click” stimulus and 4 minutes in the absence of stimuli between both. During one of the auditory stimuli exposure (either tone or click), pellets were dropped according to a 5-second variable time program (different time intervals with an average time of five seconds). The conditioned stimulus "CS+", signalled cue-delivery periods whereas the absence of pellets was signalled by the "CS-" stimulus; in both cases head entries (HEs) to the feeder were recorded. The 4-minute interval between CS + and CS- was the ISI (inter-stimulus interval), where only the white noise produced by a fan designed to isolate the sound boxes from the outside was maintained. CS+ and CS- were counterbalanced between subjects to avoid attentional biases derived from the unconditioned excitatory capacity of each stimulus, as well as the order in which they were presented. During this protocol, Skinner boxes had no visible operating levers.

The instrumental learning protocol was performed after the Pavlovian training and consisted of seven daily sessions. Animals had two levers available. Pressing on one of them produced pellet release so that this lever was called "active", while pressing the other one (“inactive”) had no consequence. Once the active lever was pressed, a 5-second timeout began with no pellets. Sessions ended after 30 minutes or after 30 pellets had been earned. Animals completed one fixed ratio 1 (FR1) session, three variable ratio-5 (VR5) sessions and three variable ratio-10 (VR10) sessions. No light cues were presented during the sessions. White noise was present during the sessions to dim the noise from outside the operant chambers.

Two-choice serial reaction time task (2-CSRTT)

Once the PND90 was reached, food was removed, and body weight was controlled so that it remained between 90-95% of free-feeding values. Food was provided after each experimental session. The 2-CSRTT employed here was an adaptation from the 5-choice serial reaction time task protocol (6). We used Skinner boxes (Med Associates) equipped with a feeder with entry detectors, two retractable levers placed at both sides of the feeder, and two small lamps to present cue lights above the levers. Before the beginning of the protocol, a brief training was implemented to associate the light stimulus placed on top of each lever with the release of one pellet when the lever was pressed. This training consisted of two sessions, one for each lever, with a 30 minute or 30 pellets limit. The actual 2-CSRTT protocol consisted of 13 different stages, and each one was composed of 100 serial trials. Light stimulus duration was progressively reduced throughout stages from 30 to 0.5 seconds. Response time (or time to respond since light stimulus appeared) was also diminished from 30 to 5 seconds, while inter-trial interval (ITI) (or time elapsed between trials) increased from 2 to 9 seconds. Trials started when the animal introduced its head inside the feeder, and the signalled lever was alternated randomly during these trials. Correct answers were rewarded with a food pellet, while incorrect answers, omissions (absence of response) and premature responses (lever presses before light stimulus presentation) were punished with a 5-second time-out, where the Skinner box remained in darkness, and no rewards were available. Each session ended after 100 trials were completed or after 30 minutes, whichever occurred first, and accuracy and omissions percentages were calculated to determine if the animal could progress to the next stage (6). Once stage 12 was reached, animals remained in this phase until six consecutive sessions were successfully completed, to stabilize performance. The criterion applied in this phase was ≥75% accuracy and ≤20% omissions. Afterwards, three long ITI sessions (inter-trial interval was increased to 9 seconds) were performed, separated by two baseline sessions each. This was done to elicit impulsive behaviour (7). Between 25 and 45 sessions were carried out until all the animals reached this point.

Neuroimaging studies

Magnetic resonance imaging (MRI) and diffusion tensor imaging (DTI)

At PND90, MRI and DTI studies were executed at the Biomedical Research Institute "Alberto Sols" (CSIC-UAM, Madrid, Spain). Experiments were performed on a Bruker PharmaScan system (Bruker Medical Gmbh, Ettlingen, Germany) using a 7.0 Tesla horizontal-bore superconducting magnet, equipped with a ^1^H selective quadrature 40 mm coil and a 90 mm-diameter gradient insert (36 G/cm maximum intensity). All data were acquired using a Hewlett-Packard console running Paravision 5.1 software (Bruker Medical Gmbh) operating on a Linux platform. Rats were placed into the centre of the radiofrequency volume coil and positioned in the magnet under continuous anaesthesia inhalation via a nose cone. A respiratory sensor connected to a monitoring system (SA Instruments, Stony Brook, NY) was placed under the abdomen to monitor respiration rate and depth. Animals were anaesthetized with a 2% isoflurane-oxygen mixture in an induction chamber and the flow of anaesthetic gas was constantly regulated to maintain a breathing rate of 50 +/- 20 beats per minute. T2-weighted (T2-W) spin-echo anatomical images were acquired with a rapid acquisition with relaxation enhancement (RARE) sequence in axial and coronal orientations and using the following parameters: TR: 3000 ms, TE: 44 ms, RARE: factor 8, Averages: 3, FOV: 3,5 cm, Acquisition matrix: 256 × 256 corresponding to an in-plane resolution of 136 × 136 μm^2^, Slice thickness: 1,5 mm, Number of slices: 18 for axial and 8 for coronal images. Diffusion-weighted images were acquired with a spin-echo single-shot echo-planar imaging (EPI) pulse sequence using the following parameters: TR: 3500 ms, TE: 40 ms, Averages: 1, Diffusion gradient duration: 3,5 ms, Diffusion gradient separation: 20 ms, Gradient directions: 7, Acquisition matrix: 96x96 and zero-filled in k-space to construct a 128 × 128 corresponding to an in-plane resolution of 273x273 μm^2^, B values: 100 s/mm^2^ and 1400 s/mm^2^, Slices thickness 1,5 mm. Fractional anisotropy, mean diffusivity, trace, eigenvalues and eigenvector maps were calculated with a homemade software application written in Matlab (R2007a). Values were extracted from maps using regions of interest (ROIs) with Image J software. The volumetric quantification of the different brain structures or ventricles was normalized to the brain volume of the MRI slice containing the analysed region, ruling out potential confounding effects of the differences in brain volume observed as a consequence of MIA.

In vivo proton magnetic resonance spectroscopy (^1^H-MRS)

Immediately after conducting the MRI and DTI analysis, an *in vivo* ^1^H MRS study was also performed in the Biomedical Research Institute "Alberto Sols" (CSIC-UAM, Madrid, Spain). Two brain regions were selected for this study: cortex and striatum. A Point-REsolved Spatially Spectroscopy (PRESS) was used, combined with a variable power radiofrequency (VAPOR) water suppression and employing the following parameters: TR: 3000 ms, TE: 35 ms, Averages: 128, Voxel volume: 3 mm^3^. First and second-order shims were automatically adjusted with a fast, automatic shimming technique by mapping along projections (FASTMAP) in a large voxel (4 mm^3^). The spectra were automatically analyzed using LCModel software (8), 6.2-OR version (Oakville, ON; Canada). Only the peak concentrations obtained with a standard deviation lower than 20% were accepted.

Positron emission tomography/computed tomography (PET-CT)

Once PND90 was reached, PET-CT studies were performed at the Radioisotopes for Biomedicine research group of the Center for Energy, Environmental and Technological Research (CIEMAT) in Madrid, Spain, using a small-animal PET-CT scanner (Argus PET/CT, SEDECAL, Madrid, Spain). PET (400–700 KeV energy window, 45 min static acquisition time) and CT studies (45 kV voltage, 150 μA current intensity, 8 shots, 360 projections, standard resolution) were performed 30 minutes after inoculation of 15.4±1.4 MBq of [^18^F]-2-fluoro-2-deoxy-d-glucose (^18^F-FDG) via the tail vein. Animals were anaesthetized by inhalation of 2–2.5% isoflurane in 100% oxygen. PET-CT images reconstruction was accomplished using a 2-dimension ordered subset expectation maximization (2D-OSEM) algorithm (16 subsets and 3 iterations), with random and scatter correction. The relatively poor spatial resolution of PET-CT imaging and the difficulty in identifying anatomical regions was minimized by co-registering PET-CT images to same-subject MR images (9). Brain masks (corresponding to hippocampus, caudate bodies, prefrontal cortex, cortex and whole-body) were manually segmented on the MR template and applied to their corresponding PET-CT study. Voxel value normalization consisted of standardizing PET intensity data to a brain region without statistically significant differences between groups obtained by an iterative method (see (10) for further details). We used a full ANOVA designed followed by pair-wise comparisons after significant main effects. Statistical comparisons (p<0.01 uncorrected) were performed with Statistical Parametric Mapping (SPM) software (http://www.fil.ion.ucl.ac.uk/spm/software/spm12/). Registered PET images were smoothed with a gaussian kernel of 2.5 times the voxel size of full width at a half maximum (FWHM) and masked in order to exclude extracerebral voxels from the analyses. Only clusters larger than 50 adjacent voxels were considered to minimize the effect of type I errors.

RNAseq

Once PND90 was reached, animals were decapitated for the dissection of different brain structures, under isoflurane anaesthesia. All the dissection material was autoclaved and treated with RNase*Zap*™ (Invitrogen) to avoid RNA degradation by RNases. In addition, water and saline solutions containing diethyl pyrocarbonate (Sigma-Aldrich) (1:500) were used to wash the dissecting material and tissue, respectively. An acrylic brain matrix for a 300 g - 600 g rat and razor blades were used for brain slicing, using (11) as a reference. The entire procedure was carried out at 4º C to prevent tissue degradation. The dissected samples were preserved in RNAlater™ Stabilization Solution (Invitrogen) for one day at -20°C and subsequently stored at -80°C. RNA extraction was performed using the RNeasy Mini Kit (Qiagen). RNA-Seq analysis was carried out in the Genomics Unit of the Madrid Science Park. RNA integrity number (RIN) and concentration were evaluated by employing an Agilent 2100 Bioanalyzer using an RNA 6000 nano LabChip kit. Libraries were prepared according to the “NEBNext Ultra Directional RNA Library Prep kit for Illumina” (New England Biolabs) instructions. “Chapter 1: Protocol for use with NEBNext Poly(A) mRNA Magnetic Isolation Module” indications were followed. Before starting the protocol, the total RNA input yield was 1 µg. A 14-cycle PCR was used to obtain the library amplification included in the mentioned protocol. Libraries were validated and quantified by an Agilent 2100 Bioanalyzer using a DNA7500 LabChip kit. An equimolecular pool of libraries was titrated by quantitative PCR using the “Kapa-SYBR FAST qPCR kit forLightCycler480” (Kapa BioSystems) and a reference standard was used for quantification. The pool of libraries was denatured before being seeded on a flowcell at a 2,2 pM density, where bunches were formed and sequenced using a “NextSeq™ 500 High Output Kit”, in a 1x75 single read sequencing run on a NextSeq500 sequencer. Once the sequencing process was finished, the Illumina Analysis Space tool was used to map and locate the different sequences in the reference genome, generating alignment files in “. bam" format. These files were used to perform a differential expression analysis using the CUFFDIFF tool, which counted the RNA expression of each gene, normalized by its size and by the global RNA expression of each sample, and made a comparison between groups applying a False Discovery Rate (FDR) correction. We then used the Metascape (https://metascape.org/gp/index.html#/main/step1) resource to analyze the enrichment in specific gene ontologies for each comparison.

All RNA-seq data sets generated and/or analyzed during the current study were added to the Gene Expression Omnibus (GEO) under the accession number GSE185195.

*Statistical Analysis*

Data were analyzed with IBM SPSS Statistics 24 for Windows. All results are expressed as the mean ± standard error of the mean (SEM) in the graphs or mean ± standard deviation (SD) in the tables. Outliers were identified by SPSS using the interquartile range criterion and a value of p < 0.05 was considered to represent a statistically significant difference. Square root, Neperian logarithmic and inverse transformations were applied when appropriate to correct the skewness in the distribution of the data and the lack of homogeneity of variances. All results were analyzed by two-way ANOVA considering “Prenatal immune activation” (saline or LPS) and “PUS exposure” (no stress or stress) as between-subject factors. The within-subject “Sessions” factor was considered in cocaine self-administration, Pavlovian and instrumental training procedures and the 2-CSRTT. Pearson's chi-squared test was performed for the categorical analysis of the acquisition phase of the cocaine self-administration program. We analyzed interactions by simple effects analysis with the Bonferroni correction for multiple comparisons. F-value, effect sizes (partial eta square, η^2^_p_) and degrees of freedom are also reported when appropriate.

**Supplementary Results**

Analysis of Pavlovian and instrumental conditioning

To rule out potential alterations in Pavlovian or instrumental conditioning that may have affected the self-administration data, we analysed if prenatal immune activation or PUS these two forms of learning. Animals progressively learnt both tasks, as revealed by the significant effects of the Pavlovian (F_1,56_=20.181; p=0.000; η^2^p=0.265) (Figure S4, A) and Instrumental (F_1,56_=39.638; p=0.000; η^2^p=0.414) (Figure S4, B) sessions factors, however, no significant differences were due to prenatal immune activation, PUS or their interaction.

Motor impulsivity

Given that impulsivity is an endophenotype that confers vulnerability to addiction, we asked if prenatal immune activation, PUS or their interaction affected impulsivity. Impulsive behaviour, as captured by the 2-CSRTT, was not modified by prenatal immune activation or PUS. Indeed, no significant effects were observed by prenatal immune activation, PUS or their interaction in % premature responses in any of the sessions (Figure S5, A), nor in the normalised responses during long-ITI sessions (Figure S5, B and C). A significant effect of the Sessions factor was found in correct responses (F_1,27_=20.177; p=0.000; η^2^p=0.428), incorrect responses (F_1,27_=4.756; p=0.001; η^2^p=0.150), omissions (F_1,27_=3.331; p=0.021; η^2^p=0.118), perseverative responses (F_1,27_=3.305; p=0.010; η^2^p=0.109) and premature responses (F_1,27_=33.707; p=0.000; η^2^p=0.555), however, no significant effects of the prenatal immune activation or PUS factors or their interaction were detected, ruling out potential deficits in sustained attention in these animals (Figure S5, C-G). In addition, no significant differences were observed in the number of sessions to reach stage 12 (Figure S5, H).

**Supplementary Discussion**

**Brain imaging alterations in the context of schizophrenia**

We will first discuss the effects of maternal immune activation (MIA) on its own and then examine MIA-PUS interactions. We found evidence for smaller whole brain volume in the rats exposed to LPS during gestation. This decrease in whole brain size (previously undocumented in LPS-induced MIA models) cannot be due to a concomitant decrease in body size because these animals had slightly increased body weight (mean±SD g: SAL+NS= 387.375±18.700; SAL+S=373.563±16.950; LPS+NS=410.937±23.420; LPS+S= 421.500±30.833). Even if decreased whole brain volume is a general landmark in schizophrenia (12), general reductions in whole brain volume are scarce in MIA experiments. Indeed, there are some MRI studies in the MIA literature that, relying on TLR3 activation via poly I:C (and hence mimicking viral infections rather than bacterial infections), have shown reductions in brain volume; however in one of them these reductions were concomitant with increases in the volume of other areas of the brain (13) and, in other study, the decrease in general brain volume was transient, emerging on PND35 and disappearing upon reaching adulthood (14). Moreover, other poly I:C MIA studies have not found evidence for decreased whole brain volume (15–17). Therefore, the results obtained with our specific parameters and immunogen provide further support for the notion that the reduction in whole-brain volume could indeed be a neurodevelopmental trait associated with schizophrenia, and not a consequence of the antipsychotic medication or other correlated variables, such as adverse life events or drug use. We also found evidence for increased MD in rats with gestational exposure to LPS. This increase in MD could point to reduced cellular and synaptic complexity (18,19) and may imply a delay in the maturational processes that occur in the hippocampus. In addition, these preclinical findings also provide support to the evidence of an increase in MD in the hippocampus of people with a diagnosis of schizophrenia (19), increasing the potential usefulness of this imaging landmark as an early diagnostic marker of the disease, even if the whole symptom clusters have not fully emerged. We did not detect any alterations due to MIA in brain metabolic activity in our PET experiments or the metabolite profile of the cortex or dorsal striatum in our ^1^H-MRS studies. These results contrast with previous literature and this divergence may be resulting from the different immunogen used in the studies. Indeed, while we used LPS in the present experiments, which mimics a bacterial infection, all previous reports have used poly I:C (16,20–25). These is important in the context of the understanding of the differential effects that viral and bacterial infections may have in the developing brain.

While the effects of MIA on its own are interesting to examine brain alterations that may underly the susceptibility to schizophrenia and associated neurodevelopmental disorders, the addition of a second hit may be more relevant for the actual onset of the condition. In this context, we have observed very interesting MIA * PUS interactions. For example, MIA induced an increase in the striatal volume only in stressed animals which, however, was paralleled by a potential decrease in neuronal density (as indicated by decreased NAA+NAAG concentrations), suggesting that the higher volume of this structure in MIA-PUS rats may be more likely due to glial proliferation. Metanalysis of striatal NAA levels have not found consistent evidence for reduced levels of this metabolite in the basal ganglia in schizophrenia (26,27), however, our findings with the animal model used here, together with our previous evidence for increased glutamate levels in the striatum of MIA+PUS rats (28), suggest that further investigations are needed to elucidate the actual role of these striatal alterations in the context of schizophrenia-spectrum disorders.

**Gene expression changes relevant to neurodevelopmental disorders**

While MIA alone did not seem to induce relevant gene expression changes in the nucleus accumbens (NAcc) for schizophrenia or autism-spectrum disorders (ASDs) (it affected genes related to the assembly of the axoneme and cilia-related genes), the effects of MIA in stressed animals shifted towards alterations of gene expression that involved a set of ontologies relevant for neurodevelopmental disorders. Indeed, categories important for the regulation of the synaptic structure, the transport of aminoacids such as glutamate, axonogenesis or cognition were significantly enriched. At the individual gene level in the NAcc, there were some genes with important associations to schizophrenia or ASDs that were affected in MIA+PUS rats and that were not modulated in rats with MIA but not stress experience. Some examples of these genes are *Rtn4r* (Reticulon 4 Receptor) (which was also modulated by MIA in the dorsolateral striatum) (29–31), *Tbr1* (T-Box Brain Transcription Factor 1) (modulated also by PUS but only among MIA-exposed rats) (32–34), *Bdnf* (Brain Derived Neurotrophic Factor) (35,36) or *Slc17a7* (Solute Carrier Family 17 Member 7 or vesicular glutamate transporter 1) (37,38). This transcriptomic landscape in the NAcc induced by the combination of MIA and PUS could therefore provide susceptibility to schizophrenia or autism even if the symptoms of the disease have not fully emerged (for example, in the present study, the documented decrease in PPI was at the threshold of significance, suggesting that the full emergence of the disease had not completed).

The dorsal striatum is also a structure that is affected in schizophrenia (39,40) and autism (41). Here, similarly to the NAcc, MIA also affected the axoneme assembly ontology, however, it also modulated other categories that are more relevant to neurodevelopmental disorders such as the regulation of neurological system process (including important genes such as *Cck* which codes for the peptide cholecystokinin) or the regulation of the excitatory postsynaptic potential (which also includes relevant genes to schizophrenia or autism such as *Slc17a7* which codes de vesicular glutamate transporter 1 protein or VgluT1). At the individual level, an important gene modulated by MIA was *Rtn4r* which, as stated above has been associated with schizophrenia and, in the NAcc, required the combination of MIA and PUS to be up-regulated. MIA also up-regulated *Cck* in the dorsolateral striatum, a peptide that has been involved in schizophrenia (42). Indeed, Adequate levels of cholecystokinin in the striatum have been suggested to control the interaction between cognition and reward circuitry, which seems to be crucial in schizophrenia (43). The evidence linking *Cck* to autism is much more scarce but a 3p22.1p21.31 microdeletion study has suggested that cholecystokinin could be a candidate gene for the disorder previously known as Asperger syndrome (44). The last gene that we would like to highlight regarding MIA effects in the striatum is *Nr4a2* which codes for the Nuclear Receptor Subfamily 4 Group A Member 2 protein (also known as *Nurr1*). This protein is a transcription factor implied in the differentiation, maturation, and survival of dopaminergic neurons and also has a role in regulating the expression of several proteins important for the synthesis and/or regulation of dopamine (DA) (45,46) and was involved in the attentional deficits of patients with schizophrenia in one study (47). As a whole, these results suggest that MIA in the striatum (as opposed to what is observed in the NAcc) is able to induce, on its own several gene expression alterations that may confer vulnerability to schizophrenia.

However, as stated in the previous section, the most notable effect in the striatum was the profound striatal alterations induced by PUS. 1938 DEGs were affected by stress in this structure. Some of the relevant ontologies, with potential implications for neurodevelopmental disorders, were those related to synapse organization, neuron projection development, trans-synaptic signalling or behaviour. Several of the genes in these categories belonged to the glutamatergic or GABAergic systems and have been previously shown to be involved in schizophrenia or autism. Some examples are *Slc1a1, Grin2a*, *Gabra2, Gabrg2*, *Gria1*, *Grik2* or *Gad2.* Another example that we would like to highlight is the *Ngln3* gene, down-regulated by PUS, that codes for the Neuroligin 3 protein, a postsynaptic adhesion molecule with a heavy association to autism-spectrum disorders (48). As a result, in the striatum, PUS induced, on its own, several genes that are known to be related to schizophrenia or autism, suggesting at stress at peripuberty could prime this structure, acting as a susceptibility factor. Interestingly, when MIA and PUS were combined, the striatal transcriptomic landscape completely changed. PUS among animals exposed to MIA no longer affected the previously mentioned categories and biased the effects toward a specific set of genes related to the regulation of axoneme assembly. Moreover, at the induvial gene level, there were no genes with a strong involvement in neurodevelopmental disorders. Hence, at least at the striatal transcriptome level, it seems that MIA is protecting against the deleterious effects of stress with regards to the susceptibility to autism of schizophrenia, two of the most severe neurodevelopmental disorders.

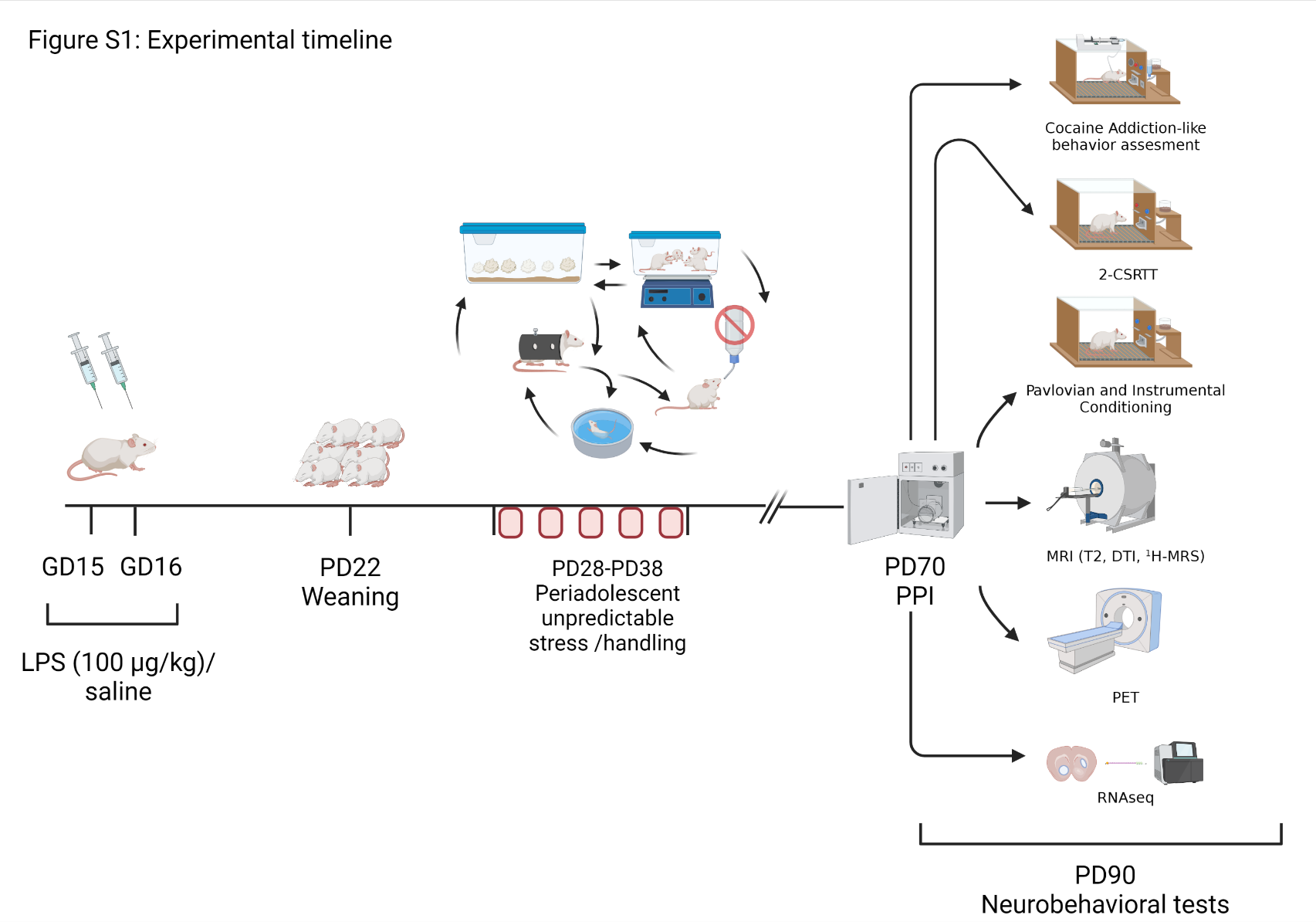

**Figure S1: Experimental Timeline.** Outline of the experimental design followed.

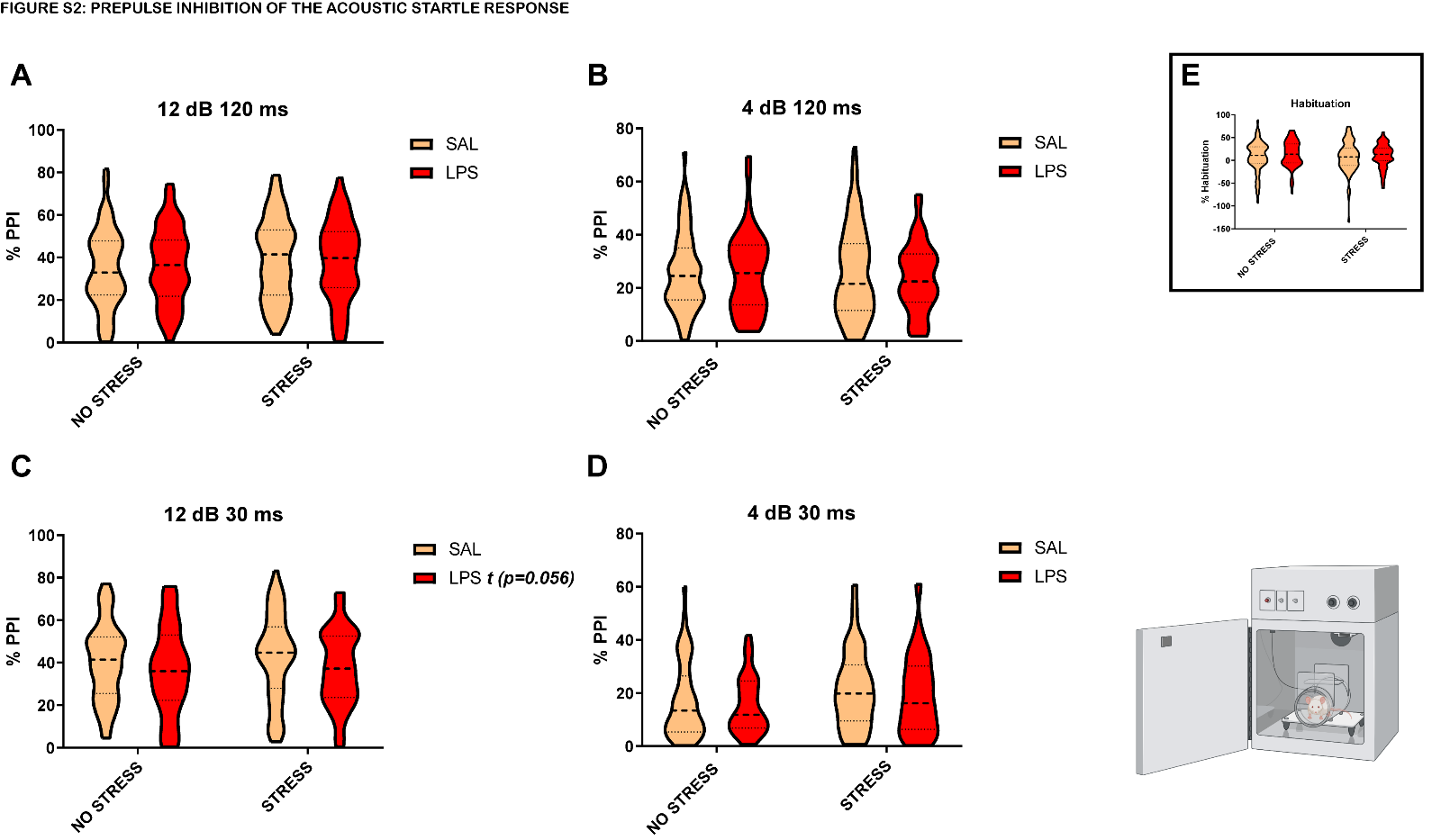

**Figure S2: Effects of prenatal LPS treatment or PUS in % PPI.** The figure shows % PPI at (A) 12 dB prepulse intensity and 120 ms interval (SAL+NS: n=82; SAL+S: n=82; LPS+NS: n=56; LPS+S: n=55); (B) 4 dB prepulse intensity and 120 ms interval (SAL+NS: n=69; SAL+S: n=69; LPS+NS: n=50; LPS+S: n=44); (C) 12 dB prepulse intensity and 30 ms interval (SAL+NS: n=80; SAL+S: n=83; LPS+NS: n=59; LPS+S: n=52). (D) 4 dB prepulse intensity and 30 ms interval (SAL+NS: n=50; SAL+S: n=46; LPS+NS: n=33; LPS+S: n=27). Note that animals with negative values (suggestive of prepulse facilitation -PPF-) were discarded, and hence the difference in sample size across prepulse intensities (lower prepulse intensities tend to yield negative PPI values indicating PPF).

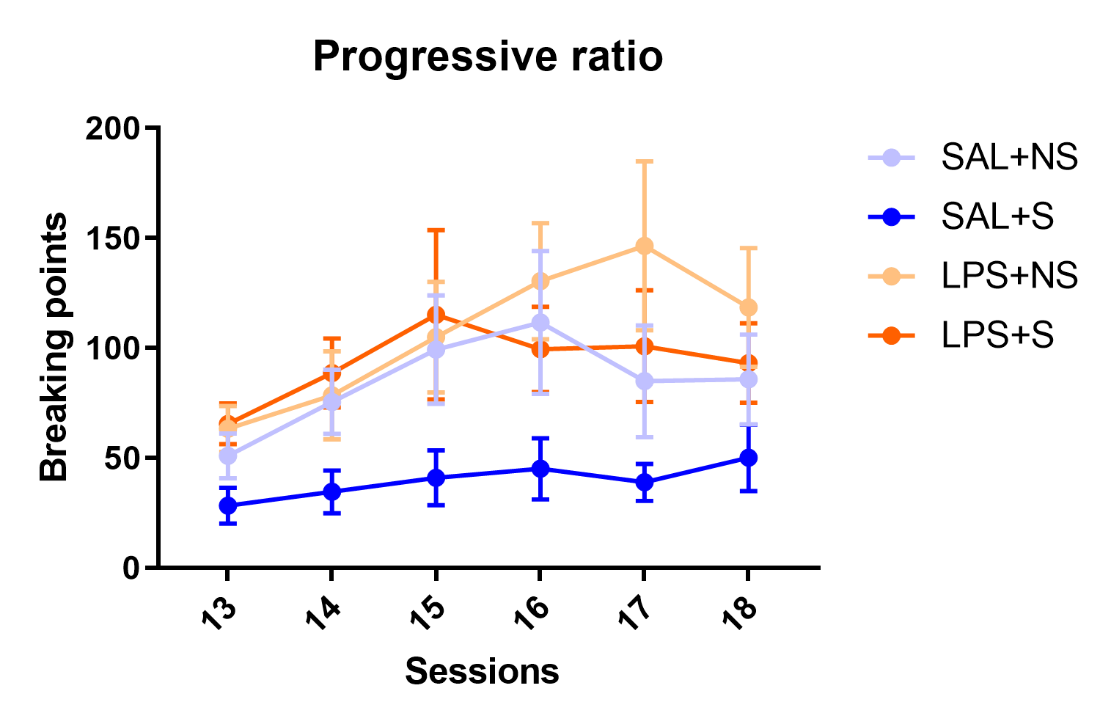

**Figure S3: Breaking points across the progressive ratio session:** Breaking points (defined as the last ordinal value of injection achieved before the rat failed to complete the requirement of an additional injection) across the progressive ration sessions. Rats showed a stable performance across sessions. No significant effects were observed as a consequence of MIA, PUS or interaction (SAL+NS: n=13; SAL+S: n=13; LPS+NS: n=13; LPS+S: n=15).

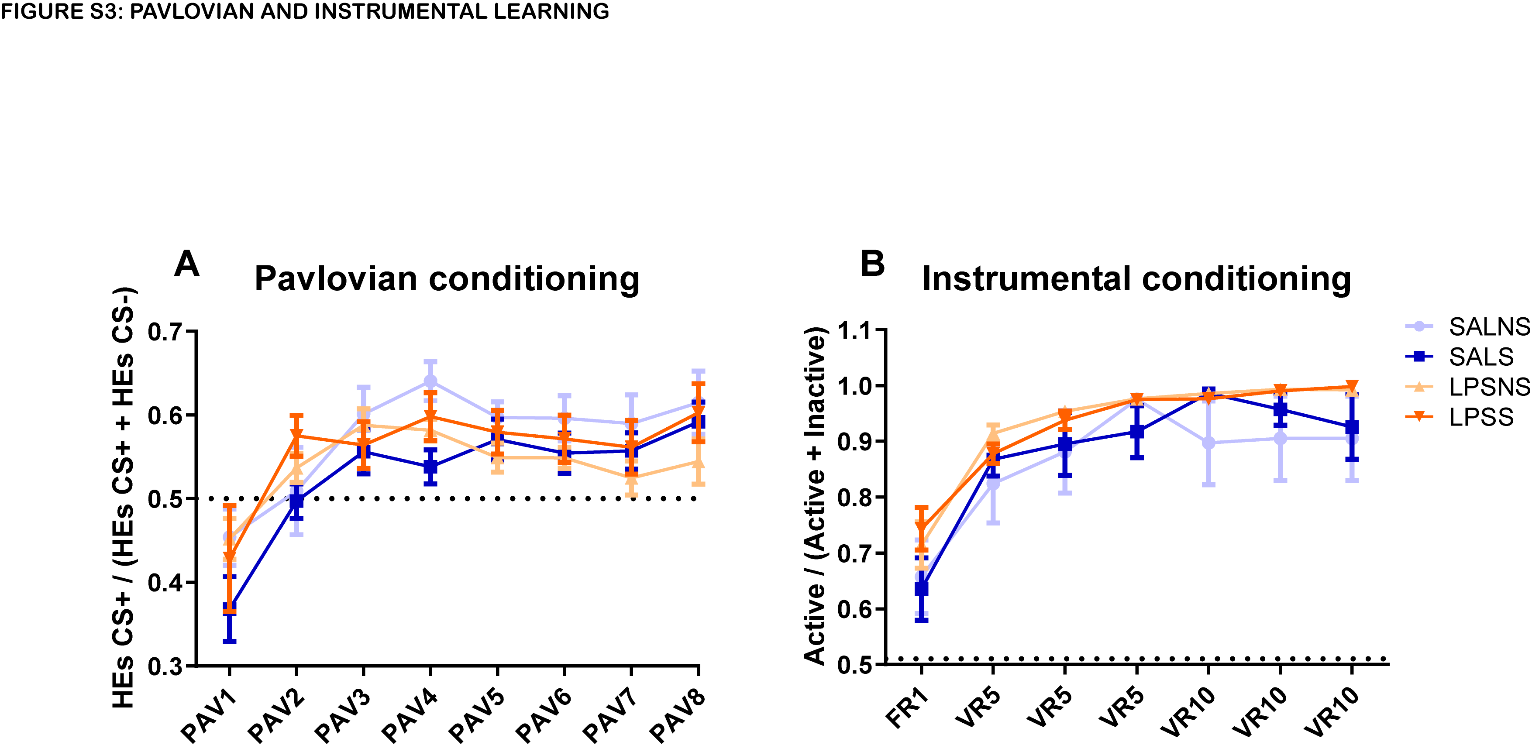

**Figure S4: Performance on Pavlovian and Instrumental Conditioning Tasks:** Ratio of head entries during the presence of the stimulus predictive of reward delivery (CS+) over the head entries during CS+ and the stimulus predictive of reward absence (CS+), across the eight conditioning sessions. Rats showed a stable performance across sessions. No significant effects were observed as a consequence of MIA, PUS or interaction (SAL+NS: n=8; SAL+S: n=8; LPS+NS: n=9; LPS+S: n=7).

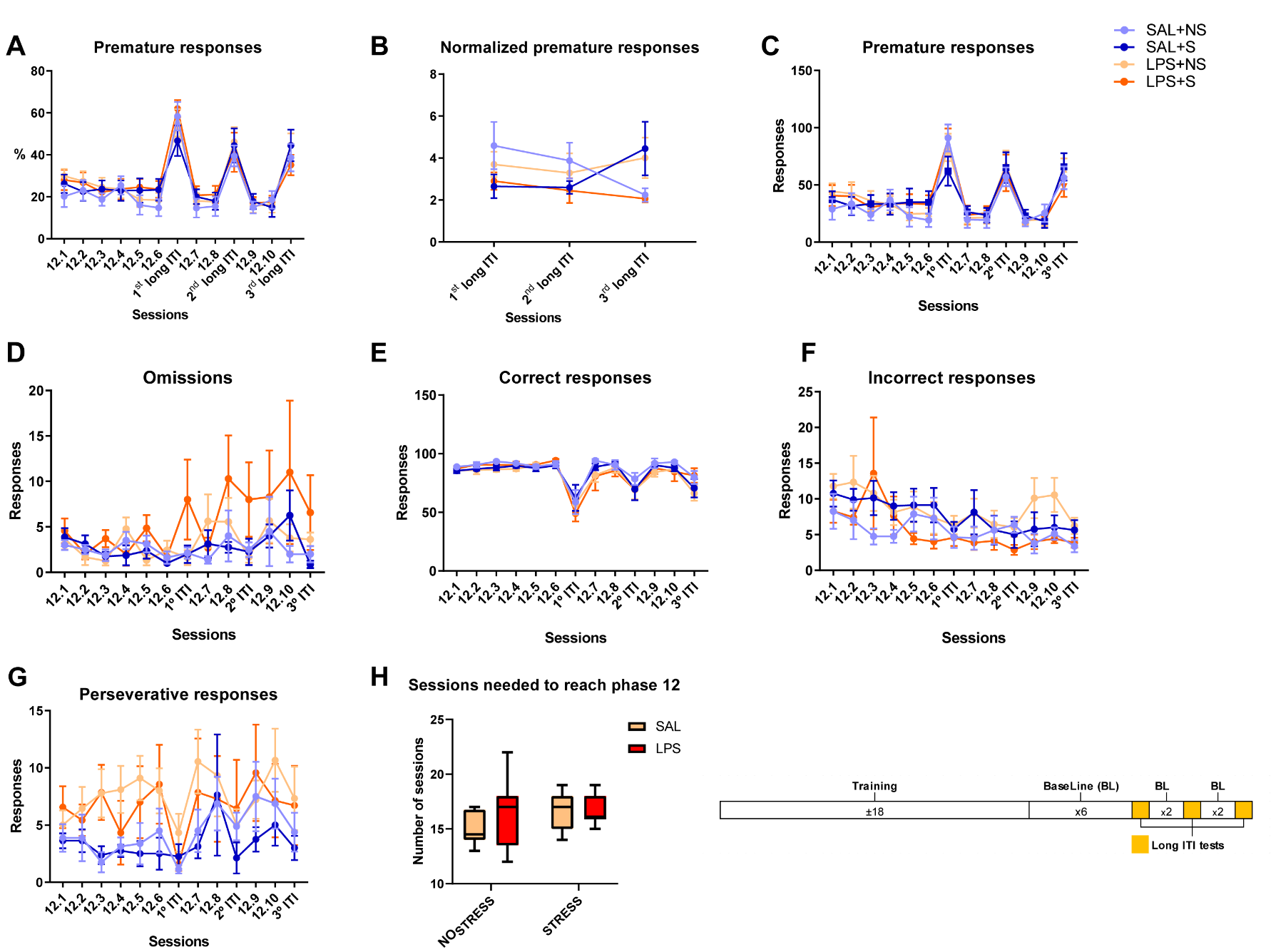

**Figure S5: Percentages of premature responses registered in the different sessions of the 2-CSRTT program.** The figure shows (A) the percentage of premature responses registered during the different sessions of the program [premature responses / (premature responses + correct responses + incorrect responses)], (B) the normalized percentage of premature responses registered during the three long ITI sessions of the program [% premature responses / mean % premature responses of the two previous sessions], (C) the number of premature responses, (D), number of omissions, (E) number of correct responses, (F) number of incorrect responses, (G) number of perseverative responses and (H) number of sessions needed to reach the final phase of training, phase 12, where the baseline was obtained. (SAL+NS: n=8; SAL+S: n=8; LPS+NS: n=9; LPS+S: n=7).

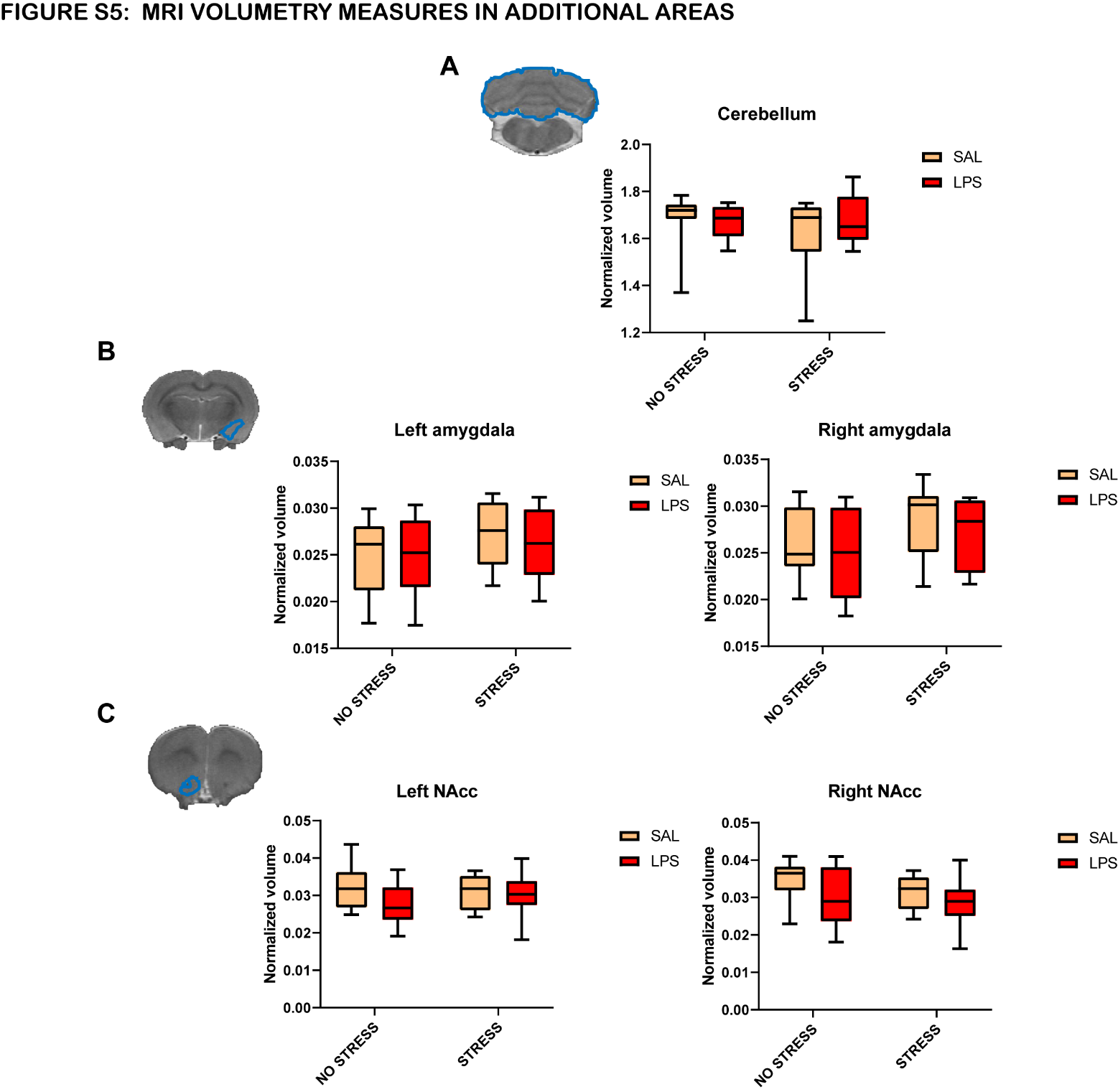

**Figure S6: MRI-assisted volumetry in additional brain areas.** The figure shows: (A) the normalized volume of the cerebellum, (B) the amygdala and (C) the NAcc. No significant effects were observed (SAL+NS: n=8; SAL+S: n=8; LPS+NS: n=8; LPS+S: n=7).

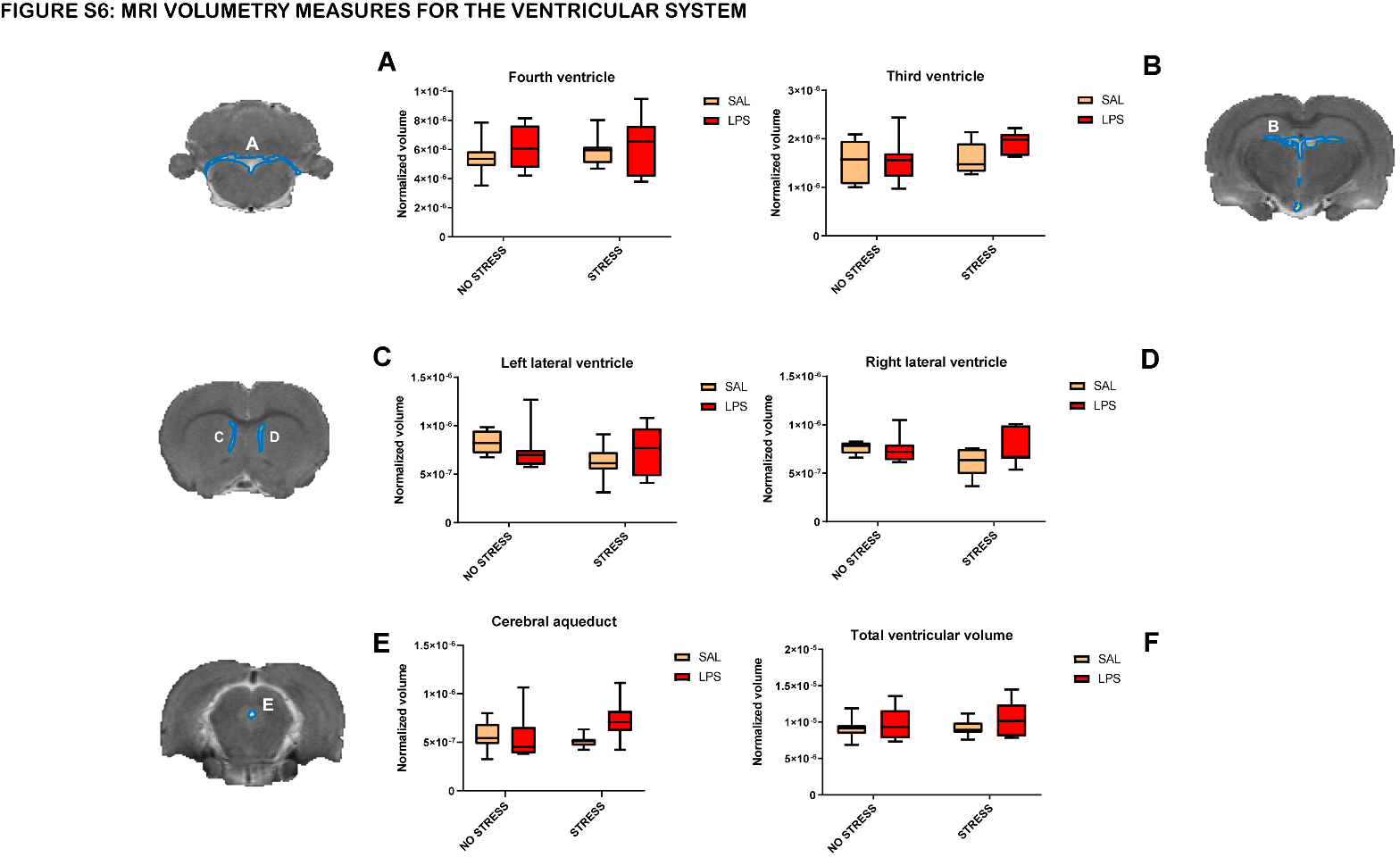
**Figure S7: MRI-assisted volumetry of the ventricular system.** The figure shows: (A) the normalized volume of the IV ventricle, (B) III ventricle and (C) the Aqueduct of Silvius or cerebral aqueduct. No significant effects were observed (SAL+NS: n=8; SAL+S: n=8; LPS+NS: n=8; LPS+S: n=7).

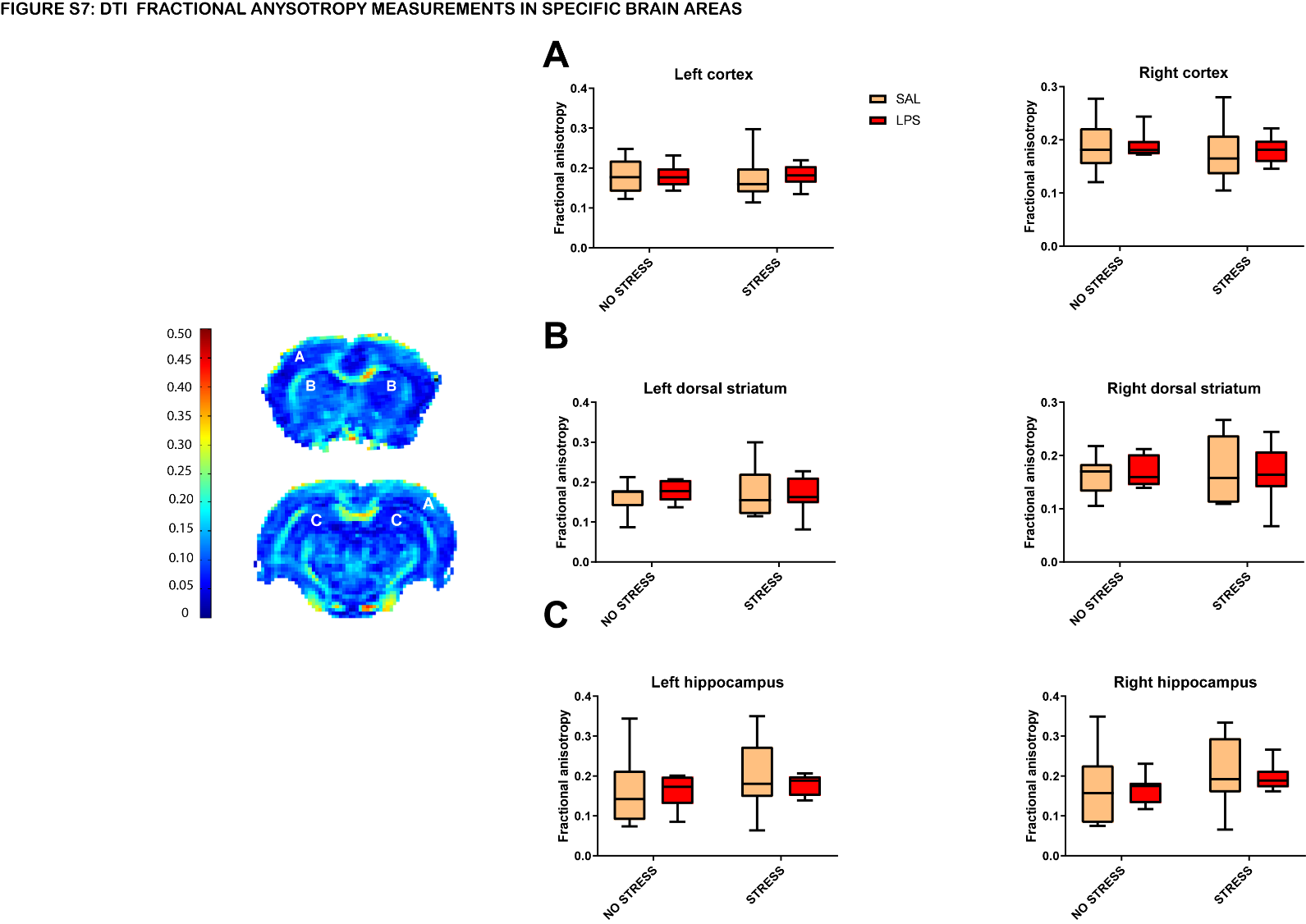

**Figure S8:** DTI FA analysis of specific brain structures. The figure shows the FA value of (A) the cortex, (B) the dorsal striatum and (C) the hippocampus. No significant effects were observed (SAL+NS: n=8; SAL+S: n=8; LPS+NS: n=8; LPS+S: n=7).

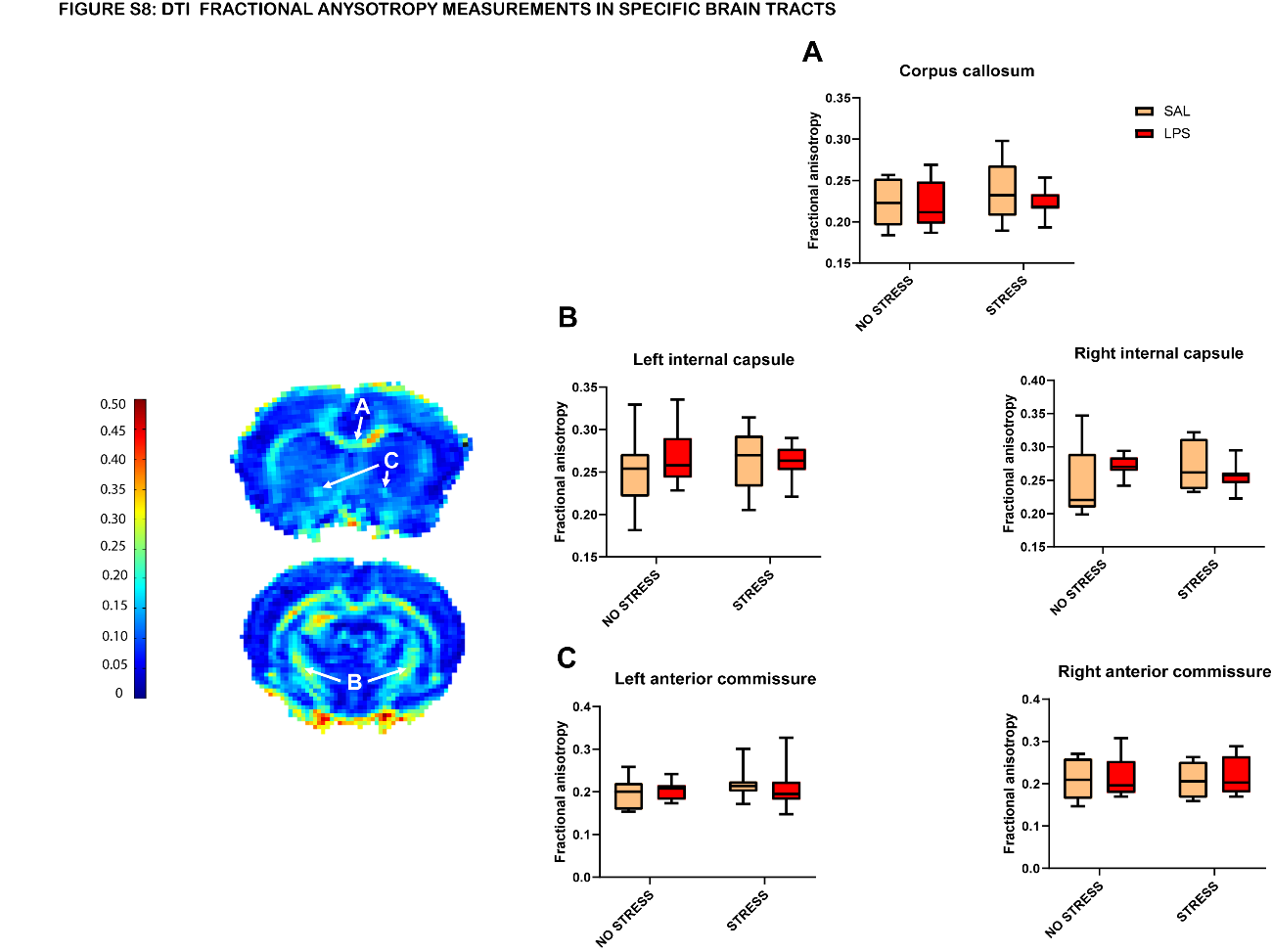

**Figure S9:** DTI FA analysis of specific brain tracts. The figure shows the FA value of (A) the corpus callosum, (B) the internal capsule and (C) the anterior commissure. No significant effects were observed (SAL+NS: n=8; SAL+S: n=8; LPS+NS: n=8; LPS+S: n=7).

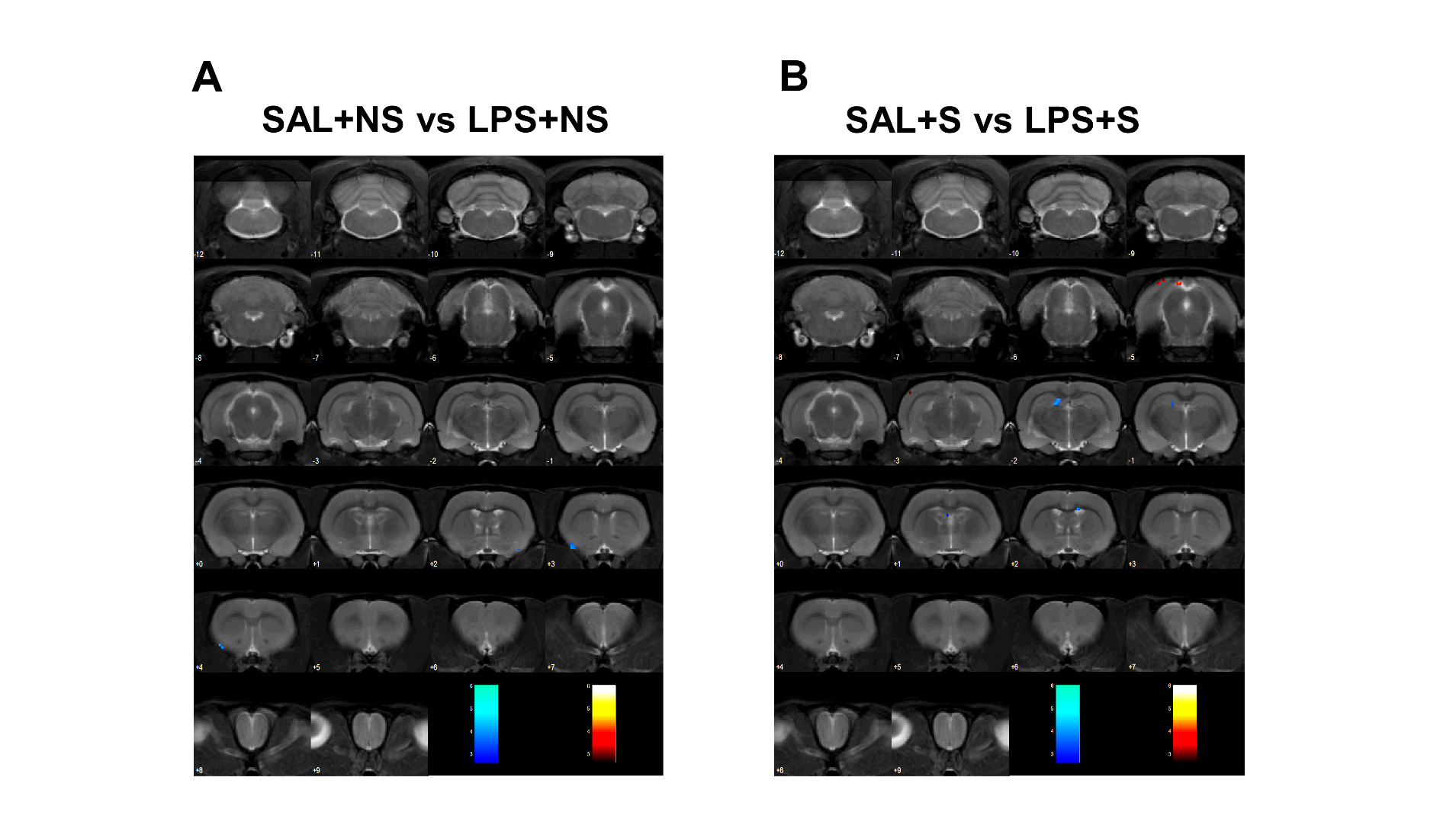
**Figure S10:** SPM analysis of PET data. No significant effects were observed as a result of MIA either in non-stressed or stressed animals (SAL+NS: n=5; SAL+S: n=5; LPS+NS: n=4; LPS+S: n=4). p<0.01 uncorrected, k>50 voxels

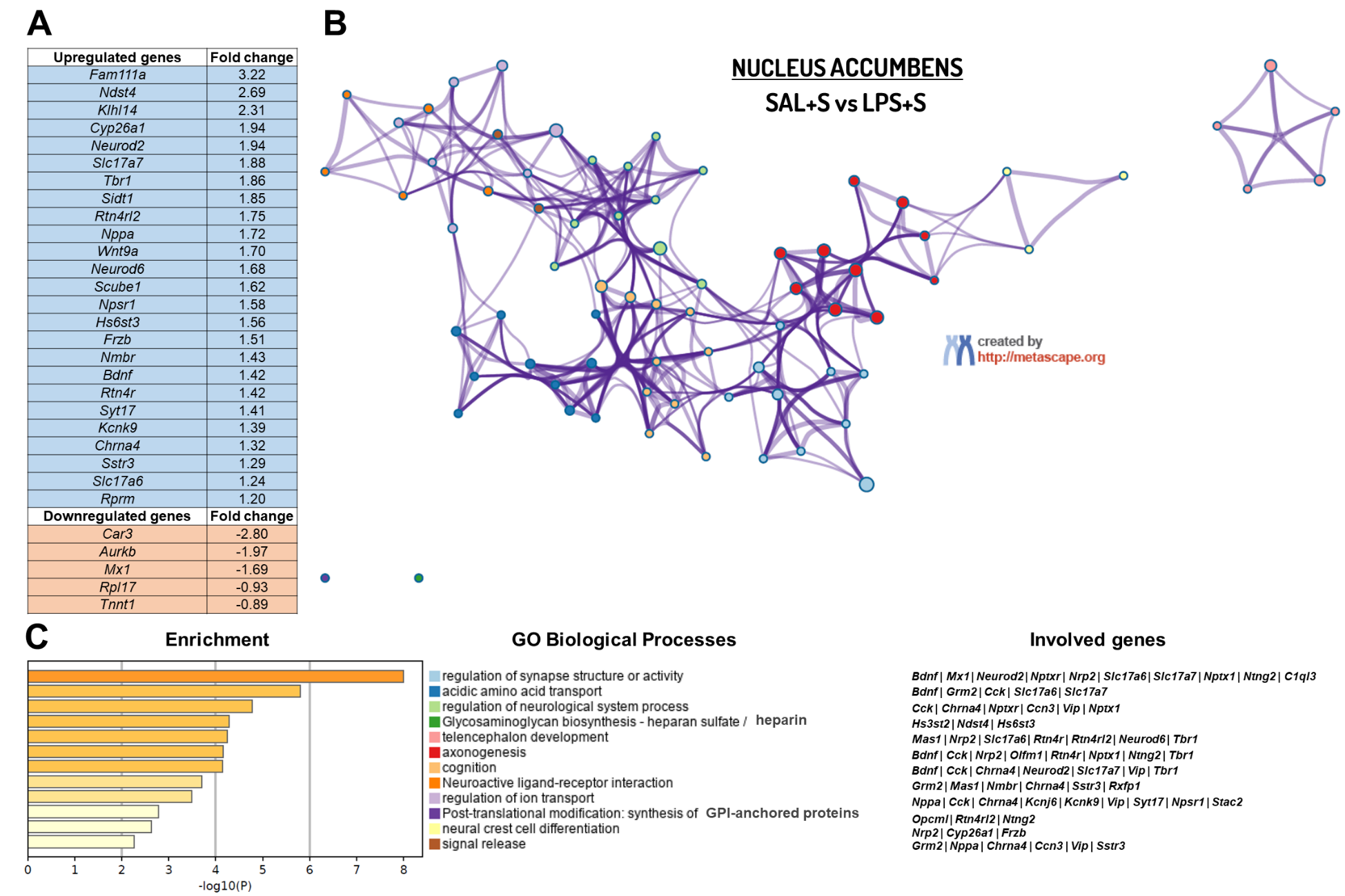

**Figure S11: RNA-Seq analysis in the NAcc for the SAL+S vs LPS+S comparison.** The figure shows (A) main upregulated (Fold change ≥ 1.2) and downregulated (Fold change ≤ -0.5) genes (B) enriched terms network coloured by “GO biological processes” cluster, where nodes that share the same cluster are represented close to each other and (C) enriched terms bar graph coloured by p-value (where terms containing more genes show more significant p-value), GO biological processes and involved genes. (SAL+NS: n=3; LPS+NS: n=3; SAL+S= 3; LPS+S=3).

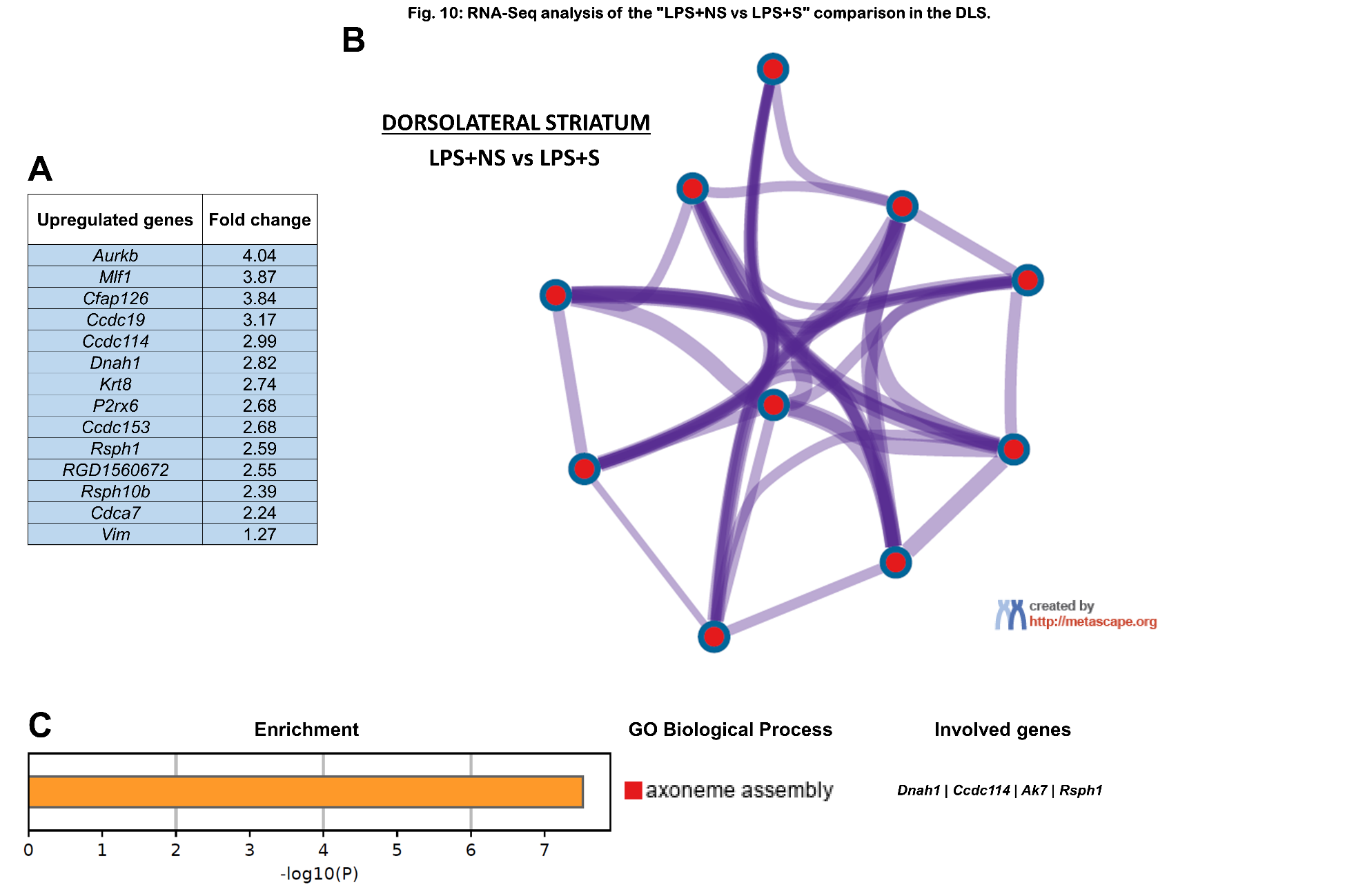

**Figure S12: RNA-Seq analysis in the dorsolateral striatum (DLS) for the SAL+NS vs LPS+S comparison.** The figure shows (A) main upregulated (Fold change ≥ 1.2) and downregulated (Fold change ≤ -0.5) genes (B) enriched terms network coloured by “GO biological processes” cluster, where nodes that share the same cluster are represented close to each other and (C) enriched terms bar graph coloured by p-value (where terms containing more genes show more significant p-value), GO biological processes and involved genes. (SAL+NS: n=3; LPS+NS: n=3; SAL+S= 3; LPS+S=3).

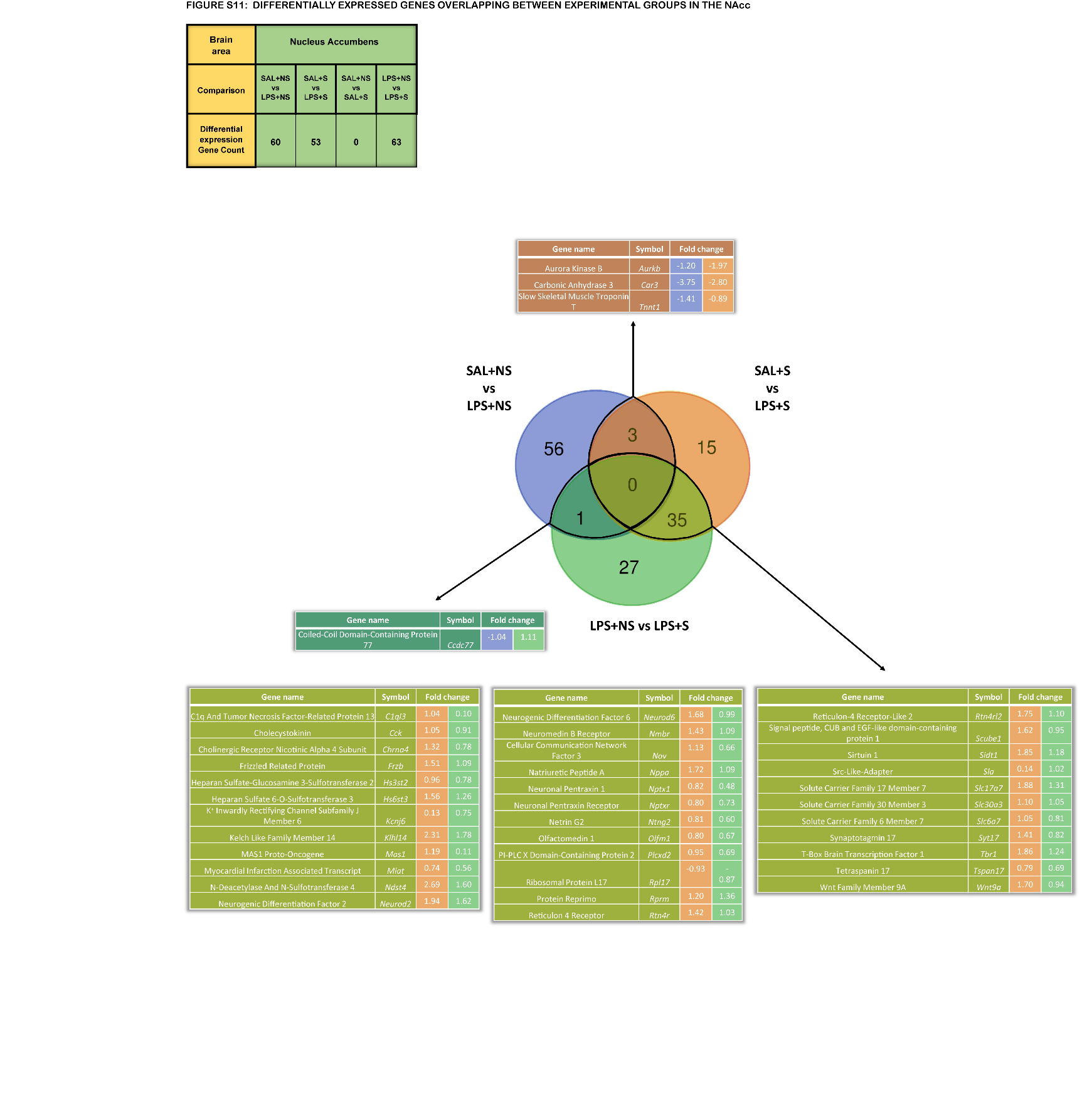
**Figure S13: Number of DEG per condition and DEG overlap between experimental groups in the NAcc.** The figure shows the number of DEG in each of the four groups and the overlap between the different conditions with tables showing the DEG for each overlap with the associated fold-change. Each colour represents the specific overlapping DEG subgroup. (SAL+NS: n=3; LPS+NS: n=3; SAL+S= 3; LPS+S=3).

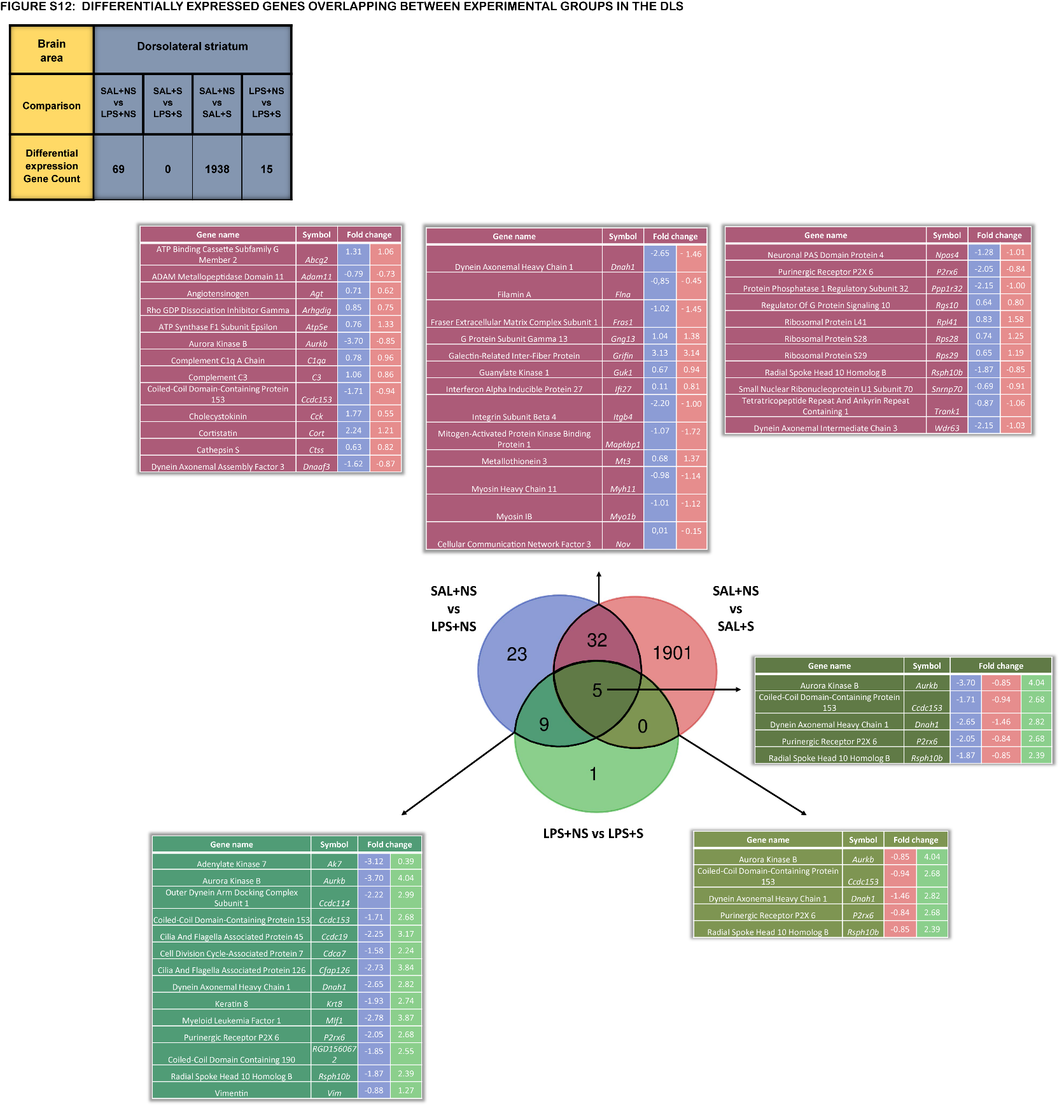
**Figure S14: Number of DEG per condition and DEG overlap between experimental groups in the DLS.** The figure shows the number of DEG in each of the four groups and the overlap between the different conditions with tables showing the DEG for each overlap with the associated fold-change. Each colour represents the specific overlapping DEG subgroup. (SAL+NS: n=3; LPS+NS: n=3; SAL+S= 3; LPS+S=3).

**Table S1: Statistical analysis of PPI and Habituation across conditions.**
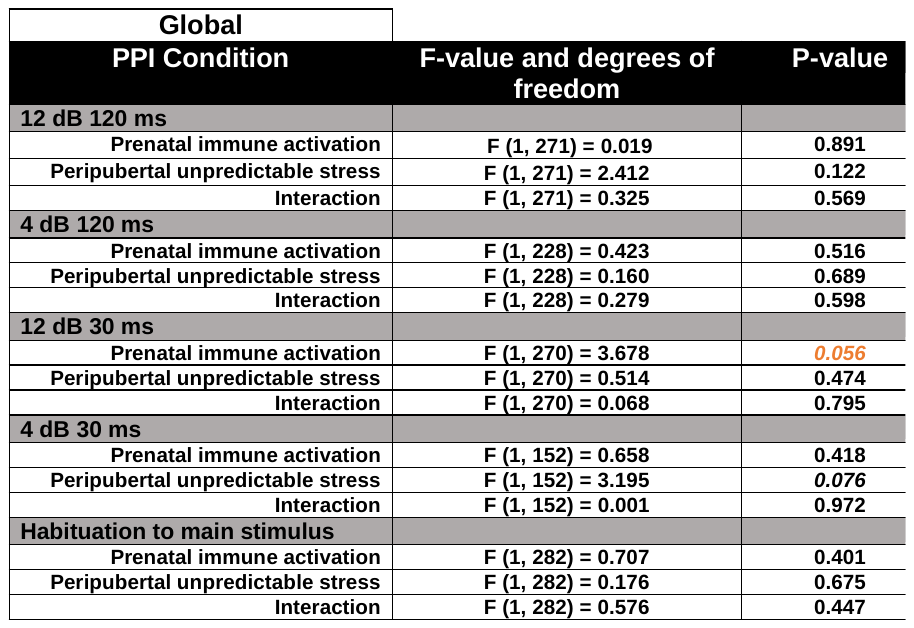

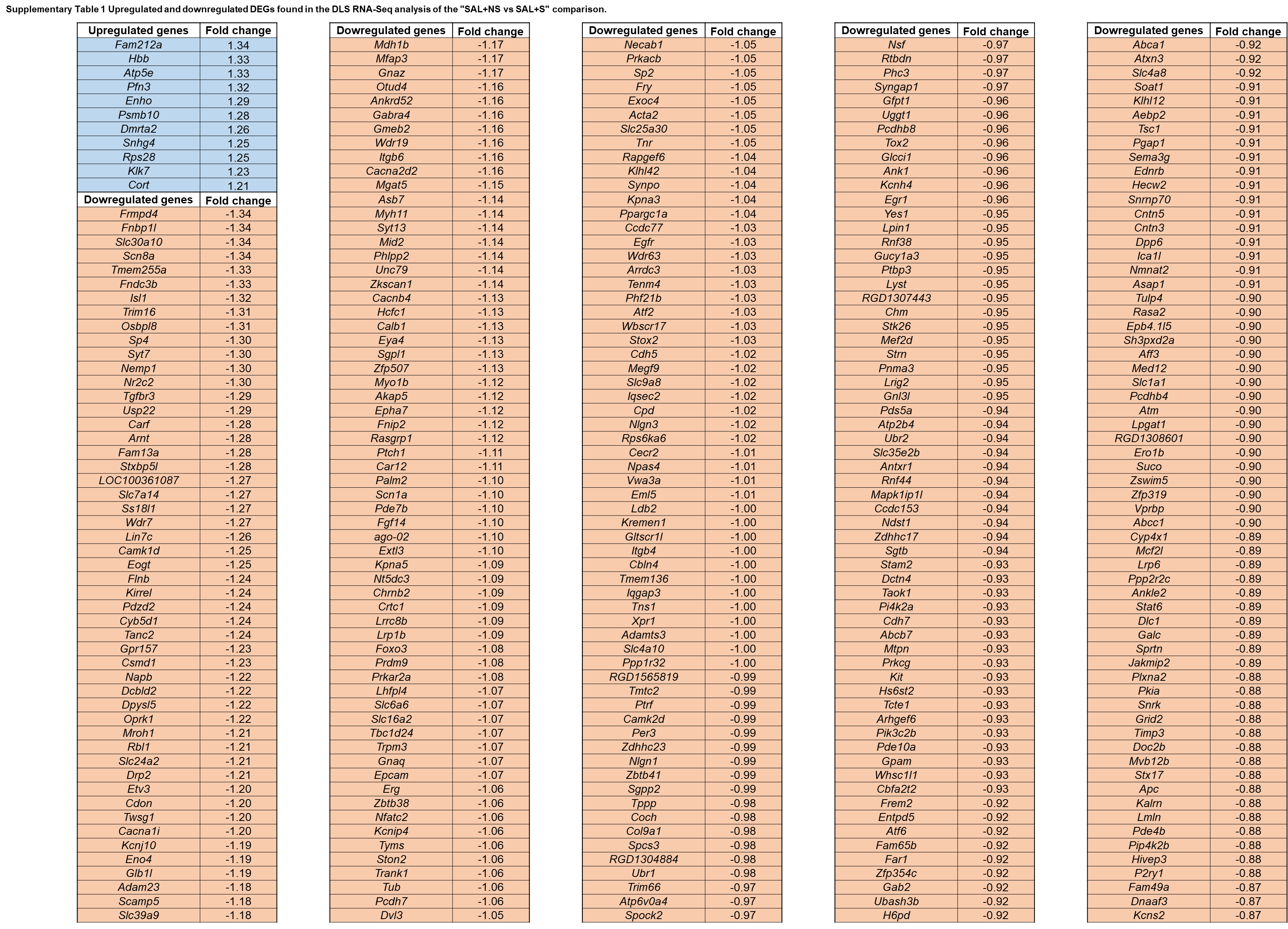

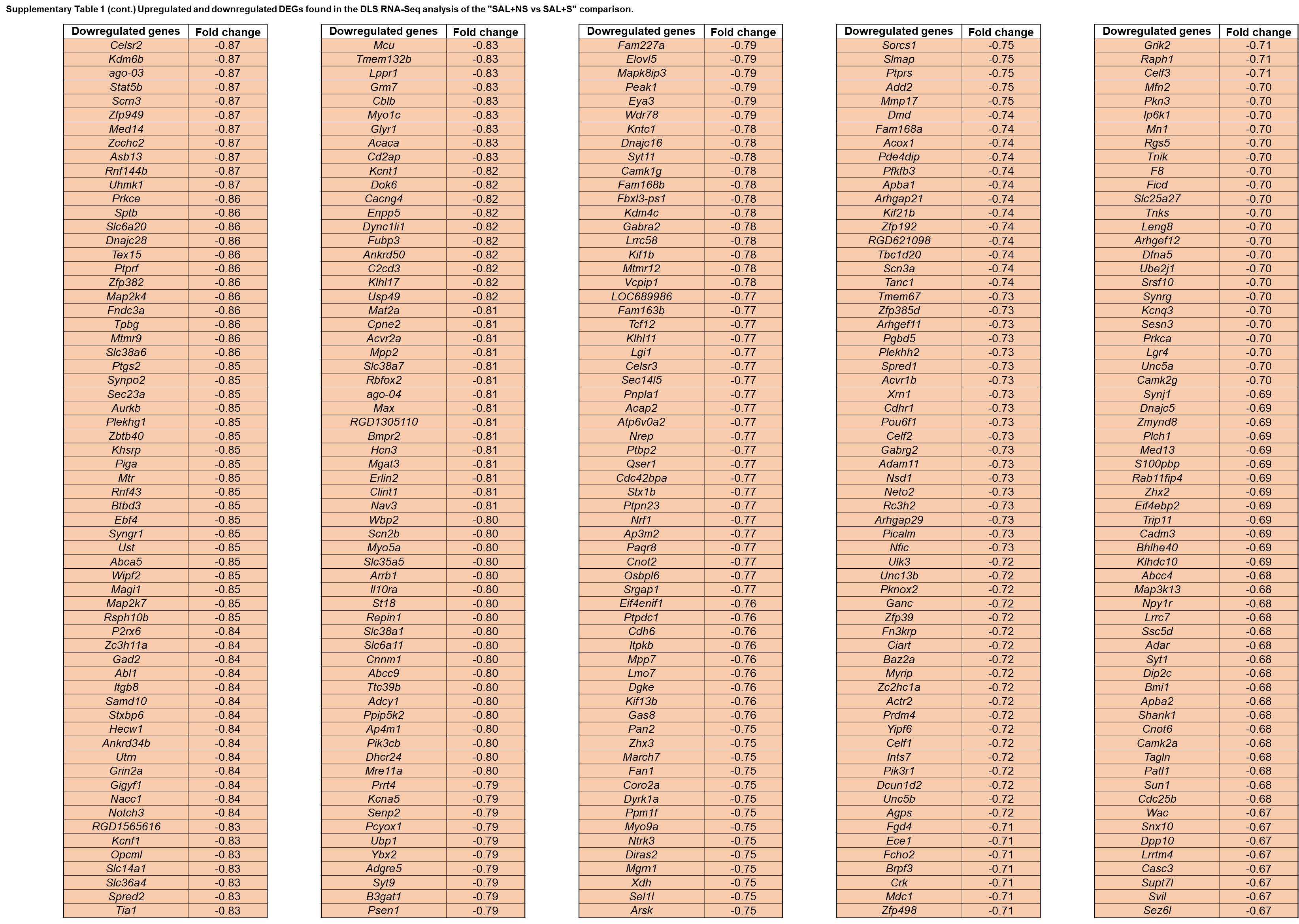

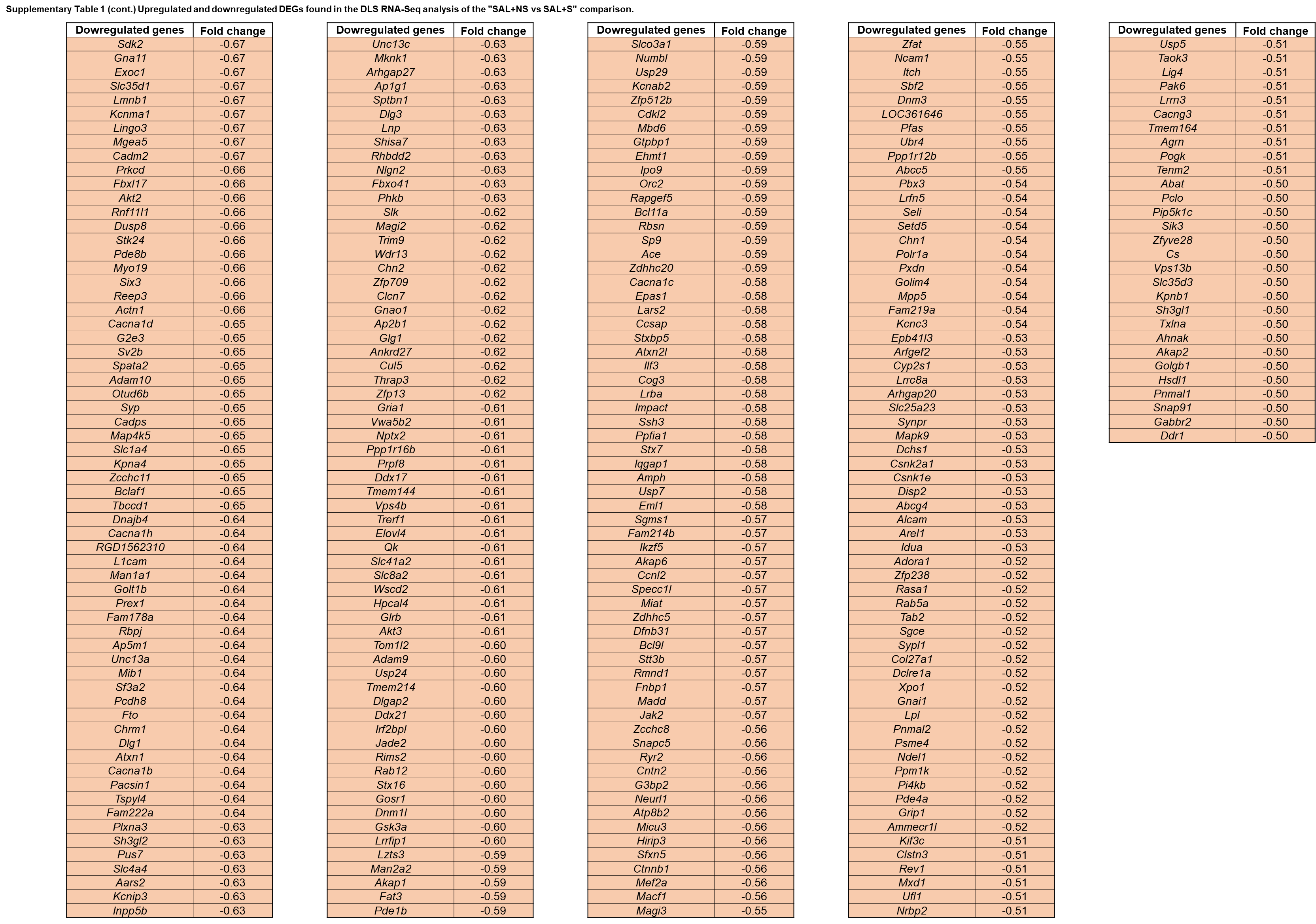

**Table S2: Up-regulated and down-regulated DEGs in the DLS for the SAL+NS vs SAL+S comparison.** For all the genes listed, q<0.05.

| **Term** | **Description** | **LogP** | **Log(q-value)** | **InTerm_InList** | **Symbols** |
| --- | --- | --- | --- | --- | --- |
| GO:0035082 | axoneme assembly | -9,171 | -4,907 | 7/64 | Dnah1,Ccdc114,Ak7,Spag6,Rsph1,Cfap206,Dnaaf3,Ropn1l,Foxj1,Aurkb,Fam107a,Dynlrb2,Tcte1 |
| GO:0035082 | axoneme assembly | -9,171 | -4,907 | 7/64 | Dnah1,Ccdc114,Ak7,Spag6,Rsph1,Cfap206,Dnaaf3 |
| GO:0001578 | microtubule bundle formation | -7,88746 | -3,924 | 7/97 | Dnah1,Ccdc114,Ak7,Spag6,Rsph1,Cfap206,Dnaaf3 |
| GO:0003341 | cilium movement | -5,95728 | -2,170 | 5/63 | Dnah1,Ccdc114,Ak7,Cfap206,Ropn1l |
| GO:0060271 | cilium assembly | -5,27028 | -1,608 | 8/334 | Foxj1,Dnah1,Ccdc114,Ak7,Spag6,Rsph1,Cfap206,Dnaaf3 |
| GO:0044782 | cilium organization | -5,04996 | -1,564 | 8/358 | Foxj1,Dnah1,Ccdc114,Ak7,Spag6,Rsph1,Cfap206,Dnaaf3 |
| GO:0007017 | microtubule-based process | -4,97995 | -1,561 | 11/762 | Aurkb,Dnah1,Ccdc114,Ak7,Spag6,Fam107a,Dynlrb2,Rsph1,Cfap206,Ropn1l,Dnaaf3 |
| GO:0000226 | microtubule cytoskeleton organization | -4,41808 | -1,160 | 9/572 | Aurkb,Dnah1,Ccdc114,Ak7,Spag6,Fam107a,Rsph1,Cfap206,Dnaaf3 |
| GO:0070286 | axonemal dynein complex assembly | -4,08411 | -0,934 | 3/29 | Dnah1,Ccdc114,Dnaaf3 |
| GO:0007018 | microtubule-based movement | -3,87345 | -0,756 | 6/270 | Dnah1,Ccdc114,Ak7,Dynlrb2,Cfap206,Ropn1l |
| GO:0120031 | plasma membrane bounded cell projection assembly | -3,8035 | -0,716 | 8/537 | Foxj1,Dnah1,Ccdc114,Ak7,Spag6,Rsph1,Cfap206,Dnaaf3 |
| GO:0030031 | cell projection assembly | -3,73788 | -0,678 | 8/549 | Foxj1,Dnah1,Ccdc114,Ak7,Spag6,Rsph1,Cfap206,Dnaaf3 |
| GO:0030317 | flagellated sperm motility | -3,58135 | -0,600 | 4/106 | Dnah1,Tcte1,Spag6,Ropn1l |
| GO:0097722 | sperm motility | -3,58135 | -0,600 | 4/106 | Dnah1,Tcte1,Spag6,Ropn1l |
| GO:0007286 | spermatid development | -2,69435 | 0,000 | 4/183 | Dnah1,Spag6,Rsph1,Ropn1l |
| GO:0048515 | spermatid differentiation | -2,6269 | 0,000 | 4/191 | Dnah1,Spag6,Rsph1,Ropn1l |
| GO:0007283 | spermatogenesis | -2,24789 | 0,000 | 6/558 | Foxj1,Dnah1,Ak7,Spag6,Rsph1,Ropn1l |
| GO:0048232 | male gamete generation | -2,17504 | 0,000 | 6/578 | Foxj1,Dnah1,Ak7,Spag6,Rsph1,Ropn1l |
| GO:0007626 | locomotory behaviour | -5,08705 | -1,564 | 7/251 | Drd3,Kcnj10,Scn1a,Calb1,Prex2,Hipk2,Ciart,Grin2a,Nrgn |
| GO:0007626 | locomotory behaviour | -5,08705 | -1,564 | 7/251 | Drd3,Kcnj10,Scn1a,Calb1,Prex2,Hipk2,Ciart |
| GO:0030534 | adult behaviour | -3,64518 | -0,612 | 5/189 | Drd3,Kcnj10,Scn1a,Prex2,Hipk2 |
| GO:0008344 | adult locomotory behaviour | -3,55029 | -0,600 | 4/108 | Kcnj10,Scn1a,Prex2,Hipk2 |
| GO:0007610 | behaviour | -3,54204 | -0,600 | 9/746 | Grin2a,Drd3,Kcnj10,Nrgn,Scn1a,Calb1,Prex2,Hipk2,Ciart |
| GO:0007628 | adult walking behaviour | -3,50929 | -0,588 | 3/45 | Kcnj10,Scn1a,Hipk2 |
| GO:0090659 | walking behaviour | -3,45315 | -0,551 | 3/47 | Kcnj10,Scn1a,Hipk2 |
| GO:1900271 | regulation of long-term synaptic potentiation | -4,76109 | -1,400 | 4/53 | Grin2a,Nrgn,Calb1,Fam107a,Erbb4,Kcnj10,Drd3 |
| GO:1900271 | regulation of long-term synaptic potentiation | -4,76109 | -1,400 | 4/53 | Grin2a,Nrgn,Calb1,Fam107a |
| GO:0050806 | positive regulation of synaptic transmission | -2,68354 | 0,000 | 5/309 | Grin2a,Erbb4,Nrgn,Calb1,Fam107a |
| GO:0048167 | regulation of synaptic plasticity | -2,57701 | 0,000 | 5/327 | Grin2a,Kcnj10,Nrgn,Calb1,Fam107a |
| GO:0060291 | long-term synaptic potentiation | -2,57042 | 0,000 | 4/198 | Grin2a,Nrgn,Calb1,Fam107a |
| GO:0050890 | cognition | -2,44522 | 0,000 | 5/351 | Grin2a,Drd3,Nrgn,Calb1,Fam107a |
| GO:0050879 | multicellular organismal movement | -4,38216 | -1,160 | 4/66 | Drd3,Tnni3,Tnnt1,Hipk2,Scn1a |
| GO:0050879 | multicellular organismal movement | -4,38216 | -1,160 | 4/66 | Drd3,Tnni3,Tnnt1,Hipk2 |
| GO:0050881 | musculoskeletal movement | -4,38216 | -1,160 | 4/66 | Drd3,Tnni3,Tnnt1,Hipk2 |
| GO:0060048 | cardiac muscle contraction | -2,22119 | 0,000 | 3/126 | Tnni3,Scn1a,Tnnt1 |
| GO:0007389 | pattern specification process | -4,19177 | -1,007 | 8/472 | C3,Erbb4,Msx1,Foxj1,Folr1,Zeb2,Hipk2,Dnaaf3 |
| GO:0007389 | pattern specification process | -4,19177 | -1,007 | 8/472 | C3,Erbb4,Msx1,Foxj1,Folr1,Zeb2,Hipk2,Dnaaf3 |
| GO:0001755 | neural crest cell migration | -3,14045 | -0,257 | 3/60 | Erbb4,Folr1,Zeb2 |
| GO:0014032 | neural crest cell development | -2,68798 | 0,000 | 3/86 | Erbb4,Folr1,Zeb2 |
| GO:0014031 | mesenchymal cell development | -2,61798 | 0,000 | 3/91 | Erbb4,Folr1,Zeb2 |
| GO:0048864 | stem cell development | -2,60447 | 0,000 | 3/92 | Erbb4,Folr1,Zeb2 |
| GO:0014033 | neural crest cell differentiation | -2,59113 | 0,000 | 3/93 | Erbb4,Folr1,Zeb2 |
| GO:0048762 | mesenchymal cell differentiation | -2,41362 | 0,000 | 4/219 | Erbb4,Msx1,Folr1,Zeb2 |
| GO:0048863 | stem cell differentiation | -2,37194 | 0,000 | 4/225 | Erbb4,Msx1,Folr1,Zeb2 |
| GO:0003002 | regionalization | -2,3237 | 0,000 | 5/375 | C3,Msx1,Foxj1,Zeb2,Hipk2 |
| GO:0060485 | mesenchyme development | -2,04626 | 0,000 | 4/279 | Erbb4,Msx1,Folr1,Zeb2 |
| GO:0034762 | regulation of transmembrane transport | -2,84404 | 0,000 | 7/579 | C3,Grin2a,Drd3,Kcnj10,Erbb4,Scn1a,Dpp10,Nrgn,Pclo,Lin7b |
| GO:0034762 | regulation of transmembrane transport | -2,84404 | 0,000 | 7/579 | C3,Grin2a,Drd3,Kcnj10,Erbb4,Scn1a,Dpp10 |
| GO:0008306 | associative learning | -2,3965 | 0,000 | 3/109 | Grin2a,Drd3,Nrgn |
| GO:0001505 | regulation of neurotransmitter levels | -2,19322 | 0,000 | 5/403 | Grin2a,Drd3,Kcnj10,Pclo,Lin7b |
| rno04728 | Dopaminergic synapse | -2,16545 | 0,000 | 3/132 | Grin2a,Drd3,Scn1a |
| GO:0050905 | neuromuscular process | -2,13862 | 0,000 | 3/135 | Grin2a,Drd3,Scn1a |
| GO:0007368 | determination of left/right symmetry | -2,24042 | 0,000 | 3/124 | Foxj1,Folr1,Dnaaf3 |
| GO:0007368 | determination of left/right symmetry | -2,24042 | 0,000 | 3/124 | Foxj1,Folr1,Dnaaf3 |
| GO:0009855 | determination of bilateral symmetry | -2,15643 | 0,000 | 3/133 | Foxj1,Folr1,Dnaaf3 |
| GO:0009799 | specification of symmetry | -2,14749 | 0,000 | 3/134 | Foxj1,Folr1,Dnaaf3 |
| GO:0048514 | blood vessel morphogenesis | -2,10192 | 0,000 | 6/599 | C3,Tnni3,Mmp14,Folr1,Hipk2,Itgb8 |
| GO:0048514 | blood vessel morphogenesis | -2,10192 | 0,000 | 6/599 | C3,Tnni3,Mmp14,Folr1,Hipk2,Itgb8 |

**Table S3: Metascape Enrichment metrics and DEG composition of each significant gene ontology in the SAL-NS vs LPS-NS comparison in the NAcc**

| Term | Description | LogP | Log(q-value) | InTerm_InList | Symbols |
| --- | --- | --- | --- | --- | --- |
| GO:0098962 | regulation of postsynaptic neurotransmitter receptor activity | -6,53957 | -2,276 | 4/17 | Chrna4,Nptxr,Nptx1,Nptx2,Agt,Cck,Ccn3,Rph3a,Nppa,Wnt9a,Neurod2,Tpbg,Cabp1,Igsf21,Ntng2,C1ql3,Slc17a7,Chrm3,Nrn1,Cd24 |
| GO:0098962 | regulation of postsynaptic neurotransmitter receptor activity | -6,53957 | -2,276 | 4/17 | Chrna4,Nptxr,Nptx1,Nptx2 |
| GO:0031644 | regulation of neurological system process | -5,60466 | -1,662 | 7/178 | Agt,Cck,Chrna4,Nptxr,Ccn3,Nptx1,Nptx2 |
| GO:0099601 | regulation of neurotransmitter receptor activity | -5,44914 | -1,662 | 5/68 | Chrna4,Nptxr,Rph3a,Nptx1,Nptx2 |
| GO:0010469 | regulation of signaling receptor activity | -4,88649 | -1,225 | 10/536 | Agt,Nppa,Cck,Chrna4,Nptxr,Ccn3,Rph3a,Nptx1,Wnt9a,Nptx2 |
| GO:0050808 | synapse organization | -4,50809 | -1,085 | 9/472 | Neurod2,Nptxr,Tpbg,Rph3a,Cabp1,Nptx1,Igsf21,Ntng2,C1ql3 |
| GO:0050803 | regulation of synapse structure or activity | -4,30713 | -1,063 | 7/282 | Neurod2,Nptxr,Tpbg,Slc17a7,Nptx1,Ntng2,C1ql3 |
| GO:0098698 | postsynaptic specialization assembly | -3,97482 | -0,936 | 3/27 | Nptxr,Nptx1,Ntng2 |
| GO:0099633 | protein localization to postsynaptic specialization membrane | -3,83561 | -0,933 | 3/30 | Nptxr,Nptx1,Nptx2 |
| GO:0099645 | neurotransmitter receptor localization to postsynaptic specialization membrane | -3,83561 | -0,933 | 3/30 | Nptxr,Nptx1,Nptx2 |
| GO:0099068 | postsynapse assembly | -3,75077 | -0,888 | 3/32 | Nptxr,Nptx1,Ntng2 |
| GO:0044057 | regulation of system process | -3,57483 | -0,842 | 9/626 | Agt,Chrm3,Nppa,Cck,Chrna4,Nptxr,Ccn3,Nptx1,Nptx2 |
| GO:0050804 | modulation of chemical synaptic transmission | -3,50908 | -0,830 | 9/639 | Agt,Chrna4,Neurod2,Nptxr,Nrn1,Cabp1,Nptx1,Nptx2,Ntng2 |
| GO:0099177 | regulation of trans-synaptic signaling | -3,50409 | -0,830 | 9/640 | Agt,Chrna4,Neurod2,Nptxr,Nrn1,Cabp1,Nptx1,Nptx2,Ntng2 |
| GO:0050807 | regulation of synapse organization | -3,47723 | -0,822 | 6/272 | Neurod2,Nptxr,Tpbg,Nptx1,Ntng2,C1ql3 |
| GO:0099084 | postsynaptic specialization organization | -3,19889 | -0,675 | 3/49 | Nptxr,Nptx1,Ntng2 |
| GO:0051963 | regulation of synapse assembly | -3,11608 | -0,623 | 4/120 | Nptxr,Tpbg,Nptx1,Ntng2 |
| GO:1903539 | protein localization to postsynaptic membrane | -3,02888 | -0,566 | 3/56 | Nptxr,Nptx1,Nptx2 |
| GO:0097120 | receptor localization to synapse | -2,73047 | -0,324 | 3/71 | Nptxr,Nptx1,Nptx2 |
| GO:0035418 | protein localization to synapse | -2,4933 | -0,188 | 3/86 | Nptxr,Nptx1,Nptx2 |
| GO:0007416 | synapse assembly | -2,4366 | -0,155 | 4/184 | Nptxr,Tpbg,Nptx1,Ntng2 |
| GO:0051668 | localization within membrane | -2,38701 | -0,116 | 4/190 | Cd24,Nptxr,Nptx1,Nptx2 |
| GO:0099072 | regulation of postsynaptic membrane neurotransmitter receptor levels | -2,34632 | -0,087 | 3/97 | Nptxr,Nptx1,Nptx2 |
| GO:0099173 | postsynapse organization | -2,25573 | -0,037 | 4/207 | Nptxr,Rph3a,Nptx1,Ntng2 |
| GO:0050890 | cognition | -4,59693 | -1,085 | 8/351 | Agt,Cck,Chrna4,Neurod2,Slc17a7,Nptx2,Cbr3,Tbr1,Slc6a7,Nr4a2,Slc16a5 |
| GO:0050890 | cognition | -4,59693 | -1,085 | 8/351 | Agt,Cck,Chrna4,Neurod2,Slc17a7,Nptx2,Cbr3,Tbr1 |
| GO:0008306 | associative learning | -4,44638 | -1,085 | 5/109 | Agt,Cck,Neurod2,Nptx2,Tbr1 |
| GO:0007612 | learning | -3,43297 | -0,822 | 5/179 | Agt,Cck,Neurod2,Nptx2,Tbr1 |
| GO:0007611 | learning or memory | -3,14816 | -0,632 | 6/314 | Agt,Cck,Neurod2,Slc17a7,Nptx2,Tbr1 |
| GO:0006865 | amino acid transport | -2,69504 | -0,300 | 4/156 | Agt,Cck,Slc17a7,Slc6a7 |
| GO:0003333 | amino acid transmembrane transport | -2,47912 | -0,184 | 3/87 | Agt,Slc17a7,Slc6a7 |
| GO:0007610 | behaviour | -2,41783 | -0,141 | 8/746 | Agt,Cck,Chrna4,Neurod2,Nr4a2,Slc17a7,Nptx2,Tbr1 |
| GO:0046942 | carboxylic acid transport | -2,26449 | -0,037 | 5/330 | Agt,Cck,Slc17a7,Slc6a7,Slc16a5 |
| GO:0015849 | organic acid transport | -2,25901 | -0,037 | 5/331 | Agt,Cck,Slc17a7,Slc6a7,Slc16a5 |
| GO:1903825 | organic acid transmembrane transport | -2,01425 | 0,000 | 3/128 | Agt,Slc17a7,Slc6a7 |
| GO:1905039 | carboxylic acid transmembrane transport | -2,01425 | 0,000 | 3/128 | Agt,Slc17a7,Slc6a7 |
| GO:0007631 | feeding behaviour | -2,00507 | 0,000 | 3/129 | Agt,Cck,Tbr1 |
| GO:0022038 | corpus callosum development | -4,44613 | -1,085 | 3/19 | Rtn4r,Rtn4rl2,Bhlhe22,Mas1,Nr2f1,Neurod6,Tbr1,Agt,Cd24,Olfm1,Syt17,Spock1,Neurod2,Nr4a2,Frzb |
| GO:0022038 | corpus callosum development | -4,44613 | -1,085 | 3/19 | Rtn4r,Rtn4rl2,Bhlhe22 |
| GO:0021537 | telencephalon development | -3,8398 | -0,933 | 7/335 | Mas1,Nr2f1,Rtn4r,Rtn4rl2,Bhlhe22,Neurod6,Tbr1 |
| GO:0010975 | regulation of neuron projection development | -3,63704 | -0,873 | 9/614 | Agt,Cd24,Nr2f1,Olfm1,Rtn4r,Syt17,Spock1,Rtn4rl2,Tbr1 |
| GO:0051961 | negative regulation of nervous system development | -3,60434 | -0,859 | 7/366 | Cd24,Neurod2,Nr2f1,Olfm1,Rtn4r,Spock1,Rtn4rl2 |
| GO:0030900 | forebrain development | -3,55364 | -0,834 | 8/496 | Mas1,Nr4a2,Nr2f1,Rtn4r,Rtn4rl2,Bhlhe22,Neurod6,Tbr1 |
| GO:0045664 | regulation of neuron differentiation | -3,47072 | -0,822 | 10/793 | Agt,Cd24,Neurod2,Nr2f1,Olfm1,Rtn4r,Syt17,Spock1,Rtn4rl2,Tbr1 |
| GO:0010721 | negative regulation of cell development | -3,39113 | -0,795 | 7/397 | Cd24,Nr2f1,Olfm1,Rtn4r,Frzb,Spock1,Rtn4rl2 |
| GO:0050768 | negative regulation of neurogenesis | -2,95606 | -0,512 | 6/342 | Cd24,Nr2f1,Olfm1,Rtn4r,Spock1,Rtn4rl2 |
| GO:0021543 | pallium development | -2,90781 | -0,470 | 5/234 | Mas1,Nr2f1,Bhlhe22,Neurod6,Tbr1 |
| GO:0045665 | negative regulation of neuron differentiation | -2,66348 | -0,286 | 5/266 | Cd24,Nr2f1,Rtn4r,Spock1,Rtn4rl2 |
| GO:0010977 | negative regulation of neuron projection development | -2,46208 | -0,171 | 4/181 | Nr2f1,Rtn4r,Spock1,Rtn4rl2 |
| GO:0031345 | negative regulation of cell projection organization | -2,19117 | 0,000 | 4/216 | Nr2f1,Rtn4r,Spock1,Rtn4rl2 |
| R-RNO-1296041 | Activation of G protein gated Potassium channels | -4,24812 | -1,063 | 3/22 | Kcnj6,Kcnj3,Kcnj9,Pde1a,Chrna4,Slc17a7,Agt,Chrm3,Neurod2,Nppa |
| R-RNO-1296041 | Activation of G protein gated Potassium channels | -4,24812 | -1,063 | 3/22 | Kcnj6,Kcnj3,Kcnj9 |
| R-RNO-1296059 | G protein gated Potassium channels | -4,24812 | -1,063 | 3/22 | Kcnj6,Kcnj3,Kcnj9 |
| R-RNO-997272 | Inhibition of voltage gated Ca2+ channels via Gbeta/gamma subunits | -4,24812 | -1,063 | 3/22 | Kcnj6,Kcnj3,Kcnj9 |
| R-RNO-1296065 | Inwardly rectifying K+ channels | -3,97482 | -0,936 | 3/27 | Kcnj6,Kcnj3,Kcnj9 |
| R-RNO-977444 | GABA B receptor activation | -3,63353 | -0,873 | 3/35 | Kcnj6,Kcnj3,Kcnj9 |
| R-RNO-991365 | Activation of GABAB receptors | -3,63353 | -0,873 | 3/35 | Kcnj6,Kcnj3,Kcnj9 |
| rno05032 | Morphine addiction | -3,53538 | -0,830 | 4/93 | Kcnj6,Kcnj3,Pde1a,Kcnj9 |
| GO:1990573 | potassium ion import across plasma membrane | -3,45998 | -0,822 | 3/40 | Kcnj6,Kcnj3,Kcnj9 |
| R-RNO-112315 | Transmission across Chemical Synapses | -3,37816 | -0,795 | 5/184 | Chrna4,Kcnj6,Kcnj3,Kcnj9,Slc17a7 |
| rno04723 | Retrograde endocannabinoid signaling | -3,35035 | -0,777 | 4/104 | Kcnj6,Kcnj3,Kcnj9,Slc17a7 |
| GO:0010107 | potassium ion import | -3,30805 | -0,743 | 3/45 | Kcnj6,Kcnj3,Kcnj9 |
| R-RNO-977443 | GABA receptor activation | -3,25224 | -0,696 | 3/47 | Kcnj6,Kcnj3,Kcnj9 |
| GO:0098739 | import across plasma membrane | -3,24333 | -0,695 | 4/111 | Agt,Kcnj6,Kcnj3,Kcnj9 |
| rno04725 | Cholinergic synapse | -3,1997 | -0,675 | 4/114 | Chrm3,Chrna4,Kcnj6,Kcnj3 |
| R-RNO-112314 | Neurotransmitter receptors and postsynaptic signal transmission | -3,01144 | -0,560 | 4/128 | Chrna4,Kcnj6,Kcnj3,Kcnj9 |
| GO:0071242 | cellular response to ammonium ion | -2,62965 | -0,263 | 3/77 | Chrm3,Chrna4,Kcnj6 |
| GO:0051602 | response to electrical stimulus | -2,59791 | -0,256 | 3/79 | Kcnj6,Kcnj3,Neurod2 |
| GO:0098659 | inorganic cation import across plasma membrane | -2,58236 | -0,253 | 3/80 | Kcnj6,Kcnj3,Kcnj9 |
| GO:0099587 | inorganic ion import across plasma membrane | -2,58236 | -0,253 | 3/80 | Kcnj6,Kcnj3,Kcnj9 |
| R-RNO-1296071 | Potassium Channels | -2,47912 | -0,184 | 3/87 | Kcnj6,Kcnj3,Kcnj9 |
| R-RNO-112316 | Neuronal System | -2,45747 | -0,171 | 5/297 | Chrna4,Kcnj6,Kcnj3,Kcnj9,Slc17a7 |
| rno04915 | Estrogen signaling pathway | -2,37164 | -0,108 | 3/95 | Kcnj6,Kcnj3,Kcnj9 |
| rno04713 | Circadian entrainment | -2,32155 | -0,070 | 3/99 | Kcnj6,Kcnj3,Kcnj9 |
| GO:0071804 | cellular potassium ion transport | -2,21947 | -0,012 | 4/212 | Nppa,Kcnj6,Kcnj3,Kcnj9 |
| GO:0071805 | potassium ion transmembrane transport | -2,21947 | -0,012 | 4/212 | Nppa,Kcnj6,Kcnj3,Kcnj9 |
| rno04726 | Serotonergic synapse | -2,03283 | 0,000 | 3/126 | Kcnj6,Kcnj3,Kcnj9 |
| GO:0006813 | potassium ion transport | -2,00879 | 0,000 | 4/244 | Nppa,Kcnj6,Kcnj3,Kcnj9 |
| rno00534 | Glycosaminoglycan biosynthesis - heparan sulfate / heparin | -4,0771 | -0,936 | 3/25 | Hs3st2,Ndst4,Hs6st3 |
| rno00534 | Glycosaminoglycan biosynthesis - heparan sulfate / heparin | -4,0771 | -0,936 | 3/25 | Hs3st2,Ndst4,Hs6st3 |
| GO:0015012 | heparan sulfate proteoglycan biosynthetic process | -3,8803 | -0,933 | 3/29 | Hs3st2,Ndst4,Hs6st3 |
| GO:0030201 | heparan sulfate proteoglycan metabolic process | -3,67138 | -0,873 | 3/34 | Hs3st2,Ndst4,Hs6st3 |
| GO:0030166 | proteoglycan biosynthetic process | -3,09881 | -0,620 | 3/53 | Hs3st2,Ndst4,Hs6st3 |
| GO:0006029 | proteoglycan metabolic process | -2,61367 | -0,263 | 3/78 | Hs3st2,Ndst4,Hs6st3 |
| GO:0048812 | neuron projection morphogenesis | -4,02217 | -0,936 | 10/678 | Cck,Nr4a2,Nrn1,Olfm1,Rtn4r,Syt17,Nptx1,Ntng2,Bhlhe22,Tbr1,Spock1 |
| GO:0048812 | neuron projection morphogenesis | -4,02217 | -0,936 | 10/678 | Cck,Nr4a2,Nrn1,Olfm1,Rtn4r,Syt17,Nptx1,Ntng2,Bhlhe22,Tbr1 |
| GO:0120039 | plasma membrane bounded cell projection morphogenesis | -3,94386 | -0,936 | 10/693 | Cck,Nr4a2,Nrn1,Olfm1,Rtn4r,Syt17,Nptx1,Ntng2,Bhlhe22,Tbr1 |
| GO:0048858 | cell projection morphogenesis | -3,92333 | -0,936 | 10/697 | Cck,Nr4a2,Nrn1,Olfm1,Rtn4r,Syt17,Nptx1,Ntng2,Bhlhe22,Tbr1 |
| GO:0032990 | cell part morphogenesis | -3,81297 | -0,929 | 10/719 | Cck,Nr4a2,Nrn1,Olfm1,Rtn4r,Syt17,Nptx1,Ntng2,Bhlhe22,Tbr1 |
| GO:0007409 | axonogenesis | -3,73715 | -0,888 | 8/466 | Cck,Nr4a2,Olfm1,Rtn4r,Nptx1,Ntng2,Bhlhe22,Tbr1 |
| GO:0061564 | axon development | -3,4442 | -0,822 | 8/515 | Cck,Nr4a2,Olfm1,Rtn4r,Nptx1,Ntng2,Bhlhe22,Tbr1 |
| GO:0048667 | cell morphogenesis involved in neuron differentiation | -2,89968 | -0,468 | 8/624 | Cck,Nr4a2,Olfm1,Rtn4r,Nptx1,Ntng2,Bhlhe22,Tbr1 |
| GO:0000904 | cell morphogenesis involved in differentiation | -2,24309 | -0,028 | 8/798 | Cck,Nr4a2,Olfm1,Rtn4r,Nptx1,Ntng2,Bhlhe22,Tbr1 |
| GO:0021953 | central nervous system neuron differentiation | -2,12301 | 0,000 | 4/226 | Nr4a2,Spock1,Bhlhe22,Tbr1 |
| GO:0006836 | neurotransmitter transport | -3,92119 | -0,936 | 7/325 | Nppa,Chrna4,Nrn1,Slc17a7,Slc6a7,Rph3a,Syt17,Agt,Cck,Cd24,Ccn3 |
| GO:0006836 | neurotransmitter transport | -3,92119 | -0,936 | 7/325 | Nppa,Chrna4,Nrn1,Slc17a7,Slc6a7,Rph3a,Syt17 |
| GO:0051954 | positive regulation of amine transport | -3,12305 | -0,623 | 3/52 | Agt,Cck,Chrna4 |
| GO:0051952 | regulation of amine transport | -3,07597 | -0,604 | 4/123 | Agt,Cck,Chrna4,Syt17 |
| GO:0015837 | amine transport | -3,02412 | -0,566 | 4/127 | Agt,Cck,Chrna4,Syt17 |
| GO:0099504 | synaptic vesicle cycle | -2,8275 | -0,402 | 5/244 | Cd24,Chrna4,Nrn1,Slc17a7,Syt17 |
| GO:0048488 | synaptic vesicle endocytosis | -2,74817 | -0,335 | 3/70 | Cd24,Slc17a7,Syt17 |
| GO:0140238 | presynaptic endocytosis | -2,74817 | -0,335 | 3/70 | Cd24,Slc17a7,Syt17 |
| GO:0050433 | regulation of catecholamine secretion | -2,67896 | -0,295 | 3/74 | Agt,Chrna4,Syt17 |
| GO:0099003 | vesicle-mediated transport in synapse | -2,67773 | -0,295 | 5/264 | Cd24,Chrna4,Nrn1,Slc17a7,Syt17 |
| GO:0050432 | catecholamine secretion | -2,61367 | -0,263 | 3/78 | Agt,Chrna4,Syt17 |
| GO:0001505 | regulation of neurotransmitter levels | -2,59557 | -0,256 | 6/403 | Agt,Chrna4,Nrn1,Slc17a7,Rph3a,Syt17 |
| GO:0036465 | synaptic vesicle recycling | -2,56703 | -0,248 | 3/81 | Cd24,Slc17a7,Syt17 |
| GO:0023061 | signal release | -2,54899 | -0,234 | 7/555 | Agt,Nppa,Chrna4,Ccn3,Nrn1,Rph3a,Syt17 |
| GO:0051937 | catecholamine transport | -2,38453 | -0,116 | 3/94 | Agt,Chrna4,Syt17 |
| GO:0007269 | neurotransmitter secretion | -2,27052 | -0,037 | 4/205 | Chrna4,Nrn1,Rph3a,Syt17 |
| GO:0099643 | signal release from synapse | -2,26311 | -0,037 | 4/206 | Chrna4,Nrn1,Rph3a,Syt17 |
| GO:1903305 | regulation of regulated secretory pathway | -2,25573 | -0,037 | 4/207 | Nppa,Chrna4,Nrn1,Syt17 |
| GO:0015844 | monoamine transport | -2,1945 | 0,000 | 3/110 | Agt,Chrna4,Syt17 |
| GO:2001023 | regulation of response to drug | -2,09063 | 0,000 | 3/120 | Agt,Chrna4,Syt17 |
| GO:0015893 | drug transport | -2,04573 | 0,000 | 4/238 | Agt,Chrna4,Slc17a7,Syt17 |
| GO:0043269 | regulation of ion transport | -3,65976 | -0,873 | 10/751 | Agt,Nppa,Cck,Chrna4,Kcnj6,Kcnj3,Kcnj9,Rph3a,Cabp1,Syt17 |
| GO:0043269 | regulation of ion transport | -3,65976 | -0,873 | 10/751 | Agt,Nppa,Cck,Chrna4,Kcnj6,Kcnj3,Kcnj9,Rph3a,Cabp1,Syt17 |
| GO:0034765 | regulation of ion transmembrane transport | -3,63148 | -0,873 | 8/483 | Agt,Nppa,Chrna4,Kcnj6,Kcnj3,Kcnj9,Rph3a,Cabp1 |
| GO:0034762 | regulation of transmembrane transport | -3,10896 | -0,623 | 8/579 | Agt,Nppa,Chrna4,Kcnj6,Kcnj3,Kcnj9,Rph3a,Cabp1 |
| GO:2001257 | regulation of cation channel activity | -2,51453 | -0,205 | 4/175 | Nppa,Chrna4,Rph3a,Cabp1 |
| GO:1904062 | regulation of cation transmembrane transport | -2,1332 | 0,000 | 5/355 | Agt,Nppa,Chrna4,Rph3a,Cabp1 |
| GO:0048588 | developmental cell growth | -3,50295 | -0,830 | 6/269 | Agt,Nppa,Nrn1,Olfm1,Rtn4r,Syt17,Ccn3,Frzb |
| GO:0048588 | developmental cell growth | -3,50295 | -0,830 | 6/269 | Agt,Nppa,Nrn1,Olfm1,Rtn4r,Syt17 |
| GO:0016049 | cell growth | -3,43857 | -0,822 | 8/516 | Agt,Nppa,Ccn3,Nrn1,Olfm1,Rtn4r,Syt17,Frzb |
| GO:0030308 | negative regulation of cell growth | -3,21335 | -0,675 | 5/200 | Agt,Nppa,Ccn3,Rtn4r,Frzb |
| GO:0001558 | regulation of cell growth | -3,13185 | -0,624 | 7/439 | Agt,Nppa,Ccn3,Olfm1,Rtn4r,Syt17,Frzb |
| GO:0045926 | negative regulation of growth | -2,64936 | -0,277 | 5/268 | Agt,Nppa,Ccn3,Rtn4r,Frzb |
| GO:1990138 | neuron projection extension | -2,33138 | -0,076 | 4/197 | Nrn1,Olfm1,Rtn4r,Syt17 |
| GO:0040008 | regulation of growth | -2,00758 | 0,000 | 7/700 | Agt,Nppa,Ccn3,Olfm1,Rtn4r,Syt17,Frzb |
| GO:0001764 | neuron migration | -3,37816 | -0,795 | 5/184 | Cck,Nr4a2,Nr2f1,Olfm1,Spock1,Neurod2 |
| GO:0001764 | neuron migration | -3,37816 | -0,795 | 5/184 | Cck,Nr4a2,Nr2f1,Olfm1,Spock1 |
| GO:0033555 | multicellular organismal response to stress | -2,26197 | -0,037 | 3/104 | Cck,Neurod2,Nr4a2 |
| GO:0019233 | sensory perception of pain | -2,71546 | -0,315 | 4/154 | Cck,Chrna4,Ephx2,Ccn3 |
| GO:0019233 | sensory perception of pain | -2,71546 | -0,315 | 4/154 | Cck,Chrna4,Ephx2,Ccn3 |
| GO:0003407 | neural retina development | -2,61367 | -0,263 | 3/78 | Slc17a7,Bhlhe22,Miat |
| GO:0003407 | neural retina development | -2,61367 | -0,263 | 3/78 | Slc17a7,Bhlhe22,Miat |
| R-RNO-163125 | Post-translational modification: synthesis of GPI-anchored proteins | -2,56703 | -0,248 | 3/81 | Nrn1,Rtn4rl2,Ntng2 |
| R-RNO-163125 | Post-translational modification: synthesis of GPI-anchored proteins | -2,56703 | -0,248 | 3/81 | Nrn1,Rtn4rl2,Ntng2 |
| GO:0032101 | regulation of response to external stimulus | -2,29517 | -0,048 | 8/782 | Agt,Nppa,Cd24,Mas1,Cck,Ccn3,Rtn4r,Tbr1,Basp1 |
| GO:0032101 | regulation of response to external stimulus | -2,29517 | -0,048 | 8/782 | Agt,Nppa,Cd24,Mas1,Cck,Ccn3,Rtn4r,Tbr1 |
| GO:1901222 | regulation of NIK/NF-kappaB signaling | -2,11062 | 0,000 | 3/118 | Agt,Mas1,Ccn3 |
| GO:0050727 | regulation of inflammatory response | -2,09845 | 0,000 | 5/362 | Agt,Nppa,Cd24,Mas1,Ccn3 |
| GO:0072009 | nephron epithelium development | -2,07101 | 0,000 | 3/122 | Agt,Cd24,Basp1 |
| GO:0038061 | NIK/NF-kappaB signaling | -2,03283 | 0,000 | 3/126 | Agt,Mas1,Ccn3 |
| GO:0008637 | apoptotic mitochondrial changes | -2,16228 | 0,000 | 3/113 | Cd24,Cck,Nptx1 |
| GO:0008637 | apoptotic mitochondrial changes | -2,16228 | 0,000 | 3/113 | Cd24,Cck,Nptx1 |

**Table S4: Metascape Enrichment metrics and DEG composition of each significant gene ontology in the LPS-NS vs LPS-S comparison in the NAcc**

| Term | Description | LogP | Log(q-value) | InTerm_InList | Symbols |
| --- | --- | --- | --- | --- | --- |
| GO:0050803 | regulation of synapse structure or activity | -6,93158 | -2,668 | 9/282 | Bdnf,Neurod2,Nptxr,Nrp2,Slc17a6,Slc17a7,Nptx1,Ntng2,C1ql3,Chrna4,Grm2 |
| GO:0050803 | regulation of synapse structure or activity | -6,93158 | -2,668 | 9/282 | Bdnf,Neurod2,Nptxr,Nrp2,Slc17a6,Slc17a7,Nptx1,Ntng2,C1ql3 |
| GO:0050807 | regulation of synapse organization | -4,86032 | -1,385 | 7/272 | Bdnf,Neurod2,Nptxr,Nrp2,Nptx1,Ntng2,C1ql3 |
| GO:0098962 | regulation of postsynaptic neurotransmitter receptor activity | -4,80357 | -1,385 | 3/17 | Chrna4,Nptxr,Nptx1 |
| GO:0098698 | postsynaptic specialization assembly | -4,17896 | -1,036 | 3/27 | Nptxr,Nptx1,Ntng2 |
| GO:0099068 | postsynapse assembly | -3,9541 | -0,981 | 3/32 | Nptxr,Nptx1,Ntng2 |
| GO:0099084 | postsynaptic specialization organization | -3,39948 | -0,713 | 3/49 | Nptxr,Nptx1,Ntng2 |
| GO:0051963 | regulation of synapse assembly | -3,37605 | -0,712 | 4/120 | Bdnf,Nptxr,Nptx1,Ntng2 |
| GO:0050808 | synapse organization | -3,36472 | -0,712 | 7/472 | Bdnf,Neurod2,Nptxr,Nrp2,Nptx1,Ntng2,C1ql3 |
| GO:0099601 | regulation of neurotransmitter receptor activity | -2,98193 | -0,466 | 3/68 | Chrna4,Nptxr,Nptx1 |
| GO:0007416 | synapse assembly | -2,68574 | -0,255 | 4/184 | Bdnf,Nptxr,Nptx1,Ntng2 |
| GO:0021675 | nerve development | -2,64547 | -0,239 | 3/89 | Bdnf,Nrp2,Nptx1 |
| GO:0050804 | modulation of chemical synaptic transmission | -2,60116 | -0,208 | 7/639 | Bdnf,Grm2,Chrna4,Neurod2,Nptxr,Nptx1,Ntng2 |
| GO:0099177 | regulation of trans-synaptic signaling | -2,59735 | -0,208 | 7/640 | Bdnf,Grm2,Chrna4,Neurod2,Nptxr,Nptx1,Ntng2 |
| GO:0099173 | postsynapse organization | -2,50102 | -0,146 | 4/207 | Nptxr,Nrp2,Nptx1,Ntng2 |
| GO:0099175 | regulation of postsynapse organization | -2,2117 | 0,000 | 3/127 | Nptxr,Nrp2,Nptx1 |
| GO:0015800 | acidic amino acid transport | -5,79093 | -1,828 | 5/68 | Bdnf,Grm2,Cck,Slc17a6,Slc17a7,Slc6a7,Nppa,Chrna4,Syt17,Miat,Kcnk9,Kcnj6 |
| GO:0015800 | acidic amino acid transport | -5,79093 | -1,828 | 5/68 | Bdnf,Grm2,Cck,Slc17a6,Slc17a7 |
| GO:0006865 | amino acid transport | -5,21038 | -1,424 | 6/156 | Bdnf,Grm2,Cck,Slc17a6,Slc17a7,Slc6a7 |
| GO:0006835 | dicarboxylic acid transport | -5,07194 | -1,410 | 5/95 | Bdnf,Grm2,Cck,Slc17a6,Slc17a7 |
| GO:0006836 | neurotransmitter transport | -4,36506 | -1,059 | 7/325 | Grm2,Nppa,Chrna4,Slc17a6,Slc17a7,Slc6a7,Syt17 |
| GO:0014047 | glutamate secretion | -3,79948 | -0,933 | 3/36 | Bdnf,Grm2,Cck |
| rno05033 | Nicotine addiction | -3,66202 | -0,888 | 3/40 | Chrna4,Slc17a6,Slc17a7 |
| GO:0046942 | carboxylic acid transport | -3,40422 | -0,713 | 6/330 | Bdnf,Grm2,Cck,Slc17a6,Slc17a7,Slc6a7 |
| GO:0015849 | organic acid transport | -3,39724 | -0,713 | 6/331 | Bdnf,Grm2,Cck,Slc17a6,Slc17a7,Slc6a7 |
| GO:0003407 | neural retina development | -2,80962 | -0,352 | 3/78 | Slc17a6,Slc17a7,Miat |
| GO:0003333 | amino acid transmembrane transport | -2,67364 | -0,255 | 3/87 | Slc17a6,Slc17a7,Slc6a7 |
| GO:0015711 | organic anion transport | -2,65894 | -0,246 | 6/460 | Bdnf,Grm2,Cck,Slc17a6,Slc17a7,Slc6a7 |
| GO:0042391 | regulation of membrane potential | -2,63545 | -0,235 | 6/465 | Bdnf,Nppa,Cck,Chrna4,Kcnk9,Slc17a7 |
| rno04723 | Retrograde endocannabinoid signaling | -2,4538 | -0,119 | 3/104 | Kcnj6,Slc17a6,Slc17a7 |
| GO:0035249 | synaptic transmission, glutamatergic | -2,41907 | -0,090 | 3/107 | Grm2,Slc17a6,Slc17a7 |
| GO:0060079 | excitatory postsynaptic potential | -2,36348 | -0,068 | 3/112 | Bdnf,Chrna4,Slc17a7 |
| rno04724 | Glutamatergic synapse | -2,32092 | -0,035 | 3/116 | Grm2,Slc17a6,Slc17a7 |
| GO:0015893 | drug transport | -2,28584 | -0,004 | 4/238 | Chrna4,Slc17a6,Slc17a7,Syt17 |
| GO:0099565 | chemical synaptic transmission, postsynaptic | -2,26992 | 0,000 | 3/121 | Bdnf,Chrna4,Slc17a7 |
| GO:0099504 | synaptic vesicle cycle | -2,24791 | 0,000 | 4/244 | Chrna4,Slc17a6,Slc17a7,Syt17 |
| GO:1903825 | organic acid transmembrane transport | -2,20229 | 0,000 | 3/128 | Slc17a6,Slc17a7,Slc6a7 |
| GO:1905039 | carboxylic acid transmembrane transport | -2,20229 | 0,000 | 3/128 | Slc17a6,Slc17a7,Slc6a7 |
| GO:0001505 | regulation of neurotransmitter levels | -2,19322 | 0,000 | 5/403 | Grm2,Chrna4,Slc17a6,Slc17a7,Syt17 |
| GO:0046717 | acid secretion | -2,18371 | 0,000 | 3/130 | Bdnf,Grm2,Cck |
| GO:0006820 | anion transport | -2,12592 | 0,000 | 6/592 | Bdnf,Grm2,Cck,Slc17a6,Slc17a7,Slc6a7 |
| GO:0031644 | regulation of neurological system process | -4,88265 | -1,385 | 6/178 | Cck,Chrna4,Nptxr,Ccn3,Vip,Nptx1,Bdnf,Nppa,Wnt9a,Tnnt1,Npsr1,Syt17,Grm2,Sstr3,Kcnj6,Cyp26a1 |
| GO:0031644 | regulation of neurological system process | -4,88265 | -1,385 | 6/178 | Cck,Chrna4,Nptxr,Ccn3,Vip,Nptx1 |
| GO:0010469 | regulation of signaling receptor activity | -4,63923 | -1,278 | 9/536 | Bdnf,Nppa,Cck,Chrna4,Nptxr,Ccn3,Vip,Nptx1,Wnt9a |
| GO:0044057 | regulation of system process | -4,11531 | -1,033 | 9/626 | Nppa,Cck,Chrna4,Nptxr,Ccn3,Vip,Tnnt1,Nptx1,Npsr1 |
| GO:0051952 | regulation of amine transport | -3,33543 | -0,695 | 4/123 | Cck,Chrna4,Vip,Syt17 |
| GO:0051954 | positive regulation of amine transport | -3,32316 | -0,693 | 3/52 | Cck,Chrna4,Vip |
| GO:0015837 | amine transport | -3,2829 | -0,683 | 4/127 | Cck,Chrna4,Vip,Syt17 |
| GO:0051930 | regulation of sensory perception of pain | -3,20577 | -0,649 | 3/57 | Cck,Ccn3,Vip |
| GO:0051931 | regulation of sensory perception | -3,1836 | -0,647 | 3/58 | Cck,Ccn3,Vip |
| GO:0019233 | sensory perception of pain | -2,96967 | -0,462 | 4/154 | Cck,Chrna4,Ccn3,Vip |
| GO:0023061 | signal release | -2,95008 | -0,450 | 7/555 | Grm2,Nppa,Chrna4,Ccn3,Vip,Sstr3,Syt17 |
| GO:0050433 | regulation of catecholamine secretion | -2,87555 | -0,390 | 3/74 | Chrna4,Vip,Syt17 |
| GO:0050432 | catecholamine secretion | -2,80962 | -0,352 | 3/78 | Chrna4,Vip,Syt17 |
| GO:0051937 | catecholamine transport | -2,57793 | -0,200 | 3/94 | Chrna4,Vip,Syt17 |
| GO:0015844 | monoamine transport | -2,38538 | -0,077 | 3/110 | Chrna4,Vip,Syt17 |
| GO:0010817 | regulation of hormone levels | -2,16085 | 0,000 | 6/582 | Nppa,Kcnj6,Ccn3,Vip,Cyp26a1,Sstr3 |
| GO:0015696 | ammonium transport | -2,14749 | 0,000 | 3/134 | Nppa,Chrna4,Syt17 |
| GO:0021537 | telencephalon development | -4,2818 | -1,059 | 7/335 | Mas1,Nrp2,Slc17a6,Rtn4r,Rtn4rl2,Neurod6,Tbr1,Sstr3 |
| GO:0021537 | telencephalon development | -4,2818 | -1,059 | 7/335 | Mas1,Nrp2,Slc17a6,Rtn4r,Rtn4rl2,Neurod6,Tbr1 |
| GO:0021761 | limbic system development | -4,04835 | -1,033 | 5/155 | Mas1,Nrp2,Slc17a6,Neurod6,Tbr1 |
| GO:0030900 | forebrain development | -4,0415 | -1,033 | 8/496 | Mas1,Nrp2,Slc17a6,Rtn4r,Sstr3,Rtn4rl2,Neurod6,Tbr1 |
| GO:0021543 | pallium development | -3,22005 | -0,649 | 5/234 | Mas1,Nrp2,Slc17a6,Neurod6,Tbr1 |
| GO:0021766 | hippocampus development | -2,25017 | 0,000 | 3/123 | Mas1,Slc17a6,Neurod6 |
| rno00534 | Glycosaminoglycan biosynthesis - heparan sulfate / heparin | -4,28155 | -1,059 | 3/25 | Hs3st2,Ndst4,Hs6st3 |
| rno00534 | Glycosaminoglycan biosynthesis - heparan sulfate / heparin | -4,28155 | -1,059 | 3/25 | Hs3st2,Ndst4,Hs6st3 |
| GO:0015012 | heparan sulfate proteoglycan biosynthetic process | -4,08411 | -1,033 | 3/29 | Hs3st2,Ndst4,Hs6st3 |
| GO:0030201 | heparan sulfate proteoglycan metabolic process | -3,87439 | -0,972 | 3/34 | Hs3st2,Ndst4,Hs6st3 |
| GO:0030166 | proteoglycan biosynthetic process | -3,29875 | -0,688 | 3/53 | Hs3st2,Ndst4,Hs6st3 |
| GO:0006029 | proteoglycan metabolic process | -2,80962 | -0,352 | 3/78 | Hs3st2,Ndst4,Hs6st3 |
| GO:0007409 | axonogenesis | -4,23073 | -1,046 | 8/466 | Bdnf,Cck,Nrp2,Olfm1,Rtn4r,Nptx1,Ntng2,Tbr1,Nppa,Ccn3,Syt17,Frzb,Neurod2,Rtn4rl2 |
| GO:0007409 | axonogenesis | -4,23073 | -1,046 | 8/466 | Bdnf,Cck,Nrp2,Olfm1,Rtn4r,Nptx1,Ntng2,Tbr1 |
| GO:0061564 | axon development | -3,92844 | -0,981 | 8/515 | Bdnf,Cck,Nrp2,Olfm1,Rtn4r,Nptx1,Ntng2,Tbr1 |
| GO:0016049 | cell growth | -3,92262 | -0,981 | 8/516 | Bdnf,Nppa,Ccn3,Nrp2,Olfm1,Rtn4r,Syt17,Frzb |
| GO:0048588 | developmental cell growth | -3,88224 | -0,972 | 6/269 | Bdnf,Nppa,Nrp2,Olfm1,Rtn4r,Syt17 |
| GO:0048812 | neuron projection morphogenesis | -3,85183 | -0,968 | 9/678 | Bdnf,Cck,Nrp2,Olfm1,Rtn4r,Syt17,Nptx1,Ntng2,Tbr1 |
| GO:0120039 | plasma membrane bounded cell projection morphogenesis | -3,78032 | -0,931 | 9/693 | Bdnf,Cck,Nrp2,Olfm1,Rtn4r,Syt17,Nptx1,Ntng2,Tbr1 |
| GO:0048858 | cell projection morphogenesis | -3,76157 | -0,929 | 9/697 | Bdnf,Cck,Nrp2,Olfm1,Rtn4r,Syt17,Nptx1,Ntng2,Tbr1 |
| GO:0032990 | cell part morphogenesis | -3,66074 | -0,888 | 9/719 | Bdnf,Cck,Nrp2,Olfm1,Rtn4r,Syt17,Nptx1,Ntng2,Tbr1 |
| GO:1990138 | neuron projection extension | -3,56188 | -0,809 | 5/197 | Bdnf,Nrp2,Olfm1,Rtn4r,Syt17 |
| GO:0001558 | regulation of cell growth | -3,55444 | -0,809 | 7/439 | Bdnf,Nppa,Ccn3,Olfm1,Rtn4r,Syt17,Frzb |
| GO:0048667 | cell morphogenesis involved in neuron differentiation | -3,36323 | -0,712 | 8/624 | Bdnf,Cck,Nrp2,Olfm1,Rtn4r,Nptx1,Ntng2,Tbr1 |
| GO:0007411 | axon guidance | -3,18676 | -0,647 | 5/238 | Bdnf,Nrp2,Rtn4r,Ntng2,Tbr1 |
| GO:0097485 | neuron projection guidance | -3,17853 | -0,647 | 5/239 | Bdnf,Nrp2,Rtn4r,Ntng2,Tbr1 |
| GO:0048675 | axon extension | -3,10082 | -0,577 | 4/142 | Bdnf,Nrp2,Olfm1,Rtn4r |
| GO:0060560 | developmental growth involved in morphogenesis | -2,93414 | -0,441 | 5/271 | Bdnf,Nrp2,Olfm1,Rtn4r,Syt17 |
| GO:0000904 | cell morphogenesis involved in differentiation | -2,67385 | -0,255 | 8/798 | Bdnf,Cck,Nrp2,Olfm1,Rtn4r,Nptx1,Ntng2,Tbr1 |
| GO:0030308 | negative regulation of cell growth | -2,55469 | -0,187 | 4/200 | Nppa,Ccn3,Rtn4r,Frzb |
| GO:0050770 | regulation of axonogenesis | -2,49352 | -0,143 | 4/208 | Bdnf,Olfm1,Rtn4r,Tbr1 |
| GO:0040008 | regulation of growth | -2,38206 | -0,077 | 7/700 | Bdnf,Nppa,Ccn3,Olfm1,Rtn4r,Syt17,Frzb |
| GO:0051961 | negative regulation of nervous system development | -2,36815 | -0,068 | 5/366 | Bdnf,Neurod2,Olfm1,Rtn4r,Rtn4rl2 |
| GO:0030516 | regulation of axon extension | -2,32092 | -0,035 | 3/116 | Bdnf,Olfm1,Rtn4r |
| GO:0010721 | negative regulation of cell development | -2,22023 | 0,000 | 5/397 | Bdnf,Olfm1,Rtn4r,Frzb,Rtn4rl2 |
| GO:0061387 | regulation of extent of cell growth | -2,17454 | 0,000 | 3/131 | Bdnf,Olfm1,Rtn4r |
| GO:0006935 | chemotaxis | -2,16438 | 0,000 | 6/581 | Bdnf,Ccn3,Nrp2,Rtn4r,Ntng2,Tbr1 |
| GO:0042330 | taxis | -2,1538 | 0,000 | 6/584 | Bdnf,Ccn3,Nrp2,Rtn4r,Ntng2,Tbr1 |
| GO:0045926 | negative regulation of growth | -2,1063 | 0,000 | 4/268 | Nppa,Ccn3,Rtn4r,Frzb |
| GO:0045664 | regulation of neuron differentiation | -2,09174 | 0,000 | 7/793 | Bdnf,Neurod2,Olfm1,Rtn4r,Syt17,Rtn4rl2,Tbr1 |
| GO:0010975 | regulation of neuron projection development | -2,05167 | 0,000 | 6/614 | Bdnf,Olfm1,Rtn4r,Syt17,Rtn4rl2,Tbr1 |
| GO:0050890 | cognition | -4,15426 | -1,036 | 7/351 | Bdnf,Cck,Chrna4,Neurod2,Slc17a7,Vip,Tbr1,Npsr1,Kcnj6 |
| GO:0050890 | cognition | -4,15426 | -1,036 | 7/351 | Bdnf,Cck,Chrna4,Neurod2,Slc17a7,Vip,Tbr1 |
| GO:0007611 | learning or memory | -3,51925 | -0,799 | 6/314 | Bdnf,Cck,Neurod2,Slc17a7,Vip,Tbr1 |
| GO:0001662 | behavioural fear response | -3,29875 | -0,688 | 3/53 | Bdnf,Cck,Neurod2 |
| GO:0002209 | behavioural defense response | -3,27483 | -0,683 | 3/54 | Bdnf,Cck,Neurod2 |
| GO:0007631 | feeding behaviour | -3,25731 | -0,675 | 4/129 | Bdnf,Cck,Npsr1,Tbr1 |
| GO:0042596 | fear response | -3,20577 | -0,649 | 3/57 | Bdnf,Cck,Neurod2 |
| GO:0007610 | behaviour | -2,85836 | -0,380 | 8/746 | Bdnf,Cck,Chrna4,Neurod2,Slc17a7,Vip,Npsr1,Tbr1 |
| GO:0051602 | response to electrical stimulus | -2,7937 | -0,343 | 3/79 | Bdnf,Kcnj6,Neurod2 |
| GO:0007612 | learning | -2,72933 | -0,291 | 4/179 | Bdnf,Cck,Neurod2,Tbr1 |
| GO:0033555 | multicellular organismal response to stress | -2,4538 | -0,119 | 3/104 | Bdnf,Cck,Neurod2 |
| GO:0008306 | associative learning | -2,3965 | -0,077 | 3/109 | Cck,Neurod2,Tbr1 |
| GO:0007218 | neuropeptide signaling pathway | -3,69524 | -0,888 | 4/99 | Nppa,Nmbr,Sstr3,Npsr1 |
| GO:0007218 | neuropeptide signaling pathway | -3,69524 | -0,888 | 4/99 | Nppa,Nmbr,Sstr3,Npsr1 |
| rno04080 | Neuroactive ligand-receptor interaction | -3,6888 | -0,888 | 6/292 | Grm2,Mas1,Nmbr,Chrna4,Sstr3,Rxfp1,Cck,Vip |
| rno04080 | Neuroactive ligand-receptor interaction | -3,6888 | -0,888 | 6/292 | Grm2,Mas1,Nmbr,Chrna4,Sstr3,Rxfp1 |
| R-RNO-500792 | GPCR ligand binding | -2,48253 | -0,138 | 5/344 | Grm2,Cck,Vip,Sstr3,Rxfp1 |
| GO:0043269 | regulation of ion transport | -3,52064 | -0,799 | 9/751 | Nppa,Cck,Chrna4,Kcnj6,Kcnk9,Vip,Syt17,Npsr1,Stac2,Slc17a6,Slc17a7,Slc30a3 |
| GO:0043269 | regulation of ion transport | -3,52064 | -0,799 | 9/751 | Nppa,Cck,Chrna4,Kcnj6,Kcnk9,Vip,Syt17,Npsr1,Stac2 |
| GO:0034765 | regulation of ion transmembrane transport | -2,55336 | -0,187 | 6/483 | Nppa,Chrna4,Kcnj6,Kcnk9,Npsr1,Stac2 |
| GO:0043270 | positive regulation of ion transport | -2,54302 | -0,182 | 5/333 | Nppa,Cck,Chrna4,Npsr1,Stac2 |
| GO:0015672 | monovalent inorganic cation transport | -2,39176 | -0,077 | 6/521 | Nppa,Kcnj6,Kcnk9,Slc17a6,Slc17a7,Vip |
| GO:0006813 | potassium ion transport | -2,24791 | 0,000 | 4/244 | Nppa,Kcnj6,Kcnk9,Vip |
| GO:0098662 | inorganic cation transmembrane transport | -2,23563 | 0,000 | 7/745 | Nppa,Kcnj6,Kcnk9,Slc17a7,Npsr1,Stac2,Slc30a3 |
| GO:0034762 | regulation of transmembrane transport | -2,17148 | 0,000 | 6/579 | Nppa,Chrna4,Kcnj6,Kcnk9,Npsr1,Stac2 |
| R-RNO-163125 | Post-translational modification: synthesis of GPI-anchored proteins | -2,7625 | -0,318 | 3/81 | Opcml,Rtn4rl2,Ntng2 |
| R-RNO-163125 | Post-translational modification: synthesis of GPI-anchored proteins | -2,7625 | -0,318 | 3/81 | Opcml,Rtn4rl2,Ntng2 |
| GO:0014033 | neural crest cell differentiation | -2,59113 | -0,208 | 3/93 | Nrp2,Cyp26a1,Frzb,Olfm1 |
| GO:0014033 | neural crest cell differentiation | -2,59113 | -0,208 | 3/93 | Nrp2,Cyp26a1,Frzb |
| GO:0048762 | mesenchymal cell differentiation | -2,41362 | -0,089 | 4/219 | Nrp2,Olfm1,Cyp26a1,Frzb |
| GO:0060485 | mesenchyme development | -2,04626 | 0,000 | 4/279 | Nrp2,Olfm1,Cyp26a1,Frzb |

**Table S5: Metascape Enrichment metrics and DEG composition of each significant gene ontology in the SAL-NS vs LPS-S comparison in the NAcc**

| Term | Description | LogP | Log(q-value) | InTerm_InList | Symbols |
| --- | --- | --- | --- | --- | --- |
| GO:0035082 | axoneme assembly | -6,65556 | -2,377 | 6/76 | Foxj1,Dnah1,Dnai3,Odad1,Ak7,Dnaaf3,Flna,Cfap126,Drc7,Itgb4,Aurkb |
| GO:0035082 | axoneme assembly | -6,65556 | -2,377 | 6/76 | Foxj1,Dnah1,Dnai3,Odad1,Ak7,Dnaaf3 |
| GO:0001578 | microtubule bundle formation | -5,7266 | -1,749 | 6/109 | Foxj1,Dnah1,Dnai3,Odad1,Ak7,Dnaaf3 |
| GO:0070286 | axonemal dynein complex assembly | -5,38878 | -1,712 | 4/32 | Dnah1,Dnai3,Odad1,Dnaaf3 |
| GO:0044782 | cilium organization | -4,35021 | -0,975 | 8/383 | Foxj1,Dnah1,Dnai3,Flna,Odad1,Ak7,Cfap126,Dnaaf3 |
| GO:0060271 | cilium assembly | -3,70404 | -0,505 | 7/355 | Foxj1,Dnah1,Dnai3,Flna,Odad1,Ak7,Dnaaf3 |
| GO:0003341 | cilium movement | -3,35971 | -0,307 | 5/187 | Dnah1,Drc7,Dnai3,Odad1,Ak7 |
| GO:0120031 | plasma membrane bounded cell projection assembly | -3,18862 | -0,212 | 8/567 | Itgb4,Foxj1,Dnah1,Dnai3,Flna,Odad1,Ak7,Dnaaf3 |
| GO:0030031 | cell projection assembly | -3,11949 | -0,209 | 8/581 | Itgb4,Foxj1,Dnah1,Dnai3,Flna,Odad1,Ak7,Dnaaf3 |
| GO:0000226 | microtubule cytoskeleton organization | -3,01031 | -0,147 | 8/604 | Aurkb,Foxj1,Dnah1,Dnai3,Flna,Odad1,Ak7,Dnaaf3 |
| GO:0031644 | regulation of nervous system process | -5,40737 | -1,712 | 7/192 | Agt,Cck,Ctss,Nptxr,Ccn3,Nptx1,Cbln1,Abcg2 |
| GO:0031644 | regulation of nervous system process | -5,40737 | -1,712 | 7/192 | Agt,Cck,Ctss,Nptxr,Ccn3,Nptx1,Cbln1 |
| GO:0051930 | regulation of sensory perception of pain | -2,97121 | -0,142 | 3/59 | Cck,Ctss,Ccn3 |
| GO:0051931 | regulation of sensory perception | -2,94998 | -0,142 | 3/60 | Cck,Ctss,Ccn3 |
| GO:0044057 | regulation of system process | -2,81958 | -0,060 | 8/647 | Agt,Cck,Ctss,Nptxr,Ccn3,Nptx1,Abcg2,Cbln1 |
| GO:0014002 | astrocyte development | -4,94907 | -1,370 | 4/41 | Agt,Vim,Mt3,C1qa,Itgb4,Cck,Rps29,Ramp3,Olfm1,Top2a,Ifi27,Abcg2,Aurkb |
| GO:0014002 | astrocyte development | -4,94907 | -1,370 | 4/41 | Agt,Vim,Mt3,C1qa |
| GO:0021782 | glial cell development | -4,06385 | -0,740 | 5/132 | Agt,Itgb4,Vim,Mt3,C1qa |
| GO:0048708 | astrocyte differentiation | -3,58279 | -0,435 | 4/91 | Agt,Vim,Mt3,C1qa |
| GO:0010942 | positive regulation of cell death | -2,61723 | 0,000 | 8/697 | Agt,Cck,Rps29,Ramp3,Olfm1,Mt3,C1qa,Top2a |
| GO:0010001 | glial cell differentiation | -2,5733 | 0,000 | 5/281 | Agt,Itgb4,Vim,Mt3,C1qa |
| GO:0007565 | female pregnancy | -2,47642 | 0,000 | 5/296 | Agt,Itgb4,Vim,Ifi27,Abcg2 |
| GO:0044706 | multi-multicellular organism process | -2,27142 | 0,000 | 5/331 | Agt,Itgb4,Vim,Ifi27,Abcg2 |
| GO:0042063 | gliogenesis | -2,03873 | 0,000 | 5/377 | Agt,Itgb4,Vim,Mt3,C1qa |
| GO:0007568 | aging | -2,02973 | 0,000 | 6/532 | Agt,Vim,Aurkb,C1qa,Abcg2,Top2a |
| GO:0032355 | response to estradiol | -4,5726 | -1,095 | 7/258 | Agt,C3,Ramp3,Vim,Foxj1,Ifi27,Abcg2,Ctss,Nr4a2,Mt3,Npas4,Mc4r |
| GO:0032355 | response to estradiol | -4,5726 | -1,095 | 7/258 | Agt,C3,Ramp3,Vim,Foxj1,Ifi27,Abcg2 |
| GO:0032870 | cellular response to hormone stimulus | -2,43298 | 0,000 | 8/747 | Agt,Ctss,Nr4a2,Ramp3,Foxj1,Mt3,Npas4,Abcg2 |
| GO:1901652 | response to peptide | -2,08552 | 0,000 | 7/681 | Agt,Mc4r,Nr4a2,Ramp3,Vim,Foxj1,Abcg2 |
| GO:0099150 | regulation of postsynaptic specialization assembly | -4,52837 | -1,095 | 3/18 | Nptxr,Nptx1,Cbln1,Agt,C3,Itgb4,Npas4,Cbln2,Flna,C1qa |
| GO:0099150 | regulation of postsynaptic specialization assembly | -4,52837 | -1,095 | 3/18 | Nptxr,Nptx1,Cbln1 |
| GO:0098698 | postsynaptic specialization assembly | -3,93526 | -0,657 | 3/28 | Nptxr,Nptx1,Cbln1 |
| GO:0099068 | postsynapse assembly | -3,719 | -0,505 | 3/33 | Nptxr,Nptx1,Cbln1 |
| GO:0034330 | cell junction organization | -3,56689 | -0,435 | 10/777 | Agt,C3,Itgb4,Nptxr,Npas4,Nptx1,Cbln2,Flna,C1qa,Cbln1 |
| GO:0050808 | synapse organization | -3,51641 | -0,414 | 8/506 | C3,Nptxr,Npas4,Nptx1,Cbln2,Flna,C1qa,Cbln1 |
| GO:0007416 | synapse assembly | -3,2768 | -0,254 | 5/195 | Nptxr,Npas4,Nptx1,Cbln2,Cbln1 |
| GO:0099084 | postsynaptic specialization organization | -3,1815 | -0,212 | 3/50 | Nptxr,Nptx1,Cbln1 |
| GO:0034329 | cell junction assembly | -3,12104 | -0,209 | 7/444 | Agt,Itgb4,Nptxr,Npas4,Nptx1,Cbln2,Cbln1 |
| GO:1901888 | regulation of cell junction assembly | -2,16001 | 0,000 | 4/222 | Agt,Nptx1,Cbln2,Cbln1 |
| GO:0051963 | regulation of synapse assembly | -2,08867 | 0,000 | 3/121 | Nptx1,Cbln2,Cbln1 |
| GO:1901890 | positive regulation of cell junction assembly | -2,0789 | 0,000 | 3/122 | Agt,Cbln2,Cbln1 |
| GO:0061564 | axon development | -3,35488 | -0,307 | 8/535 | Cck,Nr4a2,Vim,Olfm1,Rtn4r,Mt3,Nptx1,Flna,Agt,Nrn1,Necab3,Aurkb,Dnai3,Top2a,Cbln1,Ccn3 |
| GO:0061564 | axon development | -3,35488 | -0,307 | 8/535 | Cck,Nr4a2,Vim,Olfm1,Rtn4r,Mt3,Nptx1,Flna |
| GO:0007409 | axonogenesis | -2,91341 | -0,140 | 7/482 | Cck,Nr4a2,Vim,Olfm1,Rtn4r,Mt3,Nptx1 |
| GO:0048588 | developmental cell growth | -2,58667 | 0,000 | 5/279 | Agt,Nrn1,Olfm1,Rtn4r,Mt3 |
| GO:0048812 | neuron projection morphogenesis | -2,5828 | 0,000 | 8/706 | Cck,Nr4a2,Vim,Nrn1,Olfm1,Rtn4r,Mt3,Nptx1 |
| GO:0010977 | negative regulation of neuron projection development | -2,57964 | 0,000 | 4/169 | Vim,Rtn4r,Mt3,Flna |
| GO:0120039 | plasma membrane bounded cell projection morphogenesis | -2,51932 | 0,000 | 8/723 | Cck,Nr4a2,Vim,Nrn1,Olfm1,Rtn4r,Mt3,Nptx1 |
| GO:0048858 | cell projection morphogenesis | -2,50467 | 0,000 | 8/727 | Cck,Nr4a2,Vim,Nrn1,Olfm1,Rtn4r,Mt3,Nptx1 |
| GO:0032990 | cell part morphogenesis | -2,42596 | 0,000 | 8/749 | Cck,Nr4a2,Vim,Nrn1,Olfm1,Rtn4r,Mt3,Nptx1 |
| GO:0001764 | neuron migration | -2,38934 | 0,000 | 4/191 | Cck,Nr4a2,Olfm1,Flna |
| WP2433 | Spinal Cord Injury | -2,32961 | 0,000 | 3/99 | Vim,Rtn4r,Necab3 |
| GO:0051129 | negative regulation of cellular component organization | -2,32014 | 0,000 | 8/780 | Vim,Rtn4r,Aurkb,Mt3,Dnai3,Flna,Top2a,Cbln1 |
| GO:1990138 | neuron projection extension | -2,28083 | 0,000 | 4/205 | Nrn1,Olfm1,Rtn4r,Mt3 |
| GO:0030308 | negative regulation of cell growth | -2,2734 | 0,000 | 4/206 | Agt,Ccn3,Rtn4r,Mt3 |
| GO:0031345 | negative regulation of cell projection organization | -2,18745 | 0,000 | 4/218 | Vim,Rtn4r,Mt3,Flna |
| GO:0048667 | cell morphogenesis involved in neuron differentiation | -2,18517 | 0,000 | 7/652 | Cck,Nr4a2,Vim,Olfm1,Rtn4r,Mt3,Nptx1 |
| GO:0030516 | regulation of axon extension | -2,10849 | 0,000 | 3/119 | Olfm1,Rtn4r,Mt3 |
| GO:0016049 | cell growth | -2,04113 | 0,000 | 6/529 | Agt,Ccn3,Nrn1,Olfm1,Rtn4r,Mt3 |
| GO:0060079 | excitatory postsynaptic potential | -3,16832 | -0,212 | 4/117 | P2rx6,Slc17a7,Npas4,Cbln1,Agt,Cck,Flna |
| GO:0060079 | excitatory postsynaptic potential | -3,16832 | -0,212 | 4/117 | P2rx6,Slc17a7,Npas4,Cbln1 |
| GO:0099565 | chemical synaptic transmission, postsynaptic | -3,04786 | -0,167 | 4/126 | P2rx6,Slc17a7,Npas4,Cbln1 |
| GO:0042391 | regulation of membrane potential | -2,92906 | -0,142 | 7/479 | Agt,P2rx6,Cck,Slc17a7,Npas4,Flna,Cbln1 |
| GO:0060078 | regulation of postsynaptic membrane potential | -2,65589 | 0,000 | 4/161 | P2rx6,Slc17a7,Npas4,Cbln1 |
| GO:0002455 | humoral immune response mediated by circulating immunoglobulin | -3,10721 | -0,209 | 3/53 | C3,Foxj1,C1qa,Nr4a2 |
| GO:0002455 | humoral immune response mediated by circulating immunoglobulin | -3,10721 | -0,209 | 3/53 | C3,Foxj1,C1qa |
| GO:0042551 | neuron maturation | -2,92913 | -0,142 | 3/61 | C3,Nr4a2,C1qa |
| R-RNO-500792 | GPCR ligand binding | -2,93926 | -0,142 | 6/347 | Agt,C3,Cck,Mc4r,Ramp3,Gng13,Ctss,Flna,Ccn3,Smoc2 |
| R-RNO-500792 | GPCR ligand binding | -2,93926 | -0,142 | 6/347 | Agt,C3,Cck,Mc4r,Ramp3,Gng13 |
| GO:0034764 | positive regulation of transmembrane transport | -2,73428 | -0,024 | 5/258 | Agt,C3,Ctss,Ramp3,Flna |
| GO:0034767 | positive regulation of ion transmembrane transport | -2,73428 | -0,024 | 5/258 | Agt,C3,Ctss,Ramp3,Flna |
| R-RNO-375276 | Peptide ligand-binding receptors | -2,71592 | -0,017 | 4/155 | Agt,C3,Cck,Mc4r |
| GO:1904064 | positive regulation of cation transmembrane transport | -2,53402 | 0,000 | 4/174 | Agt,Ctss,Ramp3,Flna |
| GO:0048771 | tissue remodeling | -2,20841 | 0,000 | 4/215 | Agt,Mc4r,Ctss,Flna |
| R-RNO-372790 | Signaling by GPCR | -2,10338 | 0,000 | 6/513 | Agt,C3,Cck,Mc4r,Ramp3,Gng13 |
| GO:0001525 | angiogenesis | -2,07584 | 0,000 | 6/520 | Agt,C3,Ramp3,Ccn3,Smoc2,Flna |
| GO:0010043 | response to zinc ion | -2,77442 | -0,027 | 3/69 | Vim,Mt3,Slc30a3,C3,Nptx1,C1qa,Abcg2,Myo1b |
| GO:0010043 | response to zinc ion | -2,77442 | -0,027 | 3/69 | Vim,Mt3,Slc30a3 |
| GO:0010038 | response to metal ion | -2,73445 | -0,024 | 7/518 | C3,Vim,Mt3,Nptx1,C1qa,Abcg2,Slc30a3 |
| GO:0000041 | transition metal ion transport | -2,08867 | 0,000 | 3/121 | Mt3,Myo1b,Slc30a3 |
| GO:0001656 | metanephros development | -2,32961 | 0,000 | 3/99 | Ctss,Foxj1,Fras1,Agt |
| GO:0001656 | metanephros development | -2,32961 | 0,000 | 3/99 | Ctss,Foxj1,Fras1 |
| GO:0060993 | kidney morphogenesis | -2,22452 | 0,000 | 3/108 | Agt,Foxj1,Fras1 |
| GO:0050795 | regulation of behaviour | -2,28166 | 0,000 | 3/103 | Cck,Cort,Mc4r,Ramp3,Flna |
| GO:0050795 | regulation of behaviour | -2,28166 | 0,000 | 3/103 | Cck,Cort,Mc4r |
| GO:0007188 | adenylate cyclase-modulating G protein-coupled receptor signaling pathway | -2,15324 | 0,000 | 4/223 | Cort,Mc4r,Ramp3,Flna |
| WP30 | Cytoplasmic Ribosomal Proteins | -2,25845 | 0,000 | 3/105 | Rps29,Rpl41,Rps28 |
| WP30 | Cytoplasmic Ribosomal Proteins | -2,25845 | 0,000 | 3/105 | Rps29,Rpl41,Rps28 |
| GO:0002181 | cytoplasmic translation | -2,17023 | 0,000 | 3/113 | Rps29,Rpl41,Rps28 |
| GO:0007229 | integrin-mediated signaling pathway | -2,19162 | 0,000 | 3/111 | Itgb4,Flna,Adam11 |
| GO:0007229 | integrin-mediated signaling pathway | -2,19162 | 0,000 | 3/111 | Itgb4,Flna,Adam11 |
| GO:0009141 | nucleoside triphosphate metabolic process | -2,13892 | 0,000 | 3/116 | Atp5f1e,Guk1,Ak7 |
| GO:0009141 | nucleoside triphosphate metabolic process | -2,13892 | 0,000 | 3/116 | Atp5f1e,Guk1,Ak7 |

**Table S6: Metascape Enrichment metrics and DEG composition of each significant gene ontology in the SAL-NS vs LPS-NS comparison in the DLS**

| Term | Description | LogP | Log(q-value) | InTerm_InList | Symbols |
| --- | --- | --- | --- | --- | --- |
| ko03010 | Ribosome | -25,2267 | -21,248 | 69/168 | Rps29,Rps5,Rps26,Rpl32,Rpsa,Rpl9,Rps7,Rps14,Rps15,Rps17,Rps3a,Fau,Rplp0,Rpl4,Rpl27,Rpl24,Mrpl23,Rpl28,Rpl30,Rps12,Rpl21,Rpl5,Rpl13,Rpl18,Rpl19,Rpl22,Rpl36al,Rpl37,Rps10,Rps21,Rps24,Rps2,Rpl6,Rps20,Rps23,Rpl41,Rps3,Rps16,Rplp1,Rps13,Rpl26,Mrpl27,Mrps14,Mrps18c,Mrpl34,Rpl18a,Mrps12,Rpl36a,mrpl11,Mrpl16,Rps18,Mrpl19,Rpl7,Mrpl13,Rpl3,Mrps18a,Mrpl14,Mrpl12,Rpl13a,Rpl23a,Rpl22l1,Rpl34,Rpl12,Rps27l,Rpl38,Mrps21,Rps28,Rps4x,Rps27a,Mrps34,Srp19,Eif3f,Trmt112,Eef1g,Mrps18b,Eif3m,Srp14,Eif2s2,Mrpl41,Mrps33,Mrps25,Mrps35,Mrpl54,Eef1d,Eif5b,Mrps23,Chchd1,Mrpl52,Eef1b2,Srp9,Eif3j,Rps6ka6,Casc3,Rbm8a,Upf3b,Dcp1a,Upf3a,Ddx47,Fcf1,Exosc5,Imp3,Ddx21,Bysl,Dcaf13,Rrp9,Anp32a,Psmb5,Psmb6,Psmb3,Psmc3,Ubc,Adar,Psmd4,Polr2f,Sf3b1,Xpo1,Psmb7,Polr2g,Prkcd,Khsrp,Cnot6,Prpf8,Lsm4,Psmb10,Psmd8,Lsm8,Lsm3,Sf3a2,Cnot2,Xrn1,Thoc7,Hnrnph2,Zc3h11a,Thoc5,Snrnp70,Patl1,Lsm7,Cpsf7,Pan2,Psme4,Gtf2h5,Snrpd2,Sf3b5,Snrpg,Lsm2,Polr2e,Polr2k,Rwdd1,Eif3k,Mcts1,Cpeb4,Zc3h15,Dph3,Denr |
| ko03010 | Ribosome | -25,2267 | -21,248 | 69/168 | Rps29,Rps5,Rps26,Rpl32,Rpsa,Rpl9,Rps7,Rps14,Rps15,Rps17,Rps3a,Fau,Rplp0,Rpl4,Rpl27,Rpl24,Mrpl23,Rpl28,Rpl30,Rps12,Rpl21,Rpl5,Rpl13,Rpl18,Rpl19,Rpl22,Rpl36al,Rpl37,Rps10,Rps21,Rps24,Rps2,Rpl6,Rps20,Rps23,Rpl41,Rps3,Rps16,Rplp1,Rps13,Rpl26,Mrpl27,Mrps14,Mrps18c,Mrpl34,Rpl18a,Mrps12,Rpl36a,mrpl11,Mrpl16,Rps18,Mrpl19,Rpl7,Mrpl13,Rpl3,Mrps18a,Mrpl14,Mrpl12,Rpl13a,Rpl23a,Rpl22l1,Rpl34,Rpl12,Rps27l,Rpl38,Mrps21,Rps28,Rps4x,Rps27a |
| rno03010 | Ribosome | -25,2267 | -21,248 | 69/168 | Rps29,Rps5,Rps26,Rpl32,Rpsa,Rpl9,Rps7,Rps14,Rps15,Rps17,Rps3a,Fau,Rplp0,Rpl4,Rpl27,Rpl24,Mrpl23,Rpl28,Rpl30,Rps12,Rpl21,Rpl5,Rpl13,Rpl18,Rpl19,Rpl22,Rpl36al,Rpl37,Rps10,Rps21,Rps24,Rps2,Rpl6,Rps20,Rps23,Rpl41,Rps3,Rps16,Rplp1,Rps13,Rpl26,Mrpl27,Mrps14,Mrps18c,Mrpl34,Rpl18a,Mrps12,Rpl36a,mrpl11,Mrpl16,Rps18,Mrpl19,Rpl7,Mrpl13,Rpl3,Mrps18a,Mrpl14,Mrpl12,Rpl13a,Rpl23a,Rpl22l1,Rpl34,Rpl12,Rps27l,Rpl38,Mrps21,Rps28,Rps4x,Rps27a |
| R-RNO-72766 | Translation | -22,9514 | -19,149 | 78/224 | Rps29,Rpl32,Rpsa,Rpl9,Rps7,Rps14,Rps15,Rps17,Rps3a,Rplp0,Rpl4,Rpl27,Rpl24,Mrpl23,Rpl30,Rpl5,Rpl13,Rpl18,Rpl19,Rpl22,Rpl36al,Rpl37,Rps10,Rps21,Rps24,Rpl6,Rps20,Rps23,Rps3,Rps16,Rplp1,Rps13,Mrps34,Mrpl27,Mrps14,Mrps18c,Mrpl34,Rpl18a,Srp19,Mrps12,Rpl36a,Eif3f,mrpl11,Trmt112,Eef1g,Mrpl16,Mrps18b,Rps18,Eif3m,Srp14,Eif2s2,Mrpl41,Mrps33,Mrpl19,Mrps25,Mrps35,Mrpl54,Mrpl13,Eef1d,Rpl3,Mrps18a,Mrpl14,Mrpl12,Eif5b,Mrps23,Chchd1,Mrpl52,Rpl22l1,Eef1b2,Rpl12,Rps27l,Rpl38,Mrps21,Srp9,Rps28,Eif3j,Rps4x,Rps27a |
| WP30 | Cytoplasmic Ribosomal Proteins | -18,3821 | -14,705 | 46/105 | Rps29,Rps5,Rps26,Rpl32,Rpsa,Rps7,Rps14,Rps15,Rps3a,Fau,Rpl4,Rpl27,Rpl24,Rpl28,Rpl30,Rps12,Rpl13,Rpl18,Rpl19,Rpl22,Rpl37,Rps10,Rps21,Rps24,Rps2,Rpl6,Rps20,Rps23,Rpl41,Rps3,Rps16,Rplp1,Rps13,Rpl26,Rpl18a,Rpl36a,Rps18,Mrpl19,Rpl7,Rpl3,Rps6ka6,Rpl13a,Rpl23a,Rpl34,Rpl38,Rps28 |
| R-RNO-1799339 | SRP-dependent cotranslational protein targeting to membrane | -17,5387 | -13,958 | 45/105 | Rps29,Rpl32,Rpsa,Rpl9,Rps7,Rps14,Rps15,Rps17,Rps3a,Rplp0,Rpl4,Rpl27,Rpl24,Rpl30,Rpl5,Rpl13,Rpl18,Rpl19,Rpl22,Rpl36al,Rpl37,Rps10,Rps21,Rps24,Rpl6,Rps20,Rps23,Rps3,Rps16,Rplp1,Rps13,Rpl18a,Srp19,Rpl36a,Rps18,Srp14,Rpl3,Rpl22l1,Rpl12,Rps27l,Rpl38,Srp9,Rps28,Rps4x,Rps27a |
| R-RNO-72689 | Formation of a pool of free 40S subunits | -15,8797 | -12,453 | 45/114 | Rps29,Rpl32,Rpsa,Rpl9,Rps7,Rps14,Rps15,Rps17,Rps3a,Rplp0,Rpl4,Rpl27,Rpl24,Rpl30,Rpl5,Rpl13,Rpl18,Rpl19,Rpl22,Rpl36al,Rpl37,Rps10,Rps21,Rps24,Rpl6,Rps20,Rps23,Rps3,Rps16,Rplp1,Rps13,Rpl18a,Rpl36a,Eif3f,Rps18,Eif3m,Rpl3,Rpl22l1,Rpl12,Rps27l,Rpl38,Rps28,Eif3j,Rps4x,Rps27a |
| R-RNO-927802 | Nonsense-Mediated Decay (NMD) | -15,7323 | -12,453 | 47/124 | Rps29,Rpl32,Rpsa,Rpl9,Rps7,Rps14,Rps15,Rps17,Rps3a,Rplp0,Rpl4,Rpl27,Rpl24,Rpl30,Rpl5,Rpl13,Rpl18,Rpl19,Rpl22,Rpl36al,Rpl37,Rps10,Rps21,Rps24,Rpl6,Rps20,Rps23,Rps3,Rps16,Rplp1,Rps13,Casc3,Rpl18a,Rpl36a,Rps18,Rbm8a,Rpl3,Upf3b,Dcp1a,Upf3a,Rpl22l1,Rpl12,Rps27l,Rpl38,Rps28,Rps4x,Rps27a |
| R-RNO-975957 | Nonsense Mediated Decay (NMD) enhanced by the Exon Junction Complex (EJC) | -15,7323 | -12,453 | 47/124 | Rps29,Rpl32,Rpsa,Rpl9,Rps7,Rps14,Rps15,Rps17,Rps3a,Rplp0,Rpl4,Rpl27,Rpl24,Rpl30,Rpl5,Rpl13,Rpl18,Rpl19,Rpl22,Rpl36al,Rpl37,Rps10,Rps21,Rps24,Rpl6,Rps20,Rps23,Rps3,Rps16,Rplp1,Rps13,Casc3,Rpl18a,Rpl36a,Rps18,Rbm8a,Rpl3,Upf3b,Dcp1a,Upf3a,Rpl22l1,Rpl12,Rps27l,Rpl38,Rps28,Rps4x,Rps27a |
| R-RNO-72706 | GTP hydrolysis and joining of the 60S ribosomal subunit | -15,4135 | -12,194 | 47/126 | Rps29,Rpl32,Rpsa,Rpl9,Rps7,Rps14,Rps15,Rps17,Rps3a,Rplp0,Rpl4,Rpl27,Rpl24,Rpl30,Rpl5,Rpl13,Rpl18,Rpl19,Rpl22,Rpl36al,Rpl37,Rps10,Rps21,Rps24,Rpl6,Rps20,Rps23,Rps3,Rps16,Rplp1,Rps13,Rpl18a,Rpl36a,Eif3f,Rps18,Eif3m,Eif2s2,Rpl3,Eif5b,Rpl22l1,Rpl12,Rps27l,Rpl38,Rps28,Eif3j,Rps4x,Rps27a |
| R-RNO-156827 | L13a-mediated translational silencing of Ceruloplasmin expression | -14,9932 | -11,828 | 46/124 | Rps29,Rpl32,Rpsa,Rpl9,Rps7,Rps14,Rps15,Rps17,Rps3a,Rplp0,Rpl4,Rpl27,Rpl24,Rpl30,Rpl5,Rpl13,Rpl18,Rpl19,Rpl22,Rpl36al,Rpl37,Rps10,Rps21,Rps24,Rpl6,Rps20,Rps23,Rps3,Rps16,Rplp1,Rps13,Rpl18a,Rpl36a,Eif3f,Rps18,Eif3m,Eif2s2,Rpl3,Rpl22l1,Rpl12,Rps27l,Rpl38,Rps28,Eif3j,Rps4x,Rps27a |
| R-RNO-975956 | Nonsense Mediated Decay (NMD) independent of the Exon Junction Complex (EJC) | -14,5935 | -11,490 | 42/108 | Rps29,Rpl32,Rpsa,Rpl9,Rps7,Rps14,Rps15,Rps17,Rps3a,Rplp0,Rpl4,Rpl27,Rpl24,Rpl30,Rpl5,Rpl13,Rpl18,Rpl19,Rpl22,Rpl36al,Rpl37,Rps10,Rps21,Rps24,Rpl6,Rps20,Rps23,Rps3,Rps16,Rplp1,Rps13,Rpl18a,Rpl36a,Rps18,Rpl3,Rpl22l1,Rpl12,Rps27l,Rpl38,Rps28,Rps4x,Rps27a |
| R-RNO-72613 | Eukaryotic Translation Initiation | -14,5044 | -11,456 | 47/132 | Rps29,Rpl32,Rpsa,Rpl9,Rps7,Rps14,Rps15,Rps17,Rps3a,Rplp0,Rpl4,Rpl27,Rpl24,Rpl30,Rpl5,Rpl13,Rpl18,Rpl19,Rpl22,Rpl36al,Rpl37,Rps10,Rps21,Rps24,Rpl6,Rps20,Rps23,Rps3,Rps16,Rplp1,Rps13,Rpl18a,Rpl36a,Eif3f,Rps18,Eif3m,Eif2s2,Rpl3,Eif5b,Rpl22l1,Rpl12,Rps27l,Rpl38,Rps28,Eif3j,Rps4x,Rps27a |
| R-RNO-72737 | Cap-dependent Translation Initiation | -14,5044 | -11,456 | 47/132 | Rps29,Rpl32,Rpsa,Rpl9,Rps7,Rps14,Rps15,Rps17,Rps3a,Rplp0,Rpl4,Rpl27,Rpl24,Rpl30,Rpl5,Rpl13,Rpl18,Rpl19,Rpl22,Rpl36al,Rpl37,Rps10,Rps21,Rps24,Rpl6,Rps20,Rps23,Rps3,Rps16,Rplp1,Rps13,Rpl18a,Rpl36a,Eif3f,Rps18,Eif3m,Eif2s2,Rpl3,Eif5b,Rpl22l1,Rpl12,Rps27l,Rpl38,Rps28,Eif3j,Rps4x,Rps27a |
| R-RNO-6791226 | Major pathway of rRNA processing in the nucleolus and cytosol | -11,4918 | -8,718 | 50/172 | Rps29,Rpl32,Rpsa,Rpl9,Rps7,Rps14,Rps15,Rps17,Rps3a,Rplp0,Rpl4,Rpl27,Rpl24,Rpl30,Rpl5,Rpl13,Rpl18,Rpl19,Rpl22,Rpl36al,Rpl37,Rps10,Rps21,Rps24,Rpl6,Rps20,Rps23,Rps3,Rps16,Rplp1,Rps13,Rpl18a,Rpl36a,Rps18,Ddx47,Fcf1,Rpl3,Exosc5,Imp3,Ddx21,Bysl,Rpl22l1,Dcaf13,Rrp9,Rpl12,Rps27l,Rpl38,Rps28,Rps4x,Rps27a |
| R-RNO-72312 | rRNA processing | -11,4918 | -8,718 | 50/172 | Rps29,Rpl32,Rpsa,Rpl9,Rps7,Rps14,Rps15,Rps17,Rps3a,Rplp0,Rpl4,Rpl27,Rpl24,Rpl30,Rpl5,Rpl13,Rpl18,Rpl19,Rpl22,Rpl36al,Rpl37,Rps10,Rps21,Rps24,Rpl6,Rps20,Rps23,Rps3,Rps16,Rplp1,Rps13,Rpl18a,Rpl36a,Rps18,Ddx47,Fcf1,Rpl3,Exosc5,Imp3,Ddx21,Bysl,Rpl22l1,Dcaf13,Rrp9,Rpl12,Rps27l,Rpl38,Rps28,Rps4x,Rps27a |
| R-RNO-8868773 | rRNA processing in the nucleus and cytosol | -11,4918 | -8,718 | 50/172 | Rps29,Rpl32,Rpsa,Rpl9,Rps7,Rps14,Rps15,Rps17,Rps3a,Rplp0,Rpl4,Rpl27,Rpl24,Rpl30,Rpl5,Rpl13,Rpl18,Rpl19,Rpl22,Rpl36al,Rpl37,Rps10,Rps21,Rps24,Rpl6,Rps20,Rps23,Rps3,Rps16,Rplp1,Rps13,Rpl18a,Rpl36a,Rps18,Ddx47,Fcf1,Rpl3,Exosc5,Imp3,Ddx21,Bysl,Rpl22l1,Dcaf13,Rrp9,Rpl12,Rps27l,Rpl38,Rps28,Rps4x,Rps27a |
| R-RNO-72695 | Formation of the ternary complex, and subsequently, the 43S complex | -9,28237 | -6,735 | 24/58 | Rps29,Rpsa,Rps7,Rps14,Rps15,Rps17,Rps3a,Rps10,Rps21,Rps24,Rps20,Rps23,Rps3,Rps16,Rps13,Eif3f,Rps18,Eif3m,Eif2s2,Rps27l,Rps28,Eif3j,Rps4x,Rps27a |
| R-RNO-8953854 | Metabolism of RNA | -9,15233 | -6,613 | 97/510 | Rps29,Anp32a,Rpl32,Rpsa,Rpl9,Rps7,Rps14,Rps15,Rps17,Rps3a,Psmb5,Psmb6,Psmb3,Psmc3,Ubc,Rplp0,Rpl4,Rpl27,Rpl24,Rpl30,Adar,Rpl5,Rpl13,Rpl18,Rpl19,Rpl22,Rpl36al,Rpl37,Rps10,Rps21,Rps24,Psmd4,Polr2f,Sf3b1,Xpo1,Psmb7,Polr2g,Rpl6,Rps20,Rps23,Rps3,Rps16,Rplp1,Rps13,Prkcd,Khsrp,Casc3,Cnot6,Prpf8,Rpl18a,Lsm4,Psmb10,Psmd8,Rpl36a,Rps18,Rbm8a,Lsm8,Lsm3,Ddx47,Fcf1,Sf3a2,Cnot2,Rpl3,Xrn1,Thoc7,Exosc5,Hnrnph2,Upf3b,Imp3,Ddx21,Bysl,Zc3h11a,Thoc5,Dcp1a,Upf3a,Snrnp70,Patl1,Rpl22l1,Lsm7,Dcaf13,Rrp9,Cpsf7,Pan2,Psme4,Rpl12,Gtf2h5,Snrpd2,Sf3b5,Snrpg,Rps27l,Lsm2,Rpl38,Polr2e,Rps28,Polr2k,Rps4x,Rps27a |
| GO:0002181 | cytoplasmic translation | -9,06456 | -6,541 | 35/113 | Rps29,Rps26,Rpl32,Rpsa,Rpl9,Rplp0,Rpl24,Rpl30,Rpl18,Rpl19,Rpl22,Rps21,Rps2,Rpl6,Rps23,Rpl41,Rplp1,Rwdd1,Rpl26,Rpl18a,Eif3k,Rpl36a,Eif3f,Eif3m,Eif2s2,Mcts1,Cpeb4,Rpl13a,Rpl22l1,Zc3h15,Dph3,Denr,Rpl38,Rps28,Eif3j |
| R-RNO-72649 | Translation initiation complex formation | -8,0977 | -5,663 | 24/65 | Rps29,Rpsa,Rps7,Rps14,Rps15,Rps17,Rps3a,Rps10,Rps21,Rps24,Rps20,Rps23,Rps3,Rps16,Rps13,Eif3f,Rps18,Eif3m,Eif2s2,Rps27l,Rps28,Eif3j,Rps4x,Rps27a |
| R-RNO-72662 | Activation of the mRNA upon binding of the cap-binding complex and eIFs, and subsequent binding to 43S | -7,94512 | -5,529 | 24/66 | Rps29,Rpsa,Rps7,Rps14,Rps15,Rps17,Rps3a,Rps10,Rps21,Rps24,Rps20,Rps23,Rps3,Rps16,Rps13,Eif3f,Rps18,Eif3m,Eif2s2,Rps27l,Rps28,Eif3j,Rps4x,Rps27a |
| R-RNO-72702 | Ribosomal scanning and start codon recognition | -7,94512 | -5,529 | 24/66 | Rps29,Rpsa,Rps7,Rps14,Rps15,Rps17,Rps3a,Rps10,Rps21,Rps24,Rps20,Rps23,Rps3,Rps16,Rps13,Eif3f,Rps18,Eif3m,Eif2s2,Rps27l,Rps28,Eif3j,Rps4x,Rps27a |
| ko00190 | Oxidative phosphorylation | -15,7463 | -12,453 | 50/138 | Cox6a1,Ndufa5,Cox4i1,Cox7a2,Atp5mc1,Atp6v0a1,Cox6c,Cox17,Atp6v0e1,Atp5mc3,Atp6v0a2,Atp5me,Cox8a,Atp5po,Atp5f1e,Atp5f1d,Cox5a,Ndufb4,Uqcrfs1,Ndufa2,Ndufc2,Ndufab1,Ndufs8,Ndufb8,Ndufb5,Ndufs3,Ndufa8,Atp6v0a4,Ndufb6,Atp6v0b,Ndufb11,Ndufa7,Ndufb9,Atp5mg,Ndufa11,Ndufb3,Ppa2,Ndufa6,Ndufb7,Ndufb2,Ndufs5,Ndufs7,Uqcrh,Uqcrq,Atp5pd,Cox6b2,Uqcr10,Uqcr11,Cox7c,Ndufa13,Gpx1,Grm5,Sod1,Ap2s1,Gnaq,Polr2f,Ppargc1a,Dctn4,Cltb,Polr2g,Ap2b1,Dnah1,Ap2a1,Nrf1,Polr2e,Polr2k,Atp5if1,Dmac2l,Gnai1,Park7,Prkacb,Ube2l6,Ube2j1,Ube2j2,Etfb,Timmdc1,Cox16,Cacna1c,Grin2a,Lpl,Ppp3cb,Psen1,Bace1,Adam10,Cacna1d,Hsd17b10,Atf6,Akt2,Pik3r1,Akt3,Mapk9,Gsk3a,Atf4,Pik3cb,Pik3ca,Itch,Traf2,Mdh2,Cs,Adhfe1,Coa8,Uqcc2,Coa3,Uqcc3,Ndufaf8,Slc25a33,Antkmt,Iscu,Slc25a23,Fxn,Acadm,Pomc,Hsd11b1,Acadsb,Fdx1,Htr2a,Acox1,Mfn2,Cyba,Hmgcl,Cyb5b,Taldo1,Slc4a4,Mt3,Pfkfb3,Kif1b,Galk1,Zbtb20,Bloc1s1,Pgls,Eno4,Nop53,Cisd1,Gfpt1,H6pd,Eif6,Fdx2,Entpd5,Mdh1b,Phkb,Xdh,Tcf7l2,Ak3,Tmsb4x,Dnm1l,Fis1,Nudt2,Trem2,Guk1,Taf3 |
| ko00190 | Oxidative phosphorylation | -15,7463 | -12,453 | 50/138 | Cox6a1,Ndufa5,Cox4i1,Cox7a2,Atp5mc1,Atp6v0a1,Cox6c,Cox17,Atp6v0e1,Atp5mc3,Atp6v0a2,Atp5me,Cox8a,Atp5po,Atp5f1e,Atp5f1d,Cox5a,Ndufb4,Uqcrfs1,Ndufa2,Ndufc2,Ndufab1,Ndufs8,Ndufb8,Ndufb5,Ndufs3,Ndufa8,Atp6v0a4,Ndufb6,Atp6v0b,Ndufb11,Ndufa7,Ndufb9,Atp5mg,Ndufa11,Ndufb3,Ppa2,Ndufa6,Ndufb7,Ndufb2,Ndufs5,Ndufs7,Uqcrh,Uqcrq,Atp5pd,Cox6b2,Uqcr10,Uqcr11,Cox7c,Ndufa13 |
| rno00190 | Oxidative phosphorylation | -15,7463 | -12,453 | 50/138 | Cox6a1,Ndufa5,Cox4i1,Cox7a2,Atp5mc1,Atp6v0a1,Cox6c,Cox17,Atp6v0e1,Atp5mc3,Atp6v0a2,Atp5me,Cox8a,Atp5po,Atp5f1e,Atp5f1d,Cox5a,Ndufb4,Uqcrfs1,Ndufa2,Ndufc2,Ndufab1,Ndufs8,Ndufb8,Ndufb5,Ndufs3,Ndufa8,Atp6v0a4,Ndufb6,Atp6v0b,Ndufb11,Ndufa7,Ndufb9,Atp5mg,Ndufa11,Ndufb3,Ppa2,Ndufa6,Ndufb7,Ndufb2,Ndufs5,Ndufs7,Uqcrh,Uqcrq,Atp5pd,Cox6b2,Uqcr10,Uqcr11,Cox7c,Ndufa13 |
| ko05016 | Huntington's disease | -13,1193 | -10,171 | 57/195 | Gpx1,Grm5,Sod1,Cox6a1,Ndufa5,Cox4i1,Cox7a2,Atp5mc1,Cox6c,Ap2s1,Gnaq,Polr2f,Ppargc1a,Dctn4,Atp5mc3,Cltb,Polr2g,Ap2b1,Cox8a,Dnah1,Atp5po,Atp5f1e,Atp5f1d,Cox5a,Ndufb4,Uqcrfs1,Ndufa2,Ndufc2,Ndufab1,Ndufs8,Ndufb8,Ndufb5,Ndufs3,Ndufa8,Ndufb6,Ndufb11,Ndufa7,Ndufb9,Ndufa11,Ndufb3,Ap2a1,Nrf1,Ndufa6,Ndufb7,Ndufb2,Ndufs5,Ndufs7,Uqcrh,Uqcrq,Atp5pd,Cox6b2,Uqcr10,Uqcr11,Polr2e,Cox7c,Polr2k,Ndufa13 |
| rno05016 | Huntington's disease | -13,1193 | -10,171 | 57/195 | Gpx1,Grm5,Sod1,Cox6a1,Ndufa5,Cox4i1,Cox7a2,Atp5mc1,Cox6c,Ap2s1,Gnaq,Polr2f,Ppargc1a,Dctn4,Atp5mc3,Cltb,Polr2g,Ap2b1,Cox8a,Dnah1,Atp5po,Atp5f1e,Atp5f1d,Cox5a,Ndufb4,Uqcrfs1,Ndufa2,Ndufc2,Ndufab1,Ndufs8,Ndufb8,Ndufb5,Ndufs3,Ndufa8,Ndufb6,Ndufb11,Ndufa7,Ndufb9,Ndufa11,Ndufb3,Ap2a1,Nrf1,Ndufa6,Ndufb7,Ndufb2,Ndufs5,Ndufs7,Uqcrh,Uqcrq,Atp5pd,Cox6b2,Uqcr10,Uqcr11,Polr2e,Cox7c,Polr2k,Ndufa13 |
| WP59 | Electron Transport Chain | -12,8739 | -9,975 | 36/91 | Cox6a1,Atp5if1,Ndufa5,Cox4i1,Cox7a2,Cox6c,Cox17,Atp5mc3,Atp5me,Cox8a,Atp5po,Atp5f1d,Cox5a,Ndufb4,Uqcrfs1,Ndufa2,Ndufc2,Ndufab1,Ndufs8,Ndufb8,Ndufb5,Ndufs3,Ndufa8,Ndufb6,Ndufa7,Ndufb9,Atp5mg,Ndufa6,Ndufb7,Ndufb2,Dmac2l,Ndufs7,Uqcrh,Uqcrq,Atp5pd,Uqcr10 |
| rno05012 | Parkinson's disease | -12,5012 | -9,620 | 47/147 | Cox6a1,Ndufa5,Gnai1,Cox4i1,Cox7a2,Atp5mc1,Cox6c,Atp5mc3,Park7,Cox8a,Atp5po,Atp5f1e,Atp5f1d,Cox5a,Ndufb4,Uqcrfs1,Ndufa2,Ndufc2,Ndufab1,Prkacb,Ndufs8,Ndufb8,Ndufb5,Ube2l6,Ndufs3,Ndufa8,Ube2j1,Ndufb6,Ube2j2,Ndufb11,Ndufa7,Ndufb9,Ndufa11,Ndufb3,Ndufa6,Ndufb7,Ndufb2,Ndufs5,Ndufs7,Uqcrh,Uqcrq,Atp5pd,Cox6b2,Uqcr10,Uqcr11,Cox7c,Ndufa13 |
| R-RNO-163200 | Respiratory electron transport, ATP synthesis by chemiosmotic coupling, and heat production by uncoupling proteins. | -12,2298 | -9,365 | 39/109 | Cox6a1,Ndufa5,Cox4i1,Atp5mc1,Atp5me,Cox8a,Atp5po,Atp5f1d,Cox5a,Ndufb4,Uqcrfs1,Ndufa2,Etfb,Ndufc2,Ndufab1,Ndufs8,Ndufb8,Ndufb5,Ndufs3,Ndufb6,Ndufb11,Ndufa7,Ndufb9,Atp5mg,Ndufa11,Ndufb3,Timmdc1,Ndufa6,Ndufb7,Ndufb2,Ndufs5,Dmac2l,Ndufs7,Uqcrh,Uqcrq,Atp5pd,Cox7c,Cox16,Ndufa13 |
| ko05010 | Alzheimer's disease | -12,0164 | -9,184 | 52/178 | Cacna1c,Grin2a,Lpl,Ppp3cb,Cox6a1,Ndufa5,Psen1,Bace1,Cox4i1,Cox7a2,Adam10,Cacna1d,Atp5mc1,Cox6c,Hsd17b10,Gnaq,Atp5mc3,Cox8a,Atp5po,Atp5f1e,Atp5f1d,Cox5a,Ndufb4,Uqcrfs1,Ndufa2,Ndufc2,Ndufab1,Ndufs8,Ndufb8,Ndufb5,Ndufs3,Ndufa8,Ndufb6,Ndufb11,Ndufa7,Ndufb9,Ndufa11,Ndufb3,Atf6,Ndufa6,Ndufb7,Ndufb2,Ndufs5,Ndufs7,Uqcrh,Uqcrq,Atp5pd,Cox6b2,Uqcr10,Uqcr11,Cox7c,Ndufa13 |
| rno05010 | Alzheimer's disease | -12,0164 | -9,184 | 52/178 | Cacna1c,Grin2a,Lpl,Ppp3cb,Cox6a1,Ndufa5,Psen1,Bace1,Cox4i1,Cox7a2,Adam10,Cacna1d,Atp5mc1,Cox6c,Hsd17b10,Gnaq,Atp5mc3,Cox8a,Atp5po,Atp5f1e,Atp5f1d,Cox5a,Ndufb4,Uqcrfs1,Ndufa2,Ndufc2,Ndufab1,Ndufs8,Ndufb8,Ndufb5,Ndufs3,Ndufa8,Ndufb6,Ndufb11,Ndufa7,Ndufb9,Ndufa11,Ndufb3,Atf6,Ndufa6,Ndufb7,Ndufb2,Ndufs5,Ndufs7,Uqcrh,Uqcrq,Atp5pd,Cox6b2,Uqcr10,Uqcr11,Cox7c,Ndufa13 |
| WP1283 | Oxidative phosphorylation | -10,7502 | -8,002 | 26/59 | Ndufa5,Atp5mc1,Atp5mc3,Atp5me,Atp5po,Atp5f1e,Atp5f1d,Ndufa2,Ndufc2,Ndufab1,Ndufs8,Ndufb8,Ndufb5,Ndufs3,Ndufa8,Ndufb6,Ndufa7,Ndufb9,Atp5mg,Ndufa11,Ndufa6,Ndufb7,Ndufb2,Ndufs5,Dmac2l,Ndufs7 |
| ko04932 | Non-alcoholic fatty liver disease (NAFLD) | -10,3904 | -7,702 | 45/155 | Akt2,Cox6a1,Ndufa5,Pik3r1,Akt3,Cox4i1,Cox7a2,Mapk9,Gsk3a,Cox6c,Atf4,Pik3cb,Pik3ca,Cox8a,Cox5a,Ndufb4,Uqcrfs1,Ndufa2,Ndufc2,Ndufab1,Ndufs8,Ndufb8,Ndufb5,Ndufs3,Ndufa8,Ndufb6,Ndufb11,Ndufa7,Ndufb9,Ndufa11,Ndufb3,Itch,Traf2,Ndufa6,Ndufb7,Ndufb2,Ndufs5,Ndufs7,Uqcrh,Uqcrq,Cox6b2,Uqcr10,Uqcr11,Cox7c,Ndufa13 |
| rno04932 | Non-alcoholic fatty liver disease (NAFLD) | -10,3904 | -7,702 | 45/155 | Akt2,Cox6a1,Ndufa5,Pik3r1,Akt3,Cox4i1,Cox7a2,Mapk9,Gsk3a,Cox6c,Atf4,Pik3cb,Pik3ca,Cox8a,Cox5a,Ndufb4,Uqcrfs1,Ndufa2,Ndufc2,Ndufab1,Ndufs8,Ndufb8,Ndufb5,Ndufs3,Ndufa8,Ndufb6,Ndufb11,Ndufa7,Ndufb9,Ndufa11,Ndufb3,Itch,Traf2,Ndufa6,Ndufb7,Ndufb2,Ndufs5,Ndufs7,Uqcrh,Uqcrq,Cox6b2,Uqcr10,Uqcr11,Cox7c,Ndufa13 |
| R-RNO-611105 | Respiratory electron transport | -9,931 | -7,285 | 32/91 | Cox6a1,Ndufa5,Cox4i1,Cox8a,Cox5a,Ndufb4,Uqcrfs1,Ndufa2,Etfb,Ndufc2,Ndufab1,Ndufs8,Ndufb8,Ndufb5,Ndufs3,Ndufb6,Ndufb11,Ndufa7,Ndufb9,Ndufa11,Ndufb3,Timmdc1,Ndufa6,Ndufb7,Ndufb2,Ndufs5,Ndufs7,Uqcrh,Uqcrq,Cox7c,Cox16,Ndufa13 |
| R-RNO-1428517 | The citric acid (TCA) cycle and respiratory electron transport | -9,10818 | -6,577 | 42/151 | Cox6a1,Ndufa5,Cox4i1,Atp5mc1,Mdh2,Atp5me,Cs,Cox8a,Atp5po,Atp5f1d,Cox5a,Ndufb4,Uqcrfs1,Ndufa2,Etfb,Ndufc2,Ndufab1,Ndufs8,Ndufb8,Ndufb5,Ndufs3,Ndufb6,Ndufb11,Ndufa7,Ndufb9,Atp5mg,Ndufa11,Ndufb3,Timmdc1,Ndufa6,Ndufb7,Ndufb2,Adhfe1,Ndufs5,Dmac2l,Ndufs7,Uqcrh,Uqcrq,Atp5pd,Cox7c,Cox16,Ndufa13 |
| GO:0033108 | mitochondrial respiratory chain complex assembly | -8,57609 | -6,096 | 29/86 | Ndufa5,Cox17,Ndufb4,Uqcrfs1,Ndufa2,Ndufc2,Ndufab1,Ndufs8,Ndufb8,Ndufb5,Ndufa8,Ndufb6,Ndufb11,Coa8,Ndufb9,Ndufa11,Ndufb3,Ndufa6,Ndufb7,Uqcc2,Ndufb2,Ndufs5,Ndufs7,Coa3,Uqcc3,Ndufaf8,Slc25a33,Cox16,Ndufa13 |
| R-RNO-6799198 | Complex I biogenesis | -8,23153 | -5,772 | 22/55 | Ndufa5,Ndufb4,Ndufa2,Ndufc2,Ndufab1,Ndufs8,Ndufb8,Ndufb5,Ndufs3,Ndufb6,Ndufb11,Ndufa7,Ndufb9,Ndufa11,Ndufb3,Timmdc1,Ndufa6,Ndufb7,Ndufb2,Ndufs5,Ndufs7,Ndufa13 |
| GO:0010257 | NADH dehydrogenase complex assembly | -8,15586 | -5,715 | 21/51 | Ndufa5,Ndufb4,Ndufa2,Ndufc2,Ndufab1,Ndufs8,Ndufb8,Ndufb5,Ndufa8,Ndufb6,Ndufb11,Ndufb9,Ndufa11,Ndufb3,Ndufa6,Ndufb7,Ndufb2,Ndufs5,Ndufs7,Ndufaf8,Ndufa13 |
| GO:0032981 | mitochondrial respiratory chain complex I assembly | -8,15586 | -5,715 | 21/51 | Ndufa5,Ndufb4,Ndufa2,Ndufc2,Ndufab1,Ndufs8,Ndufb8,Ndufb5,Ndufa8,Ndufb6,Ndufb11,Ndufb9,Ndufa11,Ndufb3,Ndufa6,Ndufb7,Ndufb2,Ndufs5,Ndufs7,Ndufaf8,Ndufa13 |
| GO:0006119 | oxidative phosphorylation | -6,89402 | -4,559 | 33/123 | Cox6a1,Cox4i1,Cox7a2,Cox6c,Park7,Atp5me,Cox8a,Atp5po,Atp5f1e,Atp5f1d,Cox5a,Antkmt,Iscu,Uqcrfs1,Ndufc2,Ndufs8,Ndufb8,Ndufa8,Ndufb6,Ndufa7,Ndufb9,Atp5mg,Slc25a23,Uqcc2,Uqcrh,Uqcrq,Fxn,Atp5pd,Cox6b2,Uqcr10,Uqcc3,Slc25a33,Cox7c |
| GO:0006091 | generation of precursor metabolites and energy | -6,47667 | -4,210 | 79/440 | Acadm,Pomc,Hsd11b1,Akt2,Cox6a1,Ndufa5,Acadsb,Fdx1,Psen1,Cox4i1,Cox7a2,Htr2a,Acox1,Gsk3a,Cox6c,Mfn2,Cyba,Hmgcl,Cyb5b,Mdh2,Ppargc1a,Taldo1,Slc4a4,Cox17,Mt3,Pfkfb3,Park7,Kif1b,Atp5me,Cs,Pik3ca,Cox8a,Atp5po,Atp5f1e,Atp5f1d,Cox5a,Antkmt,Galk1,Zbtb20,Iscu,Bloc1s1,Pgls,Uqcrfs1,Eno4,Nop53,Etfb,Ndufc2,Ndufs8,Ndufb8,Cisd1,Ndufa8,Gfpt1,Ndufb6,H6pd,Coa8,Ndufa7,Ndufb9,Atp5mg,Slc25a23,Ndufb3,Eif6,Fdx2,Entpd5,Mdh1b,Phkb,Uqcc2,Ndufs7,Uqcrh,Xdh,Uqcrq,Fxn,Atp5pd,Cox6b2,Tcf7l2,Uqcr10,Uqcc3,Uqcr11,Slc25a33,Cox7c |
| GO:0022900 | electron transport chain | -6,46204 | -4,200 | 32/122 | Cox6a1,Ndufa5,Acadsb,Fdx1,Cox4i1,Cox7a2,Cox6c,Cyba,Cyb5b,Ppargc1a,Park7,Cox8a,Cox5a,Iscu,Uqcrfs1,Etfb,Ndufc2,Ndufs8,Ndufb8,Ndufa8,Ndufb6,Ndufa7,Ndufb9,Ndufb3,Fdx2,Uqcrh,Xdh,Uqcrq,Uqcr10,Uqcc3,Uqcr11,Cox7c |
| GO:0046034 | ATP metabolic process | -6,32916 | -4,071 | 55/272 | Cox6a1,Atp5if1,Ak3,Psen1,Cox4i1,Cox7a2,Htr2a,Atp5mc1,Cox6c,Tmsb4x,Ppargc1a,Slc4a4,Dnm1l,Atp5mc3,Pfkfb3,Park7,Atp5me,Cox8a,Atp5po,Atp5f1e,Atp5f1d,Cox5a,Antkmt,Galk1,Zbtb20,Fis1,Iscu,Uqcrfs1,Eno4,Ndufc2,Ndufs8,Ndufb8,Ndufa8,Ndufb6,Nudt2,Ndufa7,Ndufb9,Atp5mg,Slc25a23,Trem2,Guk1,Eif6,Entpd5,Uqcc2,Dmac2l,Uqcrh,Uqcrq,Fxn,Atp5pd,Cox6b2,Uqcr10,Uqcc3,Slc25a33,Cox7c,Taf3 |
| GO:0042775 | mitochondrial ATP synthesis coupled electron transport | -4,76269 | -2,743 | 18/60 | Cox6a1,Cox4i1,Park7,Cox5a,Iscu,Uqcrfs1,Ndufc2,Ndufs8,Ndufb8,Ndufa8,Ndufb6,Ndufa7,Ndufb9,Uqcrh,Uqcrq,Uqcr10,Uqcc3,Cox7c |
| GO:0042773 | ATP synthesis coupled electron transport | -4,15051 | -2,250 | 18/66 | Cox6a1,Cox4i1,Park7,Cox5a,Iscu,Uqcrfs1,Ndufc2,Ndufs8,Ndufb8,Ndufa8,Ndufb6,Ndufa7,Ndufb9,Uqcrh,Uqcrq,Uqcr10,Uqcc3,Cox7c |
| GO:0045333 | cellular respiration | -3,8621 | -2,016 | 32/160 | Cox6a1,Ndufa5,Cox4i1,Mfn2,Mdh2,Ppargc1a,Park7,Cs,Pik3ca,Atp5f1d,Cox5a,Iscu,Bloc1s1,Uqcrfs1,Nop53,Ndufc2,Ndufs8,Ndufb8,Cisd1,Ndufa8,Ndufb6,Ndufa7,Ndufb9,Slc25a23,Mdh1b,Ndufs7,Uqcrh,Uqcrq,Fxn,Uqcr10,Uqcc3,Cox7c |
| GO:0022904 | respiratory electron transport chain | -3,80331 | -1,963 | 20/82 | Cox6a1,Ndufa5,Cox4i1,Ppargc1a,Park7,Cox5a,Iscu,Uqcrfs1,Ndufc2,Ndufs8,Ndufb8,Ndufa8,Ndufb6,Ndufa7,Ndufb9,Uqcrh,Uqcrq,Uqcr10,Uqcc3,Cox7c |
| GO:0006120 | mitochondrial electron transport, NADH to ubiquinone | -3,611 | -1,820 | 9/23 | Park7,Iscu,Ndufc2,Ndufs8,Ndufb8,Ndufa8,Ndufb6,Ndufa7,Ndufb9 |
| GO:0015980 | energy derivation by oxidation of organic compounds | -2,66209 | -1,102 | 40/248 | Acadm,Pomc,Akt2,Cox6a1,Ndufa5,Cox4i1,Gsk3a,Mfn2,Mdh2,Ppargc1a,Mt3,Park7,Cs,Pik3ca,Atp5f1d,Cox5a,Iscu,Bloc1s1,Uqcrfs1,Nop53,Ndufc2,Ndufs8,Ndufb8,Cisd1,Ndufa8,Gfpt1,Ndufb6,Ndufa7,Ndufb9,Slc25a23,Mdh1b,Phkb,Ndufs7,Uqcrh,Uqcrq,Fxn,Tcf7l2,Uqcr10,Uqcc3,Cox7c |
| GO:0099504 | synaptic vesicle cycle | -15,3937 | -12,194 | 70/245 | Mx1,Ppp3cb,Prkca,Prkcg,Syp,Stx1b,Htr1b,P2ry1,Camk2a,Syt1,Cadps,Psen1,Prkce,Bace1,Htr2a,Pacsin1,Cacna1d,Cspg5,Chrnb2,Pclo,Cacnb4,Syt7,Stx7,Syt9,Syt11,Amph,Fgf14,Rab5a,Unc13a,Unc13b,Scamp5,Snap91,Dnajc5,Stxbp5,Grm7,Stx3,Doc2b,Sh3gl1,Apba1,Apba2,Ctnnb1,Clcn3,Synj1,Picalm,Magi2,Dnm1l,Cltb,Nlgn1,Sh3gl2,Rims2,Bin1,Nlgn2,Ap2b1,Trim9,Nlgn3,Dnm3,Cacna1b,Unc13c,Tbc1d24,Stxbp5l,Adcy1,Pcdh17,Fcho2,Dnajc6,Ston2,Pip5k1c,Slc4a8,Ppfia2,Napb,Dtnbp1,Arc,Grip1,Akap5,Efnb2,Htr2c,Nsf,Lin7b,Lin7c,Stxbp3,Kcnc3,Ap3m2,Rab15,Cacna1c,Anxa3,Syngr1,Rasgrp1,Vamp3,Cacna1i,Kit,Arf1,Syt13,Lat,Vamp8,Gab2,Pi4k2a,Cacna1h,Ap1g1,Rab11fip2,Plek,Agt,Egfr,Egr1,Gfap,Grin2a,Grm5,Jak2,Ncam1,Dbi,Cntn2,Ptprs,Slc1a1,Tnr,Pde4a,Gnai1,S100b,Adora1,S1pr2,Slc6a6,Ptgs2,Prkar2a,Kcnj10,Gria1,L1cam,Grik2,Synpo,Lgmn,Pcdh8,Shank1,Grid2,Atf4,Bhlhe40,Rpl22,Calb1,Kalrn,Slc24a2,Mpp2,Dlgap2,Slc8a2,Slc8a3,Cacng3,Cacng4,Fabp5,Cabp1,Shank2,Camk2g,Clstn3,Syngap1,Lgi1,Npas4,Nptx2,Tubb2b,Ntng1,Slc4a10,Zmynd8,Grik3,Cacna2d2,Sorcs2,Neto1,Neto2,Neurl1,Abl1,Tyrobp,Bscl2,Eif4ebp2,Dgke,Crtc1,Iqsec2,Shisa7,Gad2,Pde1b,Arhgef11,Slc6a11,Abat,Park7,Sv2b,Gabra2,Sdc4,Myo5a,Rab12,Exoc4,Stx17,Arfgef2,Exoc1,Vps4b,Stxbp6,Chmp2a,B2m,C2,C3,Sod1,Clu,Akt2,Apoc1,Arrb1,Tub,Cd63,Map2k2,Cyba,Ptpn23,Cdh13,Commd1,Use1,Trem2,Camk1d,Arhgap21,Mib1,Ap2a1,Inpp5f,Pacsin3,Cd2ap,Pgap1,Usp7,Myo18a,Ankrd27,Tbc1d20,Bicd1,Ccl19 |
| GO:0099504 | synaptic vesicle cycle | -15,3937 | -12,194 | 70/245 | Mx1,Ppp3cb,Prkca,Prkcg,Syp,Stx1b,Htr1b,P2ry1,Camk2a,Syt1,Cadps,Psen1,Prkce,Bace1,Htr2a,Pacsin1,Cacna1d,Cspg5,Chrnb2,Pclo,Cacnb4,Syt7,Stx7,Syt9,Syt11,Amph,Fgf14,Rab5a,Unc13a,Unc13b,Scamp5,Snap91,Dnajc5,Stxbp5,Grm7,Stx3,Doc2b,Sh3gl1,Apba1,Apba2,Ctnnb1,Clcn3,Synj1,Picalm,Magi2,Dnm1l,Cltb,Nlgn1,Sh3gl2,Rims2,Bin1,Nlgn2,Ap2b1,Trim9,Nlgn3,Dnm3,Cacna1b,Unc13c,Tbc1d24,Stxbp5l,Adcy1,Pcdh17,Fcho2,Dnajc6,Ston2,Pip5k1c,Slc4a8,Ppfia2,Napb,Dtnbp1 |
| GO:0099003 | vesicle-mediated transport in synapse | -14,8813 | -11,748 | 74/273 | Mx1,Ppp3cb,Prkca,Prkcg,Syp,Stx1b,Htr1b,P2ry1,Camk2a,Syt1,Cadps,Psen1,Prkce,Bace1,Htr2a,Pacsin1,Cacna1d,Cspg5,Chrnb2,Arc,Pclo,Cacnb4,Syt7,Stx7,Syt9,Syt11,Amph,Fgf14,Rab5a,Unc13a,Unc13b,Scamp5,Snap91,Dnajc5,Stxbp5,Grm7,Stx3,Doc2b,Sh3gl1,Apba1,Apba2,Grip1,Ctnnb1,Clcn3,Synj1,Picalm,Magi2,Dnm1l,Cltb,Nlgn1,Sh3gl2,Rims2,Bin1,Nlgn2,Ap2b1,Trim9,Akap5,Nlgn3,Dnm3,Cacna1b,Unc13c,Tbc1d24,Stxbp5l,Adcy1,Pcdh17,Efnb2,Fcho2,Dnajc6,Ston2,Pip5k1c,Slc4a8,Ppfia2,Napb,Dtnbp1 |
| GO:0007269 | neurotransmitter secretion | -10,0873 | -7,421 | 53/204 | Prkca,Prkcg,Syp,Stx1b,Htr1b,Htr2c,P2ry1,Camk2a,Syt1,Cadps,Psen1,Prkce,Bace1,Htr2a,Cacna1d,Cspg5,Chrnb2,Pclo,Cacnb4,Syt7,Nsf,Lin7b,Lin7c,Stx7,Syt9,Rab5a,Unc13a,Unc13b,Snap91,Stxbp5,Grm7,Stx3,Doc2b,Apba1,Apba2,Synj1,Stxbp3,Dnm1l,Cltb,Nlgn1,Rims2,Kcnc3,Ap3m2,Trim9,Cacna1b,Unc13c,Stxbp5l,Rab15,Adcy1,Slc4a8,Ppfia2,Napb,Dtnbp1 |
| GO:0099643 | signal release from synapse | -10,0873 | -7,421 | 53/204 | Prkca,Prkcg,Syp,Stx1b,Htr1b,Htr2c,P2ry1,Camk2a,Syt1,Cadps,Psen1,Prkce,Bace1,Htr2a,Cacna1d,Cspg5,Chrnb2,Pclo,Cacnb4,Syt7,Nsf,Lin7b,Lin7c,Stx7,Syt9,Rab5a,Unc13a,Unc13b,Snap91,Stxbp5,Grm7,Stx3,Doc2b,Apba1,Apba2,Synj1,Stxbp3,Dnm1l,Cltb,Nlgn1,Rims2,Kcnc3,Ap3m2,Trim9,Cacna1b,Unc13c,Stxbp5l,Rab15,Adcy1,Slc4a8,Ppfia2,Napb,Dtnbp1 |
| GO:0045055 | regulated exocytosis | -9,69625 | -7,076 | 65/283 | Cacna1c,Ppp3cb,Prkca,Prkcg,Syp,Stx1b,Htr1b,P2ry1,Anxa3,Camk2a,Syt1,Cadps,Psen1,Syngr1,Prkce,Bace1,Rasgrp1,Vamp3,Htr2a,Cacna1d,Cspg5,Chrnb2,Pclo,Cacna1i,Cacnb4,Syt7,Stx7,Syt9,Syt11,Kit,Arf1,Rab5a,Unc13a,Unc13b,Scamp5,Syt13,Stxbp5,Lat,Grm7,Stx3,Doc2b,Apba1,Apba2,Vamp8,Gab2,Synj1,Stxbp3,Dnm1l,Pi4k2a,Cacna1h,Nlgn1,Rims2,Trim9,Ap1g1,Cacna1b,Unc13c,Stxbp5l,Rab15,Adcy1,Rab11fip2,Slc4a8,Ppfia2,Plek,Napb,Dtnbp1 |
| GO:0099177 | regulation of trans-synaptic signaling | -9,67698 | -7,070 | 117/643 | Agt,Egfr,Egr1,Gfap,Grin2a,Grm5,Jak2,Ncam1,Prkca,Prkcg,Syp,Stx1b,Dbi,Htr1b,Htr2c,P2ry1,Cntn2,Camk2a,Ptprs,Slc1a1,Tnr,Pde4a,Gnai1,Syt1,S100b,Psen1,Syngr1,Adora1,Prkce,Bace1,S1pr2,Slc6a6,Ptgs2,Htr2a,Prkar2a,Cacna1d,Kcnj10,Cspg5,Gria1,L1cam,Chrnb2,Grik2,Arc,Cacnb4,Syt7,Synpo,Fgf14,Lgmn,Kit,Arf1,Rab5a,Unc13a,Unc13b,Pcdh8,Shank1,Grid2,Atf4,Bhlhe40,Stxbp5,Grm7,Rpl22,Stx3,Sh3gl1,Apba1,Apba2,Calb1,Kalrn,Slc24a2,Mpp2,Dnm1l,Nlgn1,Dlgap2,Rims2,Nlgn2,Kcnc3,Slc8a2,Slc8a3,Cacng3,Cacng4,Fabp5,Akap5,Cabp1,Shank2,Camk2g,Nlgn3,Clstn3,Syngap1,Lgi1,Cacna1b,Npas4,Unc13c,Tbc1d24,Stxbp5l,Nptx2,Tubb2b,Ntng1,Slc4a10,Zmynd8,Grik3,Cacna2d2,Sorcs2,Adcy1,Pcdh17,Neto1,Neto2,Neurl1,Abl1,Slc4a8,Tyrobp,Bscl2,Eif4ebp2,Ppfia2,Dgke,Dtnbp1,Crtc1,Iqsec2,Shisa7 |
| GO:0016079 | synaptic vesicle exocytosis | -9,56335 | -6,965 | 43/152 | Prkca,Prkcg,Syp,Stx1b,Htr1b,P2ry1,Camk2a,Syt1,Cadps,Psen1,Prkce,Bace1,Htr2a,Cacna1d,Cspg5,Chrnb2,Pclo,Cacnb4,Syt7,Stx7,Syt9,Rab5a,Unc13a,Unc13b,Stxbp5,Grm7,Stx3,Doc2b,Apba1,Apba2,Synj1,Dnm1l,Nlgn1,Rims2,Trim9,Cacna1b,Unc13c,Stxbp5l,Adcy1,Slc4a8,Ppfia2,Napb,Dtnbp1 |
| GO:0050804 | modulation of chemical synaptic transmission | -9,40733 | -6,827 | 116/642 | Agt,Egfr,Egr1,Gfap,Grin2a,Grm5,Jak2,Ncam1,Prkca,Prkcg,Syp,Stx1b,Dbi,Htr1b,Htr2c,P2ry1,Cntn2,Camk2a,Ptprs,Slc1a1,Tnr,Pde4a,Gnai1,Syt1,S100b,Psen1,Syngr1,Adora1,Prkce,Bace1,S1pr2,Slc6a6,Ptgs2,Htr2a,Prkar2a,Cacna1d,Kcnj10,Cspg5,Gria1,L1cam,Chrnb2,Grik2,Arc,Cacnb4,Syt7,Synpo,Fgf14,Lgmn,Kit,Arf1,Rab5a,Unc13a,Unc13b,Pcdh8,Shank1,Grid2,Atf4,Bhlhe40,Stxbp5,Grm7,Rpl22,Stx3,Sh3gl1,Apba1,Apba2,Calb1,Kalrn,Slc24a2,Mpp2,Dnm1l,Nlgn1,Dlgap2,Rims2,Nlgn2,Kcnc3,Slc8a2,Slc8a3,Cacng3,Cacng4,Akap5,Cabp1,Shank2,Camk2g,Nlgn3,Clstn3,Syngap1,Lgi1,Cacna1b,Npas4,Unc13c,Tbc1d24,Stxbp5l,Nptx2,Tubb2b,Ntng1,Slc4a10,Zmynd8,Grik3,Cacna2d2,Sorcs2,Adcy1,Pcdh17,Neto1,Neto2,Neurl1,Abl1,Slc4a8,Tyrobp,Bscl2,Eif4ebp2,Ppfia2,Dgke,Dtnbp1,Crtc1,Iqsec2,Shisa7 |
| GO:0001505 | regulation of neurotransmitter levels | -9,36967 | -6,806 | 63/275 | Gad2,Gfap,Prkca,Prkcg,Syp,Stx1b,Htr1b,Htr2c,P2ry1,Camk2a,Syt1,Cadps,Psen1,Prkce,Bace1,Htr2a,Pde1b,Cacna1d,Kcnj10,Cspg5,Chrnb2,Pclo,Cacnb4,Syt7,Nsf,Lin7b,Lin7c,Stx7,Syt9,Rab5a,Unc13a,Unc13b,Snap91,Arhgef11,Slc6a11,Stxbp5,Abat,Grm7,Stx3,Doc2b,Apba1,Apba2,Synj1,Stxbp3,Dnm1l,Cltb,Nlgn1,Rims2,Kcnc3,Park7,Sv2b,Ap3m2,Trim9,Cacna1b,Unc13c,Stxbp5l,Gabra2,Rab15,Adcy1,Slc4a8,Ppfia2,Napb,Dtnbp1 |
| GO:0006836 | neurotransmitter transport | -8,94864 | -6,440 | 60/262 | Gfap,Prkca,Prkcg,Syp,Stx1b,Htr1b,Htr2c,P2ry1,Camk2a,Syt1,Cadps,Psen1,Prkce,Bace1,Slc6a6,Htr2a,Cacna1d,Kcnj10,Cspg5,Chrnb2,Pclo,Cacnb4,Syt7,Nsf,Lin7b,Lin7c,Stx7,Syt9,Rab5a,Unc13a,Unc13b,Snap91,Slc6a11,Stxbp5,Grm7,Stx3,Doc2b,Apba1,Apba2,Synj1,Stxbp3,Dnm1l,Cltb,Nlgn1,Rims2,Kcnc3,Park7,Sv2b,Ap3m2,Trim9,Cacna1b,Unc13c,Stxbp5l,Gabra2,Rab15,Adcy1,Slc4a8,Ppfia2,Napb,Dtnbp1 |
| GO:0006887 | exocytosis | -8,2678 | -5,801 | 80/408 | Cacna1c,Ncam1,Ppp3cb,Prkca,Prkcg,Sdc4,Syp,Stx1b,Myo5a,Htr1b,P2ry1,Anxa3,Camk2a,Rab12,Syt1,Cadps,Psen1,Syngr1,Prkce,Bace1,Rasgrp1,Vamp3,Htr2a,Cacna1d,Cspg5,Chrnb2,Pclo,Cacna1i,Cacnb4,Syt7,Nsf,Lin7b,Lin7c,Stx7,Syt9,Syt11,Kit,Arf1,Rab5a,Unc13a,Unc13b,Scamp5,Syt13,Stxbp5,Lat,Grm7,Stx3,Doc2b,Apba1,Apba2,Vamp8,Gab2,Synj1,Stxbp3,Dnm1l,Pi4k2a,Cacna1h,Nlgn1,Exoc4,Rims2,Trim9,Ap1g1,Stx17,Cacna1b,Unc13c,Stxbp5l,Arfgef2,Rab15,Exoc1,Adcy1,Rab11fip2,Pip5k1c,Slc4a8,Vps4b,Stxbp6,Ppfia2,Plek,Chmp2a,Napb,Dtnbp1 |
| GO:0060627 | regulation of vesicle-mediated transport | -6,71618 | -4,428 | 103/614 | B2m,C2,C3,Ppp3cb,Prkca,Prkcg,Sdc4,Sod1,Syp,Clu,Stx1b,Myo5a,Htr1b,Akt2,P2ry1,Apoc1,Arrb1,Rab12,Tub,Syt1,Cadps,Cd63,Psen1,Prkce,Bace1,Htr2a,Pacsin1,Cacna1d,Cspg5,Chrnb2,Arc,Pclo,Cacna1i,Cacnb4,Map2k2,Syt7,Nsf,Syt9,Syt11,Amph,Arf1,Rab5a,Unc13b,Scamp5,Snap91,Cyba,Dnajc5,Syt13,Stxbp5,Grm7,Doc2b,Sh3gl1,Apba1,Apba2,Vamp8,Gab2,Synj1,Picalm,Magi2,Stxbp3,Dnm1l,Cacna1h,Nlgn1,Sh3gl2,Rims2,Bin1,Ptpn23,Ap2b1,Trim9,Akap5,Ap1g1,Cdh13,Cacna1b,Stxbp5l,Commd1,Use1,Rab15,Trem2,Adcy1,Efnb2,Camk1d,Arhgap21,Mib1,Ap2a1,Inpp5f,Pacsin3,Dnajc6,Ston2,Pip5k1c,Slc4a8,Cd2ap,Pgap1,Usp7,Myo18a,Vps4b,Ankrd27,Tbc1d20,Bicd1,Ccl19,Ppfia2,Chmp2a,Napb,Dtnbp1 |
| GO:1903305 | regulation of regulated secretory pathway | -6,25594 | -4,014 | 43/193 | Prkca,Prkcg,Syp,Htr1b,P2ry1,Syt1,Cadps,Prkce,Bace1,Htr2a,Cacna1d,Cspg5,Chrnb2,Cacna1i,Cacnb4,Syt7,Syt9,Syt11,Arf1,Rab5a,Scamp5,Syt13,Stxbp5,Grm7,Doc2b,Apba1,Apba2,Vamp8,Gab2,Stxbp3,Dnm1l,Cacna1h,Nlgn1,Rims2,Trim9,Ap1g1,Cacna1b,Stxbp5l,Rab15,Adcy1,Slc4a8,Ppfia2,Dtnbp1 |
| GO:2000300 | regulation of synaptic vesicle exocytosis | -6,2015 | -3,975 | 27/96 | Prkca,Prkcg,Syp,Htr1b,P2ry1,Syt1,Prkce,Bace1,Htr2a,Cacna1d,Cspg5,Chrnb2,Cacnb4,Rab5a,Stxbp5,Grm7,Apba1,Apba2,Dnm1l,Nlgn1,Rims2,Cacna1b,Stxbp5l,Adcy1,Slc4a8,Ppfia2,Dtnbp1 |
| GO:0017157 | regulation of exocytosis | -5,97557 | -3,761 | 52/258 | Prkca,Prkcg,Sdc4,Syp,Stx1b,Myo5a,Htr1b,P2ry1,Rab12,Syt1,Cadps,Prkce,Bace1,Htr2a,Cacna1d,Cspg5,Chrnb2,Pclo,Cacna1i,Cacnb4,Syt7,Nsf,Syt9,Syt11,Arf1,Rab5a,Unc13b,Scamp5,Syt13,Stxbp5,Grm7,Doc2b,Apba1,Apba2,Vamp8,Gab2,Stxbp3,Dnm1l,Cacna1h,Nlgn1,Rims2,Trim9,Ap1g1,Cacna1b,Stxbp5l,Rab15,Adcy1,Slc4a8,Vps4b,Ppfia2,Chmp2a,Dtnbp1 |
| GO:0046928 | regulation of neurotransmitter secretion | -5,65985 | -3,491 | 33/138 | Prkca,Prkcg,Syp,Stx1b,Htr1b,Htr2c,P2ry1,Camk2a,Syt1,Prkce,Bace1,Htr2a,Cacna1d,Cspg5,Chrnb2,Cacnb4,Rab5a,Unc13a,Stxbp5,Grm7,Apba1,Apba2,Dnm1l,Nlgn1,Rims2,Kcnc3,Cacna1b,Unc13c,Stxbp5l,Adcy1,Slc4a8,Ppfia2,Dtnbp1 |
| GO:0048167 | regulation of synaptic plasticity | -5,44564 | -3,322 | 62/340 | Agt,Egr1,Gfap,Grin2a,Grm5,Prkcg,Syp,Dbi,Cntn2,Camk2a,Slc1a1,Tnr,S100b,Psen1,Syngr1,Adora1,Ptgs2,Kcnj10,Gria1,Grik2,Arc,Syt7,Synpo,Fgf14,Lgmn,Kit,Arf1,Rab5a,Unc13a,Unc13b,Shank1,Grid2,Atf4,Bhlhe40,Stx3,Calb1,Kalrn,Slc24a2,Mpp2,Nlgn1,Rims2,Slc8a2,Slc8a3,Akap5,Cabp1,Shank2,Camk2g,Nlgn3,Syngap1,Npas4,Unc13c,Slc4a10,Sorcs2,Adcy1,Neto1,Neurl1,Abl1,Tyrobp,Eif4ebp2,Crtc1,Iqsec2,Shisa7 |
| GO:0051588 | regulation of neurotransmitter transport | -4,9957 | -2,954 | 34/154 | Gfap,Prkca,Prkcg,Syp,Stx1b,Htr1b,Htr2c,P2ry1,Camk2a,Syt1,Prkce,Bace1,Htr2a,Cacna1d,Cspg5,Chrnb2,Cacnb4,Rab5a,Unc13a,Stxbp5,Grm7,Apba1,Apba2,Dnm1l,Nlgn1,Rims2,Kcnc3,Cacna1b,Unc13c,Stxbp5l,Adcy1,Slc4a8,Ppfia2,Dtnbp1 |
| GO:0050806 | positive regulation of synaptic transmission | -4,08894 | -2,199 | 51/292 | Egfr,Egr1,Gfap,Grin2a,Prkcg,Stx1b,Dbi,Htr2c,Slc1a1,Tnr,Syt1,S100b,Adora1,Prkce,Ptgs2,Cacna1d,Gria1,Chrnb2,Grik2,Arc,Cacnb4,Lgmn,Unc13a,Shank1,Stx3,Calb1,Kalrn,Slc24a2,Mpp2,Dnm1l,Nlgn1,Rims2,Nlgn2,Slc8a2,Slc8a3,Cacng3,Cacng4,Akap5,Shank2,Nlgn3,Clstn3,Lgi1,Cacna1b,Adcy1,Abl1,Slc4a8,Tyrobp,Dtnbp1,Crtc1,Iqsec2,Shisa7 |
| GO:0050808 | synapse organization | -14,2369 | -11,213 | 110/506 | Add2,C3,Grin2a,Grm5,Prkca,Gabrb3,Myo5a,Dlg1,Fnta,Cntn2,Gabrb2,Glrb,Pik3r1,Ptprs,Slc1a1,Tnr,Agrn,Utrn,Psen1,Ntrk3,Adam10,C1qb,Gabrg2,L1cam,Chrnb2,Arc,Pclo,Cacnb4,Robo1,Dlg3,Synpo,Tsc1,Lgmn,Arf1,Mfn2,Unc13a,Unc13b,Pcdh8,Snap91,Drp2,Shank1,Grid2,Lrrn3,Actn1,Pafah1b1,Tpbg,Kalrn,Ctnnb1,Mpp2,Picalm,Magi2,Dnm1l,Ntn1,Cntn5,Sorbs2,Nlgn1,Rims2,Nlgn2,Il10ra,Slc8a2,Slc8a3,Adgrl3,Cabp1,Shank2,Opa1,Epha7,Nlgn3,Clstn3,Dnm3,Syngap1,Npas4,Lzts3,Unc13c,Gabra2,Actr2,Ppfia1,Flna,Amigo1,Ntng1,Zmynd8,Setd5,C1qa,Rer1,Flrt2,Abhd17a,Cacna2d2,Trem2,Frmpd4,Tanc2,Tiam1,Sez6l,Pcdh17,Efnb2,Neurl1,Tanc1,Abl1,Plxnd1,Lrfn5,Fzd5,Lhfpl4,Ptprf,Sdk2,Adgre5,Caprin1,C1qc,Ppfia2,Cbln4,Lrrtm4,Dtnbp1,Shisa7,Agt,Ace,Sdc4,Cdh6,Itgb4,Cdh7,Slk,Dlc1,Lin7b,Plec,Dapk3,Cldn5,Myo1c,S100a10,Gjb6,Cxadr,Epb41l3,Ptpn23,Mtdh,Myo9a,Ppm1f,Cldn10,Numbl,Lamtor2,Nphp1,Tns1,Mpp7,Cdh5,Dchs1,Whrn,Peak1,Dusp22,Iqgap1,Specc1l,Macf1,Marveld2,Camk2a,Map2,Strn,Pacsin1,Sema3a,Gsk3a,Crk,Dynlt1,Celsr2,Grip1,Mark1,Mt3,Akap5,Fat3,Ss18l1,Tbc1d24,Hecw1,Tnik,Bcl11a,Camk1d,Mef2a,Btbd3,Prex1,Asap1,Hecw2,RGD1307443,Ankrd27,Rbfox2,Trappc4,Crtc1,Mx1,Stx1b,P2ry1,Syt1,S100b,Cspg5,Mapk9,Kit,Mdk,Fgd4,Plxna2,S100a13,Bcl9l,Rnd2,Cdc42ep4,Dvl3,Map3k13,Tbccd1,Rhobtb2,Plxna3,Plekho1,Zmym3,Coch,Palm2 |
| GO:0050808 | synapse organization | -14,2369 | -11,213 | 110/506 | Add2,C3,Grin2a,Grm5,Prkca,Gabrb3,Myo5a,Dlg1,Fnta,Cntn2,Gabrb2,Glrb,Pik3r1,Ptprs,Slc1a1,Tnr,Agrn,Utrn,Psen1,Ntrk3,Adam10,C1qb,Gabrg2,L1cam,Chrnb2,Arc,Pclo,Cacnb4,Robo1,Dlg3,Synpo,Tsc1,Lgmn,Arf1,Mfn2,Unc13a,Unc13b,Pcdh8,Snap91,Drp2,Shank1,Grid2,Lrrn3,Actn1,Pafah1b1,Tpbg,Kalrn,Ctnnb1,Mpp2,Picalm,Magi2,Dnm1l,Ntn1,Cntn5,Sorbs2,Nlgn1,Rims2,Nlgn2,Il10ra,Slc8a2,Slc8a3,Adgrl3,Cabp1,Shank2,Opa1,Epha7,Nlgn3,Clstn3,Dnm3,Syngap1,Npas4,Lzts3,Unc13c,Gabra2,Actr2,Ppfia1,Flna,Amigo1,Ntng1,Zmynd8,Setd5,C1qa,Rer1,Flrt2,Abhd17a,Cacna2d2,Trem2,Frmpd4,Tanc2,Tiam1,Sez6l,Pcdh17,Efnb2,Neurl1,Tanc1,Abl1,Plxnd1,Lrfn5,Fzd5,Lhfpl4,Ptprf,Sdk2,Adgre5,Caprin1,C1qc,Ppfia2,Cbln4,Lrrtm4,Dtnbp1,Shisa7 |
| GO:0034330 | cell junction organization | -13,1078 | -10,171 | 146/777 | Add2,Agt,C3,Ace,Grin2a,Grm5,Prkca,Sdc4,Gabrb3,Myo5a,Dlg1,Fnta,Cntn2,Cdh6,Gabrb2,Glrb,Pik3r1,Ptprs,Slc1a1,Tnr,Agrn,Utrn,Itgb4,Cdh7,Psen1,Ntrk3,Adam10,C1qb,Gabrg2,L1cam,Chrnb2,Slk,Arc,Pclo,Dlc1,Cacnb4,Robo1,Dlg3,Synpo,Lin7b,Tsc1,Lgmn,Plec,Arf1,Dapk3,Mfn2,Unc13a,Unc13b,Pcdh8,Cldn5,Snap91,Myo1c,Drp2,Shank1,Grid2,Lrrn3,Actn1,S100a10,Pafah1b1,Tpbg,Kalrn,Ctnnb1,Gjb6,Mpp2,Picalm,Cxadr,Magi2,Dnm1l,Ntn1,Cntn5,Sorbs2,Nlgn1,Epb41l3,Rims2,Nlgn2,Il10ra,Ptpn23,Slc8a2,Slc8a3,Adgrl3,Mtdh,Cabp1,Shank2,Opa1,Epha7,Myo9a,Nlgn3,Clstn3,Dnm3,Syngap1,Npas4,Lzts3,Unc13c,Ppm1f,Gabra2,Actr2,Cldn10,Numbl,Ppfia1,Flna,Lamtor2,Amigo1,Ntng1,Nphp1,Zmynd8,Setd5,C1qa,Rer1,Flrt2,Abhd17a,Cacna2d2,Trem2,Tns1,Frmpd4,Tanc2,Tiam1,Sez6l,Pcdh17,Efnb2,Mpp7,Cdh5,Dchs1,Neurl1,Tanc1,Abl1,Plxnd1,Whrn,Lrfn5,Peak1,Fzd5,Lhfpl4,Ptprf,Sdk2,Dusp22,Adgre5,Iqgap1,Specc1l,Caprin1,Macf1,C1qc,Ppfia2,Marveld2,Cbln4,Lrrtm4,Dtnbp1,Shisa7 |
| GO:0016358 | dendrite development | -7,6014 | -5,214 | 65/317 | Camk2a,Ptprs,Map2,Strn,Psen1,Adam10,Pacsin1,Sema3a,Gsk3a,Chrnb2,Crk,Arc,Robo1,Arf1,Mfn2,Shank1,Dynlt1,Celsr2,Pafah1b1,Tpbg,Kalrn,Grip1,Picalm,Dnm1l,Ntn1,Nlgn1,Mark1,Mt3,Nlgn2,Akap5,Shank2,Opa1,Nlgn3,Dnm3,Fat3,Syngap1,Ss18l1,Lzts3,Tbc1d24,Actr2,Hecw1,Numbl,Tnik,Zmynd8,Tanc2,Tiam1,Bcl11a,Camk1d,Neurl1,Mef2a,Btbd3,Prex1,Abl1,Asap1,Hecw2,Ptprf,RGD1307443,Ankrd27,Iqgap1,Caprin1,Ppfia2,Rbfox2,Trappc4,Dtnbp1,Crtc1 |
| GO:0099173 | postsynapse organization | -5,99306 | -3,774 | 46/217 | Grin2a,Dlg1,Fnta,Glrb,Ptprs,Agrn,Ntrk3,Adam10,Arc,Dlg3,Lgmn,Arf1,Mfn2,Shank1,Grid2,Actn1,Pafah1b1,Kalrn,Mpp2,Magi2,Dnm1l,Sorbs2,Nlgn1,Nlgn2,Shank2,Opa1,Epha7,Nlgn3,Dnm3,Syngap1,Lzts3,Actr2,Zmynd8,Rer1,Abhd17a,Frmpd4,Tanc2,Tiam1,Tanc1,Lhfpl4,Ptprf,Caprin1,Ppfia2,Lrrtm4,Dtnbp1,Shisa7 |
| GO:0050803 | regulation of synapse structure or activity | -5,81282 | -3,623 | 54/275 | Mx1,Prkca,Stx1b,Pik3r1,Ptprs,Agrn,Ntrk3,Adam10,Chrnb2,Arc,Mfn2,Pcdh8,Snap91,Grid2,Lrrn3,Pafah1b1,Tpbg,Kalrn,Picalm,Magi2,Dnm1l,Ntn1,Nlgn1,Nlgn2,Il10ra,Adgrl3,Shank2,Opa1,Epha7,Nlgn3,Clstn3,Dnm3,Syngap1,Lzts3,Actr2,Amigo1,Zmynd8,Setd5,Flrt2,Abhd17a,Frmpd4,Tanc2,Tiam1,Neurl1,Tanc1,Abl1,Lrfn5,Lhfpl4,Ptprf,Adgre5,Caprin1,Ppfia2,Lrrtm4,Dtnbp1 |
| GO:0050807 | regulation of synapse organization | -5,36328 | -3,251 | 51/263 | Prkca,Pik3r1,Ptprs,Agrn,Ntrk3,Adam10,Chrnb2,Arc,Mfn2,Pcdh8,Snap91,Grid2,Lrrn3,Pafah1b1,Tpbg,Kalrn,Picalm,Magi2,Dnm1l,Ntn1,Nlgn1,Nlgn2,Il10ra,Adgrl3,Shank2,Opa1,Epha7,Nlgn3,Clstn3,Dnm3,Lzts3,Actr2,Amigo1,Zmynd8,Setd5,Flrt2,Abhd17a,Frmpd4,Tanc2,Tiam1,Neurl1,Tanc1,Abl1,Lrfn5,Lhfpl4,Ptprf,Adgre5,Caprin1,Ppfia2,Lrrtm4,Dtnbp1 |
| GO:0060998 | regulation of dendritic spine development | -5,35677 | -3,248 | 23/82 | Ptprs,Psen1,Arf1,Mfn2,Shank1,Pafah1b1,Kalrn,Dnm1l,Nlgn1,Nlgn2,Shank2,Opa1,Nlgn3,Dnm3,Actr2,Zmynd8,Tanc2,Tiam1,Neurl1,Asap1,Caprin1,Ppfia2,Dtnbp1 |
| GO:0048813 | dendrite morphogenesis | -5,28033 | -3,183 | 40/189 | Camk2a,Map2,Adam10,Sema3a,Chrnb2,Arc,Robo1,Mfn2,Shank1,Celsr2,Pafah1b1,Tpbg,Kalrn,Picalm,Dnm1l,Nlgn1,Mt3,Akap5,Shank2,Opa1,Nlgn3,Dnm3,Ss18l1,Lzts3,Tbc1d24,Actr2,Hecw1,Numbl,Tnik,Tanc2,Tiam1,Mef2a,Btbd3,Hecw2,Ptprf,Ankrd27,Caprin1,Ppfia2,Rbfox2,Dtnbp1 |
| GO:0050773 | regulation of dendrite development | -5,13639 | -3,058 | 32/139 | Pacsin1,Gsk3a,Chrnb2,Crk,Robo1,Mfn2,Pafah1b1,Kalrn,Dnm1l,Mark1,Mt3,Akap5,Opa1,Fat3,Ss18l1,Tbc1d24,Actr2,Hecw1,Numbl,Tnik,Tiam1,Bcl11a,Camk1d,Prex1,Abl1,Hecw2,Ptprf,RGD1307443,Ankrd27,Iqgap1,Caprin1,Crtc1 |
| GO:0061001 | regulation of dendritic spine morphogenesis | -4,72019 | -2,713 | 17/55 | Adam10,Arc,Mfn2,Pafah1b1,Kalrn,Dnm1l,Nlgn1,Opa1,Nlgn3,Dnm3,Lzts3,Actr2,Tanc2,Tiam1,Caprin1,Ppfia2,Dtnbp1 |
| GO:0097061 | dendritic spine organization | -4,24468 | -2,319 | 24/101 | Grin2a,Adam10,Arc,Lgmn,Arf1,Mfn2,Shank1,Pafah1b1,Kalrn,Dnm1l,Nlgn1,Shank2,Opa1,Nlgn3,Dnm3,Lzts3,Actr2,Zmynd8,Tanc2,Tiam1,Tanc1,Caprin1,Ppfia2,Dtnbp1 |
| GO:0022604 | regulation of cell morphogenesis | -4,20442 | -2,292 | 57/334 | Dlg1,P2ry1,Cntn2,Syt1,S100b,Cspg5,Mapk9,Crk,Arc,Dlc1,Kit,Dapk3,Mfn2,Unc13a,Mdk,S100a10,Pafah1b1,Kalrn,Grip1,Picalm,Dnm1l,Epb41l3,Rims2,Akap5,Opa1,Myo9a,Ss18l1,Fgd4,Tbc1d24,Plxna2,Actr2,Numbl,Flna,Tnik,S100a13,Ntng1,Bcl9l,Rnd2,Cdc42ep4,Dvl3,Map3k13,Tbccd1,Tiam1,Bcl11a,Rhobtb2,Plxna3,Plekho1,Prex1,Abl1,Plxnd1,Zmym3,Ptprf,Ankrd27,Caprin1,Macf1,Coch,Palm2 |
| GO:0106027 | neuron projection organization | -4,12455 | -2,229 | 25/109 | Grin2a,Psen1,Adam10,Arc,Lgmn,Arf1,Mfn2,Shank1,Pafah1b1,Kalrn,Dnm1l,Nlgn1,Shank2,Opa1,Nlgn3,Dnm3,Lzts3,Actr2,Zmynd8,Tanc2,Tiam1,Tanc1,Caprin1,Ppfia2,Dtnbp1 |
| GO:0099175 | regulation of postsynapse organization | -3,98849 | -2,117 | 25/111 | Ptprs,Ntrk3,Adam10,Arc,Mfn2,Grid2,Pafah1b1,Kalrn,Dnm1l,Nlgn1,Opa1,Epha7,Nlgn3,Dnm3,Lzts3,Actr2,Zmynd8,Abhd17a,Tanc2,Tiam1,Tanc1,Ptprf,Caprin1,Ppfia2,Dtnbp1 |
| GO:0060997 | dendritic spine morphogenesis | -3,95996 | -2,094 | 19/74 | Adam10,Arc,Mfn2,Shank1,Pafah1b1,Kalrn,Dnm1l,Nlgn1,Shank2,Opa1,Nlgn3,Dnm3,Lzts3,Actr2,Tanc2,Tiam1,Caprin1,Ppfia2,Dtnbp1 |
| GO:0060999 | positive regulation of dendritic spine development | -3,89057 | -2,039 | 17/63 | Psen1,Arf1,Mfn2,Shank1,Pafah1b1,Kalrn,Dnm1l,Nlgn1,Nlgn2,Shank2,Opa1,Nlgn3,Actr2,Zmynd8,Tiam1,Neurl1,Caprin1 |
| GO:0060996 | dendritic spine development | -3,7475 | -1,922 | 27/128 | Camk2a,Ptprs,Psen1,Adam10,Arc,Arf1,Mfn2,Shank1,Pafah1b1,Kalrn,Dnm1l,Nlgn1,Nlgn2,Shank2,Opa1,Nlgn3,Dnm3,Lzts3,Actr2,Zmynd8,Tanc2,Tiam1,Neurl1,Asap1,Caprin1,Ppfia2,Dtnbp1 |
| GO:0010770 | positive regulation of cell morphogenesis involved in differentiation | -3,59469 | -1,810 | 23/104 | Cspg5,Mapk9,Crk,Mfn2,Mdk,S100a10,Pafah1b1,Kalrn,Dnm1l,Akap5,Opa1,Ss18l1,Tbc1d24,Actr2,Numbl,Flna,Tnik,Tiam1,Prex1,Abl1,Ptprf,Ankrd27,Caprin1 |
| GO:0050775 | positive regulation of dendrite morphogenesis | -3,19948 | -1,507 | 15/59 | Mfn2,Pafah1b1,Kalrn,Dnm1l,Akap5,Opa1,Ss18l1,Tbc1d24,Actr2,Numbl,Tnik,Tiam1,Ptprf,Ankrd27,Caprin1 |
| GO:0048814 | regulation of dendrite morphogenesis | -2,91973 | -1,293 | 20/95 | Chrnb2,Robo1,Mfn2,Pafah1b1,Kalrn,Dnm1l,Mt3,Akap5,Opa1,Ss18l1,Tbc1d24,Actr2,Hecw1,Numbl,Tnik,Tiam1,Hecw2,Ptprf,Ankrd27,Caprin1 |
| GO:0010769 | regulation of cell morphogenesis involved in differentiation | -2,84236 | -1,234 | 24/124 | Cntn2,Cspg5,Mapk9,Crk,Mfn2,Mdk,S100a10,Pafah1b1,Kalrn,Dnm1l,Akap5,Opa1,Ss18l1,Tbc1d24,Actr2,Numbl,Flna,Tnik,Tiam1,Prex1,Abl1,Ptprf,Ankrd27,Caprin1 |
| GO:0061003 | positive regulation of dendritic spine morphogenesis | -2,37383 | -0,894 | 8/27 | Mfn2,Pafah1b1,Kalrn,Dnm1l,Opa1,Actr2,Tiam1,Caprin1 |
| GO:0006412 | translation | -13,1975 | -10,197 | 140/732 | Egfr,Grm5,Hbb,Prkca,Akt2,Rps29,Rps5,Rps26,Rpl32,Rpsa,Rpl9,Rps7,Tyms,Rps14,Rps15,Rps17,Rps3a,Fau,Ddx25,Tsc1,Rplp0,Rpl4,Rpl27,Rpl24,Mrpl23,Dapk3,Akap6,Rpl28,Rpl30,Rps12,Atf4,Rpl21,Rpl5,Rpl13,Rpl18,Rpl19,Rpl22,Rpl36al,Rpl37,Rps21,Rps24,Rps2,Ilf3,Abcf1,Polr2g,Rpl6,Rps20,Rps23,Rpl41,Rps3,Rps16,Rplp1,Rps13,Khsrp,Uhmk1,Casc3,Rwdd1,Mrps34,Cnot6,Rpl26,Mrpl27,Krt17,Nanos3,Mrps14,Mrps18c,Mrpl34,Rpl18a,Fto,Rmnd1,Mrps12,Eif3k,Rpl36a,Eif3f,mrpl11,Eef1g,Mrpl16,Mrps18b,Rps18,Foxo3,Rbm8a,Eif3m,Secisbp2l,Eif2s2,Mrpl41,Pus7,Mrpl19,Mrps35,Otud6b,Mrpl13,Eef1d,Gtpbp1,Rpl3,Xrn1,Mrps18a,Mrpl14,Aars2,Mcts1,Cpeb4,Ybx2,Mrpl12,Eif4enif1,Eif6,Eif5b,Neurl1,Nelfb,Tia1,Upf3b,Rpl13a,Rpl23a,Mrps23,Rcc1l,Chchd1,Mrpl52,Upf3a,Uqcc2,Eif4ebp2,Rpl22l1,Rpl34,Zc3h15,Celf1,Caprin1,Lars2,Eef1b2,Rpl37a-ps1,Impact,Coa3,Purb,Qk,Rpl12,Mknk1,Dph3,Rps27l,Denr,Rpl38,Mrps21,Srp9,Rps28,Eif3j,Rps4x,Rps27a,Dmd,Slc1a1,Clu,Hsd11b1,Gsk3a,Csnk1e,Picalm,Bin1,Prkcd,Rtn2 |
| GO:0006412 | translation | -13,1975 | -10,197 | 140/732 | Egfr,Grm5,Hbb,Prkca,Akt2,Rps29,Rps5,Rps26,Rpl32,Rpsa,Rpl9,Rps7,Tyms,Rps14,Rps15,Rps17,Rps3a,Fau,Ddx25,Tsc1,Rplp0,Rpl4,Rpl27,Rpl24,Mrpl23,Dapk3,Akap6,Rpl28,Rpl30,Rps12,Atf4,Rpl21,Rpl5,Rpl13,Rpl18,Rpl19,Rpl22,Rpl36al,Rpl37,Rps21,Rps24,Rps2,Ilf3,Abcf1,Polr2g,Rpl6,Rps20,Rps23,Rpl41,Rps3,Rps16,Rplp1,Rps13,Khsrp,Uhmk1,Casc3,Rwdd1,Mrps34,Cnot6,Rpl26,Mrpl27,Krt17,Nanos3,Mrps14,Mrps18c,Mrpl34,Rpl18a,Fto,Rmnd1,Mrps12,Eif3k,Rpl36a,Eif3f,mrpl11,Eef1g,Mrpl16,Mrps18b,Rps18,Foxo3,Rbm8a,Eif3m,Secisbp2l,Eif2s2,Mrpl41,Pus7,Mrpl19,Mrps35,Otud6b,Mrpl13,Eef1d,Gtpbp1,Rpl3,Xrn1,Mrps18a,Mrpl14,Aars2,Mcts1,Cpeb4,Ybx2,Mrpl12,Eif4enif1,Eif6,Eif5b,Neurl1,Nelfb,Tia1,Upf3b,Rpl13a,Rpl23a,Mrps23,Rcc1l,Chchd1,Mrpl52,Upf3a,Uqcc2,Eif4ebp2,Rpl22l1,Rpl34,Zc3h15,Celf1,Caprin1,Lars2,Eef1b2,Rpl37a-ps1,Impact,Coa3,Purb,Qk,Rpl12,Mknk1,Dph3,Rps27l,Denr,Rpl38,Mrps21,Srp9,Rps28,Eif3j,Rps4x,Rps27a |
| GO:0043043 | peptide biosynthetic process | -12,9292 | -10,012 | 142/752 | Egfr,Grm5,Hbb,Prkca,Dmd,Akt2,Rps29,Rps5,Slc1a1,Rps26,Rpl32,Rpsa,Rpl9,Rps7,Tyms,Rps14,Rps15,Rps17,Rps3a,Fau,Ddx25,Tsc1,Rplp0,Rpl4,Rpl27,Rpl24,Mrpl23,Dapk3,Akap6,Rpl28,Rpl30,Rps12,Atf4,Rpl21,Rpl5,Rpl13,Rpl18,Rpl19,Rpl22,Rpl36al,Rpl37,Rps21,Rps24,Rps2,Ilf3,Abcf1,Polr2g,Rpl6,Rps20,Rps23,Rpl41,Rps3,Rps16,Rplp1,Rps13,Khsrp,Uhmk1,Casc3,Rwdd1,Mrps34,Cnot6,Rpl26,Mrpl27,Krt17,Nanos3,Mrps14,Mrps18c,Mrpl34,Rpl18a,Fto,Rmnd1,Mrps12,Eif3k,Rpl36a,Eif3f,mrpl11,Eef1g,Mrpl16,Mrps18b,Rps18,Foxo3,Rbm8a,Eif3m,Secisbp2l,Eif2s2,Mrpl41,Pus7,Mrpl19,Mrps35,Otud6b,Mrpl13,Eef1d,Gtpbp1,Rpl3,Xrn1,Mrps18a,Mrpl14,Aars2,Mcts1,Cpeb4,Ybx2,Mrpl12,Eif4enif1,Eif6,Eif5b,Neurl1,Nelfb,Tia1,Upf3b,Rpl13a,Rpl23a,Mrps23,Rcc1l,Chchd1,Mrpl52,Upf3a,Uqcc2,Eif4ebp2,Rpl22l1,Rpl34,Zc3h15,Celf1,Caprin1,Lars2,Eef1b2,Rpl37a-ps1,Impact,Coa3,Purb,Qk,Rpl12,Mknk1,Dph3,Rps27l,Denr,Rpl38,Mrps21,Srp9,Rps28,Eif3j,Rps4x,Rps27a |
| GO:0034248 | regulation of cellular amide metabolic process | -5,59595 | -3,437 | 72/410 | Grm5,Hbb,Prkca,Clu,Hsd11b1,Akt2,Tyms,Rps14,Rps3a,Gsk3a,Csnk1e,Ddx25,Tsc1,Dapk3,Akap6,Rpl30,Atf4,Rpl5,Rpl22,Ilf3,Abcf1,Picalm,Polr2g,Bin1,Rps23,Rps3,Rplp1,Prkcd,Khsrp,Uhmk1,Casc3,Cnot6,Rpl26,Krt17,Nanos3,Fto,Rmnd1,Eif3k,Foxo3,Rbm8a,Secisbp2l,Pus7,Otud6b,Mrpl13,Xrn1,Cpeb4,Ybx2,Eif4enif1,Eif6,Eif5b,Rtn2,Neurl1,Tia1,Upf3b,Rpl13a,Rcc1l,Upf3a,Uqcc2,Eif4ebp2,Celf1,Caprin1,Impact,Coa3,Purb,Qk,Rpl12,Mknk1,Dph3,Rps27l,Rpl38,Srp9,Rps4x |
| GO:0006417 | regulation of translation | -5,55613 | -3,404 | 64/352 | Grm5,Hbb,Prkca,Akt2,Tyms,Rps14,Rps3a,Ddx25,Tsc1,Dapk3,Akap6,Rpl30,Atf4,Rpl5,Rpl22,Ilf3,Abcf1,Polr2g,Rps23,Rps3,Rplp1,Khsrp,Uhmk1,Casc3,Cnot6,Rpl26,Krt17,Nanos3,Fto,Rmnd1,Eif3k,Foxo3,Rbm8a,Secisbp2l,Pus7,Otud6b,Mrpl13,Xrn1,Cpeb4,Ybx2,Eif4enif1,Eif6,Eif5b,Neurl1,Tia1,Upf3b,Rpl13a,Rcc1l,Upf3a,Uqcc2,Eif4ebp2,Celf1,Caprin1,Impact,Coa3,Purb,Qk,Rpl12,Mknk1,Dph3,Rps27l,Rpl38,Srp9,Rps4x |
| GO:0034249 | negative regulation of cellular amide metabolic process | -2,58962 | -1,055 | 29/166 | Prkca,Clu,Tyms,Tsc1,Dapk3,Atf4,Ilf3,Bin1,Rps3,Khsrp,Nanos3,Fto,Pus7,Otud6b,Mrpl13,Xrn1,Cpeb4,Ybx2,Eif4enif1,Eif6,Rtn2,Tia1,Rpl13a,Eif4ebp2,Celf1,Caprin1,Purb,Qk,Srp9 |
| GO:0017148 | negative regulation of translation | -2,58177 | -1,048 | 26/144 | Prkca,Tyms,Tsc1,Dapk3,Atf4,Ilf3,Rps3,Khsrp,Nanos3,Fto,Pus7,Otud6b,Mrpl13,Xrn1,Cpeb4,Ybx2,Eif4enif1,Eif6,Tia1,Rpl13a,Eif4ebp2,Celf1,Caprin1,Purb,Qk,Srp9 |
| R-RNO-112316 | Neuronal System | -11,7859 | -8,969 | 72/298 | Camk2d,Gad2,Grin2a,Grm5,Prkcg,Gabrb3,Dlg1,Camk2a,Gabrb2,Glrb,Kcna5,Ptprs,Slc1a1,Abcc9,Gnai1,Syt1,Ntrk3,Prkar2a,Gabrg2,Kcnj10,Kcnab2,Gria1,Chrnb2,Grik2,Cacnb4,Dlg3,Nsf,Lin7b,Lin7c,Unc13b,Ap2s1,Kcnmb4,Kcns2,Shank1,Dnajc5,Gabrg3,Slc6a11,Abat,Apba1,Gabbr2,Kcnma1,Grip1,Kcnh4,Gng10,Hcn3,Nlgn1,Dlgap2,Epb41l3,Nlgn2,Lrrc7,Ap2b1,Gabra4,Cacng3,Cacng4,Slc38a1,Kcnq2,Shank2,Camk2g,Nlgn3,Gng8,Prkacb,Ppfia1,Grik3,Kcnf1,Cacna2d2,Adcy1,Ap2a1,Panx1,Rps6ka6,Ptprf,Lrrtm4,Gng13,Pde4b,Prkca,Arrb1,Pde4a,Adora1,Pde1b,Gnao1,Pde10a,Pde7b,Cacna1b,Gabra2,Pde8b,Cacna1c,Cacna1d,Npas4,Lhfpl4,Cbln4,Dbi,Htr1b,Prkce,Tpbg,Clcn3,Clstn3,Ptgs2,Mapk9,Gnaq,Atf4,Shisa7 |
| R-RNO-112316 | Neuronal System | -11,7859 | -8,969 | 72/298 | Camk2d,Gad2,Grin2a,Grm5,Prkcg,Gabrb3,Dlg1,Camk2a,Gabrb2,Glrb,Kcna5,Ptprs,Slc1a1,Abcc9,Gnai1,Syt1,Ntrk3,Prkar2a,Gabrg2,Kcnj10,Kcnab2,Gria1,Chrnb2,Grik2,Cacnb4,Dlg3,Nsf,Lin7b,Lin7c,Unc13b,Ap2s1,Kcnmb4,Kcns2,Shank1,Dnajc5,Gabrg3,Slc6a11,Abat,Apba1,Gabbr2,Kcnma1,Grip1,Kcnh4,Gng10,Hcn3,Nlgn1,Dlgap2,Epb41l3,Nlgn2,Lrrc7,Ap2b1,Gabra4,Cacng3,Cacng4,Slc38a1,Kcnq2,Shank2,Camk2g,Nlgn3,Gng8,Prkacb,Ppfia1,Grik3,Kcnf1,Cacna2d2,Adcy1,Ap2a1,Panx1,Rps6ka6,Ptprf,Lrrtm4,Gng13 |
| R-RNO-112315 | Transmission across Chemical Synapses | -9,69245 | -7,076 | 49/185 | Camk2d,Gad2,Grin2a,Prkcg,Gabrb3,Dlg1,Camk2a,Gabrb2,Glrb,Slc1a1,Gnai1,Syt1,Prkar2a,Gabrg2,Kcnj10,Gria1,Chrnb2,Grik2,Cacnb4,Dlg3,Nsf,Lin7b,Lin7c,Unc13b,Ap2s1,Dnajc5,Gabrg3,Slc6a11,Abat,Apba1,Gabbr2,Grip1,Gng10,Lrrc7,Ap2b1,Gabra4,Cacng3,Cacng4,Slc38a1,Camk2g,Gng8,Prkacb,Ppfia1,Grik3,Cacna2d2,Adcy1,Ap2a1,Rps6ka6,Gng13 |
| R-RNO-112314 | Neurotransmitter receptors and postsynaptic signal transmission | -7,59246 | -5,211 | 35/127 | Camk2d,Grin2a,Prkcg,Gabrb3,Dlg1,Camk2a,Gabrb2,Glrb,Gnai1,Prkar2a,Gabrg2,Kcnj10,Gria1,Chrnb2,Grik2,Dlg3,Nsf,Ap2s1,Gabrg3,Gabbr2,Grip1,Gng10,Lrrc7,Ap2b1,Gabra4,Cacng3,Cacng4,Camk2g,Gng8,Prkacb,Grik3,Adcy1,Ap2a1,Rps6ka6,Gng13 |
| ko05032 | Morphine addiction | -5,49176 | -3,349 | 25/92 | Pde4b,Prkca,Prkcg,Gabrb3,Arrb1,Gabrb2,Pde4a,Gnai1,Adora1,Pde1b,Gabrg2,Gnao1,Pde10a,Gabrg3,Gabbr2,Gng10,Gabra4,Pde7b,Gng8,Cacna1b,Gabra2,Prkacb,Adcy1,Pde8b,Gng13 |
| rno05032 | Morphine addiction | -5,49176 | -3,349 | 25/92 | Pde4b,Prkca,Prkcg,Gabrb3,Arrb1,Gabrb2,Pde4a,Gnai1,Adora1,Pde1b,Gabrg2,Gnao1,Pde10a,Gabrg3,Gabbr2,Gng10,Gabra4,Pde7b,Gng8,Cacna1b,Gabra2,Prkacb,Adcy1,Pde8b,Gng13 |
| ko04727 | GABAergic synapse | -5,14354 | -3,063 | 24/90 | Cacna1c,Gad2,Prkca,Prkcg,Gabrb3,Gabrb2,Gnai1,Gabrg2,Cacna1d,Gnao1,Nsf,Gabrg3,Slc6a11,Abat,Gabbr2,Gng10,Gabra4,Slc38a1,Gng8,Cacna1b,Gabra2,Prkacb,Adcy1,Gng13 |
| rno04727 | GABAergic synapse | -5,14354 | -3,063 | 24/90 | Cacna1c,Gad2,Prkca,Prkcg,Gabrb3,Gabrb2,Gnai1,Gabrg2,Cacna1d,Gnao1,Nsf,Gabrg3,Slc6a11,Abat,Gabbr2,Gng10,Gabra4,Slc38a1,Gng8,Cacna1b,Gabra2,Prkacb,Adcy1,Gng13 |
| GO:1904862 | inhibitory synapse assembly | -4,42907 | -2,488 | 8/15 | Gabrb3,Gabrb2,Gabrg2,Nlgn2,Npas4,Gabra2,Lhfpl4,Cbln4 |
| GO:0051932 | synaptic transmission, GABAergic | -3,89057 | -2,039 | 17/63 | Gabrb3,Dbi,Htr1b,Gabrb2,Adora1,Prkce,Gabrg2,Cacnb4,Gabrg3,Tpbg,Clcn3,Nlgn1,Nlgn2,Gabra4,Clstn3,Npas4,Gabra2 |
| ko04723 | Retrograde endocannabinoid signaling | -3,66009 | -1,862 | 23/103 | Cacna1c,Grm5,Prkca,Prkcg,Gabrb3,Gabrb2,Gnai1,Ptgs2,Gabrg2,Cacna1d,Gria1,Mapk9,Gnao1,Gabrg3,Gnaq,Gng10,Gabra4,Gng8,Cacna1b,Gabra2,Prkacb,Adcy1,Gng13 |
| rno04723 | Retrograde endocannabinoid signaling | -3,66009 | -1,862 | 23/103 | Cacna1c,Grm5,Prkca,Prkcg,Gabrb3,Gabrb2,Gnai1,Ptgs2,Gabrg2,Cacna1d,Gria1,Mapk9,Gnao1,Gabrg3,Gnaq,Gng10,Gabra4,Gng8,Cacna1b,Gabra2,Prkacb,Adcy1,Gng13 |
| GO:0007214 | gamma-aminobutyric acid signaling pathway | -3,42952 | -1,682 | 10/29 | Gabrb3,Gabrb2,Gabrg2,Cacnb4,Gabrg3,Atf4,Gabbr2,Gabra4,Gabra2,Shisa7 |
| ko05033 | Nicotine addiction | -2,25233 | -0,819 | 10/40 | Grin2a,Gabrb3,Gabrb2,Gabrg2,Gria1,Chrnb2,Gabrg3,Gabra4,Cacna1b,Gabra2 |
| rno05033 | Nicotine addiction | -2,25233 | -0,819 | 10/40 | Grin2a,Gabrb3,Gabrb2,Gabrg2,Gria1,Chrnb2,Gabrg3,Gabra4,Cacna1b,Gabra2 |
| R-RNO-977443 | GABA receptor activation | -2,22447 | -0,798 | 12/53 | Gabrb3,Gabrb2,Gnai1,Gabrg2,Kcnj10,Gabrg3,Gabbr2,Gng10,Gabra4,Gng8,Adcy1,Gng13 |
| GO:0010975 | regulation of neuron projection development | -11,053 | -8,292 | 110/563 | Agt,B2m,Gfap,Dmd,Stx1b,RT1-A2,Cntn2,Ptprs,Tnr,Map2,Psen1,Klk6,Ntrk3,Adam10,Pacsin1,Sema3a,Atp5mc1,Gsk3a,L1cam,Ntm,Chrnb2,Crk,Arc,Robo1,Map2k2,Tsc1,Rpl4,Mfn2,Snap91,Grid2,Mdk,Nr2f1,Dynlt1,Pafah1b1,Nme2,Kalrn,Chn1,Kif3c,Picalm,Magi2,Kremen1,Dnm1l,Ntn1,Nlgn1,Mark1,Mt3,Lrrc7,Bmpr2,Ndel1,Akap5,Opa1,Camk2g,Epha7,Nlgn3,Dnm3,Fat3,Nme1,Syngap1,Ss18l1,Lzts3,Tbc1d24,Metrn,Plxna2,Actr2,Sema3g,Hecw1,Slc39a12,Numbl,Flna,Tnik,Amigo1,Ntng1,Cbfa2t2,Sf3a2,Rnd2,Tanc2,Map3k13,Tiam1,Rrn3,Bcl11a,Kif13b,Efnb2,Camk1d,Ap2a1,Inpp5f,Plxna3,Lrig2,Prex1,Abl1,Plxnd1,Lrp6,Plk5,Hecw2,Neu4,Ptprf,Stk24,RGD1307443,Ust,Ankrd27,Iqgap1,Caprin1,Macf1,Ppfia2,Fbxo7,Qk,Fxn,Dtnbp1,Rbpj,Twf2,Crtc1,Apc,Egfr,Ncam1,Prkca,Clu,Cck,Camk2a,Nog,Agrn,Ddr1,Syt1,Stmn1,Cspg5,Notch3,Unc5a,Unc5b,Rpl24,Isl1,Unc13a,Dpysl5,Shank1,Alcam,Celsr2,Celsr3,Tpbg,Grip1,Ctnnb1,Ptch1,Cntn4,Epb41l3,Sh3gl2,Rims2,Tenm2,Gli3,Shank2,Myo9a,Nexn,Lgi1,Taok1,Lgr4,Tubb2b,Pak6,Lhx2,Flrt2,Mapk8ip3,Jade2,Dvl3,Taok3,Adcy1,Mef2a,Plekho1,Btbd3,Zswim5,Pip5k1c,Etv4,Rbfox2,Raph1,Zdhhc17,Impact,Nrn1l,Fis1,Ace,Grm5,Atxn1,Pik3r1,Kit,Lrrn3,Synj1,Bin1,Nlgn2,Adgrl3,Clstn3,Lig4,Smarcd3,Plag1,Tenm4,Nkx6-2,Neurl1,Ufl1,Dmrta2,Adgre5,Lrrtm4,Bex1,Rab5a,Prkcd,Ift20,Tbc1d17,Zmynd8,Trem2,Cdc42ep4,Ripor2,Fnbp1l,Tmem67,Mien1,Tbc1d20,Ccl19,Dynlt2b,Dlg1,S100b,Bhlhe40,Atf5,Tmem98,Kdm4a,Lpin1,Lingo1,Tppp,Tcf7l2,Sod1,Cdhr1,Hexa,Whrn,S1pr2,Mapk9,Cldn5,S100a10,Cdh5,Tyrobp,Jak2,Mtr,Nrep,Svbp,Camk2d,Akap6,Fdps,Sorbs2,Sox9,Map2k4,Med12,Extl3,Cyba,Mtpn,Csnk2a1,Dnph1,Krt17,Hyal1,Slc25a33 |
| GO:0010975 | regulation of neuron projection development | -11,053 | -8,292 | 110/563 | Agt,B2m,Gfap,Dmd,Stx1b,RT1-A2,Cntn2,Ptprs,Tnr,Map2,Psen1,Klk6,Ntrk3,Adam10,Pacsin1,Sema3a,Atp5mc1,Gsk3a,L1cam,Ntm,Chrnb2,Crk,Arc,Robo1,Map2k2,Tsc1,Rpl4,Mfn2,Snap91,Grid2,Mdk,Nr2f1,Dynlt1,Pafah1b1,Nme2,Kalrn,Chn1,Kif3c,Picalm,Magi2,Kremen1,Dnm1l,Ntn1,Nlgn1,Mark1,Mt3,Lrrc7,Bmpr2,Ndel1,Akap5,Opa1,Camk2g,Epha7,Nlgn3,Dnm3,Fat3,Nme1,Syngap1,Ss18l1,Lzts3,Tbc1d24,Metrn,Plxna2,Actr2,Sema3g,Hecw1,Slc39a12,Numbl,Flna,Tnik,Amigo1,Ntng1,Cbfa2t2,Sf3a2,Rnd2,Tanc2,Map3k13,Tiam1,Rrn3,Bcl11a,Kif13b,Efnb2,Camk1d,Ap2a1,Inpp5f,Plxna3,Lrig2,Prex1,Abl1,Plxnd1,Lrp6,Plk5,Hecw2,Neu4,Ptprf,Stk24,RGD1307443,Ust,Ankrd27,Iqgap1,Caprin1,Macf1,Ppfia2,Fbxo7,Qk,Fxn,Dtnbp1,Rbpj,Twf2,Crtc1 |
| GO:0048858 | cell projection morphogenesis | -10,4806 | -7,758 | 131/727 | Apc,Egfr,Ncam1,Prkca,Clu,Dmd,Cck,Cntn2,Camk2a,Nog,Ptprs,Tnr,Agrn,Map2,Ddr1,Syt1,Psen1,Stmn1,Ntrk3,Adam10,Pacsin1,Sema3a,Atp5mc1,Cspg5,L1cam,Chrnb2,Arc,Notch3,Robo1,Map2k2,Unc5a,Unc5b,Rpl4,Rpl24,Isl1,Mfn2,Unc13a,Snap91,Dpysl5,Shank1,Alcam,Mdk,Dynlt1,Celsr2,Celsr3,Pafah1b1,Tpbg,Kalrn,Grip1,Chn1,Ctnnb1,Picalm,Ptch1,Dnm1l,Ntn1,Nlgn1,Cntn4,Epb41l3,Sh3gl2,Rims2,Mt3,Tenm2,Gli3,Bmpr2,Ndel1,Akap5,Shank2,Opa1,Epha7,Myo9a,Nlgn3,Dnm3,Syngap1,Ss18l1,Nexn,Lgi1,Lzts3,Taok1,Lgr4,Tbc1d24,Metrn,Plxna2,Actr2,Sema3g,Tubb2b,Hecw1,Numbl,Tnik,Amigo1,Ntng1,Pak6,Lhx2,Flrt2,Mapk8ip3,Jade2,Rnd2,Tanc2,Dvl3,Map3k13,Tiam1,Taok3,Adcy1,Bcl11a,Kif13b,Efnb2,Plxna3,Mef2a,Plekho1,Btbd3,Abl1,Plxnd1,Zswim5,Pip5k1c,Hecw2,Ptprf,Etv4,RGD1307443,Ust,Ankrd27,Iqgap1,Caprin1,Macf1,Ppfia2,Rbfox2,Raph1,Zdhhc17,Impact,Fxn,Dtnbp1,Twf2,Nrn1l |
| GO:0048812 | neuron projection morphogenesis | -10,4344 | -7,723 | 128/706 | Apc,Egfr,Ncam1,Prkca,Clu,Dmd,Cck,Cntn2,Camk2a,Nog,Ptprs,Tnr,Agrn,Map2,Ddr1,Syt1,Psen1,Stmn1,Ntrk3,Adam10,Pacsin1,Sema3a,Atp5mc1,L1cam,Chrnb2,Arc,Notch3,Robo1,Map2k2,Unc5a,Unc5b,Rpl4,Rpl24,Isl1,Mfn2,Unc13a,Snap91,Dpysl5,Shank1,Alcam,Dynlt1,Celsr2,Celsr3,Pafah1b1,Tpbg,Kalrn,Grip1,Chn1,Ctnnb1,Picalm,Ptch1,Dnm1l,Ntn1,Nlgn1,Cntn4,Epb41l3,Sh3gl2,Rims2,Mt3,Tenm2,Gli3,Bmpr2,Ndel1,Akap5,Shank2,Opa1,Epha7,Myo9a,Nlgn3,Dnm3,Syngap1,Ss18l1,Nexn,Lgi1,Lzts3,Taok1,Lgr4,Tbc1d24,Metrn,Plxna2,Actr2,Sema3g,Tubb2b,Hecw1,Numbl,Tnik,Amigo1,Ntng1,Pak6,Lhx2,Flrt2,Mapk8ip3,Jade2,Rnd2,Tanc2,Dvl3,Map3k13,Tiam1,Taok3,Adcy1,Bcl11a,Kif13b,Efnb2,Plxna3,Mef2a,Btbd3,Abl1,Plxnd1,Zswim5,Pip5k1c,Hecw2,Ptprf,Etv4,RGD1307443,Ust,Ankrd27,Iqgap1,Caprin1,Macf1,Ppfia2,Rbfox2,Raph1,Zdhhc17,Impact,Fxn,Dtnbp1,Twf2,Nrn1l |
| GO:0120039 | plasma membrane bounded cell projection morphogenesis | -10,0322 | -7,376 | 129/723 | Apc,Egfr,Ncam1,Prkca,Clu,Dmd,Cck,Cntn2,Camk2a,Nog,Ptprs,Tnr,Agrn,Map2,Ddr1,Syt1,Psen1,Stmn1,Ntrk3,Adam10,Pacsin1,Sema3a,Atp5mc1,L1cam,Chrnb2,Arc,Notch3,Robo1,Map2k2,Unc5a,Unc5b,Rpl4,Rpl24,Isl1,Mfn2,Unc13a,Snap91,Dpysl5,Shank1,Alcam,Dynlt1,Celsr2,Celsr3,Pafah1b1,Tpbg,Kalrn,Grip1,Chn1,Ctnnb1,Picalm,Ptch1,Dnm1l,Ntn1,Nlgn1,Cntn4,Epb41l3,Sh3gl2,Rims2,Mt3,Tenm2,Gli3,Bmpr2,Ndel1,Akap5,Shank2,Opa1,Epha7,Myo9a,Nlgn3,Dnm3,Syngap1,Ss18l1,Nexn,Lgi1,Lzts3,Taok1,Lgr4,Tbc1d24,Metrn,Plxna2,Actr2,Sema3g,Tubb2b,Hecw1,Numbl,Tnik,Amigo1,Ntng1,Pak6,Lhx2,Flrt2,Mapk8ip3,Jade2,Rnd2,Tanc2,Dvl3,Map3k13,Tiam1,Taok3,Adcy1,Bcl11a,Kif13b,Efnb2,Plxna3,Mef2a,Plekho1,Btbd3,Abl1,Plxnd1,Zswim5,Pip5k1c,Hecw2,Ptprf,Etv4,RGD1307443,Ust,Ankrd27,Iqgap1,Caprin1,Macf1,Ppfia2,Rbfox2,Raph1,Zdhhc17,Impact,Fxn,Dtnbp1,Twf2,Nrn1l |
| GO:0032990 | cell part morphogenesis | -9,88347 | -7,248 | 132/749 | Apc,Egfr,Ncam1,Prkca,Clu,Dmd,Cck,Cntn2,Camk2a,Nog,Ptprs,Tnr,Agrn,Map2,Ddr1,Syt1,Psen1,Stmn1,Ntrk3,Adam10,Pacsin1,Sema3a,Atp5mc1,Cspg5,L1cam,Chrnb2,Arc,Notch3,Robo1,Map2k2,Unc5a,Unc5b,Rpl4,Rpl24,Isl1,Mfn2,Unc13a,Snap91,Dpysl5,Shank1,Alcam,Mdk,Dynlt1,Celsr2,Celsr3,Pafah1b1,Tpbg,Kalrn,Grip1,Chn1,Ctnnb1,Picalm,Ptch1,Dnm1l,Ntn1,Nlgn1,Cntn4,Epb41l3,Sh3gl2,Rims2,Mt3,Tenm2,Gli3,Bmpr2,Ndel1,Akap5,Shank2,Opa1,Epha7,Myo9a,Nlgn3,Dnm3,Syngap1,Ss18l1,Nexn,Lgi1,Lzts3,Taok1,Lgr4,Tbc1d24,Metrn,Fis1,Plxna2,Actr2,Sema3g,Tubb2b,Hecw1,Numbl,Tnik,Amigo1,Ntng1,Pak6,Lhx2,Flrt2,Mapk8ip3,Jade2,Rnd2,Tanc2,Dvl3,Map3k13,Tiam1,Taok3,Adcy1,Bcl11a,Kif13b,Efnb2,Plxna3,Mef2a,Plekho1,Btbd3,Abl1,Plxnd1,Zswim5,Pip5k1c,Hecw2,Ptprf,Etv4,RGD1307443,Ust,Ankrd27,Iqgap1,Caprin1,Macf1,Ppfia2,Rbfox2,Raph1,Zdhhc17,Impact,Fxn,Dtnbp1,Twf2,Nrn1l |
| GO:0051962 | positive regulation of nervous system development | -8,83741 | -6,336 | 74/356 | Ace,Egfr,Gfap,Grm5,Prkca,Atxn1,Nog,Pik3r1,Agrn,Ntrk3,Atp5mc1,L1cam,Robo1,Map2k2,Kit,Rpl4,Mfn2,Snap91,Grid2,Lrrn3,Mdk,Pafah1b1,Tpbg,Kalrn,Ctnnb1,Synj1,Picalm,Dnm1l,Ntn1,Nlgn1,Bin1,Nlgn2,Gli3,Bmpr2,Adgrl3,Ndel1,Akap5,Opa1,Nlgn3,Clstn3,Ss18l1,Tbc1d24,Metrn,Plxna2,Actr2,Lig4,Numbl,Tnik,Amigo1,Smarcd3,Plag1,Flrt2,Jade2,Rnd2,Map3k13,Tiam1,Bcl11a,Tenm4,Nkx6-2,Plxna3,Neurl1,Plxnd1,Ufl1,Dmrta2,Ptprf,Adgre5,Ankrd27,Caprin1,Macf1,Qk,Fxn,Lrrtm4,Bex1,Twf2 |
| GO:0031344 | regulation of cell projection organization | -8,44236 | -5,969 | 133/795 | Agt,Apc,B2m,Gfap,Dmd,Stx1b,RT1-A2,Cntn2,Pik3r1,Ptprs,Tnr,Agrn,Map2,Psen1,Klk6,Ntrk3,Adam10,Pacsin1,Sema3a,Atp5mc1,Gsk3a,L1cam,Ntm,Chrnb2,Crk,Arc,Robo1,Map2k2,Tsc1,Kit,Rpl4,Mfn2,Rab5a,Snap91,Grid2,Mdk,Nr2f1,Dynlt1,Pafah1b1,Nme2,Kalrn,Grip1,Chn1,Kif3c,Picalm,Magi2,Kremen1,Dnm1l,Ntn1,Nlgn1,Mark1,Mt3,Tenm2,Lrrc7,Bmpr2,Prkcd,Ndel1,Akap5,Opa1,Camk2g,Epha7,Myo9a,Nlgn3,Dnm3,Fat3,Nme1,Syngap1,Ss18l1,Lzts3,Tbc1d24,Metrn,Ift20,Plxna2,Actr2,Sema3g,Hecw1,Slc39a12,Numbl,Tbc1d17,Flna,Tnik,Amigo1,Ntng1,Cbfa2t2,Zmynd8,Sf3a2,Trem2,Rnd2,Tanc2,Cdc42ep4,Dvl3,Map3k13,Tiam1,Rrn3,Bcl11a,Kif13b,Efnb2,Ripor2,Camk1d,Ap2a1,Inpp5f,Plxna3,Neurl1,Lrig2,Fnbp1l,Prex1,Abl1,Plxnd1,Lrp6,Tmem67,Plk5,Hecw2,Neu4,Ptprf,Mien1,Stk24,RGD1307443,Ust,Ankrd27,Iqgap1,Caprin1,Tbc1d20,Ccl19,Macf1,Ppfia2,Fbxo7,Dynlt2b,Qk,Fxn,Dtnbp1,Rbpj,Twf2,Crtc1 |
| GO:0120035 | regulation of plasma membrane bounded cell projection organization | -8,19471 | -5,741 | 130/779 | Agt,Apc,B2m,Gfap,Dmd,Stx1b,RT1-A2,Cntn2,Pik3r1,Ptprs,Tnr,Agrn,Map2,Psen1,Klk6,Ntrk3,Adam10,Pacsin1,Sema3a,Atp5mc1,Gsk3a,L1cam,Ntm,Chrnb2,Crk,Arc,Robo1,Map2k2,Tsc1,Kit,Rpl4,Mfn2,Rab5a,Snap91,Grid2,Mdk,Nr2f1,Dynlt1,Pafah1b1,Nme2,Kalrn,Chn1,Kif3c,Picalm,Magi2,Kremen1,Dnm1l,Ntn1,Nlgn1,Mark1,Mt3,Tenm2,Lrrc7,Bmpr2,Prkcd,Ndel1,Akap5,Opa1,Camk2g,Epha7,Nlgn3,Dnm3,Fat3,Nme1,Syngap1,Ss18l1,Lzts3,Tbc1d24,Metrn,Ift20,Plxna2,Actr2,Sema3g,Hecw1,Slc39a12,Numbl,Tbc1d17,Flna,Tnik,Amigo1,Ntng1,Cbfa2t2,Zmynd8,Sf3a2,Trem2,Rnd2,Tanc2,Cdc42ep4,Map3k13,Tiam1,Rrn3,Bcl11a,Kif13b,Efnb2,Ripor2,Camk1d,Ap2a1,Inpp5f,Plxna3,Neurl1,Lrig2,Fnbp1l,Prex1,Abl1,Plxnd1,Lrp6,Tmem67,Plk5,Hecw2,Neu4,Ptprf,Mien1,Stk24,RGD1307443,Ust,Ankrd27,Iqgap1,Caprin1,Tbc1d20,Ccl19,Macf1,Ppfia2,Fbxo7,Dynlt2b,Qk,Fxn,Dtnbp1,Rbpj,Twf2,Crtc1 |
| GO:0051960 | regulation of nervous system development | -7,75706 | -5,347 | 98/549 | B2m,Ace,Egfr,Gfap,Grm5,Prkca,Atxn1,Dlg1,Nog,Pik3r1,Ptprs,Tnr,Agrn,Map2,S100b,Psen1,Ntrk3,Sema3a,Atp5mc1,L1cam,Robo1,Map2k2,Kit,Rpl4,Mfn2,Snap91,Grid2,Bhlhe40,Lrrn3,Mdk,Dynlt1,Pafah1b1,Tpbg,Kalrn,Ctnnb1,Synj1,Picalm,Dnm1l,Ntn1,Nlgn1,Bin1,Mt3,Nlgn2,Gli3,Bmpr2,Adgrl3,Ndel1,Akap5,Opa1,Epha7,Nlgn3,Clstn3,Syngap1,Ss18l1,Atf5,Tbc1d24,Metrn,Plxna2,Actr2,Sema3g,Lig4,Numbl,Tnik,Amigo1,Lhx2,Smarcd3,Plag1,Flrt2,Jade2,Tmem98,Rnd2,Map3k13,Tiam1,Bcl11a,Tenm4,Nkx6-2,Plxna3,Neurl1,Plxnd1,Lrp6,Ufl1,Dmrta2,Kdm4a,Lpin1,Lingo1,Ptprf,RGD1307443,Adgre5,Tppp,Ankrd27,Caprin1,Macf1,Qk,Fxn,Lrrtm4,Bex1,Tcf7l2,Twf2 |
| GO:0048667 | cell morphogenesis involved in neuron differentiation | -7,54245 | -5,166 | 111/652 | Apc,Ncam1,Prkca,Sod1,Cck,Cntn2,Camk2a,Nog,Ptprs,Tnr,Agrn,Map2,Psen1,Stmn1,Ntrk3,Adam10,Sema3a,Atp5mc1,L1cam,Chrnb2,Arc,Notch3,Robo1,Map2k2,Unc5a,Unc5b,Rpl4,Rpl24,Isl1,Mfn2,Snap91,Dpysl5,Shank1,Alcam,Celsr2,Celsr3,Pafah1b1,Tpbg,Kalrn,Chn1,Picalm,Ptch1,Cdhr1,Dnm1l,Ntn1,Nlgn1,Cntn4,Mt3,Tenm2,Gli3,Bmpr2,Ndel1,Akap5,Shank2,Opa1,Epha7,Nlgn3,Dnm3,Syngap1,Ss18l1,Nexn,Lgi1,Lzts3,Lgr4,Tbc1d24,Metrn,Plxna2,Actr2,Sema3g,Tubb2b,Hecw1,Numbl,Tnik,Amigo1,Ntng1,Lhx2,Flrt2,Hexa,Mapk8ip3,Rnd2,Tanc2,Map3k13,Tiam1,Adcy1,Bcl11a,Kif13b,Efnb2,Ripor2,Plxna3,Mef2a,Btbd3,Abl1,Plxnd1,Whrn,Zswim5,Pip5k1c,Hecw2,Ptprf,Etv4,RGD1307443,Ust,Ankrd27,Caprin1,Macf1,Ppfia2,Rbfox2,Raph1,Zdhhc17,Fxn,Dtnbp1,Twf2 |
| GO:0050769 | positive regulation of neurogenesis | -6,93718 | -4,597 | 60/295 | Ace,Egfr,Gfap,Grm5,Atxn1,Nog,Ntrk3,Atp5mc1,L1cam,Robo1,Map2k2,Kit,Rpl4,Mfn2,Snap91,Mdk,Pafah1b1,Kalrn,Ctnnb1,Synj1,Picalm,Dnm1l,Ntn1,Bin1,Gli3,Bmpr2,Ndel1,Akap5,Opa1,Ss18l1,Tbc1d24,Metrn,Plxna2,Actr2,Lig4,Numbl,Tnik,Amigo1,Smarcd3,Plag1,Jade2,Rnd2,Map3k13,Tiam1,Bcl11a,Tenm4,Nkx6-2,Plxna3,Neurl1,Plxnd1,Ufl1,Dmrta2,Ptprf,Ankrd27,Caprin1,Macf1,Qk,Fxn,Bex1,Twf2 |
| GO:0010720 | positive regulation of cell development | -6,64592 | -4,367 | 71/378 | Ace,Egfr,Gfap,Grm5,Atxn1,Nog,S1pr2,Ntrk3,Atp5mc1,Cspg5,Mapk9,L1cam,Crk,Robo1,Map2k2,Kit,Rpl4,Mfn2,Cldn5,Snap91,Mdk,S100a10,Pafah1b1,Kalrn,Ctnnb1,Synj1,Picalm,Dnm1l,Ntn1,Bin1,Gli3,Bmpr2,Ndel1,Akap5,Opa1,Ss18l1,Tbc1d24,Metrn,Plxna2,Actr2,Lig4,Numbl,Flna,Tnik,Amigo1,Smarcd3,Plag1,Jade2,Rnd2,Map3k13,Tiam1,Bcl11a,Cdh5,Tenm4,Nkx6-2,Plxna3,Neurl1,Prex1,Abl1,Plxnd1,Ufl1,Dmrta2,Ptprf,Tyrobp,Ankrd27,Caprin1,Macf1,Qk,Fxn,Bex1,Twf2 |
| GO:0061564 | axon development | -6,27059 | -4,017 | 91/535 | Apc,Jak2,Ncam1,Prkca,Cck,Cntn2,Nog,Ptprs,Tnr,Agrn,Map2,Ddr1,Psen1,Stmn1,Ntrk3,Sema3a,Atp5mc1,Cspg5,L1cam,Chrnb2,Notch3,Robo1,Map2k2,Unc5a,Unc5b,Rpl4,Rpl24,Isl1,Snap91,Dpysl5,Alcam,Mtr,Dynlt1,Celsr3,Pafah1b1,Kalrn,Chn1,Picalm,Ptch1,Kremen1,Ntn1,Cntn4,Mt3,Tenm2,Gli3,Bmpr2,Ndel1,Epha7,Nlgn3,Syngap1,Nexn,Lgi1,Lgr4,Tbc1d24,Metrn,Plxna2,Sema3g,Tubb2b,Numbl,Flna,Amigo1,Ntng1,Lhx2,Flrt2,Mapk8ip3,Rnd2,Map3k13,Tiam1,Adcy1,Bcl11a,Kif13b,Efnb2,Inpp5f,Plxna3,Lrig2,Abl1,Plxnd1,Zswim5,Pip5k1c,Nrep,Ptprf,Etv4,Stk24,RGD1307443,Ust,Svbp,Macf1,Raph1,Zdhhc17,Fxn,Twf2 |
| GO:0050767 | regulation of neurogenesis | -5,78318 | -3,601 | 78/450 | B2m,Ace,Egfr,Gfap,Grm5,Atxn1,Nog,Ptprs,Tnr,Map2,Psen1,Ntrk3,Sema3a,Atp5mc1,L1cam,Robo1,Map2k2,Kit,Rpl4,Mfn2,Snap91,Bhlhe40,Mdk,Dynlt1,Pafah1b1,Kalrn,Ctnnb1,Synj1,Picalm,Dnm1l,Ntn1,Bin1,Mt3,Gli3,Bmpr2,Ndel1,Akap5,Opa1,Epha7,Syngap1,Ss18l1,Atf5,Tbc1d24,Metrn,Plxna2,Actr2,Sema3g,Lig4,Numbl,Tnik,Amigo1,Lhx2,Smarcd3,Plag1,Jade2,Tmem98,Rnd2,Map3k13,Tiam1,Bcl11a,Tenm4,Nkx6-2,Plxna3,Neurl1,Plxnd1,Ufl1,Dmrta2,Kdm4a,Lingo1,Ptprf,RGD1307443,Ankrd27,Caprin1,Macf1,Qk,Fxn,Bex1,Twf2 |
| GO:0031346 | positive regulation of cell projection organization | -5,53734 | -3,388 | 77/449 | Agt,Apc,Dmd,Pik3r1,Agrn,Ntrk3,Pacsin1,Atp5mc1,L1cam,Robo1,Map2k2,Kit,Rpl4,Mfn2,Snap91,Mdk,Dynlt1,Pafah1b1,Nme2,Kalrn,Grip1,Kif3c,Picalm,Magi2,Dnm1l,Ntn1,Nlgn1,Tenm2,Lrrc7,Bmpr2,Ndel1,Akap5,Opa1,Dnm3,Nme1,Ss18l1,Tbc1d24,Metrn,Plxna2,Actr2,Numbl,Flna,Tnik,Amigo1,Cbfa2t2,Zmynd8,Sf3a2,Rnd2,Cdc42ep4,Dvl3,Map3k13,Tiam1,Rrn3,Bcl11a,Ripor2,Camk1d,Ap2a1,Plxna3,Neurl1,Fnbp1l,Abl1,Plxnd1,Lrp6,Tmem67,Plk5,Ptprf,Mien1,Stk24,Ankrd27,Iqgap1,Caprin1,Ccl19,Macf1,Qk,Fxn,Twf2,Crtc1 |
| GO:0050770 | regulation of axonogenesis | -4,46657 | -2,520 | 38/190 | Cntn2,Ptprs,Tnr,Map2,Psen1,Ntrk3,Sema3a,Atp5mc1,L1cam,Robo1,Map2k2,Rpl4,Snap91,Pafah1b1,Chn1,Picalm,Ntn1,Mt3,Bmpr2,Ndel1,Epha7,Syngap1,Metrn,Plxna2,Sema3g,Amigo1,Rnd2,Map3k13,Tiam1,Bcl11a,Kif13b,Plxna3,Plxnd1,RGD1307443,Ust,Macf1,Fxn,Twf2 |
| GO:0007409 | axonogenesis | -4,41182 | -2,474 | 77/482 | Apc,Ncam1,Prkca,Cck,Cntn2,Nog,Ptprs,Tnr,Agrn,Map2,Psen1,Stmn1,Ntrk3,Sema3a,Atp5mc1,L1cam,Chrnb2,Notch3,Robo1,Map2k2,Unc5a,Unc5b,Rpl4,Rpl24,Isl1,Snap91,Dpysl5,Alcam,Celsr3,Pafah1b1,Kalrn,Chn1,Picalm,Ptch1,Ntn1,Cntn4,Mt3,Tenm2,Gli3,Bmpr2,Ndel1,Epha7,Nlgn3,Syngap1,Nexn,Lgi1,Lgr4,Metrn,Plxna2,Sema3g,Tubb2b,Numbl,Amigo1,Ntng1,Lhx2,Flrt2,Mapk8ip3,Rnd2,Map3k13,Tiam1,Adcy1,Bcl11a,Kif13b,Efnb2,Plxna3,Abl1,Plxnd1,Zswim5,Pip5k1c,Etv4,RGD1307443,Ust,Macf1,Raph1,Zdhhc17,Fxn,Twf2 |
| GO:0050772 | positive regulation of axonogenesis | -4,09961 | -2,208 | 24/103 | Ntrk3,Atp5mc1,L1cam,Robo1,Map2k2,Rpl4,Snap91,Pafah1b1,Picalm,Ntn1,Bmpr2,Ndel1,Metrn,Plxna2,Amigo1,Rnd2,Map3k13,Tiam1,Bcl11a,Plxna3,Plxnd1,Macf1,Fxn,Twf2 |
| GO:1990138 | neuron projection extension | -4,06671 | -2,184 | 39/205 | Ptprs,Tnr,Map2,Ddr1,Syt1,Ntrk3,Sema3a,Atp5mc1,L1cam,Rpl4,Unc13a,Alcam,Pafah1b1,Ctnnb1,Picalm,Ntn1,Sh3gl2,Rims2,Mt3,Bmpr2,Ndel1,Nlgn3,Sema3g,Pak6,Lhx2,Jade2,Map3k13,Tiam1,Bcl11a,Plxna3,Abl1,RGD1307443,Iqgap1,Macf1,Raph1,Impact,Fxn,Twf2,Nrn1l |
| GO:0048588 | developmental cell growth | -4,01454 | -2,140 | 49/279 | Agt,Camk2d,Ptprs,Tnr,Map2,Ddr1,Syt1,Ntrk3,Sema3a,Atp5mc1,Gsk3a,L1cam,Rpl4,Akap6,Unc13a,Alcam,Pafah1b1,Fdps,Ctnnb1,Picalm,Ntn1,Sorbs2,Sh3gl2,Rims2,Mt3,Sox9,Bmpr2,Ndel1,Epha7,Nlgn3,Map2k4,Sema3g,Pak6,Lhx2,Jade2,Rnd2,Map3k13,Tiam1,Bcl11a,Plxna3,Abl1,RGD1307443,Iqgap1,Macf1,Raph1,Impact,Fxn,Twf2,Nrn1l |
| GO:0060560 | developmental growth involved in morphogenesis | -2,9202 | -1,293 | 45/279 | Ptprs,Tnr,Map2,Ddr1,Syt1,Ntrk3,Sema3a,Atp5mc1,L1cam,Robo1,Rpl4,Unc13a,Alcam,Pafah1b1,Ctnnb1,Picalm,Ntn1,Sh3gl2,Rims2,Mt3,Sox9,Bmpr2,Ndel1,Epha7,Nlgn3,Sema3g,Pak6,Lhx2,Jade2,Rnd2,Map3k13,Tiam1,Bcl11a,Plxna3,Abl1,Lrp6,RGD1307443,Iqgap1,Macf1,Raph1,Impact,Fxn,Med12,Twf2,Nrn1l |
| GO:0030307 | positive regulation of cell growth | -2,39419 | -0,913 | 32/194 | Egfr,Syt1,Ntrk3,Adam10,Atp5mc1,L1cam,Crk,Extl3,Rpl4,Akap6,Unc13a,Cyba,Mtpn,Pafah1b1,Fdps,Picalm,Ntn1,Csnk2a1,Rims2,Bmpr2,Ndel1,Dnph1,Lgi1,Krt17,Rnd2,Map3k13,Bcl11a,Macf1,Hyal1,Fxn,Twf2,Slc25a33 |
| GO:0061387 | regulation of extent of cell growth | -2,38083 | -0,900 | 24/134 | Ptprs,Tnr,Map2,Ntrk3,Sema3a,Atp5mc1,L1cam,Rpl4,Pafah1b1,Ntn1,Mt3,Bmpr2,Ndel1,Epha7,Sema3g,Rnd2,Map3k13,Bcl11a,Plxna3,Abl1,RGD1307443,Macf1,Fxn,Twf2 |
| GO:0048675 | axon extension | -2,23595 | -0,808 | 25/145 | Ptprs,Tnr,Map2,Ntrk3,Sema3a,Atp5mc1,L1cam,Rpl4,Alcam,Pafah1b1,Ntn1,Mt3,Bmpr2,Ndel1,Nlgn3,Sema3g,Lhx2,Map3k13,Plxna3,Abl1,RGD1307443,Macf1,Raph1,Fxn,Twf2 |
| GO:0030516 | regulation of axon extension | -2,07852 | -0,691 | 21/119 | Ptprs,Tnr,Map2,Ntrk3,Sema3a,Atp5mc1,L1cam,Rpl4,Pafah1b1,Ntn1,Mt3,Bmpr2,Ndel1,Sema3g,Map3k13,Plxna3,Abl1,RGD1307443,Macf1,Fxn,Twf2 |
| GO:0042391 | regulation of membrane potential | -10,7153 | -7,980 | 97/479 | Agt,B2m,Cacna1c,Camk2d,Grin2a,Grm5,Sod1,Dmd,Gabrb3,Stx1b,P2rx6,Atxn1,Chrm1,Akt2,Dlg1,Cck,Scn2b,Atp1b3,Atp5if1,Gabrb2,Glrb,Kcna5,Agrn,Psen1,Dpp6,Adora1,Oprk1,Prkce,S1pr2,Ntrk3,Kcnq3,Gabrg2,Scn8a,Cacna1d,Kcnj10,Kcnab2,Atp5mc1,Bok,Gria1,Chrnb2,Grik2,Cacna1i,Cacnb4,Npff,Fgf14,Mfn2,Akap6,Unc13b,Snap91,Kcnmb4,Shank1,Gabrg3,Grid2,Scn1a,Abat,Gna11,Gnaq,Sh3gl1,Kcnma1,Slc4a4,Slc25a27,Mpp2,Cxadr,Kcnh4,Hcn3,Cacna1h,Nlgn1,Rims2,Bin1,Nlgn2,Park7,Slc8a2,Slc8a3,Gabra4,Oga,Kcnq2,Shank2,Nlgn3,Npas4,Tbc1d24,Gabra2,Slmap,Ndufc2,Flna,Zmynd8,Grik3,Trem2,Neto1,Neto2,Ppa2,Slc4a8,Bscl2,Scn3a,Crtc1,Tafa4,Ryr2,Slc25a33,Ace,Egfr,Jak2,Ncam1,Pde4b,Pomc,Prkca,Calcrl,Sp4,Htr2c,Pik3r1,Slc1a1,Abcc9,Tnr,Utrn,Hba-a1,S100b,Gal,Tnni3,Acvr2a,Npy1r,Aif1,Epas1,Ptgs2,Htr2a,Atp2b4,Sema3a,Ctss,Gnao1,Ednrb,Gsk3a,Cacna1e,Glrx3,Kit,Isl1,Cyba,Mtpn,Ccn3,Ppargc1a,Fdps,Foxo1,Ece1,Akap1,Dlgap2,Bmpr2,Cacng4,Fabp5,Camk2g,Tnnt1,Cacna1b,Nptx2,Fto,Foxo3,Snx5,Tnnc2,Cacna2d2,Tmem98,Zfhx2,Tenm4,Mef2a,Abl1,Pbx3,Abcg2,Lpin1,Tppp,Rbfox2,Ecrg4,Gucy1a1,Shisa7 |
| GO:0042391 | regulation of membrane potential | -10,7153 | -7,980 | 97/479 | Agt,B2m,Cacna1c,Camk2d,Grin2a,Grm5,Sod1,Dmd,Gabrb3,Stx1b,P2rx6,Atxn1,Chrm1,Akt2,Dlg1,Cck,Scn2b,Atp1b3,Atp5if1,Gabrb2,Glrb,Kcna5,Agrn,Psen1,Dpp6,Adora1,Oprk1,Prkce,S1pr2,Ntrk3,Kcnq3,Gabrg2,Scn8a,Cacna1d,Kcnj10,Kcnab2,Atp5mc1,Bok,Gria1,Chrnb2,Grik2,Cacna1i,Cacnb4,Npff,Fgf14,Mfn2,Akap6,Unc13b,Snap91,Kcnmb4,Shank1,Gabrg3,Grid2,Scn1a,Abat,Gna11,Gnaq,Sh3gl1,Kcnma1,Slc4a4,Slc25a27,Mpp2,Cxadr,Kcnh4,Hcn3,Cacna1h,Nlgn1,Rims2,Bin1,Nlgn2,Park7,Slc8a2,Slc8a3,Gabra4,Oga,Kcnq2,Shank2,Nlgn3,Npas4,Tbc1d24,Gabra2,Slmap,Ndufc2,Flna,Zmynd8,Grik3,Trem2,Neto1,Neto2,Ppa2,Slc4a8,Bscl2,Scn3a,Crtc1,Tafa4,Ryr2,Slc25a33 |
| GO:0044057 | regulation of system process | -8,01191 | -5,584 | 112/647 | Agt,Cacna1c,Camk2d,Ace,Egfr,Grin2a,Jak2,Ncam1,Pde4b,Pomc,Prkca,Sod1,Dmd,Stx1b,Calcrl,Sp4,Htr2c,Dlg1,Cck,Scn2b,Kcna5,Pik3r1,Slc1a1,Abcc9,Tnr,Agrn,Utrn,Hba-a1,S100b,Gal,Tnni3,Acvr2a,Adora1,Oprk1,Npy1r,S1pr2,Aif1,Epas1,Ptgs2,Htr2a,Atp2b4,Cacna1d,Kcnj10,Sema3a,Ctss,Gnao1,Ednrb,Gsk3a,Cacna1e,Glrx3,Npff,Kit,Isl1,Mfn2,Akap6,Unc13b,Shank1,Cyba,Mtpn,Ccn3,Abat,Sh3gl1,Ppargc1a,Kcnma1,Fdps,Foxo1,Cxadr,Ece1,Akap1,Cacna1h,Nlgn1,Dlgap2,Rims2,Bin1,Nlgn2,Slc8a2,Slc8a3,Bmpr2,Cacng4,Fabp5,Oga,Shank2,Camk2g,Nlgn3,Tnnt1,Cacna1b,Tbc1d24,Nptx2,Fto,Foxo3,Snx5,Tnnc2,Zmynd8,Cacna2d2,Tmem98,Zfhx2,Neto1,Neto2,Tenm4,Mef2a,Abl1,Pbx3,Abcg2,Lpin1,Tppp,Bscl2,Rbfox2,Ecrg4,Gucy1a1,Tafa4,Ryr2,Shisa7 |
| GO:0060078 | regulation of postsynaptic membrane potential | -7,68921 | -5,291 | 41/161 | Cacna1c,Grin2a,Grm5,Gabrb3,Stx1b,P2rx6,Atxn1,Chrm1,Gabrb2,Glrb,Adora1,S1pr2,Gabrg2,Gria1,Chrnb2,Grik2,Npff,Fgf14,Unc13b,Shank1,Gabrg3,Grid2,Abat,Sh3gl1,Mpp2,Nlgn1,Rims2,Nlgn2,Slc8a2,Slc8a3,Gabra4,Shank2,Nlgn3,Npas4,Tbc1d24,Gabra2,Zmynd8,Grik3,Neto1,Neto2,Bscl2 |
| GO:0031644 | regulation of nervous system process | -6,75189 | -4,441 | 44/192 | Agt,Grin2a,Ncam1,Stx1b,Htr2c,Cck,Tnr,Hba-a1,S100b,Adora1,Oprk1,S1pr2,Kcnj10,Ctss,Ednrb,Npff,Unc13b,Shank1,Ccn3,Abat,Sh3gl1,Nlgn1,Dlgap2,Rims2,Nlgn2,Slc8a2,Slc8a3,Cacng4,Fabp5,Shank2,Nlgn3,Tbc1d24,Nptx2,Zmynd8,Tmem98,Zfhx2,Neto1,Neto2,Tenm4,Lpin1,Tppp,Bscl2,Tafa4,Shisa7 |
| GO:0099565 | chemical synaptic transmission, postsynaptic | -5,16475 | -3,079 | 30/126 | Grin2a,Gabrb3,Stx1b,P2rx6,Atxn1,Glrb,Adora1,S1pr2,Chrnb2,Grik2,Npff,Unc13b,Shank1,Grid2,Abat,Sh3gl1,Mpp2,Nlgn1,Rims2,Nlgn2,Slc8a2,Slc8a3,Shank2,Nlgn3,Npas4,Tbc1d24,Zmynd8,Neto1,Neto2,Bscl2 |
| GO:0098815 | modulation of excitatory postsynaptic potential | -4,49485 | -2,538 | 17/57 | Grin2a,Stx1b,S1pr2,Shank1,Sh3gl1,Nlgn1,Rims2,Nlgn2,Slc8a2,Slc8a3,Shank2,Nlgn3,Tbc1d24,Zmynd8,Neto1,Neto2,Bscl2 |
| GO:0060079 | excitatory postsynaptic potential | -4,4542 | -2,509 | 27/117 | Grin2a,Stx1b,P2rx6,Atxn1,Glrb,Adora1,S1pr2,Chrnb2,Grik2,Npff,Shank1,Grid2,Sh3gl1,Mpp2,Nlgn1,Rims2,Nlgn2,Slc8a2,Slc8a3,Shank2,Nlgn3,Npas4,Tbc1d24,Zmynd8,Neto1,Neto2,Bscl2 |
| GO:2000463 | positive regulation of excitatory postsynaptic potential | -2,88252 | -1,264 | 11/39 | Grin2a,Stx1b,Shank1,Nlgn1,Rims2,Nlgn2,Shank2,Nlgn3,Tbc1d24,Neto1,Neto2 |
| GO:0007610 | behaviour | -9,51079 | -6,922 | 133/766 | Agt,B2m,Cacna1c,Cebpb,Ace,Egfr,Egr1,Grin2a,Grm5,Ncam1,Pomc,Ppp3cb,Prkca,Prkcg,Sod1,Gabrb3,Myo5a,Dbi,Atxn1,Htr1b,Htr2c,P2ry1,Cck,Cort,Cstb,Cntn2,Glrb,Nog,Slc1a1,Tnr,S100b,Gal,Strn,Psen1,Penk,Adora1,Oprk1,Prkce,Npy1r,Selenop,Ptgs2,Htr2a,Pde1b,Gabrg2,Scn8a,Cacna1d,Kcnj10,Gria1,Nr2c2,Gnao1,Cacna1e,Chrnb2,Grik2,Arc,Csnk1e,Cacnb4,Synpo,Atxn3,Tsc1,Syt11,Amph,Fgf14,Lgmn,Kit,Pcdh8,Kcnip3,Shank1,Per3,Six3,Mdk,Scn1a,Abat,Gnaq,Grm7,Pafah1b1,Apba1,Apba2,Tpbg,Ncoa2,Wfs1,Kcnma1,Calb1,Kalrn,Clcn3,Slc24a2,Synj1,Picalm,Nlgn1,Nlgn2,Park7,Slc8a2,Slc8a3,Gli3,Adgrl3,Shank2,Nlgn3,Syngap1,Gng8,Cacna1b,Npas4,Ift20,Nptx2,Mmp17,Actr2,Fuom,Slc4a10,Pak6,Tcf15,Nts,Myg1,Hexa,Sez6l,Adcy1,Zfhx2,Pcdh17,Neto1,Inpp5f,Vps13a,Arrdc3,Pde8b,Tanc1,Abl1,Pbx3,Gpr157,Unc79,Idua,Eif4ebp2,Csmd1,Ciart,Fxn,Crtc1,Tpgs1,Shisa7,Chrm1,Mgat3,Setd5,Cbr3,Slc1a4 |
| GO:0007610 | behaviour | -9,51079 | -6,922 | 133/766 | Agt,B2m,Cacna1c,Cebpb,Ace,Egfr,Egr1,Grin2a,Grm5,Ncam1,Pomc,Ppp3cb,Prkca,Prkcg,Sod1,Gabrb3,Myo5a,Dbi,Atxn1,Htr1b,Htr2c,P2ry1,Cck,Cort,Cstb,Cntn2,Glrb,Nog,Slc1a1,Tnr,S100b,Gal,Strn,Psen1,Penk,Adora1,Oprk1,Prkce,Npy1r,Selenop,Ptgs2,Htr2a,Pde1b,Gabrg2,Scn8a,Cacna1d,Kcnj10,Gria1,Nr2c2,Gnao1,Cacna1e,Chrnb2,Grik2,Arc,Csnk1e,Cacnb4,Synpo,Atxn3,Tsc1,Syt11,Amph,Fgf14,Lgmn,Kit,Pcdh8,Kcnip3,Shank1,Per3,Six3,Mdk,Scn1a,Abat,Gnaq,Grm7,Pafah1b1,Apba1,Apba2,Tpbg,Ncoa2,Wfs1,Kcnma1,Calb1,Kalrn,Clcn3,Slc24a2,Synj1,Picalm,Nlgn1,Nlgn2,Park7,Slc8a2,Slc8a3,Gli3,Adgrl3,Shank2,Nlgn3,Syngap1,Gng8,Cacna1b,Npas4,Ift20,Nptx2,Mmp17,Actr2,Fuom,Slc4a10,Pak6,Tcf15,Nts,Myg1,Hexa,Sez6l,Adcy1,Zfhx2,Pcdh17,Neto1,Inpp5f,Vps13a,Arrdc3,Pde8b,Tanc1,Abl1,Pbx3,Gpr157,Unc79,Idua,Eif4ebp2,Csmd1,Ciart,Fxn,Crtc1,Tpgs1,Shisa7 |
| GO:0050890 | cognition | -7,71613 | -5,312 | 73/370 | Agt,B2m,Cacna1c,Cebpb,Egfr,Egr1,Grin2a,Grm5,Ncam1,Prkca,Prkcg,Gabrb3,Dbi,Atxn1,Chrm1,Cck,Cntn2,Nog,Slc1a1,Tnr,S100b,Psen1,Adora1,Oprk1,Ptgs2,Mgat3,Htr2a,Pde1b,Cacna1d,Gria1,Cacna1e,Chrnb2,Arc,Synpo,Syt11,Amph,Lgmn,Kit,Pcdh8,Shank1,Mdk,Grm7,Pafah1b1,Tpbg,Calb1,Kalrn,Slc24a2,Synj1,Picalm,Slc8a2,Slc8a3,Shank2,Nlgn3,Syngap1,Npas4,Ift20,Nptx2,Actr2,Pak6,Setd5,Nts,Cbr3,Adcy1,Slc1a4,Neto1,Pde8b,Tanc1,Abl1,Idua,Eif4ebp2,Csmd1,Crtc1,Shisa7 |
| GO:0007611 | learning or memory | -7,63803 | -5,245 | 67/330 | Agt,B2m,Cacna1c,Cebpb,Egfr,Egr1,Grin2a,Grm5,Ncam1,Prkca,Prkcg,Gabrb3,Dbi,Atxn1,Cck,Cntn2,Nog,Slc1a1,Tnr,S100b,Psen1,Oprk1,Ptgs2,Htr2a,Pde1b,Cacna1d,Gria1,Cacna1e,Chrnb2,Arc,Synpo,Syt11,Amph,Lgmn,Kit,Pcdh8,Shank1,Mdk,Grm7,Pafah1b1,Tpbg,Calb1,Kalrn,Slc24a2,Synj1,Picalm,Slc8a2,Slc8a3,Shank2,Nlgn3,Syngap1,Npas4,Ift20,Nptx2,Actr2,Pak6,Nts,Adcy1,Neto1,Pde8b,Tanc1,Abl1,Idua,Eif4ebp2,Csmd1,Crtc1,Shisa7 |
| GO:0007612 | learning | -7,26684 | -4,912 | 45/191 | Agt,Cacna1c,Grin2a,Grm5,Gabrb3,Atxn1,Cck,Cntn2,Nog,Slc1a1,Tnr,Oprk1,Ptgs2,Pde1b,Cacna1e,Chrnb2,Arc,Synpo,Syt11,Amph,Lgmn,Kit,Shank1,Grm7,Tpbg,Kalrn,Slc24a2,Synj1,Slc8a2,Slc8a3,Shank2,Nlgn3,Syngap1,Npas4,Ift20,Nptx2,Actr2,Pak6,Nts,Neto1,Pde8b,Tanc1,Abl1,Idua,Csmd1 |
| GO:0008306 | associative learning | -5,87441 | -3,667 | 30/117 | Agt,Cacna1c,Grin2a,Atxn1,Cck,Nog,Slc1a1,Tnr,Oprk1,Pde1b,Cacna1e,Chrnb2,Synpo,Lgmn,Kit,Shank1,Grm7,Tpbg,Nlgn3,Syngap1,Ift20,Nptx2,Actr2,Nts,Neto1,Pde8b,Tanc1,Abl1,Idua,Csmd1 |
| GO:0007613 | memory | -5,48092 | -3,341 | 37/166 | Cebpb,Egr1,Grin2a,Gabrb3,Atxn1,Cck,Nog,Slc1a1,S100b,Psen1,Ptgs2,Htr2a,Cacna1d,Gria1,Chrnb2,Arc,Syt11,Lgmn,Pcdh8,Shank1,Mdk,Grm7,Calb1,Kalrn,Slc24a2,Slc8a2,Slc8a3,Shank2,Npas4,Pak6,Adcy1,Neto1,Idua,Eif4ebp2,Csmd1,Crtc1,Shisa7 |
| GO:0008542 | visual learning | -3,62539 | -1,830 | 18/72 | Cacna1c,Grin2a,Atxn1,Cck,Nog,Pde1b,Cacna1e,Chrnb2,Synpo,Kit,Nlgn3,Syngap1,Ift20,Nts,Neto1,Pde8b,Tanc1,Idua |
| GO:0007632 | visual behaviour | -3,31546 | -1,595 | 18/76 | Cacna1c,Grin2a,Atxn1,Cck,Nog,Pde1b,Cacna1e,Chrnb2,Synpo,Kit,Nlgn3,Syngap1,Ift20,Nts,Neto1,Pde8b,Tanc1,Idua |
| GO:0036465 | synaptic vesicle recycling | -9,39746 | -6,826 | 30/85 | Mx1,Ppp3cb,Syp,Stx1b,Syt1,Pacsin1,Syt7,Syt11,Amph,Fgf14,Rab5a,Scamp5,Snap91,Sh3gl1,Synj1,Picalm,Dnm1l,Cltb,Nlgn1,Sh3gl2,Bin1,Nlgn2,Ap2b1,Nlgn3,Dnm3,Tbc1d24,Fcho2,Dnajc6,Ston2,Pip5k1c,Arrb1,Nedd8,Vamp3,Ubc,Syt9,Ap2s1,Vamp8,Fnbp1,Actr2,Arpc4,Ap2a1,Fnbp1l,Stam2,Pacsin3,RGD1307443,Rps27a,Inpp5f,Arf1,Clint1 |
| GO:0036465 | synaptic vesicle recycling | -9,39746 | -6,826 | 30/85 | Mx1,Ppp3cb,Syp,Stx1b,Syt1,Pacsin1,Syt7,Syt11,Amph,Fgf14,Rab5a,Scamp5,Snap91,Sh3gl1,Synj1,Picalm,Dnm1l,Cltb,Nlgn1,Sh3gl2,Bin1,Nlgn2,Ap2b1,Nlgn3,Dnm3,Tbc1d24,Fcho2,Dnajc6,Ston2,Pip5k1c |
| GO:0048488 | synaptic vesicle endocytosis | -8,74374 | -6,257 | 27/75 | Mx1,Ppp3cb,Syp,Syt1,Pacsin1,Syt7,Syt11,Amph,Scamp5,Snap91,Sh3gl1,Synj1,Picalm,Dnm1l,Cltb,Nlgn1,Sh3gl2,Bin1,Nlgn2,Ap2b1,Nlgn3,Dnm3,Tbc1d24,Fcho2,Dnajc6,Ston2,Pip5k1c |
| GO:0140238 | presynaptic endocytosis | -8,74374 | -6,257 | 27/75 | Mx1,Ppp3cb,Syp,Syt1,Pacsin1,Syt7,Syt11,Amph,Scamp5,Snap91,Sh3gl1,Synj1,Picalm,Dnm1l,Cltb,Nlgn1,Sh3gl2,Bin1,Nlgn2,Ap2b1,Nlgn3,Dnm3,Tbc1d24,Fcho2,Dnajc6,Ston2,Pip5k1c |
| R-RNO-8856828 | Clathrin-mediated endocytosis | -6,2049 | -3,975 | 32/125 | Arrb1,Nedd8,Syt1,Vamp3,Pacsin1,Ubc,Syt9,Syt11,Amph,Ap2s1,Snap91,Sh3gl1,Vamp8,Synj1,Picalm,Cltb,Sh3gl2,Bin1,Ap2b1,Dnm3,Fnbp1,Actr2,Arpc4,Ap2a1,Fnbp1l,Stam2,Pacsin3,Dnajc6,Ston2,Pip5k1c,RGD1307443,Rps27a |
| GO:1903421 | regulation of synaptic vesicle recycling | -5,19652 | -3,105 | 13/32 | Ppp3cb,Stx1b,Syt7,Syt11,Fgf14,Scamp5,Snap91,Sh3gl1,Picalm,Dnm1l,Nlgn1,Dnm3,Pip5k1c |
| GO:1900242 | regulation of synaptic vesicle endocytosis | -4,052 | -2,171 | 10/25 | Ppp3cb,Syt7,Syt11,Scamp5,Snap91,Sh3gl1,Picalm,Dnm1l,Nlgn1,Pip5k1c |
| R-RNO-8856825 | Cargo recognition for clathrin-mediated endocytosis | -3,23193 | -1,529 | 20/90 | Arrb1,Nedd8,Syt1,Vamp3,Ubc,Syt9,Syt11,Ap2s1,Snap91,Sh3gl1,Vamp8,Picalm,Cltb,Sh3gl2,Ap2b1,Ap2a1,Stam2,Ston2,RGD1307443,Rps27a |
| GO:0072583 | clathrin-dependent endocytosis | -2,95466 | -1,319 | 12/44 | Syt11,Ap2s1,Snap91,Picalm,Cltb,Sh3gl2,Ap2b1,Ap2a1,Inpp5f,Fcho2,Fnbp1l,Dnajc6 |
| GO:1903423 | positive regulation of synaptic vesicle recycling | -2,7985 | -1,201 | 6/14 | Snap91,Sh3gl1,Picalm,Dnm1l,Nlgn1,Dnm3 |
| GO:0016185 | synaptic vesicle budding from presynaptic endocytic zone membrane | -2,75764 | -1,172 | 5/10 | Mx1,Snap91,Picalm,Cltb,Dnm3 |
| GO:0070142 | synaptic vesicle budding | -2,6154 | -1,068 | 6/15 | Mx1,Arf1,Snap91,Picalm,Cltb,Dnm3 |
| GO:1900244 | positive regulation of synaptic vesicle endocytosis | -2,33573 | -0,871 | 5/12 | Snap91,Sh3gl1,Picalm,Dnm1l,Nlgn1 |
| GO:0048268 | clathrin coat assembly | -2,0354 | -0,678 | 6/19 | Snap91,Picalm,Cltb,Ap2b1,Fcho2,Clint1 |
| GO:0072657 | protein localization to membrane | -9,32414 | -6,769 | 110/600 | Camk2d,Egfr,Grin2a,Stx1b,Chm,Myo5a,Akt2,Dlg1,P2ry1,Fnta,Atp1b3,Camk2a,Glrb,Pik3r1,Rab12,Scp2,Slc1a1,Agrn,Tub,Dpp6,Adora1,Vamp3,Atp2b4,Adam10,Pacsin1,Grik2,Sec23a,Dlg3,Stx8,Nsf,Lin7b,Lin7c,Stx7,Timm10,Kcnip3,Myo1c,S100a10,Stx3,Tpbg,Vamp8,Kalrn,Grip1,Picalm,Ptch1,Magi2,Sorbs2,Csnk2a1,Nlgn1,Exoc4,Pgap2,Epb41l3,Nlgn2,Lrrc7,Ap2b1,Cacng3,Cacng4,Oga,Akap5,Camk2g,Syngap1,Phaf1,Lgi1,Timm13,Kcnip4,Ift20,Nptx2,Fis1,Rilpl2,Commd1,Slmap,Srp19,Ppfia1,Flna,Sgtb,Sec62,Tnik,Srp14,Zmynd8,Rer1,Rab15,Abhd17a,Trem2,Rapgef6,Tiam1,Ap4m1,Wdr19,Sptbn1,Zdhhc20,Ank1,Rab11fip2,Dchs1,Fcho2,Ssna1,Ift122,Lrp6,Pals1,Fam126b,Lhfpl4,Emc8,Zdhhc5,Macf1,Zdhhc23,Dpp10,Fam126a,Lrrtm4,Iqsec2,Sec61g,Srp9,Shisa7,Ndufa13,Gnai1,Kif1b,Numa1,Apc,Jak2,Chrm1,Atp5if1,Oaz1,Map2,Psen1,Cct4,Ptgs2,Mgat3,Kcnab2,Gsk3a,Arc,Csnk1e,Cacnb4,Syt11,Cdk5rap3,Polr1a,Ctnnb1,Ran,Synj1,Xpo1,Akap1,Pkia,Park7,Gli3,Ifi27,Prkcd,Anp32b,Ndel1,Dkc1,Cabp1,Uhmk1,Vcpip1,Ppm1f,B3gat3,Ddrgk1,Ube2j1,Ube2j2,Tmem98,Dvl3,Efnb2,Ripor2,Ctdspl2,Pip5k1c,Cd2ap,Fzd5,Gas8,Tyrobp,Iqgap1,Bicd1,Rangap1,Ufm1,Dynlt2b,Ubl5,Tcf7l2,Tomm7,Larp7,Gnl3l |
| GO:0072657 | protein localization to membrane | -9,32414 | -6,769 | 110/600 | Camk2d,Egfr,Grin2a,Stx1b,Chm,Myo5a,Akt2,Dlg1,P2ry1,Fnta,Atp1b3,Camk2a,Glrb,Pik3r1,Rab12,Scp2,Slc1a1,Agrn,Tub,Dpp6,Adora1,Vamp3,Atp2b4,Adam10,Pacsin1,Grik2,Sec23a,Dlg3,Stx8,Nsf,Lin7b,Lin7c,Stx7,Timm10,Kcnip3,Myo1c,S100a10,Stx3,Tpbg,Vamp8,Kalrn,Grip1,Picalm,Ptch1,Magi2,Sorbs2,Csnk2a1,Nlgn1,Exoc4,Pgap2,Epb41l3,Nlgn2,Lrrc7,Ap2b1,Cacng3,Cacng4,Oga,Akap5,Camk2g,Syngap1,Phaf1,Lgi1,Timm13,Kcnip4,Ift20,Nptx2,Fis1,Rilpl2,Commd1,Slmap,Srp19,Ppfia1,Flna,Sgtb,Sec62,Tnik,Srp14,Zmynd8,Rer1,Rab15,Abhd17a,Trem2,Rapgef6,Tiam1,Ap4m1,Wdr19,Sptbn1,Zdhhc20,Ank1,Rab11fip2,Dchs1,Fcho2,Ssna1,Ift122,Lrp6,Pals1,Fam126b,Lhfpl4,Emc8,Zdhhc5,Macf1,Zdhhc23,Dpp10,Fam126a,Lrrtm4,Iqsec2,Sec61g,Srp9,Shisa7,Ndufa13 |
| GO:1990778 | protein localization to cell periphery | -8,96986 | -6,454 | 77/374 | Camk2d,Egfr,Grin2a,Stx1b,Myo5a,Akt2,Dlg1,P2ry1,Atp1b3,Camk2a,Pik3r1,Rab12,Scp2,Slc1a1,Tub,Gnai1,Dpp6,Vamp3,Atp2b4,Adam10,Pacsin1,Sec23a,Stx8,Nsf,Lin7b,Lin7c,Stx7,Kcnip3,S100a10,Stx3,Tpbg,Vamp8,Kalrn,Grip1,Picalm,Ptch1,Magi2,Csnk2a1,Pgap2,Epb41l3,Kif1b,Cacng3,Cacng4,Akap5,Camk2g,Phaf1,Lgi1,Kcnip4,Ift20,Nptx2,Rilpl2,Commd1,Slmap,Ppfia1,Flna,Tnik,Rer1,Rab15,Trem2,Rapgef6,Ap4m1,Wdr19,Sptbn1,Ank1,Rab11fip2,Numa1,Dchs1,Fcho2,Lrp6,Pals1,Fam126b,Zdhhc5,Macf1,Zdhhc23,Dpp10,Fam126a,Iqsec2 |
| GO:1903827 | regulation of cellular protein localization | -6,69331 | -4,410 | 102/607 | Apc,Camk2d,Egfr,Jak2,Myo5a,Chrm1,Akt2,Dlg1,Atp5if1,Camk2a,Oaz1,Pik3r1,Slc1a1,Map2,Gnai1,Psen1,Dpp6,Cct4,Ptgs2,Mgat3,Atp2b4,Adam10,Pacsin1,Kcnab2,Gsk3a,Arc,Csnk1e,Cacnb4,Stx8,Stx7,Syt11,Myo1c,Cdk5rap3,Stx3,Polr1a,Vamp8,Kalrn,Ctnnb1,Ran,Synj1,Xpo1,Picalm,Magi2,Akap1,Sorbs2,Pkia,Csnk2a1,Nlgn2,Park7,Gli3,Ap2b1,Ifi27,Prkcd,Anp32b,Ndel1,Dkc1,Akap5,Cabp1,Camk2g,Uhmk1,Vcpip1,Ppm1f,Fis1,Commd1,Ppfia1,B3gat3,Flna,Tnik,Ddrgk1,Zmynd8,Ube2j1,Rer1,Ube2j2,Abhd17a,Trem2,Tmem98,Dvl3,Sptbn1,Efnb2,Ripor2,Rab11fip2,Numa1,Ctdspl2,Pip5k1c,Cd2ap,Fzd5,Gas8,Tyrobp,Iqgap1,Zdhhc5,Bicd1,Rangap1,Dpp10,Ufm1,Dynlt2b,Ubl5,Tcf7l2,Iqsec2,Tomm7,Larp7,Shisa7,Gnl3l |
| GO:0072659 | protein localization to plasma membrane | -6,2627 | -4,014 | 59/301 | Camk2d,Egfr,Myo5a,Akt2,Dlg1,P2ry1,Atp1b3,Camk2a,Pik3r1,Rab12,Scp2,Dpp6,Vamp3,Atp2b4,Pacsin1,Sec23a,Stx8,Nsf,Stx7,Kcnip3,S100a10,Stx3,Tpbg,Vamp8,Kalrn,Grip1,Picalm,Ptch1,Csnk2a1,Pgap2,Epb41l3,Akap5,Camk2g,Phaf1,Kcnip4,Ift20,Rilpl2,Commd1,Slmap,Ppfia1,Flna,Tnik,Rer1,Rab15,Trem2,Rapgef6,Sptbn1,Ank1,Rab11fip2,Dchs1,Fcho2,Lrp6,Pals1,Fam126b,Zdhhc5,Macf1,Zdhhc23,Dpp10,Fam126a |
| GO:1904375 | regulation of protein localization to cell periphery | -4,57464 | -2,602 | 31/141 | Camk2d,Egfr,Myo5a,Dlg1,Camk2a,Pik3r1,Gnai1,Dpp6,Atp2b4,Adam10,Stx8,Stx7,Stx3,Vamp8,Kalrn,Picalm,Magi2,Csnk2a1,Akap5,Camk2g,Commd1,Ppfia1,Tnik,Rer1,Trem2,Sptbn1,Rab11fip2,Numa1,Zdhhc5,Dpp10,Iqsec2 |
| GO:1905475 | regulation of protein localization to membrane | -3,78095 | -1,950 | 38/204 | Camk2d,Egfr,Myo5a,Akt2,Dlg1,Camk2a,Pik3r1,Slc1a1,Dpp6,Atp2b4,Adam10,Stx8,Stx7,Myo1c,Stx3,Vamp8,Kalrn,Picalm,Magi2,Sorbs2,Csnk2a1,Nlgn2,Ap2b1,Akap5,Camk2g,Fis1,Commd1,Ppfia1,Tnik,Zmynd8,Rer1,Trem2,Sptbn1,Rab11fip2,Zdhhc5,Dpp10,Iqsec2,Shisa7 |
| GO:1903076 | regulation of protein localization to plasma membrane | -3,72954 | -1,908 | 25/115 | Camk2d,Egfr,Myo5a,Dlg1,Camk2a,Pik3r1,Dpp6,Atp2b4,Stx8,Stx7,Stx3,Vamp8,Kalrn,Picalm,Csnk2a1,Akap5,Camk2g,Commd1,Ppfia1,Rer1,Trem2,Sptbn1,Rab11fip2,Zdhhc5,Dpp10 |
| GO:1903829 | positive regulation of cellular protein localization | -3,50163 | -1,738 | 51/308 | Apc,Egfr,Jak2,Myo5a,Akt2,Dlg1,Pik3r1,Gnai1,Psen1,Dpp6,Cct4,Ptgs2,Mgat3,Atp2b4,Gsk3a,Cacnb4,Syt11,Myo1c,Cdk5rap3,Stx3,Ran,Nlgn2,Park7,Gli3,Ap2b1,Prkcd,Dkc1,Akap5,Fis1,Commd1,Flna,Ddrgk1,Rer1,Ube2j2,Abhd17a,Trem2,Sptbn1,Rab11fip2,Numa1,Cd2ap,Fzd5,Gas8,Tyrobp,Iqgap1,Zdhhc5,Bicd1,Dpp10,Ubl5,Tomm7,Larp7,Gnl3l |
| GO:1904377 | positive regulation of protein localization to cell periphery | -2,83496 | -1,230 | 16/70 | Egfr,Myo5a,Dlg1,Gnai1,Dpp6,Atp2b4,Stx3,Akap5,Commd1,Rer1,Trem2,Sptbn1,Rab11fip2,Numa1,Zdhhc5,Dpp10 |
| GO:1903078 | positive regulation of protein localization to plasma membrane | -2,42949 | -0,932 | 14/63 | Egfr,Myo5a,Dlg1,Dpp6,Atp2b4,Stx3,Akap5,Commd1,Rer1,Trem2,Sptbn1,Rab11fip2,Zdhhc5,Dpp10 |
| GO:0048168 | regulation of neuronal synaptic plasticity | -7,5099 | -5,139 | 26/79 | Agt,Egr1,Grin2a,Grm5,Syp,Cntn2,Camk2a,S100b,Syngr1,Kcnj10,Grik2,Arc,Synpo,Kit,Rab5a,Unc13a,Unc13b,Bhlhe40,Kalrn,Slc8a2,Camk2g,Syngap1,Slc4a10,Neto1,Neurl1,Shisa7 |
| GO:0048168 | regulation of neuronal synaptic plasticity | -7,5099 | -5,139 | 26/79 | Agt,Egr1,Grin2a,Grm5,Syp,Cntn2,Camk2a,S100b,Syngr1,Kcnj10,Grik2,Arc,Synpo,Kit,Rab5a,Unc13a,Unc13b,Bhlhe40,Kalrn,Slc8a2,Camk2g,Syngap1,Slc4a10,Neto1,Neurl1,Shisa7 |
| GO:0048169 | regulation of long-term neuronal synaptic plasticity | -5,89481 | -3,684 | 15/37 | Agt,Egr1,Grin2a,Grm5,Syp,Syngr1,Kcnj10,Grik2,Synpo,Kit,Rab5a,Camk2g,Syngap1,Neto1,Neurl1 |
| GO:0048172 | regulation of short-term neuronal synaptic plasticity | -3,52104 | -1,754 | 8/19 | Syp,Syngr1,Grik2,Unc13a,Unc13b,Slc8a2,Slc4a10,Shisa7 |
| GO:0023061 | signal release | -7,49095 | -5,125 | 105/608 | Agt,Cacna1c,Egfr,Jak2,Pomc,Ppp3cb,Prkca,Prkcg,Syp,Stx1b,Myo5a,Htr1b,Htr2c,P2ry1,Arrb1,Camk2a,Rbp4,Syt1,Cadps,Gal,Psen1,Acvr2a,Adora1,Oprk1,Prkce,Npy1r,Bace1,Vamp3,Htr2a,Cacna1d,Cspg5,Mapk9,Cacna1e,Chrnb2,Pclo,Cacnb4,Syt7,Ensa,Npff,Nsf,Lin7b,Lin7c,Stx7,Syt9,Syt11,Isl1,Mfn2,Rab5a,Unc13a,Unc13b,Snap91,Syt13,Stxbp5,Ccn3,Abat,Grm7,Stx3,Doc2b,Apba1,Apba2,Vamp8,Kalrn,Synj1,Madd,Stxbp3,Dnm1l,Cltb,Nlgn1,Rims2,Nlgn2,Kcnc3,Park7,Ptpn23,Ap3m2,Oga,Trim9,Abcc4,Acvr1c,Cacna1b,Slc16a2,Unc13c,Stxbp5l,RT1-Db1,Mcu,Ddrgk1,Phpt1,Rab15,Tiam1,Adcy1,Rab11fip2,Pde8b,Abca1,Slc4a8,Myrip,Pck2,Uqcc2,Ppfia2,Ecrg4,Ufm1,Selenom,Napb,Dtnbp1,Tcf7l2,Mpc2,Enho,Sdc4,Cck,Cacna1i,Arf1,Scamp5,Gab2,Cacna1h,Rgcc,Zfp384,Ap1g1,Adam9,Trem2,Exoc1,Cd2ap,Myo18a,Vps4b,Chmp2a,Cyp51,Rab12,Rhbdd3,Idua,Dph3,Pafah1b1,Lyst,Akap5,Copg1,Sel1l,C1qtnf5,Svbp,Plek,Cbln4,Cd74,Ednrb,Cyba,Snx5,Chrm1,Akt2,Atp5if1,Oaz1,Pik3r1,Slc1a1,Ptgs2,Pacsin1,Gsk3a,Arc,Myo1c,Ran,Xpo1,Picalm,Akap1,Pkia,Exoc4,Gli3,Ifi27,Prkcd,Anp32b,Ndel1,Cabp1,Uhmk1,Ppm1f,Fis1,Commd1,B3gat3,Flna,Ube2j1,Ube2j2,Efnb2,Ctdspl2,Pip5k1c,Fzd5,Rangap1,Dynlt2b,Ubl5,Tomm7,Cct4,Dkc1,Nsd2 |
| GO:0023061 | signal release | -7,49095 | -5,125 | 105/608 | Agt,Cacna1c,Egfr,Jak2,Pomc,Ppp3cb,Prkca,Prkcg,Syp,Stx1b,Myo5a,Htr1b,Htr2c,P2ry1,Arrb1,Camk2a,Rbp4,Syt1,Cadps,Gal,Psen1,Acvr2a,Adora1,Oprk1,Prkce,Npy1r,Bace1,Vamp3,Htr2a,Cacna1d,Cspg5,Mapk9,Cacna1e,Chrnb2,Pclo,Cacnb4,Syt7,Ensa,Npff,Nsf,Lin7b,Lin7c,Stx7,Syt9,Syt11,Isl1,Mfn2,Rab5a,Unc13a,Unc13b,Snap91,Syt13,Stxbp5,Ccn3,Abat,Grm7,Stx3,Doc2b,Apba1,Apba2,Vamp8,Kalrn,Synj1,Madd,Stxbp3,Dnm1l,Cltb,Nlgn1,Rims2,Nlgn2,Kcnc3,Park7,Ptpn23,Ap3m2,Oga,Trim9,Abcc4,Acvr1c,Cacna1b,Slc16a2,Unc13c,Stxbp5l,RT1-Db1,Mcu,Ddrgk1,Phpt1,Rab15,Tiam1,Adcy1,Rab11fip2,Pde8b,Abca1,Slc4a8,Myrip,Pck2,Uqcc2,Ppfia2,Ecrg4,Ufm1,Selenom,Napb,Dtnbp1,Tcf7l2,Mpc2,Enho |
| GO:1903532 | positive regulation of secretion by cell | -5,66695 | -3,495 | 66/364 | Agt,Egfr,Jak2,Ppp3cb,Prkca,Sdc4,Stx1b,Htr2c,P2ry1,Cck,Arrb1,Rbp4,Syt1,Cadps,Gal,Acvr2a,Oprk1,Prkce,Cacna1d,Mapk9,Chrnb2,Cacna1i,Cacnb4,Syt7,Syt9,Arf1,Isl1,Mfn2,Rab5a,Unc13a,Unc13b,Scamp5,Stxbp5,Abat,Doc2b,Vamp8,Gab2,Dnm1l,Cacna1h,Nlgn1,Nlgn2,Rgcc,Ptpn23,Oga,Zfp384,Ap1g1,Cacna1b,Stxbp5l,Adam9,RT1-Db1,Mcu,Phpt1,Rab15,Trem2,Exoc1,Slc4a8,Cd2ap,Myrip,Myo18a,Vps4b,Pck2,Ecrg4,Chmp2a,Dtnbp1,Tcf7l2,Mpc2 |
| GO:1903530 | regulation of secretion by cell | -5,6369 | -3,475 | 110/702 | Agt,Egfr,Jak2,Pomc,Ppp3cb,Prkca,Prkcg,Sdc4,Syp,Stx1b,Myo5a,Htr1b,Htr2c,P2ry1,Cck,Arrb1,Camk2a,Cyp51,Rab12,Rbp4,Syt1,Cadps,Gal,Acvr2a,Adora1,Oprk1,Prkce,Npy1r,Bace1,Vamp3,Htr2a,Cacna1d,Cspg5,Mapk9,Cacna1e,Chrnb2,Pclo,Cacna1i,Cacnb4,Syt7,Ensa,Npff,Nsf,Syt9,Syt11,Arf1,Isl1,Mfn2,Rab5a,Unc13a,Unc13b,Scamp5,Syt13,Stxbp5,Ccn3,Abat,Grm7,Doc2b,Apba1,Apba2,Vamp8,Kalrn,Gab2,Madd,Stxbp3,Dnm1l,Cacna1h,Nlgn1,Rims2,Nlgn2,Kcnc3,Rgcc,Ptpn23,Oga,Trim9,Zfp384,Ap1g1,Acvr1c,Cacna1b,Unc13c,Stxbp5l,Rhbdd3,Adam9,RT1-Db1,Mcu,Ddrgk1,Phpt1,Rab15,Trem2,Tiam1,Exoc1,Adcy1,Pde8b,Slc4a8,Cd2ap,Myrip,Myo18a,Vps4b,Idua,Pck2,Uqcc2,Ppfia2,Ecrg4,Chmp2a,Ufm1,Dtnbp1,Tcf7l2,Dph3,Mpc2,Enho |
| GO:0009306 | protein secretion | -5,07851 | -3,008 | 69/401 | Cacna1c,Egfr,Jak2,Ppp3cb,Myo5a,Arrb1,Cyp51,Rab12,Rbp4,Prkce,Cacna1d,Cacna1e,Pclo,Syt7,Ensa,Npff,Syt9,Arf1,Isl1,Unc13b,Stxbp5,Ccn3,Abat,Doc2b,Pafah1b1,Vamp8,Lyst,Madd,Stxbp3,Dnm1l,Rims2,Nlgn2,Park7,Ptpn23,Oga,Zfp384,Akap5,Acvr1c,Stxbp5l,Rhbdd3,Adam9,RT1-Db1,Mcu,Ddrgk1,Phpt1,Copg1,Rab15,Trem2,Tiam1,Exoc1,Rab11fip2,Pde8b,Abca1,Sel1l,C1qtnf5,Cd2ap,Myrip,Myo18a,Idua,Pck2,Uqcc2,Svbp,Plek,Ufm1,Cbln4,Tcf7l2,Dph3,Mpc2,Enho |
| GO:0002790 | peptide secretion | -5,04707 | -2,988 | 76/455 | Cacna1c,Egfr,Jak2,Ppp3cb,Myo5a,Htr2c,Arrb1,Cyp51,Rab12,Cd74,Rbp4,Gal,Adora1,Prkce,Cacna1d,Cacna1e,Pclo,Syt7,Ensa,Npff,Syt9,Arf1,Isl1,Unc13b,Stxbp5,Ccn3,Abat,Doc2b,Pafah1b1,Vamp8,Kalrn,Lyst,Madd,Stxbp3,Dnm1l,Rims2,Nlgn2,Park7,Ptpn23,Oga,Zfp384,Akap5,Acvr1c,Slc16a2,Stxbp5l,Rhbdd3,Adam9,RT1-Db1,Mcu,Ddrgk1,Phpt1,Copg1,Rab15,Trem2,Tiam1,Exoc1,Rab11fip2,Pde8b,Abca1,Sel1l,C1qtnf5,Cd2ap,Myrip,Myo18a,Idua,Pck2,Uqcc2,Svbp,Ecrg4,Plek,Ufm1,Cbln4,Tcf7l2,Dph3,Mpc2,Enho |
| GO:0035592 | establishment of protein localization to extracellular region | -5,00385 | -2,955 | 69/403 | Cacna1c,Egfr,Jak2,Ppp3cb,Myo5a,Arrb1,Cyp51,Rab12,Rbp4,Prkce,Cacna1d,Cacna1e,Pclo,Syt7,Ensa,Npff,Syt9,Arf1,Isl1,Unc13b,Stxbp5,Ccn3,Abat,Doc2b,Pafah1b1,Vamp8,Lyst,Madd,Stxbp3,Dnm1l,Rims2,Nlgn2,Park7,Ptpn23,Oga,Zfp384,Akap5,Acvr1c,Stxbp5l,Rhbdd3,Adam9,RT1-Db1,Mcu,Ddrgk1,Phpt1,Copg1,Rab15,Trem2,Tiam1,Exoc1,Rab11fip2,Pde8b,Abca1,Sel1l,C1qtnf5,Cd2ap,Myrip,Myo18a,Idua,Pck2,Uqcc2,Svbp,Plek,Ufm1,Cbln4,Tcf7l2,Dph3,Mpc2,Enho |
| GO:0051047 | positive regulation of secretion | -4,87857 | -2,847 | 70/414 | Agt,Egfr,Jak2,Ppp3cb,Prkca,Sdc4,Stx1b,Htr2c,P2ry1,Cck,Arrb1,Rbp4,Syt1,Cadps,Gal,Acvr2a,Adora1,Oprk1,Prkce,Cacna1d,Mapk9,Ednrb,Chrnb2,Cacna1i,Cacnb4,Syt7,Syt9,Arf1,Isl1,Mfn2,Rab5a,Unc13a,Unc13b,Scamp5,Cyba,Stxbp5,Abat,Doc2b,Vamp8,Gab2,Dnm1l,Cacna1h,Nlgn1,Nlgn2,Rgcc,Ptpn23,Oga,Zfp384,Ap1g1,Cacna1b,Stxbp5l,Adam9,RT1-Db1,Mcu,Snx5,Phpt1,Rab15,Trem2,Exoc1,Slc4a8,Cd2ap,Myrip,Myo18a,Vps4b,Pck2,Ecrg4,Chmp2a,Dtnbp1,Tcf7l2,Mpc2 |
| GO:0071692 | protein localization to extracellular region | -4,74943 | -2,735 | 69/410 | Cacna1c,Egfr,Jak2,Ppp3cb,Myo5a,Arrb1,Cyp51,Rab12,Rbp4,Prkce,Cacna1d,Cacna1e,Pclo,Syt7,Ensa,Npff,Syt9,Arf1,Isl1,Unc13b,Stxbp5,Ccn3,Abat,Doc2b,Pafah1b1,Vamp8,Lyst,Madd,Stxbp3,Dnm1l,Rims2,Nlgn2,Park7,Ptpn23,Oga,Zfp384,Akap5,Acvr1c,Stxbp5l,Rhbdd3,Adam9,RT1-Db1,Mcu,Ddrgk1,Phpt1,Copg1,Rab15,Trem2,Tiam1,Exoc1,Rab11fip2,Pde8b,Abca1,Sel1l,C1qtnf5,Cd2ap,Myrip,Myo18a,Idua,Pck2,Uqcc2,Svbp,Plek,Ufm1,Cbln4,Tcf7l2,Dph3,Mpc2,Enho |
| GO:0090087 | regulation of peptide transport | -4,67056 | -2,672 | 100/656 | Egfr,Jak2,Ppp3cb,Htr2c,Chrm1,Akt2,Arrb1,Atp5if1,Cyp51,Oaz1,Pik3r1,Slc1a1,Cd74,Rbp4,Gal,Psen1,Adora1,Prkce,Ptgs2,Pacsin1,Cacna1d,Gsk3a,Cacna1e,Arc,Syt7,Ensa,Npff,Syt9,Arf1,Isl1,Unc13b,Snap91,Myo1c,Stxbp5,Ccn3,Abat,Doc2b,Vamp8,Kalrn,Ran,Synj1,Xpo1,Picalm,Madd,Dnm1l,Akap1,Pkia,Exoc4,Nlgn2,Park7,Ptpn23,Gli3,Oga,Ifi27,Prkcd,Anp32b,Ndel1,Zfp384,Akap5,Cabp1,Acvr1c,Uhmk1,Ppm1f,Stxbp5l,Fis1,Rhbdd3,Commd1,Adam9,B3gat3,Flna,RT1-Db1,Mcu,Ddrgk1,Phpt1,Ube2j1,Ube2j2,Trem2,Tiam1,Exoc1,Efnb2,Pde8b,Ctdspl2,Pip5k1c,Cd2ap,Fzd5,Myrip,Myo18a,Idua,Pck2,Uqcc2,Rangap1,Ecrg4,Ufm1,Dynlt2b,Ubl5,Tcf7l2,Dph3,Tomm7,Mpc2,Enho |
| GO:0051223 | regulation of protein transport | -4,63882 | -2,645 | 94/609 | Egfr,Jak2,Ppp3cb,Chrm1,Akt2,Arrb1,Atp5if1,Cyp51,Oaz1,Pik3r1,Slc1a1,Rbp4,Psen1,Prkce,Ptgs2,Pacsin1,Cacna1d,Gsk3a,Cacna1e,Arc,Syt7,Ensa,Npff,Syt9,Arf1,Isl1,Unc13b,Snap91,Myo1c,Stxbp5,Ccn3,Abat,Doc2b,Vamp8,Ran,Synj1,Xpo1,Picalm,Madd,Dnm1l,Akap1,Pkia,Exoc4,Nlgn2,Park7,Ptpn23,Gli3,Oga,Ifi27,Prkcd,Anp32b,Ndel1,Zfp384,Akap5,Cabp1,Acvr1c,Uhmk1,Ppm1f,Stxbp5l,Fis1,Rhbdd3,Commd1,Adam9,B3gat3,Flna,RT1-Db1,Mcu,Ddrgk1,Phpt1,Ube2j1,Ube2j2,Trem2,Tiam1,Exoc1,Efnb2,Pde8b,Ctdspl2,Pip5k1c,Cd2ap,Fzd5,Myrip,Myo18a,Idua,Pck2,Uqcc2,Rangap1,Ufm1,Dynlt2b,Ubl5,Tcf7l2,Dph3,Tomm7,Mpc2,Enho |
| GO:0070201 | regulation of establishment of protein localization | -4,557 | -2,589 | 97/636 | Egfr,Jak2,Ppp3cb,Chrm1,Akt2,Arrb1,Atp5if1,Cyp51,Oaz1,Pik3r1,Slc1a1,Rbp4,Psen1,Prkce,Cct4,Ptgs2,Pacsin1,Cacna1d,Gsk3a,Cacna1e,Arc,Syt7,Ensa,Npff,Syt9,Arf1,Isl1,Unc13b,Snap91,Myo1c,Stxbp5,Ccn3,Abat,Doc2b,Vamp8,Ran,Synj1,Xpo1,Picalm,Madd,Dnm1l,Akap1,Pkia,Exoc4,Nlgn2,Park7,Ptpn23,Gli3,Oga,Ifi27,Prkcd,Anp32b,Ndel1,Dkc1,Zfp384,Akap5,Cabp1,Acvr1c,Uhmk1,Ppm1f,Stxbp5l,Fis1,Rhbdd3,Commd1,Adam9,B3gat3,Flna,RT1-Db1,Mcu,Ddrgk1,Phpt1,Ube2j1,Ube2j2,Trem2,Tiam1,Exoc1,Efnb2,Pde8b,Ctdspl2,Pip5k1c,Cd2ap,Fzd5,Myrip,Myo18a,Idua,Pck2,Uqcc2,Rangap1,Ufm1,Dynlt2b,Ubl5,Tcf7l2,Nsd2,Dph3,Tomm7,Mpc2,Enho |
| GO:0050714 | positive regulation of protein secretion | -3,35847 | -1,621 | 31/163 | Egfr,Jak2,Ppp3cb,Arrb1,Rbp4,Prkce,Cacna1d,Arf1,Isl1,Unc13b,Abat,Doc2b,Vamp8,Dnm1l,Nlgn2,Ptpn23,Oga,Zfp384,Stxbp5l,Adam9,RT1-Db1,Mcu,Phpt1,Trem2,Exoc1,Cd2ap,Myrip,Myo18a,Pck2,Tcf7l2,Mpc2 |
| GO:0002791 | regulation of peptide secretion | -3,24071 | -1,535 | 56/355 | Egfr,Jak2,Ppp3cb,Htr2c,Arrb1,Cyp51,Cd74,Rbp4,Gal,Adora1,Prkce,Cacna1d,Cacna1e,Syt7,Ensa,Npff,Syt9,Arf1,Isl1,Unc13b,Stxbp5,Ccn3,Abat,Doc2b,Vamp8,Kalrn,Madd,Dnm1l,Nlgn2,Ptpn23,Oga,Zfp384,Acvr1c,Stxbp5l,Rhbdd3,Adam9,RT1-Db1,Mcu,Ddrgk1,Phpt1,Trem2,Tiam1,Exoc1,Pde8b,Cd2ap,Myrip,Myo18a,Idua,Pck2,Uqcc2,Ecrg4,Ufm1,Tcf7l2,Dph3,Mpc2,Enho |
| GO:0050708 | regulation of protein secretion | -3,17712 | -1,486 | 50/310 | Egfr,Jak2,Ppp3cb,Arrb1,Cyp51,Rbp4,Prkce,Cacna1d,Cacna1e,Syt7,Ensa,Npff,Syt9,Arf1,Isl1,Unc13b,Stxbp5,Ccn3,Abat,Doc2b,Vamp8,Madd,Dnm1l,Nlgn2,Ptpn23,Oga,Zfp384,Acvr1c,Stxbp5l,Rhbdd3,Adam9,RT1-Db1,Mcu,Ddrgk1,Phpt1,Trem2,Tiam1,Exoc1,Pde8b,Cd2ap,Myrip,Myo18a,Idua,Pck2,Uqcc2,Ufm1,Tcf7l2,Dph3,Mpc2,Enho |
| GO:0002793 | positive regulation of peptide secretion | -2,91937 | -1,293 | 34/195 | Egfr,Jak2,Ppp3cb,Arrb1,Rbp4,Gal,Adora1,Prkce,Cacna1d,Arf1,Isl1,Unc13b,Abat,Doc2b,Vamp8,Dnm1l,Nlgn2,Ptpn23,Oga,Zfp384,Stxbp5l,Adam9,RT1-Db1,Mcu,Phpt1,Trem2,Exoc1,Cd2ap,Myrip,Myo18a,Pck2,Ecrg4,Tcf7l2,Mpc2 |
| GO:0030073 | insulin secretion | -2,6695 | -1,107 | 39/240 | Cacna1c,Jak2,Ppp3cb,Myo5a,Arrb1,Rbp4,Prkce,Cacna1d,Cacna1e,Pclo,Syt7,Ensa,Npff,Syt9,Isl1,Ccn3,Abat,Doc2b,Stxbp3,Rims2,Nlgn2,Park7,Oga,Acvr1c,Stxbp5l,RT1-Db1,Mcu,Ddrgk1,Phpt1,Tiam1,Rab11fip2,Pde8b,Myrip,Pck2,Uqcc2,Ufm1,Tcf7l2,Mpc2,Enho |
| GO:0030072 | peptide hormone secretion | -2,49448 | -0,980 | 46/301 | Cacna1c,Egfr,Jak2,Ppp3cb,Myo5a,Htr2c,Arrb1,Rbp4,Gal,Prkce,Cacna1d,Cacna1e,Pclo,Syt7,Ensa,Npff,Syt9,Isl1,Ccn3,Abat,Doc2b,Kalrn,Madd,Stxbp3,Rims2,Nlgn2,Park7,Oga,Acvr1c,Slc16a2,Stxbp5l,RT1-Db1,Mcu,Ddrgk1,Phpt1,Tiam1,Rab11fip2,Pde8b,Myrip,Pck2,Uqcc2,Ecrg4,Ufm1,Tcf7l2,Mpc2,Enho |
| GO:0050796 | regulation of insulin secretion | -2,16095 | -0,754 | 32/201 | Jak2,Ppp3cb,Arrb1,Rbp4,Prkce,Cacna1d,Cacna1e,Syt7,Ensa,Npff,Syt9,Isl1,Ccn3,Abat,Doc2b,Nlgn2,Oga,Acvr1c,Stxbp5l,RT1-Db1,Mcu,Ddrgk1,Phpt1,Tiam1,Pde8b,Myrip,Pck2,Uqcc2,Ufm1,Tcf7l2,Mpc2,Enho |
| GO:0032024 | positive regulation of insulin secretion | -2,12497 | -0,725 | 18/96 | Jak2,Ppp3cb,Arrb1,Rbp4,Prkce,Cacna1d,Isl1,Abat,Doc2b,Nlgn2,Oga,RT1-Db1,Mcu,Phpt1,Myrip,Pck2,Tcf7l2,Mpc2 |
| GO:0034765 | regulation of ion transmembrane transport | -7,29937 | -4,939 | 103/598 | Agt,C3,Cacna1c,Camk2d,Ace,Grin2a,Grm5,Gstm2,Pde4b,Prkca,Yes1,Dmd,Myo5a,Dbi,Akt2,Dlg1,Scn2b,Atp1b3,Kcna5,Oaz1,Agrn,Rasa1,Gal,Cd63,Psen1,Dpp6,Oprk1,Prkce,Kcnq3,Scn8a,Cacna1d,Kcnj10,Kcnab2,Ctss,Gsk3a,Cacna1e,Arc,Cacna1i,Cacnb4,Fgf14,Arf1,Kcna6,Mfn2,Akap6,Kcnip3,Kcns2,Shank1,Cyba,Scn1a,Tmsb4x,Ppargc1a,Kcnma1,Slc25a27,Cox17,Acsl5,Kcnh4,Stxbp3,Hcn3,Cacna1h,Nlgn1,Bin1,Nlgn2,Kcnc3,Park7,Cacng3,Cacng4,Fabp5,Oga,Prkcd,Kcnq2,Cabp1,Shank2,Nlgn3,Ahnak,Lmbrd1,Cacna1b,Kcnip4,Antkmt,Iscu,Commd1,Slmap,Hecw1,Flna,Clic2,Amigo1,Kcnf1,Coa8,Cacna2d2,Trem2,Neto1,Neto2,Rtn2,Mef2a,Abl1,Osbpl8,Ubash3b,Hecw2,Dpp10,Catsper2,Repin1,Scn3a,Ryr2,Shisa7,Htr2c,Cox6a1,Slc1a1,Abcc9,Cox4i1,Slc6a6,Cox7a2,Htr2a,Atp2b4,Atp5mc1,Atp6v0a1,Cox6c,Rnase4,Kcnt1,Kcnmb4,Slc6a11,Grm7,Atox1,Clcn3,Slc4a4,Slc24a2,Atp6v0e1,Slc6a20,Atp5mc3,Atp6v0a2,Slc8a2,Slc8a3,Atp5me,Opa1,Cox8a,Atp5po,Atp5f1e,Atp5f1d,Cox5a,Slc30a10,Slc39a12,Mcu,Slc4a10,Atp6v0a4,Atp6v0b,Atp5mg,Slc25a23,Kcnt2,Ank1,Slc9a8,Slc4a8,Panx1,Slc39a1,Ccl19,Ndufs7,Micu3,Zdhhc17,Atp5pd,Cox7c,Taf3,Adora1,Atf4,Gnaq,Akap5,Nsf,Tsc1,Vps4b |
| GO:0034765 | regulation of ion transmembrane transport | -7,29937 | -4,939 | 103/598 | Agt,C3,Cacna1c,Camk2d,Ace,Grin2a,Grm5,Gstm2,Pde4b,Prkca,Yes1,Dmd,Myo5a,Dbi,Akt2,Dlg1,Scn2b,Atp1b3,Kcna5,Oaz1,Agrn,Rasa1,Gal,Cd63,Psen1,Dpp6,Oprk1,Prkce,Kcnq3,Scn8a,Cacna1d,Kcnj10,Kcnab2,Ctss,Gsk3a,Cacna1e,Arc,Cacna1i,Cacnb4,Fgf14,Arf1,Kcna6,Mfn2,Akap6,Kcnip3,Kcns2,Shank1,Cyba,Scn1a,Tmsb4x,Ppargc1a,Kcnma1,Slc25a27,Cox17,Acsl5,Kcnh4,Stxbp3,Hcn3,Cacna1h,Nlgn1,Bin1,Nlgn2,Kcnc3,Park7,Cacng3,Cacng4,Fabp5,Oga,Prkcd,Kcnq2,Cabp1,Shank2,Nlgn3,Ahnak,Lmbrd1,Cacna1b,Kcnip4,Antkmt,Iscu,Commd1,Slmap,Hecw1,Flna,Clic2,Amigo1,Kcnf1,Coa8,Cacna2d2,Trem2,Neto1,Neto2,Rtn2,Mef2a,Abl1,Osbpl8,Ubash3b,Hecw2,Dpp10,Catsper2,Repin1,Scn3a,Ryr2,Shisa7 |
| GO:0098662 | inorganic cation transmembrane transport | -7,24388 | -4,894 | 127/784 | Cacna1c,Camk2d,Grin2a,Gstm2,Pde4b,Dmd,Myo5a,Dbi,Htr2c,Dlg1,Cox6a1,Scn2b,Atp1b3,Kcna5,Slc1a1,Abcc9,Agrn,Gal,Cd63,Psen1,Dpp6,Oprk1,Prkce,Cox4i1,Slc6a6,Cox7a2,Htr2a,Atp2b4,Kcnq3,Scn8a,Cacna1d,Kcnj10,Kcnab2,Atp5mc1,Atp6v0a1,Cacna1e,Cox6c,Rnase4,Cacna1i,Cacnb4,Kcnt1,Fgf14,Arf1,Kcna6,Akap6,Kcnip3,Kcnmb4,Kcns2,Cyba,Slc6a11,Scn1a,Grm7,Tmsb4x,Kcnma1,Atox1,Clcn3,Slc4a4,Slc24a2,Cox17,Atp6v0e1,Slc6a20,Kcnh4,Hcn3,Atp5mc3,Cacna1h,Atp6v0a2,Bin1,Kcnc3,Park7,Slc8a2,Slc8a3,Atp5me,Cacng3,Cacng4,Kcnq2,Cabp1,Opa1,Cox8a,Ahnak,Atp5po,Atp5f1e,Atp5f1d,Cox5a,Cacna1b,Kcnip4,Antkmt,Iscu,Slc30a10,Commd1,Slmap,Hecw1,Slc39a12,Flna,Clic2,Mcu,Amigo1,Slc4a10,Atp6v0a4,Atp6v0b,Kcnf1,Coa8,Atp5mg,Cacna2d2,Slc25a23,Trem2,Kcnt2,Ank1,Neto1,Neto2,Slc9a8,Abl1,Slc4a8,Panx1,Ubash3b,Hecw2,Slc39a1,Ccl19,Ndufs7,Dpp10,Micu3,Catsper2,Zdhhc17,Scn3a,Atp5pd,Ryr2,Cox7c,Taf3 |
| GO:0034762 | regulation of transmembrane transport | -7,15002 | -4,805 | 103/602 | Agt,C3,Cacna1c,Camk2d,Ace,Grin2a,Grm5,Gstm2,Pde4b,Prkca,Yes1,Dmd,Myo5a,Dbi,Akt2,Dlg1,Scn2b,Atp1b3,Kcna5,Oaz1,Agrn,Rasa1,Gal,Cd63,Psen1,Dpp6,Oprk1,Prkce,Kcnq3,Scn8a,Cacna1d,Kcnj10,Kcnab2,Ctss,Gsk3a,Cacna1e,Arc,Cacna1i,Cacnb4,Fgf14,Arf1,Kcna6,Mfn2,Akap6,Kcnip3,Kcns2,Shank1,Cyba,Scn1a,Tmsb4x,Ppargc1a,Kcnma1,Slc25a27,Cox17,Acsl5,Kcnh4,Stxbp3,Hcn3,Cacna1h,Nlgn1,Bin1,Nlgn2,Kcnc3,Park7,Cacng3,Cacng4,Fabp5,Oga,Prkcd,Kcnq2,Cabp1,Shank2,Nlgn3,Ahnak,Lmbrd1,Cacna1b,Kcnip4,Antkmt,Iscu,Commd1,Slmap,Hecw1,Flna,Clic2,Amigo1,Kcnf1,Coa8,Cacna2d2,Trem2,Neto1,Neto2,Rtn2,Mef2a,Abl1,Osbpl8,Ubash3b,Hecw2,Dpp10,Catsper2,Repin1,Scn3a,Ryr2,Shisa7 |
| GO:0043266 | regulation of potassium ion transport | -4,1526 | -2,250 | 26/115 | Dlg1,Atp1b3,Kcna5,Agrn,Gal,Cd63,Dpp6,Adora1,Oprk1,Htr2a,Cacna1d,Kcnab2,Akap6,Kcnip3,Kcns2,Atf4,Gnaq,Bin1,Akap5,Kcnip4,Flna,Amigo1,Trem2,Neto1,Neto2,Dpp10 |
| GO:1904062 | regulation of cation transmembrane transport | -4,03423 | -2,156 | 62/377 | Agt,Cacna1c,Camk2d,Grin2a,Gstm2,Pde4b,Dmd,Myo5a,Dbi,Dlg1,Scn2b,Atp1b3,Agrn,Gal,Cd63,Dpp6,Oprk1,Prkce,Cacna1d,Kcnab2,Ctss,Arc,Cacnb4,Fgf14,Arf1,Akap6,Kcnip3,Kcns2,Shank1,Cyba,Tmsb4x,Ppargc1a,Cox17,Nlgn1,Bin1,Nlgn2,Park7,Cacng3,Cacng4,Cabp1,Shank2,Nlgn3,Ahnak,Kcnip4,Antkmt,Iscu,Commd1,Slmap,Hecw1,Flna,Clic2,Amigo1,Coa8,Trem2,Neto1,Neto2,Abl1,Ubash3b,Hecw2,Dpp10,Ryr2,Shisa7 |
| GO:0006813 | potassium ion transport | -3,85326 | -2,009 | 44/246 | Dlg1,Atp1b3,Kcna5,Abcc9,Agrn,Gal,Cd63,Dpp6,Adora1,Oprk1,Htr2a,Kcnq3,Cacna1d,Kcnj10,Kcnab2,Nsf,Kcnt1,Tsc1,Kcna6,Akap6,Kcnip3,Kcnmb4,Kcns2,Atf4,Gnaq,Kcnma1,Slc24a2,Kcnh4,Hcn3,Bin1,Kcnc3,Kcnq2,Akap5,Kcnip4,Flna,Amigo1,Kcnf1,Trem2,Kcnt2,Neto1,Neto2,Slc9a8,Vps4b,Dpp10 |
| GO:1901379 | regulation of potassium ion transmembrane transport | -3,10325 | -1,425 | 20/92 | Dlg1,Atp1b3,Agrn,Gal,Cd63,Dpp6,Oprk1,Cacna1d,Kcnab2,Akap6,Kcnip3,Kcns2,Bin1,Kcnip4,Flna,Amigo1,Trem2,Neto1,Neto2,Dpp10 |
| GO:0032412 | regulation of ion transmembrane transporter activity | -3,08585 | -1,411 | 45/274 | Camk2d,Grin2a,Grm5,Gstm2,Pde4b,Prkca,Dmd,Myo5a,Dlg1,Scn2b,Atp1b3,Agrn,Gal,Cacna1d,Ctss,Arc,Cacnb4,Fgf14,Akap6,Kcns2,Shank1,Tmsb4x,Ppargc1a,Cox17,Nlgn1,Nlgn2,Park7,Cacng3,Cacng4,Cabp1,Shank2,Nlgn3,Ahnak,Antkmt,Slmap,Hecw1,Clic2,Amigo1,Coa8,Trem2,Neto1,Neto2,Hecw2,Ryr2,Shisa7 |
| GO:2001257 | regulation of cation channel activity | -3,02385 | -1,365 | 33/185 | Grin2a,Gstm2,Pde4b,Dmd,Myo5a,Dlg1,Gal,Ctss,Arc,Cacnb4,Fgf14,Akap6,Kcns2,Shank1,Tmsb4x,Ppargc1a,Nlgn1,Nlgn2,Park7,Cacng3,Cacng4,Cabp1,Shank2,Nlgn3,Ahnak,Antkmt,Slmap,Clic2,Amigo1,Trem2,Neto1,Neto2,Shisa7 |
| GO:0071805 | potassium ion transmembrane transport | -2,73528 | -1,158 | 36/215 | Dlg1,Atp1b3,Kcna5,Abcc9,Agrn,Gal,Cd63,Dpp6,Oprk1,Kcnq3,Cacna1d,Kcnj10,Kcnab2,Kcnt1,Kcna6,Akap6,Kcnip3,Kcnmb4,Kcns2,Kcnma1,Slc24a2,Kcnh4,Hcn3,Bin1,Kcnc3,Kcnq2,Kcnip4,Flna,Amigo1,Kcnf1,Trem2,Kcnt2,Neto1,Neto2,Slc9a8,Dpp10 |
| GO:0022898 | regulation of transmembrane transporter activity | -2,73101 | -1,156 | 45/285 | Camk2d,Grin2a,Grm5,Gstm2,Pde4b,Prkca,Dmd,Myo5a,Dlg1,Scn2b,Atp1b3,Agrn,Gal,Cacna1d,Ctss,Arc,Cacnb4,Fgf14,Akap6,Kcns2,Shank1,Tmsb4x,Ppargc1a,Cox17,Nlgn1,Nlgn2,Park7,Cacng3,Cacng4,Cabp1,Shank2,Nlgn3,Ahnak,Antkmt,Slmap,Hecw1,Clic2,Amigo1,Coa8,Trem2,Neto1,Neto2,Hecw2,Ryr2,Shisa7 |
| GO:0032409 | regulation of transporter activity | -2,63831 | -1,085 | 46/296 | Camk2d,Grin2a,Grm5,Gstm2,Pde4b,Prkca,Dmd,Myo5a,Dlg1,Scn2b,Atp1b3,Agrn,Gal,Cacna1d,Ctss,Arc,Cacnb4,Fgf14,Akap6,Kcns2,Shank1,Tmsb4x,Ppargc1a,Cox17,Nlgn1,Nlgn2,Park7,Cacng3,Cacng4,Prkcd,Cabp1,Shank2,Nlgn3,Ahnak,Antkmt,Slmap,Hecw1,Clic2,Amigo1,Coa8,Trem2,Neto1,Neto2,Hecw2,Ryr2,Shisa7 |
| GO:0035249 | synaptic transmission, glutamatergic | -6,84855 | -4,521 | 31/112 | Egfr,Grm5,Htr1b,Tnr,Syt1,Psen1,Adora1,Ptgs2,Htr2a,Gria1,Grik2,Cacnb4,Unc13a,Unc13b,Shank1,Grid2,Grm7,Kalrn,Clcn3,Nlgn1,Nlgn2,Cacng3,Cacng4,Shank2,Nlgn3,Clstn3,Unc13c,Grik3,Napb,Dtnbp1,Iqsec2,Grin2a,Fnta,Agrn,Arc,Ppargc1a,Dlgap2,Park7,Nptx2,Neto1,Neto2,Shisa7,Jak2,P2ry1,Nog,Cblb,Zfyve28,Neurl1,Bicd1,Tafa4 |
| GO:0035249 | synaptic transmission, glutamatergic | -6,84855 | -4,521 | 31/112 | Egfr,Grm5,Htr1b,Tnr,Syt1,Psen1,Adora1,Ptgs2,Htr2a,Gria1,Grik2,Cacnb4,Unc13a,Unc13b,Shank1,Grid2,Grm7,Kalrn,Clcn3,Nlgn1,Nlgn2,Cacng3,Cacng4,Shank2,Nlgn3,Clstn3,Unc13c,Grik3,Napb,Dtnbp1,Iqsec2 |
| GO:0051966 | regulation of synaptic transmission, glutamatergic | -4,99974 | -2,955 | 22/80 | Egfr,Grm5,Htr1b,Tnr,Syt1,Psen1,Adora1,Ptgs2,Htr2a,Grik2,Shank1,Grm7,Kalrn,Nlgn1,Nlgn2,Cacng3,Cacng4,Shank2,Nlgn3,Grik3,Dtnbp1,Iqsec2 |
| GO:0051968 | positive regulation of synaptic transmission, glutamatergic | -3,4785 | -1,720 | 12/39 | Egfr,Tnr,Ptgs2,Shank1,Nlgn1,Nlgn2,Cacng3,Cacng4,Shank2,Nlgn3,Dtnbp1,Iqsec2 |
| GO:0099601 | regulation of neurotransmitter receptor activity | -3,24243 | -1,536 | 18/77 | Grin2a,Fnta,Agrn,Arc,Shank1,Ppargc1a,Nlgn1,Dlgap2,Nlgn2,Park7,Cacng3,Cacng4,Shank2,Nlgn3,Nptx2,Neto1,Neto2,Shisa7 |
| GO:2000311 | regulation of AMPA receptor activity | -3,01527 | -1,361 | 9/27 | Arc,Shank1,Nlgn1,Nlgn2,Cacng3,Cacng4,Shank2,Nlgn3,Shisa7 |
| GO:0010469 | regulation of signaling receptor activity | -2,12207 | -0,725 | 28/171 | Grin2a,Jak2,P2ry1,Fnta,Nog,Agrn,Psen1,Adora1,Arc,Shank1,Ppargc1a,Nlgn1,Dlgap2,Nlgn2,Park7,Cacng3,Cacng4,Shank2,Cblb,Nlgn3,Nptx2,Zfyve28,Neto1,Neto2,Neurl1,Bicd1,Tafa4,Shisa7 |
| R-RNO-5389840 | Mitochondrial translation elongation | -6,84622 | -4,521 | 24/74 | Mrpl23,Mrps34,Mrpl27,Mrps14,Mrps18c,Mrpl34,Mrps12,mrpl11,Mrpl16,Mrps18b,Mrpl41,Mrps33,Mrpl19,Mrps25,Mrps35,Mrpl54,Mrpl13,Mrps18a,Mrpl14,Mrpl12,Mrps23,Chchd1,Mrpl52,Mrps21,Hsd17b10,Rmnd1,Foxo3,Trmt10b,Aars2,Twnk,Rcc1l,Uqcc2,Lars2,Coa3,Mtres1,Slc25a33 |
| R-RNO-5389840 | Mitochondrial translation elongation | -6,84622 | -4,521 | 24/74 | Mrpl23,Mrps34,Mrpl27,Mrps14,Mrps18c,Mrpl34,Mrps12,mrpl11,Mrpl16,Mrps18b,Mrpl41,Mrps33,Mrpl19,Mrps25,Mrps35,Mrpl54,Mrpl13,Mrps18a,Mrpl14,Mrpl12,Mrps23,Chchd1,Mrpl52,Mrps21 |
| R-RNO-5419276 | Mitochondrial translation termination | -6,72236 | -4,430 | 24/75 | Mrpl23,Mrps34,Mrpl27,Mrps14,Mrps18c,Mrpl34,Mrps12,mrpl11,Mrpl16,Mrps18b,Mrpl41,Mrps33,Mrpl19,Mrps25,Mrps35,Mrpl54,Mrpl13,Mrps18a,Mrpl14,Mrpl12,Mrps23,Chchd1,Mrpl52,Mrps21 |
| R-RNO-5368287 | Mitochondrial translation | -6,60118 | -4,326 | 24/76 | Mrpl23,Mrps34,Mrpl27,Mrps14,Mrps18c,Mrpl34,Mrps12,mrpl11,Mrpl16,Mrps18b,Mrpl41,Mrps33,Mrpl19,Mrps25,Mrps35,Mrpl54,Mrpl13,Mrps18a,Mrpl14,Mrpl12,Mrps23,Chchd1,Mrpl52,Mrps21 |
| GO:0140053 | mitochondrial gene expression | -3,39696 | -1,655 | 21/94 | Hsd17b10,Mrpl23,Mrps34,Mrps14,Rmnd1,Mrpl16,Mrps18b,Foxo3,Mrps35,Trmt10b,Aars2,Mrpl12,Twnk,Rcc1l,Chchd1,Mrpl52,Uqcc2,Lars2,Coa3,Mtres1,Slc25a33 |
| GO:0032543 | mitochondrial translation | -2,42949 | -0,932 | 14/63 | Mrpl23,Mrps34,Mrps14,Rmnd1,Mrpl16,Mrps18b,Mrps35,Aars2,Rcc1l,Chchd1,Mrpl52,Uqcc2,Lars2,Coa3 |
| R-RNO-5653656 | Vesicle-mediated transport | -6,82342 | -4,503 | 88/498 | Hbb,Chm,Akt2,Cpd,Arrb1,Nedd8,Rab12,Hba-a1,Syt1,Akt3,Vamp3,Bet1,Pacsin1,Ubc,Gria1,Sec23a,Nsf,Tsc1,Syt9,Syt11,Amph,Arl1,Arf1,Ap2s1,Snap91,Arf5,Sh3gl1,Pafah1b1,Vamp8,Gjb6,Dctn4,Synj1,Kif3c,Picalm,Gosr1,Madd,Cltb,Sh3gl2,Bin1,Kif1b,Ap2b1,Dnm3,Dnase2,Golgb1,Fnbp1,Stx17,Dync1li1,Tbc1d24,Trappc1,Bloc1s1,Kif21b,Actr2,Use1,Cope,Tubb2b,Trappc2l,Tbc1d17,Snx5,Copg1,Arpc4,Tmed3,Ap4m1,Ank1,Ap2a1,Fnbp1l,Stam2,Pacsin3,Dnajc6,Sptb,Ston2,Trip11,Pip5k1c,Hba-a2,Vps4b,Cog3,RGD1307443,Kdelr1,Mvb12b,Tbc1d20,Stx16,Bicd1,Yipf6,Chmp2a,Trappc4,Tuba1b,Dtnbp1,Sbf2,Rps27a,Asgr1,Fuca1,Mgat3,Mgat5,Renbp,Manea,Fuom,Man1a1,Gfpt1,Pmm1,Dpagt1,Mpdu1,Glb1,Neu4,St3gal1,St6galnac6,Myo5a,Rab2a,Exoc4,Myo1b,Ap1g1,Phaf1,Klhl12,Commd1,Arfgef2,Tex261,Rer1,Exoc1,Sptbn1,Sorcs1,Rbsn,Pgap1,Ccdc22,Myo18a,Dnajc28,Macf1,Stxbp6,Nrbp2,Trappc6a |
| R-RNO-5653656 | Vesicle-mediated transport | -6,82342 | -4,503 | 88/498 | Hbb,Chm,Akt2,Cpd,Arrb1,Nedd8,Rab12,Hba-a1,Syt1,Akt3,Vamp3,Bet1,Pacsin1,Ubc,Gria1,Sec23a,Nsf,Tsc1,Syt9,Syt11,Amph,Arl1,Arf1,Ap2s1,Snap91,Arf5,Sh3gl1,Pafah1b1,Vamp8,Gjb6,Dctn4,Synj1,Kif3c,Picalm,Gosr1,Madd,Cltb,Sh3gl2,Bin1,Kif1b,Ap2b1,Dnm3,Dnase2,Golgb1,Fnbp1,Stx17,Dync1li1,Tbc1d24,Trappc1,Bloc1s1,Kif21b,Actr2,Use1,Cope,Tubb2b,Trappc2l,Tbc1d17,Snx5,Copg1,Arpc4,Tmed3,Ap4m1,Ank1,Ap2a1,Fnbp1l,Stam2,Pacsin3,Dnajc6,Sptb,Ston2,Trip11,Pip5k1c,Hba-a2,Vps4b,Cog3,RGD1307443,Kdelr1,Mvb12b,Tbc1d20,Stx16,Bicd1,Yipf6,Chmp2a,Trappc4,Tuba1b,Dtnbp1,Sbf2,Rps27a |
| R-RNO-199991 | Membrane Trafficking | -6,77401 | -4,458 | 85/477 | Chm,Akt2,Cpd,Arrb1,Nedd8,Rab12,Syt1,Akt3,Vamp3,Bet1,Pacsin1,Ubc,Gria1,Sec23a,Nsf,Tsc1,Syt9,Syt11,Amph,Arl1,Arf1,Ap2s1,Snap91,Arf5,Sh3gl1,Pafah1b1,Vamp8,Gjb6,Dctn4,Synj1,Kif3c,Picalm,Gosr1,Madd,Cltb,Sh3gl2,Bin1,Kif1b,Ap2b1,Dnm3,Dnase2,Golgb1,Fnbp1,Stx17,Dync1li1,Tbc1d24,Trappc1,Bloc1s1,Kif21b,Actr2,Use1,Cope,Tubb2b,Trappc2l,Tbc1d17,Snx5,Copg1,Arpc4,Tmed3,Ap4m1,Ank1,Ap2a1,Fnbp1l,Stam2,Pacsin3,Dnajc6,Sptb,Ston2,Trip11,Pip5k1c,Vps4b,Cog3,RGD1307443,Kdelr1,Mvb12b,Tbc1d20,Stx16,Bicd1,Yipf6,Chmp2a,Trappc4,Tuba1b,Dtnbp1,Sbf2,Rps27a |
| R-RNO-446203 | Asparagine N-linked glycosylation | -4,31923 | -2,388 | 42/221 | Asgr1,Fuca1,Mgat3,Bet1,Ubc,Gria1,Sec23a,Nsf,Arf1,Mgat5,Arf5,Renbp,Dctn4,Gosr1,Manea,Golgb1,Stx17,Dync1li1,Trappc1,Cope,Tubb2b,Trappc2l,Fuom,Man1a1,Gfpt1,Copg1,Pmm1,Dpagt1,Tmed3,Mpdu1,Ank1,Sptb,Glb1,Neu4,Cog3,Kdelr1,Tbc1d20,St3gal1,Trappc4,St6galnac6,Tuba1b,Rps27a |
| R-RNO-948021 | Transport to the Golgi and subsequent modification | -3,35193 | -1,615 | 29/149 | Fuca1,Mgat3,Bet1,Gria1,Sec23a,Nsf,Arf1,Mgat5,Arf5,Dctn4,Gosr1,Manea,Golgb1,Stx17,Dync1li1,Trappc1,Cope,Tubb2b,Trappc2l,Man1a1,Copg1,Tmed3,Ank1,Sptb,Cog3,Kdelr1,Tbc1d20,Trappc4,Tuba1b |
| GO:0048193 | Golgi vesicle transport | -3,25909 | -1,549 | 45/269 | Myo5a,Vamp3,Bet1,Sec23a,Nsf,Arl1,Arf1,Rab2a,Arf5,Gosr1,Exoc4,Myo1b,Ap1g1,Phaf1,Stx17,Klhl12,Trappc1,Commd1,Use1,Cope,Trappc2l,Arfgef2,Tex261,Copg1,Rer1,Tmed3,Ap4m1,Exoc1,Sptbn1,Ank1,Sorcs1,Rbsn,Trip11,Pgap1,Ccdc22,Myo18a,Dnajc28,Cog3,Kdelr1,Tbc1d20,Macf1,Stxbp6,Trappc4,Nrbp2,Trappc6a |
| R-RNO-199977 | ER to Golgi Anterograde Transport | -2,74398 | -1,160 | 24/126 | Bet1,Gria1,Sec23a,Nsf,Arf1,Arf5,Dctn4,Gosr1,Golgb1,Stx17,Dync1li1,Trappc1,Cope,Tubb2b,Trappc2l,Copg1,Tmed3,Ank1,Sptb,Cog3,Kdelr1,Tbc1d20,Trappc4,Tuba1b |
| GO:0006888 | endoplasmic reticulum to Golgi vesicle-mediated transport | -2,64157 | -1,087 | 23/121 | Bet1,Sec23a,Arf1,Rab2a,Gosr1,Stx17,Klhl12,Trappc1,Use1,Cope,Trappc2l,Tex261,Copg1,Tmed3,Ank1,Trip11,Pgap1,Cog3,Kdelr1,Tbc1d20,Trappc4,Nrbp2,Trappc6a |
| R-RNO-6807878 | COPI-mediated anterograde transport | -2,43533 | -0,934 | 17/83 | Bet1,Nsf,Arf1,Arf5,Dctn4,Gosr1,Golgb1,Dync1li1,Cope,Tubb2b,Copg1,Tmed3,Ank1,Sptb,Cog3,Kdelr1,Tuba1b |
| GO:0007626 | locomotory behaviour | -6,72883 | -4,430 | 54/258 | Cacna1c,Egr1,Grm5,Ppp3cb,Sod1,Myo5a,Atxn1,Htr2c,Cstb,Cntn2,Glrb,Slc1a1,Tnr,Strn,Penk,Oprk1,Prkce,Npy1r,Selenop,Pde1b,Scn8a,Kcnj10,Gnao1,Cacna1e,Chrnb2,Cacnb4,Tsc1,Fgf14,Scn1a,Abat,Pafah1b1,Apba1,Apba2,Ncoa2,Kcnma1,Calb1,Kalrn,Clcn3,Nlgn2,Park7,Adgrl3,Cacna1b,Slc4a10,Pak6,Myg1,Hexa,Sez6l,Inpp5f,Vps13a,Arrdc3,Pbx3,Idua,Ciart,Fxn,Htr2a,Gabrg2,Shank1,Grm7,Shank2,Nlgn3,Zfhx2,Pcdh17,Unc79,Tpgs1 |
| GO:0007626 | locomotory behaviour | -6,72883 | -4,430 | 54/258 | Cacna1c,Egr1,Grm5,Ppp3cb,Sod1,Myo5a,Atxn1,Htr2c,Cstb,Cntn2,Glrb,Slc1a1,Tnr,Strn,Penk,Oprk1,Prkce,Npy1r,Selenop,Pde1b,Scn8a,Kcnj10,Gnao1,Cacna1e,Chrnb2,Cacnb4,Tsc1,Fgf14,Scn1a,Abat,Pafah1b1,Apba1,Apba2,Ncoa2,Kcnma1,Calb1,Kalrn,Clcn3,Nlgn2,Park7,Adgrl3,Cacna1b,Slc4a10,Pak6,Myg1,Hexa,Sez6l,Inpp5f,Vps13a,Arrdc3,Pbx3,Idua,Ciart,Fxn |
| GO:0030534 | adult behaviour | -4,62723 | -2,638 | 38/187 | Cacna1c,Atxn1,Htr2c,Cstb,Cntn2,Glrb,Slc1a1,Oprk1,Htr2a,Gabrg2,Scn8a,Kcnj10,Chrnb2,Cacnb4,Tsc1,Fgf14,Shank1,Scn1a,Abat,Grm7,Pafah1b1,Kcnma1,Kalrn,Clcn3,Nlgn2,Park7,Shank2,Nlgn3,Hexa,Sez6l,Zfhx2,Pcdh17,Inpp5f,Pbx3,Unc79,Idua,Fxn,Tpgs1 |
| GO:0008344 | adult locomotory behaviour | -3,34443 | -1,609 | 23/108 | Cacna1c,Atxn1,Cstb,Cntn2,Glrb,Scn8a,Kcnj10,Cacnb4,Tsc1,Fgf14,Scn1a,Pafah1b1,Kcnma1,Kalrn,Clcn3,Nlgn2,Park7,Hexa,Sez6l,Inpp5f,Pbx3,Idua,Fxn |
| GO:0007628 | adult walking behaviour | -2,26901 | -0,826 | 11/46 | Cacna1c,Cntn2,Glrb,Scn8a,Kcnj10,Cacnb4,Scn1a,Kcnma1,Hexa,Idua,Fxn |
| GO:0090659 | walking behaviour | -2,12186 | -0,725 | 11/48 | Cacna1c,Cntn2,Glrb,Scn8a,Kcnj10,Cacnb4,Scn1a,Kcnma1,Hexa,Idua,Fxn |
| R-RNO-399719 | Trafficking of AMPA receptors | -6,72468 | -4,430 | 13/25 | Camk2d,Prkcg,Dlg1,Camk2a,Gria1,Nsf,Ap2s1,Grip1,Ap2b1,Cacng3,Cacng4,Camk2g,Ap2a1,Cltb,Fzd5,Prkca,Atp1b3,Gnaq,Calb1,Dnm3,Prkacb,Soat1,Npc2 |
| R-RNO-399719 | Trafficking of AMPA receptors | -6,72468 | -4,430 | 13/25 | Camk2d,Prkcg,Dlg1,Camk2a,Gria1,Nsf,Ap2s1,Grip1,Ap2b1,Cacng3,Cacng4,Camk2g,Ap2a1 |
| R-RNO-399721 | Glutamate binding, activation of AMPA receptors and synaptic plasticity | -6,72468 | -4,430 | 13/25 | Camk2d,Prkcg,Dlg1,Camk2a,Gria1,Nsf,Ap2s1,Grip1,Ap2b1,Cacng3,Cacng4,Camk2g,Ap2a1 |
| R-RNO-416993 | Trafficking of GluR2-containing AMPA receptors | -3,70248 | -1,894 | 7/14 | Prkcg,Gria1,Nsf,Ap2s1,Grip1,Ap2b1,Ap2a1 |
| R-RNO-5140745 | WNT5A-dependent internalization of FZD2, FZD5 and ROR2 | -2,75764 | -1,172 | 5/10 | Ap2s1,Cltb,Ap2b1,Ap2a1,Fzd5 |
| R-RNO-5099900 | WNT5A-dependent internalization of FZD4 | -2,33573 | -0,871 | 5/12 | Prkcg,Ap2s1,Cltb,Ap2b1,Ap2a1 |
| ko04961 | Endocrine and other factor-regulated calcium reabsorption | -2,05222 | -0,688 | 11/49 | Prkca,Prkcg,Atp1b3,Ap2s1,Gnaq,Calb1,Cltb,Ap2b1,Dnm3,Prkacb,Ap2a1 |
| rno04961 | Endocrine and other factor-regulated calcium reabsorption | -2,05222 | -0,688 | 11/49 | Prkca,Prkcg,Atp1b3,Ap2s1,Gnaq,Calb1,Cltb,Ap2b1,Dnm3,Prkacb,Ap2a1 |
| R-RNO-8964038 | LDL clearance | -2,00795 | -0,660 | 5/14 | Ap2s1,Soat1,Ap2b1,Npc2,Ap2a1 |
| GO:0009144 | purine nucleoside triphosphate metabolic process | -6,50595 | -4,235 | 27/93 | Ak3,Atp5mc1,Tmsb4x,Ppargc1a,Nme2,Ran,Nme3,Atp5mc3,Atp5me,Oga,Opa1,Nme1,Atp5po,Atp5f1e,Atp5f1d,Antkmt,Ndufc2,Nudt2,Gtpbp1,Atp5mg,Trem2,Guk1,Dmac2l,Nudt16,Atp5pd,Uqcc3,Taf3,Tyms,Dtymk,Pde4b,Dbi,Acadsb,Pde4a,Psen1,Htr2a,Gpam,Cacnb4,Acaca,Pde10a,Hmgcl,Gda,Slc4a4,Acsl5,Dlgap2,Pfkfb3,Acaa2,Cs,Elovl5,Pfas,Galk1,Zbtb20,Gpat4,Eno4,Far1,Pank1,Nudt9,Eif6,Adcy1,Pde8b,Pmvk,Entpd5,Elovl4,Gucy1a1,Xdh,Mpc2,Ubp1,Rfk,Park7,Dnph1,Nmnat2,Naxe,Naxd,Cox6a1,Cox4i1,Cox7a2,Atp6v0a1,Cox6c,Rnase4,Clcn3,Cox17,Atp6v0e1,Atp6v0a2,Cox8a,Cox5a,Slc4a10,Atp6v0a4,Atp6v0b,Coa8,Slc9a8,Ndufs7,Cox7c,Plaat3,Htr2c,P2ry1,Pdgfa,Pik3r1,Scp2,Ip6k1,Itpkb,Fdps,Pik3cb,Pip4k2b,Pi4k2a,Pgap2,Fabp5,Pik3ca,Dhrs7b,Pigp,Pik3c2b,Dpagt1,Atm,Abhd8,Ppip5k1,Etnk1,Lpin1,Pip5k1c,Pgap1,Sgms1,Bscl2,Selenoi,Piga,Plek,Dgke,Ppip5k2,Gamt,Atp5if1 |
| GO:0009144 | purine nucleoside triphosphate metabolic process | -6,50595 | -4,235 | 27/93 | Ak3,Atp5mc1,Tmsb4x,Ppargc1a,Nme2,Ran,Nme3,Atp5mc3,Atp5me,Oga,Opa1,Nme1,Atp5po,Atp5f1e,Atp5f1d,Antkmt,Ndufc2,Nudt2,Gtpbp1,Atp5mg,Trem2,Guk1,Dmac2l,Nudt16,Atp5pd,Uqcc3,Taf3 |
| GO:0009205 | purine ribonucleoside triphosphate metabolic process | -5,71837 | -3,539 | 24/84 | Ak3,Atp5mc1,Tmsb4x,Ppargc1a,Nme2,Ran,Nme3,Atp5mc3,Atp5me,Opa1,Nme1,Atp5po,Atp5f1e,Atp5f1d,Antkmt,Ndufc2,Nudt2,Gtpbp1,Atp5mg,Trem2,Dmac2l,Atp5pd,Uqcc3,Taf3 |
| GO:0009206 | purine ribonucleoside triphosphate biosynthetic process | -5,56518 | -3,410 | 21/69 | Ak3,Atp5mc1,Tmsb4x,Ppargc1a,Nme2,Nme3,Atp5mc3,Atp5me,Nme1,Atp5po,Atp5f1e,Atp5f1d,Antkmt,Ndufc2,Nudt2,Atp5mg,Trem2,Dmac2l,Atp5pd,Uqcc3,Taf3 |
| GO:0009141 | nucleoside triphosphate metabolic process | -5,46168 | -3,332 | 29/116 | Ak3,Tyms,Atp5mc1,Tmsb4x,Ppargc1a,Nme2,Ran,Nme3,Atp5mc3,Atp5me,Oga,Opa1,Nme1,Atp5po,Atp5f1e,Atp5f1d,Antkmt,Ndufc2,Nudt2,Gtpbp1,Atp5mg,Trem2,Dtymk,Guk1,Dmac2l,Nudt16,Atp5pd,Uqcc3,Taf3 |
| GO:0009145 | purine nucleoside triphosphate biosynthetic process | -5,45359 | -3,327 | 21/70 | Ak3,Atp5mc1,Tmsb4x,Ppargc1a,Nme2,Nme3,Atp5mc3,Atp5me,Nme1,Atp5po,Atp5f1e,Atp5f1d,Antkmt,Ndufc2,Nudt2,Atp5mg,Trem2,Dmac2l,Atp5pd,Uqcc3,Taf3 |
| GO:0009142 | nucleoside triphosphate biosynthetic process | -5,07012 | -3,003 | 23/85 | Ak3,Tyms,Atp5mc1,Tmsb4x,Ppargc1a,Nme2,Nme3,Atp5mc3,Atp5me,Nme1,Atp5po,Atp5f1e,Atp5f1d,Antkmt,Ndufc2,Nudt2,Atp5mg,Trem2,Dtymk,Dmac2l,Atp5pd,Uqcc3,Taf3 |
| GO:0009199 | ribonucleoside triphosphate metabolic process | -5,05404 | -2,992 | 24/91 | Ak3,Atp5mc1,Tmsb4x,Ppargc1a,Nme2,Ran,Nme3,Atp5mc3,Atp5me,Opa1,Nme1,Atp5po,Atp5f1e,Atp5f1d,Antkmt,Ndufc2,Nudt2,Gtpbp1,Atp5mg,Trem2,Dmac2l,Atp5pd,Uqcc3,Taf3 |
| GO:0009201 | ribonucleoside triphosphate biosynthetic process | -4,93296 | -2,894 | 21/75 | Ak3,Atp5mc1,Tmsb4x,Ppargc1a,Nme2,Nme3,Atp5mc3,Atp5me,Nme1,Atp5po,Atp5f1e,Atp5f1d,Antkmt,Ndufc2,Nudt2,Atp5mg,Trem2,Dmac2l,Atp5pd,Uqcc3,Taf3 |
| GO:0009150 | purine ribonucleotide metabolic process | -4,91015 | -2,876 | 62/353 | Pde4b,Dbi,Acadsb,Pde4a,Ak3,Psen1,Htr2a,Gpam,Atp5mc1,Cacnb4,Acaca,Pde10a,Hmgcl,Tmsb4x,Ppargc1a,Gda,Nme2,Slc4a4,Ran,Nme3,Acsl5,Atp5mc3,Dlgap2,Pfkfb3,Atp5me,Acaa2,Cs,Opa1,Elovl5,Nme1,Atp5po,Atp5f1e,Atp5f1d,Antkmt,Pfas,Galk1,Zbtb20,Gpat4,Eno4,Ndufc2,Far1,Pank1,Nudt2,Gtpbp1,Atp5mg,Trem2,Guk1,Nudt9,Eif6,Adcy1,Pde8b,Pmvk,Entpd5,Elovl4,Dmac2l,Nudt16,Gucy1a1,Xdh,Atp5pd,Uqcc3,Mpc2,Taf3 |
| GO:0006754 | ATP biosynthetic process | -4,87465 | -2,846 | 18/59 | Ak3,Atp5mc1,Tmsb4x,Ppargc1a,Atp5mc3,Atp5me,Atp5po,Atp5f1e,Atp5f1d,Antkmt,Ndufc2,Nudt2,Atp5mg,Trem2,Dmac2l,Atp5pd,Uqcc3,Taf3 |
| GO:0009259 | ribonucleotide metabolic process | -4,75467 | -2,738 | 64/372 | Pde4b,Dbi,Acadsb,Pde4a,Ak3,Psen1,Htr2a,Gpam,Atp5mc1,Cacnb4,Acaca,Pde10a,Hmgcl,Tmsb4x,Ppargc1a,Gda,Nme2,Slc4a4,Ran,Nme3,Acsl5,Atp5mc3,Dlgap2,Pfkfb3,Atp5me,Acaa2,Cs,Opa1,Elovl5,Nme1,Atp5po,Atp5f1e,Atp5f1d,Antkmt,Pfas,Galk1,Zbtb20,Gpat4,Eno4,Ndufc2,Far1,Pank1,Nudt2,Gtpbp1,Atp5mg,Ubp1,Trem2,Guk1,Nudt9,Eif6,Adcy1,Pde8b,Pmvk,Entpd5,Elovl4,Dmac2l,Nudt16,Gucy1a1,Xdh,Rfk,Atp5pd,Uqcc3,Mpc2,Taf3 |
| GO:0019693 | ribose phosphate metabolic process | -4,39071 | -2,456 | 64/382 | Pde4b,Dbi,Acadsb,Pde4a,Ak3,Psen1,Htr2a,Gpam,Atp5mc1,Cacnb4,Acaca,Pde10a,Hmgcl,Tmsb4x,Ppargc1a,Gda,Nme2,Slc4a4,Ran,Nme3,Acsl5,Atp5mc3,Dlgap2,Pfkfb3,Atp5me,Acaa2,Cs,Opa1,Elovl5,Nme1,Atp5po,Atp5f1e,Atp5f1d,Antkmt,Pfas,Galk1,Zbtb20,Gpat4,Eno4,Ndufc2,Far1,Pank1,Nudt2,Gtpbp1,Atp5mg,Ubp1,Trem2,Guk1,Nudt9,Eif6,Adcy1,Pde8b,Pmvk,Entpd5,Elovl4,Dmac2l,Nudt16,Gucy1a1,Xdh,Rfk,Atp5pd,Uqcc3,Mpc2,Taf3 |
| GO:0015985 | energy coupled proton transport, down electrochemical gradient | -4,23253 | -2,311 | 10/24 | Atp5mc1,Atp5mc3,Atp5me,Atp5po,Atp5f1e,Atp5f1d,Antkmt,Atp5mg,Atp5pd,Taf3 |
| GO:0015986 | ATP synthesis coupled proton transport | -4,23253 | -2,311 | 10/24 | Atp5mc1,Atp5mc3,Atp5me,Atp5po,Atp5f1e,Atp5f1d,Antkmt,Atp5mg,Atp5pd,Taf3 |
| GO:0006163 | purine nucleotide metabolic process | -4,22837 | -2,309 | 63/379 | Pde4b,Dbi,Acadsb,Pde4a,Ak3,Psen1,Htr2a,Gpam,Atp5mc1,Cacnb4,Acaca,Pde10a,Hmgcl,Tmsb4x,Ppargc1a,Gda,Nme2,Slc4a4,Ran,Nme3,Acsl5,Atp5mc3,Dlgap2,Pfkfb3,Atp5me,Oga,Acaa2,Cs,Opa1,Elovl5,Nme1,Atp5po,Atp5f1e,Atp5f1d,Antkmt,Pfas,Galk1,Zbtb20,Gpat4,Eno4,Ndufc2,Far1,Pank1,Nudt2,Gtpbp1,Atp5mg,Trem2,Guk1,Nudt9,Eif6,Adcy1,Pde8b,Pmvk,Entpd5,Elovl4,Dmac2l,Nudt16,Gucy1a1,Xdh,Atp5pd,Uqcc3,Mpc2,Taf3 |
| GO:0009117 | nucleotide metabolic process | -3,94292 | -2,080 | 72/458 | Pde4b,Dbi,Acadsb,Pde4a,Ak3,Psen1,Tyms,Htr2a,Gpam,Atp5mc1,Cacnb4,Acaca,Pde10a,Hmgcl,Tmsb4x,Ppargc1a,Gda,Nme2,Slc4a4,Ran,Nme3,Acsl5,Atp5mc3,Dlgap2,Pfkfb3,Park7,Atp5me,Oga,Acaa2,Cs,Dnph1,Opa1,Elovl5,Nme1,Atp5po,Atp5f1e,Atp5f1d,Antkmt,Pfas,Galk1,Zbtb20,Nmnat2,Gpat4,Eno4,Ndufc2,Far1,Pank1,Naxe,Nudt2,Gtpbp1,Atp5mg,Ubp1,Trem2,Dtymk,Guk1,Nudt9,Eif6,Adcy1,Pde8b,Pmvk,Entpd5,Elovl4,Naxd,Dmac2l,Nudt16,Gucy1a1,Xdh,Rfk,Atp5pd,Uqcc3,Mpc2,Taf3 |
| GO:1902600 | proton transmembrane transport | -3,92301 | -2,064 | 31/152 | Cox6a1,Cox4i1,Cox7a2,Atp5mc1,Atp6v0a1,Cox6c,Rnase4,Tmsb4x,Clcn3,Cox17,Atp6v0e1,Atp5mc3,Atp6v0a2,Park7,Atp5me,Cox8a,Atp5po,Atp5f1e,Atp5f1d,Cox5a,Antkmt,Slc4a10,Atp6v0a4,Atp6v0b,Coa8,Atp5mg,Slc9a8,Ndufs7,Atp5pd,Cox7c,Taf3 |
| GO:0090407 | organophosphate biosynthetic process | -3,88148 | -2,032 | 73/468 | Plaat3,Htr2c,P2ry1,Pdgfa,Pik3r1,Scp2,Ak3,Tyms,Htr2a,Gpam,Atp5mc1,Ip6k1,Itpkb,Acaca,Tmsb4x,Ppargc1a,Nme2,Fdps,Pik3cb,Nme3,Pip4k2b,Acsl5,Pi4k2a,Atp5mc3,Pgap2,Atp5me,Fabp5,Pik3ca,Elovl5,Nme1,Atp5po,Atp5f1e,Atp5f1d,Antkmt,Dhrs7b,Pfas,Pigp,Pik3c2b,Nmnat2,Gpat4,Ndufc2,Far1,Pank1,Nudt2,Dpagt1,Atp5mg,Atm,Trem2,Dtymk,Guk1,Adcy1,Abhd8,Pmvk,Ppip5k1,Etnk1,Lpin1,Pip5k1c,Elovl4,Pgap1,Sgms1,Bscl2,Selenoi,Dmac2l,Piga,Plek,Gucy1a1,Dgke,Rfk,Ppip5k2,Atp5pd,Uqcc3,Mpc2,Taf3 |
| GO:0006753 | nucleoside phosphate metabolic process | -3,70933 | -1,894 | 72/466 | Pde4b,Dbi,Acadsb,Pde4a,Ak3,Psen1,Tyms,Htr2a,Gpam,Atp5mc1,Cacnb4,Acaca,Pde10a,Hmgcl,Tmsb4x,Ppargc1a,Gda,Nme2,Slc4a4,Ran,Nme3,Acsl5,Atp5mc3,Dlgap2,Pfkfb3,Park7,Atp5me,Oga,Acaa2,Cs,Dnph1,Opa1,Elovl5,Nme1,Atp5po,Atp5f1e,Atp5f1d,Antkmt,Pfas,Galk1,Zbtb20,Nmnat2,Gpat4,Eno4,Ndufc2,Far1,Pank1,Naxe,Nudt2,Gtpbp1,Atp5mg,Ubp1,Trem2,Dtymk,Guk1,Nudt9,Eif6,Adcy1,Pde8b,Pmvk,Entpd5,Elovl4,Naxd,Dmac2l,Nudt16,Gucy1a1,Xdh,Rfk,Atp5pd,Uqcc3,Mpc2,Taf3 |
| R-RNO-163210 | Formation of ATP by chemiosmotic coupling | -3,70248 | -1,894 | 7/14 | Atp5mc1,Atp5me,Atp5po,Atp5f1d,Atp5mg,Dmac2l,Atp5pd |
| R-RNO-8949613 | Cristae formation | -3,70248 | -1,894 | 7/14 | Atp5mc1,Atp5me,Atp5po,Atp5f1d,Atp5mg,Dmac2l,Atp5pd |
| GO:0072521 | purine-containing compound metabolic process | -3,60701 | -1,818 | 64/406 | Pde4b,Dbi,Gamt,Acadsb,Pde4a,Ak3,Psen1,Htr2a,Gpam,Atp5mc1,Cacnb4,Acaca,Pde10a,Hmgcl,Tmsb4x,Ppargc1a,Gda,Nme2,Slc4a4,Ran,Nme3,Acsl5,Atp5mc3,Dlgap2,Pfkfb3,Atp5me,Oga,Acaa2,Cs,Opa1,Elovl5,Nme1,Atp5po,Atp5f1e,Atp5f1d,Antkmt,Pfas,Galk1,Zbtb20,Gpat4,Eno4,Ndufc2,Far1,Pank1,Nudt2,Gtpbp1,Atp5mg,Trem2,Guk1,Nudt9,Eif6,Adcy1,Pde8b,Pmvk,Entpd5,Elovl4,Dmac2l,Nudt16,Gucy1a1,Xdh,Atp5pd,Uqcc3,Mpc2,Taf3 |
| GO:0009152 | purine ribonucleotide biosynthetic process | -3,60518 | -1,817 | 31/158 | Ak3,Atp5mc1,Acaca,Tmsb4x,Ppargc1a,Nme2,Nme3,Acsl5,Atp5mc3,Atp5me,Elovl5,Nme1,Atp5po,Atp5f1e,Atp5f1d,Antkmt,Pfas,Ndufc2,Pank1,Nudt2,Atp5mg,Trem2,Guk1,Adcy1,Elovl4,Dmac2l,Gucy1a1,Atp5pd,Uqcc3,Mpc2,Taf3 |
| CORUM:372 | F0F1 ATP synthase, mitochondrial | -3,46928 | -1,712 | 7/15 | Atp5if1,Atp5mc1,Atp5me,Atp5f1e,Atp5f1d,Atp5mg,Atp5pd |
| GO:0009260 | ribonucleotide biosynthetic process | -3,31829 | -1,595 | 32/171 | Ak3,Atp5mc1,Acaca,Tmsb4x,Ppargc1a,Nme2,Nme3,Acsl5,Atp5mc3,Atp5me,Elovl5,Nme1,Atp5po,Atp5f1e,Atp5f1d,Antkmt,Pfas,Ndufc2,Pank1,Nudt2,Atp5mg,Trem2,Guk1,Adcy1,Elovl4,Dmac2l,Gucy1a1,Rfk,Atp5pd,Uqcc3,Mpc2,Taf3 |
| CORUM:158 | ATP synthasome | -3,06836 | -1,398 | 7/17 | Atp5mc1,Atp5mc3,Atp5me,Atp5po,Atp5f1e,Atp5f1d,Atp5pd |
| GO:0046390 | ribose phosphate biosynthetic process | -3,00892 | -1,356 | 32/178 | Ak3,Atp5mc1,Acaca,Tmsb4x,Ppargc1a,Nme2,Nme3,Acsl5,Atp5mc3,Atp5me,Elovl5,Nme1,Atp5po,Atp5f1e,Atp5f1d,Antkmt,Pfas,Ndufc2,Pank1,Nudt2,Atp5mg,Trem2,Guk1,Adcy1,Elovl4,Dmac2l,Gucy1a1,Rfk,Atp5pd,Uqcc3,Mpc2,Taf3 |
| GO:0042776 | mitochondrial ATP synthesis coupled proton transport | -2,89407 | -1,272 | 7/18 | Atp5me,Atp5po,Atp5f1e,Atp5f1d,Antkmt,Atp5mg,Atp5pd |
| GO:0006164 | purine nucleotide biosynthetic process | -2,70604 | -1,138 | 31/178 | Ak3,Atp5mc1,Acaca,Tmsb4x,Ppargc1a,Nme2,Nme3,Acsl5,Atp5mc3,Atp5me,Elovl5,Nme1,Atp5po,Atp5f1e,Atp5f1d,Antkmt,Pfas,Ndufc2,Pank1,Nudt2,Atp5mg,Trem2,Guk1,Adcy1,Elovl4,Dmac2l,Gucy1a1,Atp5pd,Uqcc3,Mpc2,Taf3 |
| GO:0072522 | purine-containing compound biosynthetic process | -2,44147 | -0,940 | 31/185 | Ak3,Atp5mc1,Acaca,Tmsb4x,Ppargc1a,Nme2,Nme3,Acsl5,Atp5mc3,Atp5me,Elovl5,Nme1,Atp5po,Atp5f1e,Atp5f1d,Antkmt,Pfas,Ndufc2,Pank1,Nudt2,Atp5mg,Trem2,Guk1,Adcy1,Elovl4,Dmac2l,Gucy1a1,Atp5pd,Uqcc3,Mpc2,Taf3 |
| R-RNO-1592230 | Mitochondrial biogenesis | -2,09354 | -0,704 | 7/24 | Atp5mc1,Atp5me,Atp5po,Atp5f1d,Atp5mg,Dmac2l,Atp5pd |

**Table S7: Metascape Enrichment metrics and DEG composition of each significant gene ontology in the SAL-NS vs SAL-S comparison in the DLS**

| Term | Description | LogP | Log(q-value) | InTerm_InList | Symbols |
| --- | --- | --- | --- | --- | --- |
| GO:0035082 | axoneme assembly | -7,20911 | -2,931 | 4/76 | Dnah1,Odad1,Ak7,Rsph1,Cfap126,Aurkb |
| GO:0035082 | axoneme assembly | -7,20911 | -2,931 | 4/76 | Dnah1,Odad1,Ak7,Rsph1 |
| GO:0001578 | microtubule bundle formation | -6,57586 | -2,598 | 4/109 | Dnah1,Odad1,Ak7,Rsph1 |
| GO:0044782 | cilium organization | -5,99666 | -2,195 | 5/383 | Dnah1,Odad1,Ak7,Rsph1,Cfap126 |
| GO:0000226 | microtubule cytoskeleton organization | -5,0261 | -1,350 | 5/604 | Aurkb,Dnah1,Odad1,Ak7,Rsph1 |
| GO:0060271 | cilium assembly | -4,53674 | -0,957 | 4/355 | Dnah1,Odad1,Ak7,Rsph1 |
| GO:0003341 | cilium movement | -3,89435 | -0,394 | 3/187 | Dnah1,Odad1,Ak7 |
| GO:0120031 | plasma membrane bounded cell projection assembly | -3,74566 | -0,329 | 4/567 | Dnah1,Odad1,Ak7,Rsph1 |
| GO:0030031 | cell projection assembly | -3,70484 | -0,329 | 4/581 | Dnah1,Odad1,Ak7,Rsph1 |
| GO:0007018 | microtubule-based movement | -2,9623 | 0,000 | 3/389 | Dnah1,Odad1,Ak7 |
| GO:0007283 | spermatogenesis | -2,44021 | 0,000 | 3/592 | Dnah1,Ak7,Rsph1 |
| GO:0048232 | male gamete generation | -2,39743 | 0,000 | 3/613 | Dnah1,Ak7,Rsph1 |
| GO:0007276 | gamete generation | -2,10471 | 0,000 | 3/780 | Dnah1,Ak7,Rsph1 |

**Table S8: Metascape Enrichment metrics and DEG composition of each significant gene ontology in the LPS-NS vs LPS-S comparison in the DLS**
